# A phylogenomic approach, combined with morphological characters gleaned via machine learning, uncovers the hybrid origin and biogeographic diversification of the plum genus

**DOI:** 10.1101/2023.09.13.557598

**Authors:** Richard G. J. Hodel, Sundre K. Winslow, Si-Yu Xie, Bin-Bin Liu, Liang Zhao, Gabriel Johnson, Michael Trizna, Alex E. White, Rebecca B. Dikow, Daniel Potter, Elizabeth A. Zimmer, Jun Wen

**Affiliations:** Department of Biological Sciences, Northern Arizona University, 617 S. Beaver St., Flagstaff AZ 86011, USA; Department of Botany, National Museum of Natural History, MRC 166, Smithsonian Institution, Washington, DC, 20013-7012, USA; Data Science Lab, Office of the Chief Information Officer, Smithsonian Institution, Washington, DC, 20560, USA; College of Life Sciences, Northwest A&F University, Yangling 712100, China; State Key Laboratory of Systematic and Evolutionary Botany, Institute of Botany, Chinese Academy of Sciences, Beijing 100093, China; Department of Plant Sciences, University of California, Davis, California, 95616, USA

**Keywords:** allopolyploidy, herbarium specimens, hybridization, machine learning, niche conservatism, tropical-to-temperate biogeographic transitions

## Abstract

The evolutionary histories of species have been shaped by genomic, environmental, and morphological variation. Understanding the interactions among these sources of variation is critical to infer accurately the biogeographic history of lineages. Here, using the geographically widely distributed plum genus (*Prunus*, Rosaceae) as a model, we investigate how changes in genomic and environmental variation drove the diversification of this group, and we quantify the morphological features that facilitated or resulted from diversification. We sequenced 587 nuclear loci and complete chloroplast genomes from 99 species representing all major lineages in *Prunus*, with a special focus on the understudied tropical racemose group. The environmental variation in extant species was quantified by synthesizing bioclimatic variables into principal components of environmental variation using thousands of georeferenced herbarium specimens. We used machine learning algorithms to classify and measure morphological variation present in thousands of digitized herbarium sheet images. Our phylogenomic and biogeographic analyses revealed that ancient hybridization and/or allopolyploidy spurred the initial rapid diversification of the genus in the early Eocene, with subsequent diversification in the north temperate zone, neotropics, and paleotropics. This diversification involved successful transitions between tropical and temperate biomes, an exceedingly rare event in woody plant lineages, accompanied by morphological changes in leaf and reproductive morphology. The machine learning approach detected morphological variation associated with ancient hybridization and quantified the breadth of morphospace occupied by major lineages within the genus. The paleotropical lineages of *Prunus* have diversified steadily since the late Eocene/early Oligocene, while the neotropical lineages diversified much later. Critically, both the tropical and temperate lineages have continued to diversify. We conclude that the genomic rearrangements created by reticulation deep in the phylogeny of *Prunus* may explain why this group has been more successful than other groups with tropical origins that currently persist only in either tropical or temperate regions, but not both.

## INTRODUCTION

Bursts of speciation associated with changing environmental conditions have occurred throughout the Tree of Life. Environmental factors, such as changes in climate, tectonic shifts, mountain uplift, as well as biotic interactions, can all drive the diversification of lineages. In particular, speciation may occur when lineages diversify to occupy newly available niches resulting from changing environmental conditions. Classic evolutionary studies such as those focused on Galapagos finches found that new ecological opportunities drove the rapid evolution of new lineages (Schluter 2000). Climatic fluctuations have been implicated in promoting both speciation and extinction in a variety of fauna (Weir and Schulter 2007). Changing environmental conditions can have a particularly strong impact on plant lineages. For example, increased global aridity likely promoted the diversification of both C4 grasses and succulent lineages (Arakaki et al. 2011). Synthesis of current evidence suggests that many plant lineages diversified in the tropics, sometimes spreading into temperate biomes (Spriggs et al. 2015), and that many temperate lineages had a Boreotropical origin (Zhang et al. 2021, Nie et al. 2023). Despite some evidence of shared biogeographic patterns, however, surprisingly few lineages have successfully made transitions from tropical to temperate regions (Kerkhoff et al. 2014). In fact, in the neotropics, the descendants of tropical ancestral lineages remained tropical in 94% of woody angiosperms (Kerkhoff et al. 2014). Similarly, temperate lineages were quite conserved, with 90% of temperate descendants arising from temperate ancestors (Kerkhoff et al. 2014). These patterns of diversification are often attributed to the tropical conservatism hypothesis (TCH) (Wiens & Donoghue 2004), which postulates that many lineages of tropical origin could not adapt readily to the cooler, drier, and more seasonal temperate regions, and either were extirpated (i.e., migrated to track tropical environments), or went extinct (Donoghue 2008).

In addition to environmental conditions, genomic changes can spark the diversification of lineages. In some cases, reshuffling of genetic material may facilitate rapid diversification in response to environmental change (Schenk 2021). A variety of genomic mechanisms, including hybridization, gene duplication or loss, and/or genome doubling, can rearrange genomic material such that new traits and adaptations develop to allow plant lineages to spread and diversify into new niches and habitats (Xu et al. 2017, Hodel et al. 2022). Genome doubling may provide extensive genetic variation and novelty upon which selection may act, driving adaptive diversification (Seehausen 2004, Soltis and Soltis 2016, Doyle and Coate 2019, Griffiths et al. 2019, reviewed in Schenk 2021). Moreover, genomic changes in response to novel environments may be coupled with morphological innovation (García-Verdugo et al. 2013). When lineages encounter new environments, they may already possess phenotypes that facilitate their occupancy of available niches in the community they enter, or the available niches in newly encountered ecological conditions may drive diversifying selection on traits that are shaped to promote survival in the novel niche space (Wellborn & Langerhans 2014). In some lineages, success of diversification events in response to new ecological opportunities is mediated by biological features as well as idiosyncrasies of the environmental conditions at the time (Wellborn & Langerhans 2014). In summary, both evolutionary events and environmental circumstances interact to shape the speciation and extinction rates that characterize diversification (Spriggs et al. 2015).

One lineage that has been able to readily diversify in response to changing environments was the genus *Prunus* (the plum genus, Rosaceae). This group contains approximately 250-400 evergreen and deciduous species that occur throughout the temperate regions of the northern hemisphere and in the tropics and subtropics of both the Old and New Worlds (Rehder 1940, Wen et al. 2008, Perez Zabala 2022). Species in this genus are key elements of both temperate and tropical broadleaf forests. Despite its ecological and economic significance—*Prunus* contains important crop species such as cherries, peaches, almonds, apricots, and plums—the phylogenetic relationships of the major lineages in the genus are still unresolved. Historically, the major groups within the genus were defined by inflorescence morphology—with three major groups identified—solitary flower (e.g., peach), corymbose (i.e., producing a flat-topped indeterminate cluster of flowers; e.g., cherries), and racemose (i.e., producing an indeterminate inflorescence with the main axis not terminating in a flower; e.g., bird-cherries) (Rehder 1940; Su et al. 2023). Multiple genetic studies using chloroplast DNA markers have inferred that the solitary and corymbose groups form a clade that is sister to the racemose group (Bortiri et al. 2001, Wen et al. 2008, Chin et al. 2014, Zhao et al. 2016). However, nuclear markers have been unable to resolve the backbone phylogenetic relationships of the genus (Lee & Wen 2001, Bortiri et al. 2001, Wen et al. 2008, Chin et al. 2014, Zhao et al. 2016; Fig. 1). Furthermore, studies using nuclear markers either relied on few nuclear loci (Wen et al. 2008, Chin et al. 2014, Zhao et al. 2016) or had poor sampling in the understudied racemose group (Hodel et al. 2021).

**Figure 1.**
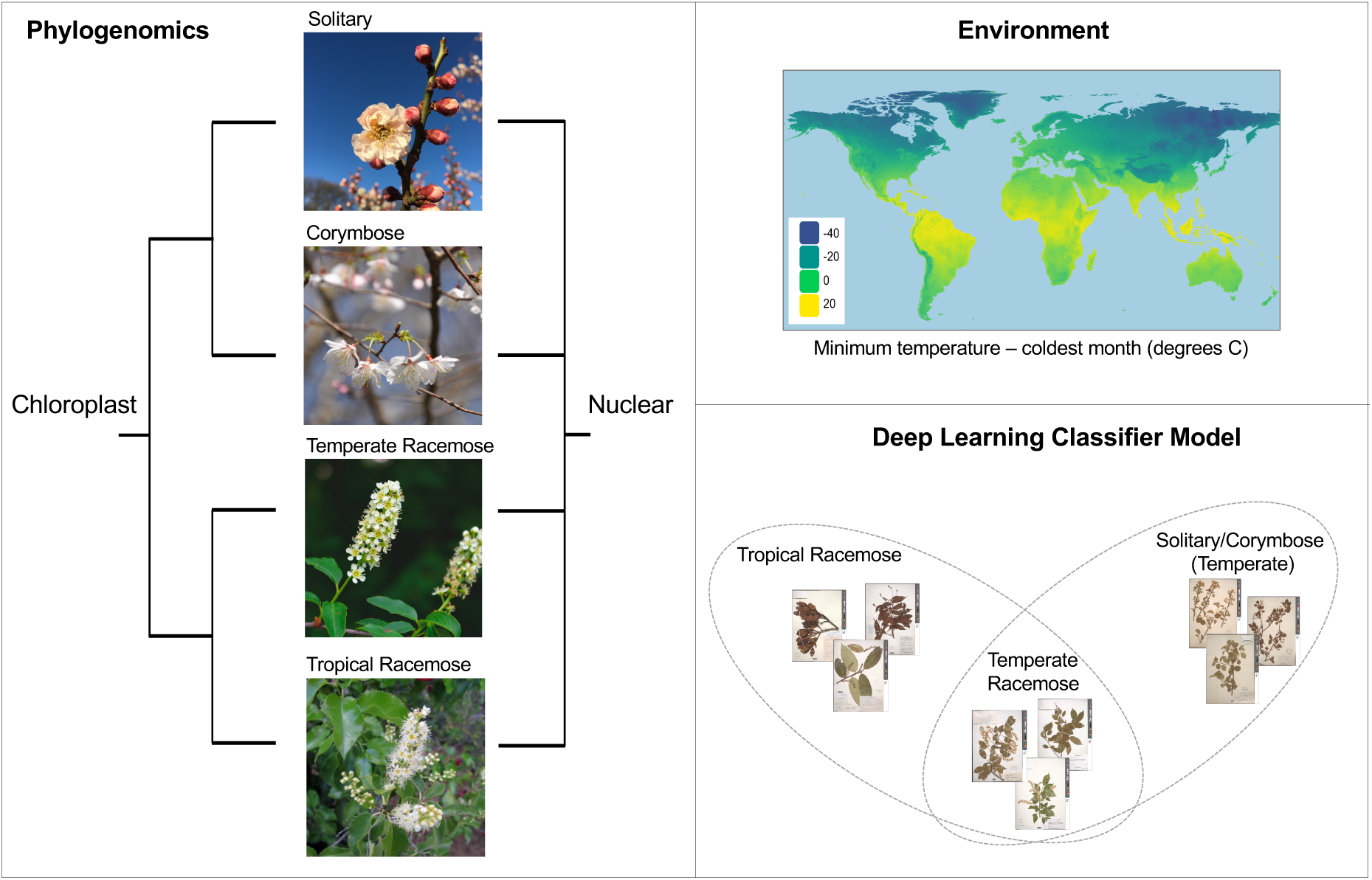
Synthesizing genomic, environmental, and phenomic data can improve our understanding of the evolutionary processes that drive diversification. The summary of our phylogenetic understanding of the major groups in the plum genus (*Prunus*) based on chloroplast and nuclear markers (left). One environmental variable, minimum temperature of the coldest month, often determines the distributional limits of tropical taxa (upper right). Machine learning approaches, when applied to digitized museum specimens, can be used to infer how characters associated with reproduction and/or environmental adaptation correspond to phylogeny (lower right).

Attempts to characterize the backbone of the *Prunus* phylogeny have been obscured by cytonuclear discord (Chin et al. 2014). One hypothesis that has been invoked to explain this cytonuclear discord is that the genus diversified via an ancient allopolyploidy and/or hybridization event (Zhao et al. 2016). Based on chromosome count data, the corymbose and solitary groups are diploid, whereas nearly all racemose lineages are polyploid. In summary, the evolutionary history of the plum genus remains unclear. To address phylogenomic uncertainty, herein we assemble a phylogenomic dataset with hundreds of nuclear loci and entire chloroplast genomes for each of 99 *Prunus* species representing all major lineages in the genus. Our taxon sampling is the most complete to date, especially in the understudied racemose group. We use the phylogeny of *Prunus* as a framework to investigate reticulation events, to reconstruct the biogeographic history, and test hypotheses regarding the morphological basis of key biogeographic transitions and reticulation. The morphological characters associated with ancient biogeographic transitions and reticulation are notoriously difficult to measure in extant species (e.g., McVay et al. 2017). Accordingly, we developed a novel approach to quantify and categorize morphological variation: leveraging machine learning approaches with digitized herbarium sheet image data (Fig. 1). Our specific objectives in this study are to:

1. Resolve the phylogenetic relationships of major groups within the genus.
2. Assess the role of ancient genomic rearrangements—specifically allopolyploidy and/or hybridization—in shaping the evolutionary history of *Prunus*.
3. Clarify the biogeographic history of *Prunus*, especially concerning the timing and frequency of transitions between tropical and temperate regions.
4. Implement a machine learning approach with thousands of digitized herbarium specimen images to test for morphological evidence associated with biogeographic transitions and reticulation events such as allopolyploidy and hybridization.

## MATERIALS AND METHODS

### Genomic data collection

#### Hyb-Seq probe design

We designed a 610-locus custom Hyb-Seq probe set to target nuclear genes in *Prunus*. Our approach aimed to obtain genes with the highest probability of being strictly single-copy (i.e., single-copy nuclear genes; hereafter SCNs). We used an iterative process to BLAST publicly available genomes against themselves to obtain a candidate pool of putatively SCN loci (Supplemental Fig. S1); the subsequent analyses were conducted in Geneious Prime 2020.0.5 (https://www.geneious.com). We first conducted BLAST searches of the annotated genomes of *P. persica* (L.) Batsch (peach, PRJNA31227; Verde et al. 2013), *P. avium* (L.) L. (sweet cherry, PRJDB4877; Shirasawa et al. 2017), and *Malus domestica* (Suckow) Borkh. (PRJNA339703, Daccord et al. 2017) against themselves using an e-value of 1e-10, which yielded 11,305, 10,703, and 968 candidate SCNs, respectively. We used *Malus domestica* (apple), which is in the same subfamily as *Prunus* – the Amygdaloideae – to expand the phylogenetic breadth of the baits. We then BLASTed the candidate SCNs from *P. avium* against the SCNs from *P. persica*, and filtered out loci that had multiple hits, were fewer than 300 nucleotides, and had less than 95.4% pairwise identity. Loci with multiple hits were presumably not strictly single-copy, loci with fewer than than 300 nucleotides have less phylogenetic information than longer loci, and loci with pairwise identity close to 100% may not be variable enough to be informative (Weitemier et al. 2014). This left 318 loci with a total of 262,684 nucleotides of sequence, and we retained the sequences from *P. avium* for bait design. To obtain additional loci more closely related to racemose *Prunus* species, we also BLASTed the candidate loci that passed the above length and pairwise identity filter, and had two or fewer hits, against the *P. serotina* Ehrh. (black cherry) transcriptome (Swenson et al. 2017; available via the Hardwood Genomics Project, hardwoodgenomics.org). This yielded 96 additional SCNs with 72,172 nucleotides of sequence, and the *P. serotina* sequence was retained for bait design. We also BLASTed the candidate SCNs from *P. avium* against the SCNs from *Malus domestica*; after filtering out loci with multiple hits, fewer than 600 nucleotides, and less than 80% or greater than 87.5% pairwise identity, we retained 160 loci totaling 213,990 nucleotides. Here, we used sequences from *P. avium* for bait design. Finally, because inflorescence architecture has historically been used to define phylogroups in *Prunus*, we selected 36 functional genes that may be associated with flowering (e.g., APETALA, FT; Yao et al. 2022). This gene set included 36 loci totaling 137,252 nucleotides, and we used *P. avium* sequences for bait design. In total, we selected 610 loci with a total of 686,098 nucleotides for our custom *Prunus* bait set (Arbor Biosciences, Ann Arbor, MI). In the Rosaceae, a custom bait set may perform better than a universal set (Ufimov et al. 2021).

#### Field collection, DNA extraction, library preparation, and hybridization reactions

Specimens were collected from the field (Supplemental Table S1) and stored in silica gel until DNA was extracted from leaf tissue using a modified CTAB protocol (Doyle & Doyle 1987). Next, DNA libraries were constructed using a KAPA HyperPrep kit (available from Roche, Basel, Switzerland) following the manufacturer’s protocol except with quarter-volume reactions for each individual. Briefly, DNA extractions were fragmented via sonication using the QSonica ultrasonicator (Newtown, Connecticut, USA), uneven ends were repaired and A-tailed, and Illumina adaptors were ligated to the DNA fragments. Next, AMPure magnetic beads were used to purify and size-select the adaptor ligated DNA, and the DNA libraries were PCR amplified to add unique i5 and i7 indexed oligonucleotides (i.e., barcodes). Libraries were pooled in groups of eight with the criteria of balancing samples among presumed phylogenetic distance and concentration prior to the hybridization capture reaction with custom-designed baits. DNA libraries were hybridized with the baits for 48 hours at 60°C following the MyBaits v4 manual (Arbor Biosciences, Ann Arbor, Michigan, USA). The hybridization enriched libraries were combined with unenriched DNA libraries in a 60:40 ratio to enable generation of plastomes from off target reads. All enriched and unenriched libraries were combined into a single tube and sent for 2x150bp sequencing on the Illumina HiSeq 4000 at Novogene (Sacramento, California, USA). We also acquired sequence data for one species from NCBI GenBank (Supplemental Table S1). In total, we generated or obtained data for 119 accessions (Supplemental Table S1) representing 101 species, which include 99 *Prunus* species and two outgroups (*Lyonothamnus floribundus* A.Gray and *Physocarpus opulifolius* (L.) Maxim.). These outgroups were selected because *Lyonothamnus* is likely the sister lineage of *Prunus* (Xiang et al. 2017), and *Physocarpus* is more distantly related but within the Amygdaloideae (Xiang et al. 2017, Zhang et al. 2017).

#### Plastome assembly and phylogenetic inference

First, raw sequencing reads were quality filtered and Illumina adapters were removed using bbduk (https://sourceforge.net/projects/bbmap/). GetOrganelle (Jin et al. 2020) was used to generate plastomes for each species using the reads cleaned by bbduk. Of the 119 accessions included in the final dataset, 75 produced complete, circular plastomes after an initial run through GetOrganelle. For the remaining 44, we used minimap2 (Li 2018) implemented in Geneious Prime 2020.0.5 (https://www.geneious.com) to assemble the contigs and scaffolds from GetOrganelle, using the *Prunus avium* plastome (NCBI accession number MK622380) as a reference to generate as complete as possible plastomes. After using minimap2 on the non-circularized plastomes, we obtained nearly complete plastomes for all accessions. To ensure that the use of *Prunus avium* as a reference did not bias the plastome results, we constructed a phylogeny using the plastomes we generated, as well as all publicly available *Prunus* plastomes (N = 17), to check that each newly sequenced accession is placed reasonably in the phylogeny (Supplemental Table S2). The plastome phylogeny was inferred using RAxML with 100 rapid bootstrap replicates and 20 independent maximum likelihood searches (i.e., the following parameter settings: “-f a -m GTRGAMMA -p 12345 -x 12345 -# 100”). All analyses, unless otherwise stated, were run on the Smithsonian Institution High Performance Cluster (SI/HPC, “Hydra”) (https://doi.org/10.25572/SIHPC).

#### Hyb-Seq locus assembly

We used the software package HybPiper v2.16 (Johnson et al. 2016) to assemble nuclear loci. The same quality filtering and Illumina adapter removal strategy as with the plastome pipeline was used with the nuclear data (i.e., cleaning and trimming using bbduk). We followed the core scripts of the HybPiper pipeline; we first used hybpiper assemble –run_intronerate to map reads using BWA (Li & Durbin 2009), sorted reads into fasta files, and ran the SPADES assembler (Bankevich et al. 2012). Next, the command hybpiper stats was used to get the lengths of the recovered gene sequences, visualize the success of each gene for each species, and generate a file of descriptive statistics for the assembly (Supplemental Table S3, Supplemental Fig. S2). The command hybpiper retrieve_sequences was used to retrieve exons, introns, and supercontigs from each gene region for every species, and put them in an unaligned fasta file. Next, for each type of locus (i.e., exons, introns, and supercontigs), the unaligned gene files were aligned using MAFFT (Katoh & Standley 2013) with the following parameter settings: “--maxiterate 5000 --auto --adjustdirectionaccurately –leavegappyregion.” The resulting alignments were trimmed using the phyx command pxclsq with the proportion of sites required to have data (-p option) set at 0.5 (Brown et al, 2017). For each of the trimmed alignments, gene trees were estimated using RAxML (Stamatakis 2014) with 100 rapid bootstrap replicates and 20 independent maximum likelihood searches (i.e., the following parameter settings: “-f a -m GTRGAMMA -p 12345 -x 12345 -# 100”). After confirming that exons and supercontigs produced similar species tree topologies (Supplemental Fig. S3), we used supercontigs for all subsequent analyses.

#### Species tree estimation

The nuclear species tree was estimated using ASTRAL-III (Zhang et al. 2018), an approach consistent with the coalescent, to summarize the 587 gene trees. We used the ASTRAL quartet scores to assess nodewise support for inferred phylogenetic relationships. A concatenation approach implemented in RaXML, with 100 rapid bootstrap replicates and 20 independent maximum likelihood searches, was also used for comparative purposes. For some downstream analyses (e.g., divergence dating), it was necessary to have a phylogeny with meaningful branch lengths. For these analyses, we used the ASTRAL topology as a constraint tree for a RAxML tree search to obtain a species tree with the ASTRAL topology but with proportional branch lengths.

#### Gene tree discordance analysis

To assess congruence between gene trees and the species tree, we used the program phyparts (Smith et al. 2015). At each node in the tree, phyparts compares the rooted gene tree topologies with the rooted species tree to label the number of genes that are concordant, discordant, and uninformative relative to the species tree. Gene trees were rooted for the phyparts analysis and we considered any gene trees with less than 50% bootstrap support at a given node to be uninformative for that node (i.e., -s 50 option).

#### Paralogs

We used the tree-based paralog detection implemented in HypPiper 2.0+ to distinguish between true orthologs and paralogs. The HybPiper post-processing command hybpiper paralog_retriever was used to retrieve the multiple sequences for putative paralogous genes. Next, the unaligned retrieved gene sequences were inputted into a phylogenetic pipeline that included alignment with MAFFT (Katoh & Standley 2013) and phylogeny construction with FastTree (Price et al. 2009). We manually examined the resulting phylogeny for each gene. In some cases, genes suspected to be paralogs formed a clade, indicating that the putative paralogs may be alleles and not the result of duplication. Alternatively, genes that were truly paralogs would present as multiple copies dispersed throughout the phylogeny inferred by FastTree. For all 610 genes, we classified paralogy status based on the presence or absence of paralogs. If paralogs were detected, we manually inspected the paralog tree for such genes to determine if the suspected paralogs clustered, or were in fact true paralogs dispersed throughout the gene tree. In the cases where true paralogs were detected, we discarded that gene from the analysis. In the cases where the main contig and additional contigs clustered and formed a clade, we retained the primary copy outputted by HybPiper. We discarded 23 genes that had true paralogs, and retained 587 genes for downstream phylogenetic analyses.

#### Hybridization and allopolyploidy analyses

Because hypotheses of ancient hybridization and allopolyploidy have been proposed to explain the origin and diversification of this group, we used phylonet v3.8.2 (Than et al. 2008) to investigate potential reticulation. We assembled 20 haphazardly sampled datasets each consisting of seven taxa, representing the major lineages—one species each from the corymbose, solitary-flower, and temperate racemose clades, three from the tropical racemose clade, and an outgroup (i.e., *Physocarpus opulifolius*). We constructed rooted gene trees for each gene in each of the 20 datasets using RAxML (Stamatakis 2014) with 100 rapid bootstrap replicates and 20 independent maximum likelihood searches. We used the maximum pseudolikelihood approach (Yu & Nakhleh 2015) to infer networks with maximum reticulations set to 1, 2, and 3. We ran multiple replicates (n=10) for each dataset with each of the three maximum reticulation values. For each dataset, the networks with the optimal pseudolikelihood scores were retained and visualized using Dendroscope (Huson & Scornavacca 2012).

To investigate histories of allopolyploidy, and for comparison with reticulation results from phylonet, we used the program GRAMPA (Thomas et al. 2017). This approach uses a least common ancestor mapping algorithm to reconcile gene trees and species trees (Goodman et al. 1979; Page 1994) so that polyploidy events can be placed on a phylogeny. GRAMPA can identify modes of polyploidy as well as place whole genome duplications (WGDs) on a phylogeny, and infer parental participants in instances of polyploidy. The user can input clades of interest that are suspected to have a polyploid origin, or use a global search that considers if all clades could be the result of WGD. We used the former approach to target specific major clades that were hypothesized to have an ancient polyploid origin. Specifically, we tested the nodes defining the following clades: temperate racemose, solitary-flower, corymbose, all temperate (‘Temp’), tropical (‘Trop’), core tropical (‘CTrop’), Australasia paleotropical (‘APal’), South American neotropical (‘SNeo’), and North American neotropical (‘NNeo’) (see Fig. 2A for clade definitions). In the analysis, we allowed any other node in the tree to be a parental lineage to any clades determined to be of polyploid origin. For hypothesized instance of polyploidy, GRAMPA assesses if a standard singly-labeled species tree or a multi-labeled tree is the most parsimonious explanation of the data.

**Figure 2A.**
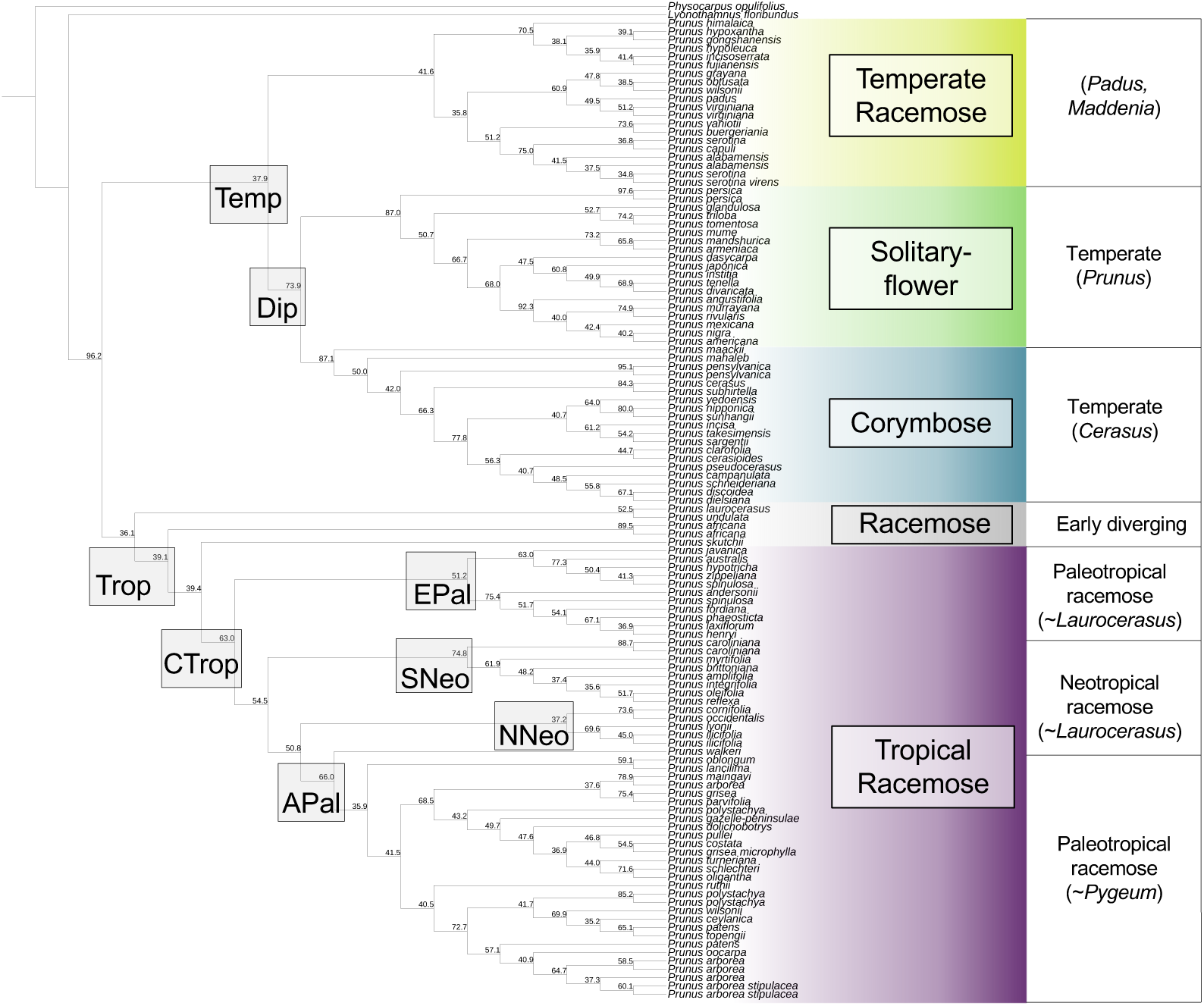
The phylogenetic species tree inferred using ASTRAL to summarize 587 nuclear gene trees. Important nodes in the phylogeny are highlighted and named: temperate (‘Temp’), diploid (‘Dip’), tropical (‘Trop’), core tropical (‘CTrop’), East Asian paleotropical (‘EPal’), Australasia paleotropical (‘APal’), South American neotropical (‘SNeo’), and North American neotropical (‘NNeo’). The numbers at nodes are ASTRAL quartet support scores.

To complement the GRAMPA analysis and investigate all nodes in the phylogeny, we used the approach of Yang et al. (2017) to map duplications to nodes in the species tree. Briefly, we mapped duplication events within orthogroups to the ASTRAL species tree. For a given subclade, if two or more taxa overlapped between daughter clades, a gene duplication event was counted at the node which was defined as the most recent common ancestor of the subclade on the ASTRAL species tree. We required the average bootstrap percentage for each orthogroup to be >50.

#### Environmental characterization

To quantify the environment for major groups, we used a Principal Components Analysis (PCA) to synthesize variation in the 19 WorldClim bioclimatic variables (https://www.worldclim.org/) at a 2.5-minute resolution. The values for each of the 19 variables were extracted for georeferenced herbarium specimens that represented each group using the R packages ‘dismo’ (Hijmans et al. 2017) and ‘raster’ (Hijmans 2016). Two approaches were used to quantify the environmental variation, first between two groups—temperate versus tropical—as well as a comparison of four groups: temperate diploid (solitary + corymbose), temperate racemose, neotropical racemose, and paleotropical racemose. We distilled the variation in 19 bioclimatic variables into 3 PC axes using the R package ‘vegan’ (Oksanen et al. 2017). For each of the four groups, we defined 95% confidence interval ellipses, which indicate the region with 95% probability that the centroid is contained within the ellipse. For this analysis, we only used the species for which we had genomic data, and a total of 10,011 herbarium specimen records were included. Although this approach only captures present environmental variation, synthesis has revealed that niches are more phylogenetically conserved than expected (Donoghue 2008), and accordingly surveying all extant species in a lineage can give a reasonable approximation for the environment in which a given lineage evolved. We also used the GPS data to collect elevation data of each specimen using the R package ‘rgbif’ (Chamberlain et al. 2023). This was done primarily to investigate whether tropical species occurred at lower or higher elevations, because some high elevation tropical regions may resemble temperate environments more closely than tropical ones, and this may impact our interpretation of biogeographic results.

#### Dating and biogeographic analysis

To investigate the timing and biogeographic history of *Prunus*, we used a biogeographic ancestral range estimation analysis implemented in BioGeoBears (Matzke 2012, 2013). First, we used treePL, a method for estimating divergence times on a phylogeny using penalized likelihood, to infer divergence dates for all nodes in the *Prunus* phylogeny. Three calibrations were used to date the phylogeny. We used *Prunus cathybrownae* (Benedict et al. 2011) from the Early Eocene of North America as the most recent common ancestor (MRCA) of the diploid (i.e., corymbose+solitary) lineages (stem lineage leading to node Dip; Fig. 2A), with a minimum age of 50 million years ago (Mya). The fossil *Prunus wutuensis* (Li et al. 2011) from the early Eocene of East Asia (Shandong, China) was used to fix the MRCA of crown *Prunus* to a minimum age of 58 Mya. Next, the crown of the Amygdaloideae was fixed at a minimum age of 90 Mya based on a calibrated Rosaceae phylogeny from Xiang et al. (2017). Similarly, we constrained the age of the stem Amygdaloideae to have a minimum age of 100 Mya using the calibrated Rosaceae phylogeny from Xiang et al. (2017) as a guide. Using BioGeoBears (Matzke 2013), we considered six possible biogeographic models (DEC, DEC + J, DIVALIKE, DIVALIKE + J, BAYAREALIKE, BAYAREALIKE + J) and selected the optimal model for the data using AIC and AICc comparisons. Likelihood ratio tests were also used to determine if the more parameter-rich +J models were preferred compared to the equivalent models without the +J term.

We defined seven biogeographic regions based on geography: North America, South America, Africa, Europe, West Asia, East Asia, and Australasia. In this analysis, the maximum areas was set to two. We also used a biome-based approaches to delimit biogeographic regions. Using the georeferenced herbarium data from the previous section (‘*Environmental analysis*’), we coded each species’ presence in each of the 14 biomes delineated in the Ecoregions-17 dataset (Dinerstein et al. 2017). We combined similar biomes such that the BioGeoBears analysis would be computationally feasible and the results readily interpretable. We defined four biome-based biogeographic regions: 1) Dry (Desert & Xeric Shrubland/Mediterranean Forests, Woodlands & Scrub); 2) Cold (Boreal Forests & Taiga/Montane Grasslands & Shrublands); 3) Temperate (Temperate Broadleaf & Mixed Forests/Temperate Conifer Forests/Temperate Grasslands, Savannas & Shrublands); 4) Tropical (Tropical & Subtropical Coniferous Forests/Tropical & Subtropical Dry Broadleaf Forests/Tropical & Subtropical Grasslands, Savannas & Shrublands/Tropical & Subtropical Moist Broadleaf Forests). In this analysis, we set the maximum areas to four. The frequency and type of biogeographic events were estimated using Biogeographic Stochastic Mapping (BSM) in BioGeoBears (Matzke 2013). For each of the sets of biogeographic regions (i.e., continent-based, biome-based), we selected the optimal model based on AIC and AICc comparisons and ran the BSM analysis on each biogeographic model, with 50 independent mappings. The BSM approach can differentiate between biogeographic events that occur at speciation nodes (i.e., cladogenesis events such as sympatric speciation, vicariance, or founder events) versus transitions occurring along branches (e.g., anagenetic dispersal).

#### Diversification analysis

To investigate the variation in diversification rates across the phylogeny, we implemented an episodic birth-death model (EBD; Höhna 2015) and a branch-specific diversification model (LSBDS; Höhna et al. 2019) in RevBayes (Höhna et al. 2016). The EBD model assumes speciation and extinction rates are constant within each time interval but are allowed to vary between time intervals. We used log-transformed rates following a Horseshoe Markov random field prior distribution; this approach assumes that rates are autocorrelated. The LSBDS model assumes rate heterogeneity across branches and does not require *a priori* assignment of shifts. In other words, this model allows for a birth-death process with diversification rates that can vary among branches. We used exponential priors for both diversification and extinction rates and used 50,000 generations of reversible-jump Markov chain Monte Carlo, with 25% burnin, to sample models with a variety of rate shift placements.

The dated species tree inferred from ASTRAL, with node ages calibrated using penalized likelihood in *treePL*, was used as input for the EBD and LSBDS models. We report the speciation rate, extinction rate, relative extinction rate, and net diversification rate. The software CRABS (Congruent Rate Analyses in Birth–death Scenarios; Höhna et al. 2022) was used to assess heterogeneity in diversification rates, and if rate patterns were robust to the non-identifiability of the birth**–**death model. Specifically, we assessed the estimated posterior median from the EBD model to compare speciation and extinction rate over time. The posterior median is considered a robust estimate for such rates (Magee et al. 2020).

### Morphological data collection

#### Herbarium image analysis

Because of the rich representation of *Prunus* species in museum collections, we used a deep learning approach with digitized herbarium specimen sheets to test the degree to which morphological classifications of species and/or lineages corresponded to phylogenetic hypotheses. We assembled a dataset of 4,228 images representing the 99 *Prunus* species with genomic data in this study. These images were downloaded using the ‘idig_search_media’ function in the ‘ridigbio’ R package (Michonneau et al. 2016). We used exiftool (https://exiftool.org/) to remove EXIF metadata from the images using the command “exiftool - overwrite_original -EXIF= *.jpg”). This approach removes metadata often incompatible with image processing libraries commonly used in machine learning. We also trimmed the edges of the digitized images before using them in downstream machine learning analyses. This was done to remove non-biological information, such as herbarium labels, color bars, and scale bars, from the digitized herbarium sheet. We removed the top 20% and bottom 20% of every image used in machine learning analyses. This unnecessarily removed some biologically relevant features, but was conservative for ensuring the removal of non-biologically meaningful information on the images. We investigated both trimmed and untrimmed sheets to test the impact of removing the bottom 20% and top 20% of each sheet; hereafter these sheets are referred to as ‘trimmed’ or ‘whole sheet’, respectively. Furthermore, because we had thousands of digitized herbarium sheets available, this approach represented a reasonable balance between removing too much data and ensuring that any data with the potential to bias results was excluded.

#### Classification Algorithms

To assess if morphological variation corresponded to genetic variation, we developed a supervised machine learning classifier algorithm using the fastai (https://github.com/fastai/fastai) deep learning library, which is built using PyTorch libraries (https://pytorch.org), to classify herbarium sheet images into categories. The labels for each category were assigned based on clades inferred using genomic data. We used two approaches for the classification analysis. First, we assigned labels of ‘diploid species,’ corresponding to all species with solitary and corymbose inflorescences, and ‘tropical racemose,’ which referred to all tropical species with racemes. These two groups were selected because the phylogenetic relationship between these two groups was the same in the nuclear and chloroplast phylogenies. This model was trained and validated using a total of 5,045 herbarium sheet images representing 81 species, with 2,746 in the solitary/corymbose group, and 2,301 in the tropical racemose group. In each group, 80% of the labeled input data were used for training, and 20% for validation. The model was run for 24 epochs until an accuracy rate of 97.2% was achieved. As an additional check that the model was performing well, we tested hundreds of specimens from species in either the solitary/corymbose groups (N = 452), or the tropical racemose groups (N = 426), which were not species used to train or validate the model. Next, the additional ‘test’ data in this implementation were species from the temperate racemose group (N = 676). This group was selected as test data because its phylogenetic position was different between the nuclear and chloroplast topologies. Essentially, this machine learning classification approach was used to test if there were morphological signatures of hybridization in the temperate racemose group that correspond to the genomic signatures of hybridization detected. Although the model was not trained to classify temperate racemose species, we expect that if the temperate racemose group arose via hybridization, species in this group may retain some morphological features from both the solitary/corymbose group and the tropical racemose group, if these two groups are the parental participants in ancient hybridization. The second classifier model we developed had the goal of distinguishing between four major groups: temperate diploid, temperate racemose, neotropical racemose, and paleotropical racemose. This second model was trained and validated using a total of 4,117 herbarium sheet images representing 81 species. In each group, 80% of the labeled input data were used for training, and 20% for validation. The model was run for 24 epochs until an accuracy rate of 91.3% was achieved. This second classification model was built so we could ultimately measure the breadth of morphological variation across specimens that could be successfully categorized in one of the four groups using a dimensionality reduction approach called IVIS (described below).

#### Gradient CAM

As a further ground-truthing step to ensure that the classification algorithm was assessing biologically meaningful variation, as opposed to learning biologically meaningless features for identification (e.g., labels, rulers), we used Gradient Class Activation Mapping (Grad-CAM), an approach for visualizing how deep neural networks make their predictions (Selvaraju et al. 2017). Briefly, Grad-CAM generates a heatmap highlighting the regions in an input image most important for why a particular prediction was made. A Grad-CAM algorithm takes the output of a convolutional layer in the neural network and computes the gradient of target category in relation to that layer’s activations, which are the pixels that are ‘activated’ by the model—the pixels that are most important for determining the class. The gradients of target category are then unsampled and overlaid onto the query image to generate a heatmap highlighting pixels and regions of the image most important for determining the prediction. Grad-CAM specifically excels when applied to Convolutional Neural Networks (CNNs), due to their ability to learn and extract useful features from images. By interrogating the pixels of an image that the model considers important, Grad-CAM can inform why and how a CNN makes its predictions. We sampled the final layer of the CNN, as well as the second-to-last and third-to-last layers. We categorized the heatmap images as classifying based on leaf features, floral features, or neither/ambiguous. Each image with both leaf and floral structures present was classified as ‘floral’, ‘mixed’, or ‘leaf’. We also assigned images that did not fall into the above categories as ‘leaf with no floral structure present’, ‘ambiguous’, or ‘problematic’. The last two categories were for images where it was difficult to tell if the heatmap favored either floral structures or leaves, or if the heatmap was overly focused on non-biological information, respectively.

#### Morphological space in tropical and temperate groups

IVIS (Implicit Variational Information Sharing) is a deep learning approach with the goal of mapping high-dimensional data into low-dimensional space. Critically, IVIS works well with image data and can preserve distances between samples in a global sense, which is not always true using other dimensionality reduction methods such as PCA, t-SNE, or umap (Szubert et al. 2019, Chari et al. 2021). In the case of image data, IVIS works by encoding the input image into a lower-dimensional representation, and then using a decoder to reconstruct the image. During training, the network learns to form a compact representation by minimizing the reconstruction loss and encouraging the low-dimensional space to be continuous and well-structured through the use of a regularization term. The low-dimensional representation learned by IVIS can be used for various tasks, such as clustering, visualization, or further fine-tuning for classification tasks. IVIS has been shown to be effective for image data, outperforming traditional methods such as PCA and t-SNE in terms of clustering and visualization quality. Here, we use IVIS to reduce the highly-dimensional variation present in image data into 2-dimensional space to make comparisons among specimens and groups.

## RESULTS

### Phylogenomic analyses

For 99 *Prunus* species representing all major lineages, we assembled a dataset of 587 nuclear genes and complete chloroplast genomes to infer phylogeny. The nuclear coalescent species tree inferred using ASTRAL (Fig. 2A) recovered monophyletic corymbose and solitary groups, but a paraphyletic racemose group. With our robust sampling, the solitary, corymbose, and temperate racemose species form a clade, although with modest support (37.9% ASTRAL quartet support (QS) score; see node Temp (“Temperate”), Fig. 2A). This clade of solitary, corymbose, and temperate racemose taxa was sister to a large clade composed of tropical racemose species, including representatives from both the pics and paleotropics. The monophyly of each of the solitary and corymbose clades was strongly supported (i.e., > 70% QS; Fig. 2A). The sister relationship between the solitary and corymbose clades also had strong support (73.9% QS score; node Dip (“Diploid”), Fig. 2A). There were also several other strongly supported clades, including the paleotropical subgenus *Pygeum*, and the clade of neotropical racemose species (see nodes APal (“Australasian paleotropical”), EPal (“East Asian paleotropical”) and SNeo (“Neotropical, predominantly South America”), respectively; Fig. 2A).

The paleotropical racemose species did not form a clade – the neotropical racemose species were nested within the paleotropical group (Fig. 2A). A clade composed of nearly all tropical racemose species, except for a few early-diverging lineages (*Prunus undulata* Buch.-Ham. ex D.Don, *P. laurocerasus* L., *P. africana* (Hook.f.) Kalkman, and *P. skutchii* I.M.Johnst.), had relatively strong support (63.0% QS score; see node CTrop (“Core Tropical”), Fig. 2A). Additionally, the New World (primarily neotropical) racemose species were not monophyletic. Whereas the exclusively South American neotropical racemose species formed a strongly supported clade, four other New World racemose species (*P. ilicifolia* (Nutt. ex Hook. & Arn.) D.Dietr., *P. lyonni* (Eastw.) Sarg., *P. occidentalis* Sw., and *P. cornifolia* Koehne) formed a clade with low support that was sister to the Australasian paleotropical racemose clade (Fig. 2A). The concatenation tree also indicated that all temperate species formed a clade, albeit with low support (42% bootstrap support; Supplemental Fig. S4). The concatenation tree was largely congruent with the ASTRAL tree; one notable exception is that the early-diverging racemose species *P. laurocerasus* and *P. undulata* were sister to the temperate clade in the concatenation phylogeny (Supplemental Fig. S4).

In contrast to the nuclear phylogeny, the chloroplast tree indicated that each group defined by inflorescence morphology formed a clade (Fig. 2B). Each of the solitary-flower and corymbose clades had high bootstrap support (100% and 99%, respectively), and the sister relationship between these two clades was also strongly supported (99%; Fig. 2B). Within the major clades, many of the relationships in the chloroplast phylogeny were similar to the nuclear phylogeny. The neotropical racemose group was similarly non-monophyletic in the chloroplast tree, and additionally the New World racemose species did not form a clade in the chloroplast phylogeny (Fig. 2B). Notably, two species that were early-branching racemose lineages in the nuclear tree (the European *P. laurocerasus* and the East and southeast Asian *P. undulata*), were nested within the temperate racemose species (Fig. 2B).

**Figure 2B.**
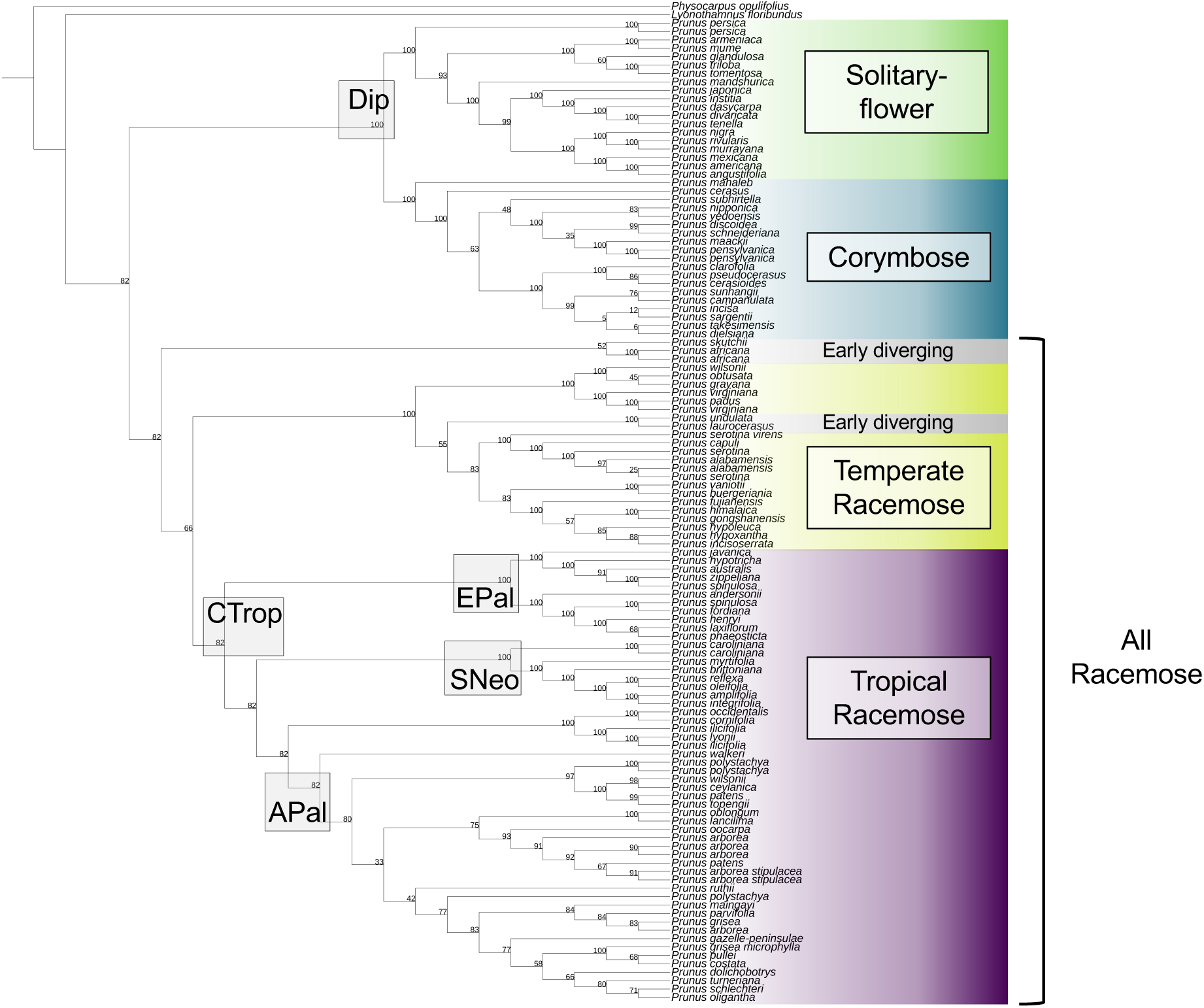
The phylogenetic tree based on chloroplast genomes. Several key nodes that are congruent with the nuclear species tree are highlighted and labeled. The numbers at each node represent bootstrap percentage scores from the RAxML analysis.

#### Gene tree discordance

Many nodes in the nuclear species tree were characterized by rampant gene tree discordance, as assessed by *phyparts* (Supplemental Fig. S5). At most nodes, there were far more gene trees that were discordant with the species tree topology than were congruent (Supplemental Fig. S5). Of note, fewer than 1% of gene trees were concordant with the species tree at both the node defining the clade temperate racemose + solitary + corymbose, and the node defining the tropical racemose clade (Fig. 2A). Within major clades, some nodes had high concordance between gene trees and the species tree (e.g., the node defining a group of neotropical racemose species), whereas others such as the node that defined the temperate racemose species, had low concordance (Supplemental Fig. S5).

#### Reticulate topology

Given existing hypotheses of multiple hybridization events in *Prunus*, we investigated multiple sets of species representing all major clades to better understand the variation in reticulation events in this clade. The 20 networks with one reticulation often demonstrated hybridization edges originating with the outgroup (Supplemental Fig. S6). The 20 networks with two reticulation events frequently indicated hybridization with the outgroup, but also reticulation within the focal clade (Fig. 3, Supplemental Fig. S6). Finally, the 20 networks with three reticulations all indicated at least two reticulation events among the major lineages of *Prunus* (Supplemental Fig. S6). Six of the 20 networks with one reticulation supported a hybrid origin of the temperate racemose group. The other 14 one-reticulation networks had reticulation events leading to the corymbose clade, tropical racemose clade, or corymbose/solitary-flower clade (Supplemental Fig. S6). In contrast, 16/20 and 17/20 networks with two and three reticulations, respectively, displayed reticulation implying a hybrid origin of the temperate racemose group (Supplemental Fig. S6).

**Figure 3.**
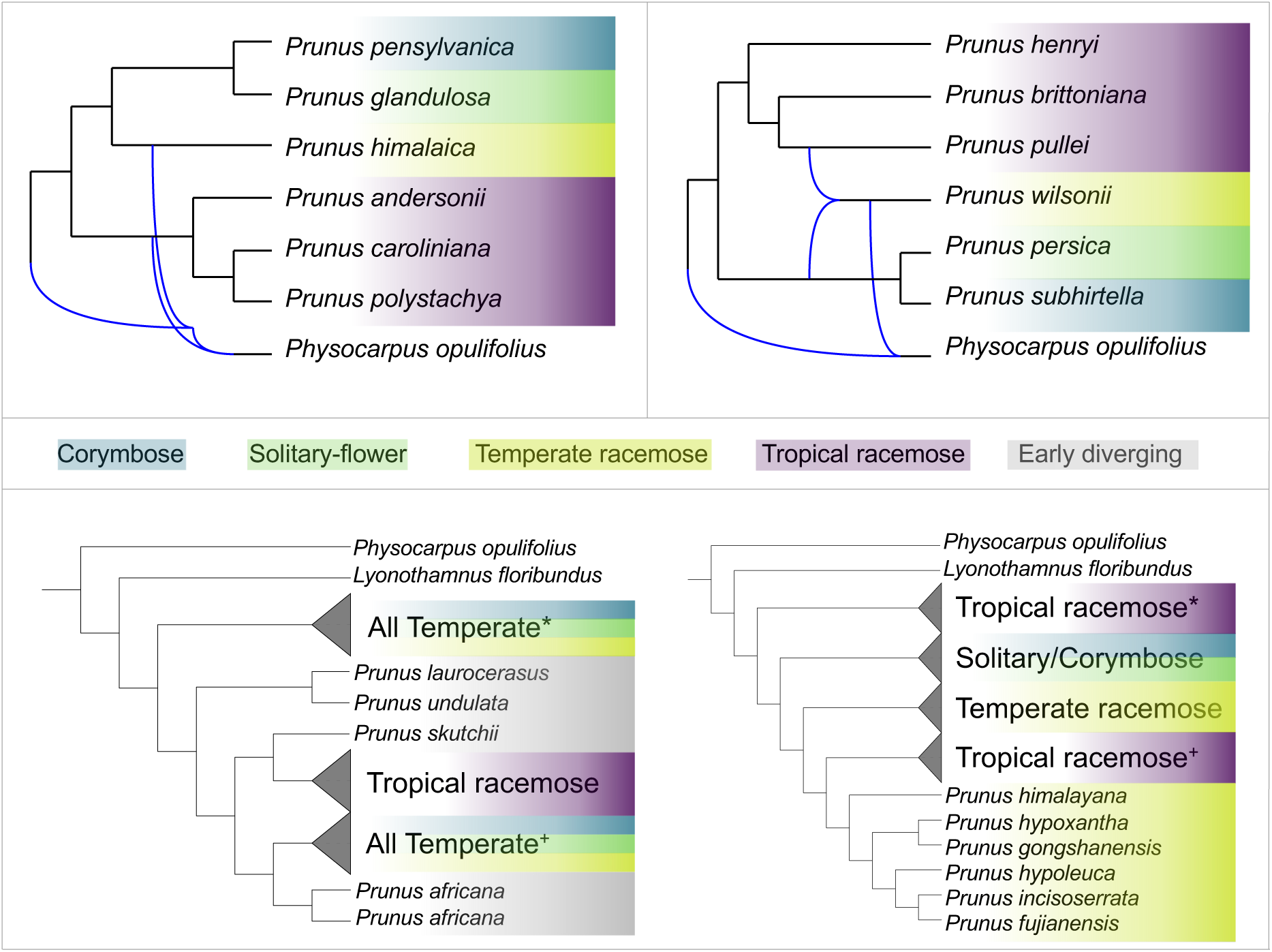
Two representative optimal 2-hybridization edge phylogenetic networks estimated using phylonet for seven species representing every major lineage in *Prunus* (top). In the network on the left, reticulation edges from the outgroup connect to the species representing the temperate racemose clade (*P. himalaica*) and the tropical racemose clade (*P. andersonii*, *P. caroliniana*, *P. polystachya*). In the network on the right, reticulation edges connect the temperate racemose species (*P. wilsonii*) to the outbroup, and also indicate that the temperate racemose species is sister to both the tropical racemose group (*P. henryi*, *P. brittoniana*, *P. pullei*) and the solitary-corymbose group (*P. persica*, *P. subhirtella*), suggesting hybridization leading to the temperate racemose group. The bottom portion shows the two most parsimonious GRAMPA results, which suggest an allopolyploid origin of the entire temperate clade (i.e., temperate racemose, solitary-flower, corymbose groups) (left; most parsimonious). The second-most parsimonious GRAMPA tree indicates an allopolyploid origin of the tropical racemose clade (right).

#### Allopolyploid origin: GRAMPA

The tests for signatures of allopolyploidy using the program GRAMPA indicated that at many of the nodes we investigated, a MUL-tree was more parsimonious than a singly-labeled tree. The most parsimonious MUL-tree, with a parsimony score of 93,501, was characterized by a duplicated clade that comprised all species in the temperate group (Fig. 3). The second most parsimonious MUL-tree (score = 107,277) indicated an allopolyploidy event gave rise to the tropical racemose clade (Fig. 3). We also detected evidence of allopolyploidy leading to the temperate racemose clade (score = 115,580), tropical racemose clade (excluding early-diverging species; score = 118,993), to the East Asian paleotropical clade (score = 126,618), and to the Australasia paleotropical clade (score = 117,855) (Supplemental Fig. S7). Notably, when we separately investigated the nodes defining the South American neotropical clade, the solitary-flower clade, and the corymbose clade, the singly-labeled tree was most parsimonious in each case, implying no evidence of allopolyploidy was detected at these nodes (parsimony scores of 126,710, 124,649, and 125,303, respectively; Supplemental Fig. S7).

The analysis to map WGDs to the species tree identified several nodes with relatively high percentages of duplications. The crown *Prunus* node indicated a duplication percentage of 79.66% (Supplemental Fig. S8). The node with the second highest duplication percentage is the one defining the tropical racemose species excluding the early diverging species (i.e., ‘CTrop’ node from Fig. 2A) at 8.47% (Supplemental Fig. S8). Three other nodes within the tropical racemose clade had duplication percentages >4%: the South American neotropical species (‘SNeo’ node from Fig. 2A), the Australasian paleotropical species (‘APal’ node from Fig. 2A), and a small clade comprised of *P. australis*, *P. hypotricha*, *P. zippeliana*, and *P. spinulosa* (Supplemental Fig. S8). Within the temperate clade, the node defining the solitary-flower group had a duplication percentage of 4.24% (Supplemental Fig. S8).

#### Environmental characterization

The PCA of environmental variation based on the 19 bioclimatic variables captured 78.8% of variation on the first three principal components, with 46.5% of the variation encapsulated in PC1, 17.3% in PC2, and 15.0% in PC3 (Fig. 4). One precipitation variable—annual precipitation—was strongly positively correlated with PC1, and four temperature variables—mean annual temperature, minimum temperature of the coldest month, mean temperature of the driest quarter, and mean temperature of the coldest quarter—were also strongly associated with positive PC1 space. Meanwhile, two temperature variables—temperature seasonality, and mean annual temperature range—were strongly and negatively associated with PC1 values. No bioclimatic variables were strongly correlated with PC2 or PC3. In summary, tropical species occupy more positive regions of PC1, which correspond to warmer and wetter conditions, with higher minimum temperatures in colder seasons, and less temperature seasonality (Fig. 4), whereas temperate species trend towards negative values of PC1. When considering the two types of tropical species—neotropical versus paleotropical—the paleotropical species extended further towards both extremes of PC1, and further towards the negative extent of PC2. Neotropical species extended slightly further into positive PC2 space than paleotropical species. The temperate racemose species had a greater affinity for positive values of PC1 whereas corymbose/solitary species trended further towards extreme negative values of PC1. The temperate racemose species occupied a larger range of PC2 space than the corymbose/solitary species (Fig. 4). There were clear elevational differences among species representing the four major groups. In the temperate solitary/corymbose group, only two out of 28 species had a median elevation greater than 1,000m (Supplemental Fig. S9). In contrast, half of the 10 temperate racemose species occurred at a median elevation greater than 1,000m, and also half of the 10 neotropical racemose species were found at a median elevation greater than 1,000m. In the paleotropical racemose species, 13 of 28 species had a median elevation greater than 1,000m (Supplemental Fig. S9).

**Figure 4.**
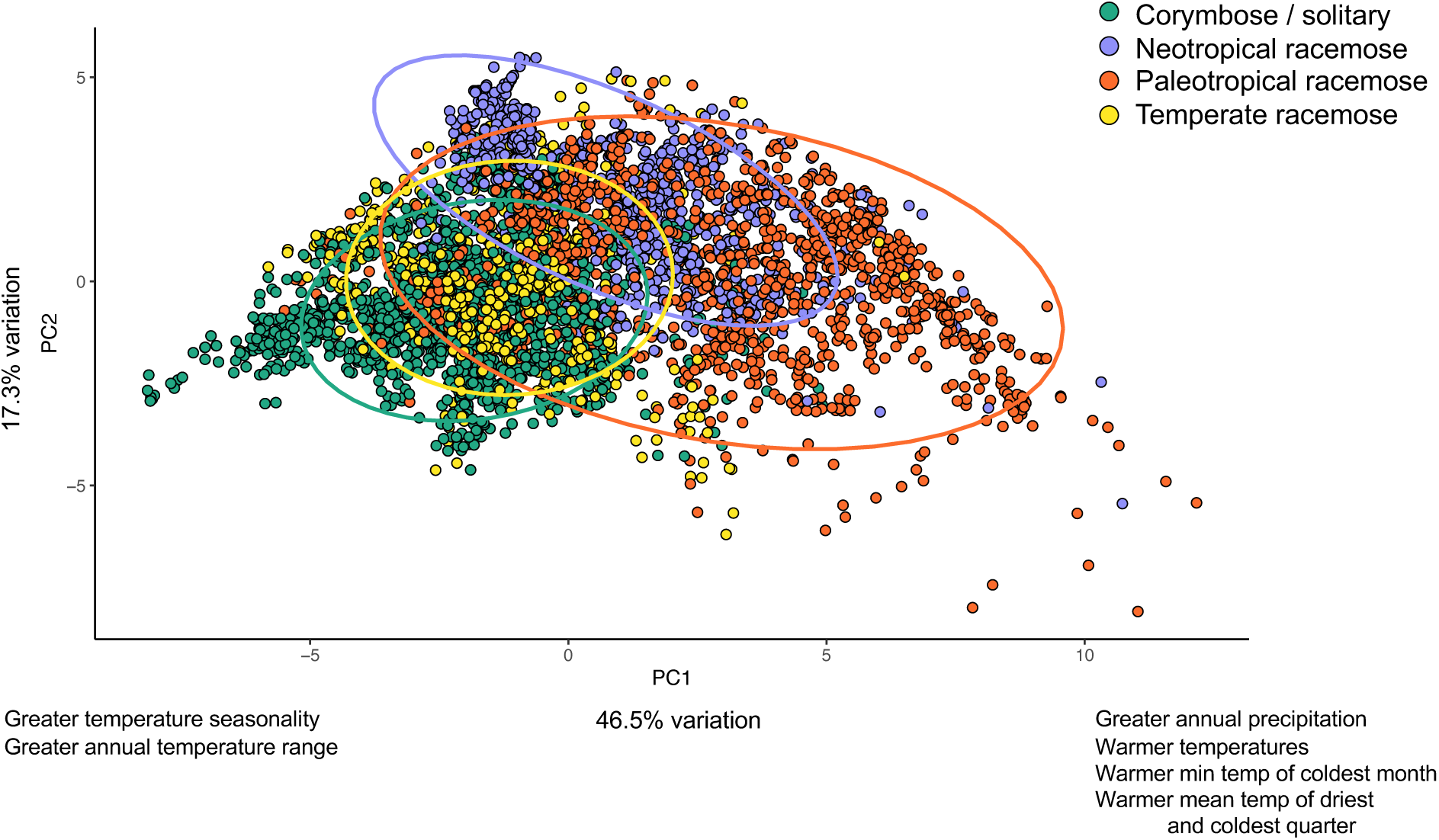
The summary of environmental variation using a Principal Components Analysis to distill the 19 bioclimatic variables into 3 dimensions. PC1 accounts for 46.5% of variation while PC2 explains 17.3% of environmental variation. Several bioclimatic variables strongly correlated with PC1 are indicated on the X-axis. Negative PC1 space is associated with increased temperature seasonality and annual temperature range. Meanwhile, positive PC1 space corresponds to higher annual precipitation, annual mean temperature, minimum temperature of the coldest month, and mean temperature of the driest quarter and coldest quarter.

#### Biogeographic analysis

The DEC+J model was the preferred model after comparing AIC scores of competing geography-based BioGeoBears models, which were based on the time-calibrated tree generated with TreePL (Supplemental Table S4). Likelihood ratio tests concluded that the addition of the J parameter, despite producing a more parameter-rich model, led to a significantly better model than DEC (Supplemental Table S5). In contrast, the BAYAREALIKE model was favored by AIC scores for the biome-based BioGeoBears model (Supplemental Table S4). Here, likelihood ratio tests did not indicate that the +J model was significantly better (Supplemental Table S5). The geography-based analysis indicated an East Asian origin of the genus, with the initial diversification occurring in the Paleocene ca. 60-55 Mya (Fig. 5). Subsequently, the temperate racemose, solitary/corymbose, and tropical racemose clades all diverged by the early-mid Eocene (ca. 50 Mya). This diversification occurred predominantly in East Asia and North America, which were characterized by the Boreotropical climate during the Late Cretaceous – early Eocene (ca. 65-50 Mya). The temperate racemose group steadily diversified in East Asia and North America beginning in the mid-late Eocene (ca. 40 Mya). Both the solitary and corymbose lineages began diversifying in the early Miocene (ca. 25-20 Mya), first in East Asia and subsequently in North America and Europe. These two lineages currently occupy temperate environments, and based on our sampling, they were more successful in speciating in East Asia versus North America (Fig. 5).

**Figure 5.**
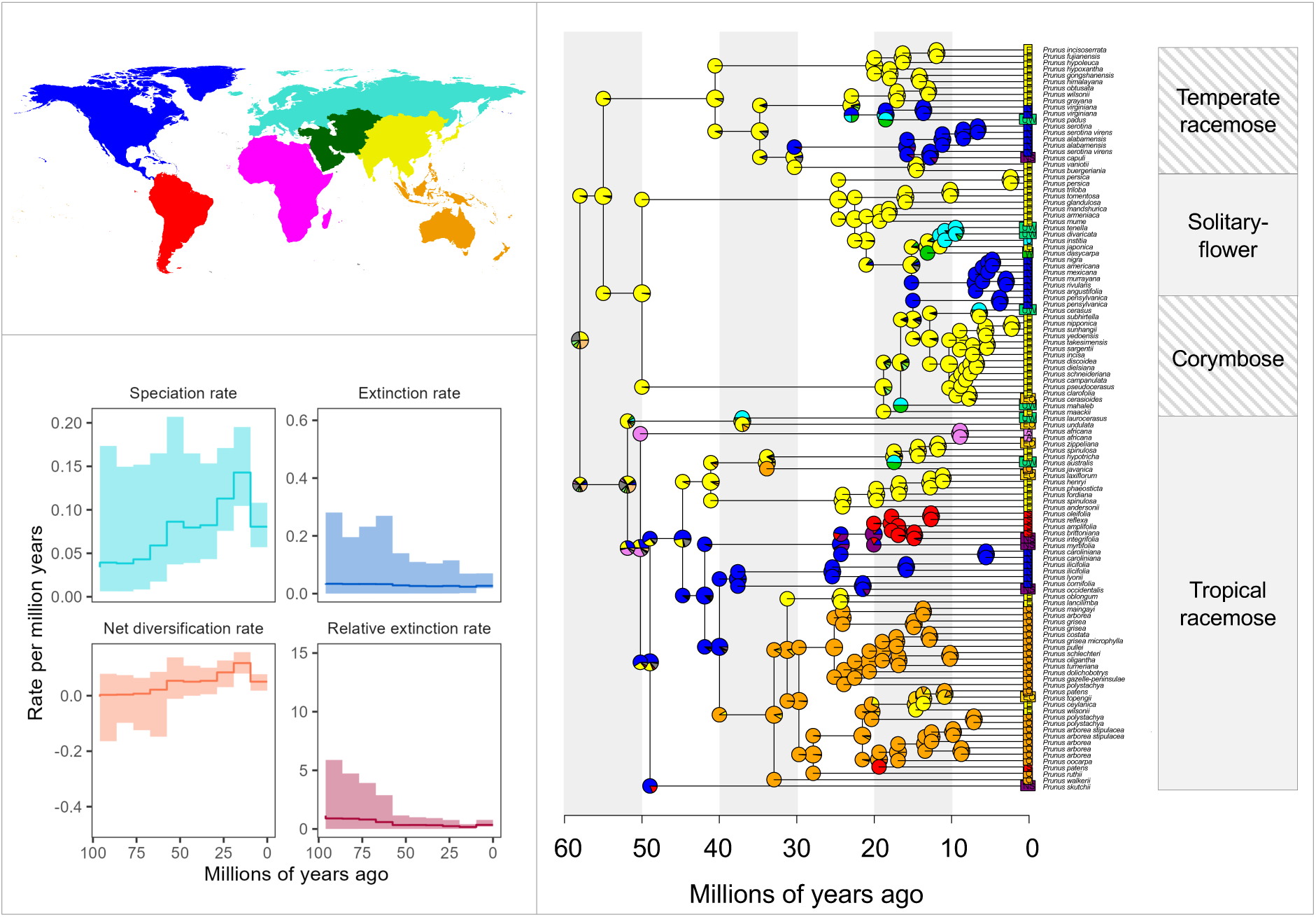
The BioGeoBears analysis (right) of the biogeographic history of *Prunus* using the DEC + J model based on seven biogeographic regions shown in the upper left: North America (blue), South America (red), Africa (magenta), Europe (turquoise), West Asia (green), East Asia (yellow), and Southeast Asia (orange). Color-coded pie charts at each node indicate the probability of the ancestral state occurring in a given biogeographic region. The episodic birth-death (EBD) model diversification analysis is shown in the lower left. The speciation rate and net diversification rate of the clade are highest during the Miocene, between approximately 20-10 Mya.

Broadly, the tropical racemose clade’s early diversification occurred in several regions, including East Asia, West Asia, Africa, North America, and Europe, between 55-40 Mya. Early-diverging lineages, including species such as *Prunus laurocerasus*, *P. undulata*, *P. africana*, and *P. skutchii*, split off during the Eocene between ca. 55-40 Mya. Notably, the ancestors of these few early-diverging lineages occupy a wide geographic range, including Europe, West Asia, East Asia, Southeast Asia, Africa, North America, and South America. Subsequently, the patterns of diversification occurred differently in the major lineages within the tropical racemose group. The neotropical lineages, occurring in tropical North and South America, are characterized by a long period of stasis, followed by rapid diversification in the Miocene beginning between 25-20 Mya and lasting until ca. 10 Mya. In contrast, two major paleotropical lineages, in East Asia and Southeast Asia, respectively, began steadily diversifying in the Oligocene (ca. 35 Mya), and continued until approximately 10 Mya. Notably, there were multiple clades in both the temperate and tropical groups of *Prunus* that demonstrated rapid diversification beginning in the early Miocene.

The biome-based biogeographic analysis indicated that the ancestral biome state of the clade consisted of both tropical and temperate biomes (Supplemental Fig. S10). In the portion of the clade that presently occupies temperate environments (top clade; Supplemental Fig. S10), there were transitions to other biome categories, including temperate (many corymbose species) and dry/temperate (many solitary-flower species) beginning approximately 20-15 Mya (Supplemental Fig. S10). In the temperate racemose clade, the biome-based analysis indicated that part of this lineage retained its ancestral biome-affinity (green pies), whereas other species transitioned from the tropical/temperate biomes to dry/temperate/tropical, dry/temperate, temperate, or cold/temperate beginning around 15 Mya (Supplemental Fig. S10). In the tropical racemose clade, the ancestral state of temperate/tropical transitioned to predominately tropical environments from approximately 50 to 30 Mya (Supplemental Fig. S10). Then, portions of the paleotropical racemose group in subgenus *Laurocerasus* shifted back to temperate/tropical environments (top portion of tropical racemose clade; Supplemental Fig. S10). Meanwhile, neotropical species either remained tropical, or transitioned to dry/temperate, dry/temperate/tropical, or cold/tropical between 25-10 Mya (Supplemental Fig. S10). In the bottom portion of the clade, lineages either remained tropical, or transitioned to cold/tropical or temperate/tropical approximately 25-10 Mya (Supplemental Fig. S10).

The stochastic mapping analysis clarified the timing of transitions between geographic regions and biomes (Supplemental Fig. S11). The vast majority of transitions have taken place since 25 Mya—either as cladogenesis events at nodes—or anagenetic shifts along branches (Supplemental Fig. S11). The anagenetic transitions in particular frequently occurred in the past 15 million years (Supplemental Fig. S11). In the geography-based analysis, the majority of biogeographic transitions between regions was sympatric speciation, with smaller proportions of founder events and anagenetic dispersal (Supplemental Table S6). Most biogeographic events in the biome-based analysis were sympatric, meaning that there were speciation events without a corresponding biome shift. Approximately 30% of the biogeographic events were anagenetic dispersal—or transitions between biomes along a branch (Supplemental Table S6). In the geography-based biogeographic analysis, East Asia acted as a major source, with especially strong dispersal to N. America, Europe, and Oceania (Supplemental Table S7). Europe and West Asia underwent substantial reciprocal dispersal, and North America acted as a source to South America’s sink (Supplemental Table S7). In the biome-based biogeographic analysis, both the temperate and tropical biomes acted as key sources for other biomes (dry, cold). The dispersal rate from tropical to temperate was larger than the rate from temperate to tropical (Supplemental Table S7).

#### Diversification analysis

To investigate the variation in diversification rates across the phylogeny, we implemented an episodic birth-death model (EBD; Höhna 2015) and a branch-specific diversification model (LSBDS; Höhna et al. 2019) in RevBayes (Höhna et al. 2016). The EBD model estimated that the net diversification rate peaked approximately 20-10 Mya; this was driven by the speciation rate, which also was at its maximum between approximately 20-10 Mya (Fig. 5). The LSBDS model, which shows branch-specific diversification, generally showed lower net diversification in the early-diverging tropical racemose species, with similar net diversification in most other lineages (Supplemental Fig. S12). The CRABS analysis confirmed results from the EBD model; the extinction rate remained low, whereas the speciation rate increased to a peak approximately 10 Mya, followed by a decline (Supplemental Fig. S12).

#### Morphological analyses

We ran two experiments using computer vision-based machine learning analyses of digitized herbarium sheets. The assessment of morphological characters associated with hybridization revealed that whole sheet phenotypes of temperate racemose species, with the top 20% and bottom 20% of the sheet trimmed, were nearly evenly divided between classification as ‘solitary/corymbose’ (N = 310 observations with >90% probability) and ‘tropical racemose’ (N = 229 observations with >90% probability) (Fig 6). When using whole sheets (i.e., untrimmed), there was a similar trend: there were N = 279 observations classified as ‘solitary/corymbose’ with >90% probability and N = 223 observations classified as ‘tropical racemose’ with >90% probability. The Grad-CAM analysis of internal model layers indicated that inflorescence vs. leaf morphology was the driving factor in determining if the machine learning model inferred whether to classify temperate racemose species as corymbose/solitary or tropical racemose (Fig. 6). Manual visual inspection indicated that the second-to-last and third-to-last layers were most informative for determining classifications. The Grad-CAM results for the last three model layers for every herbarium sheet specimen image are shown in Supplemental Tables S8 and S9. Training and test data for all models are listed in Supplemental Tables S10 and S11.

**Figure 6.**
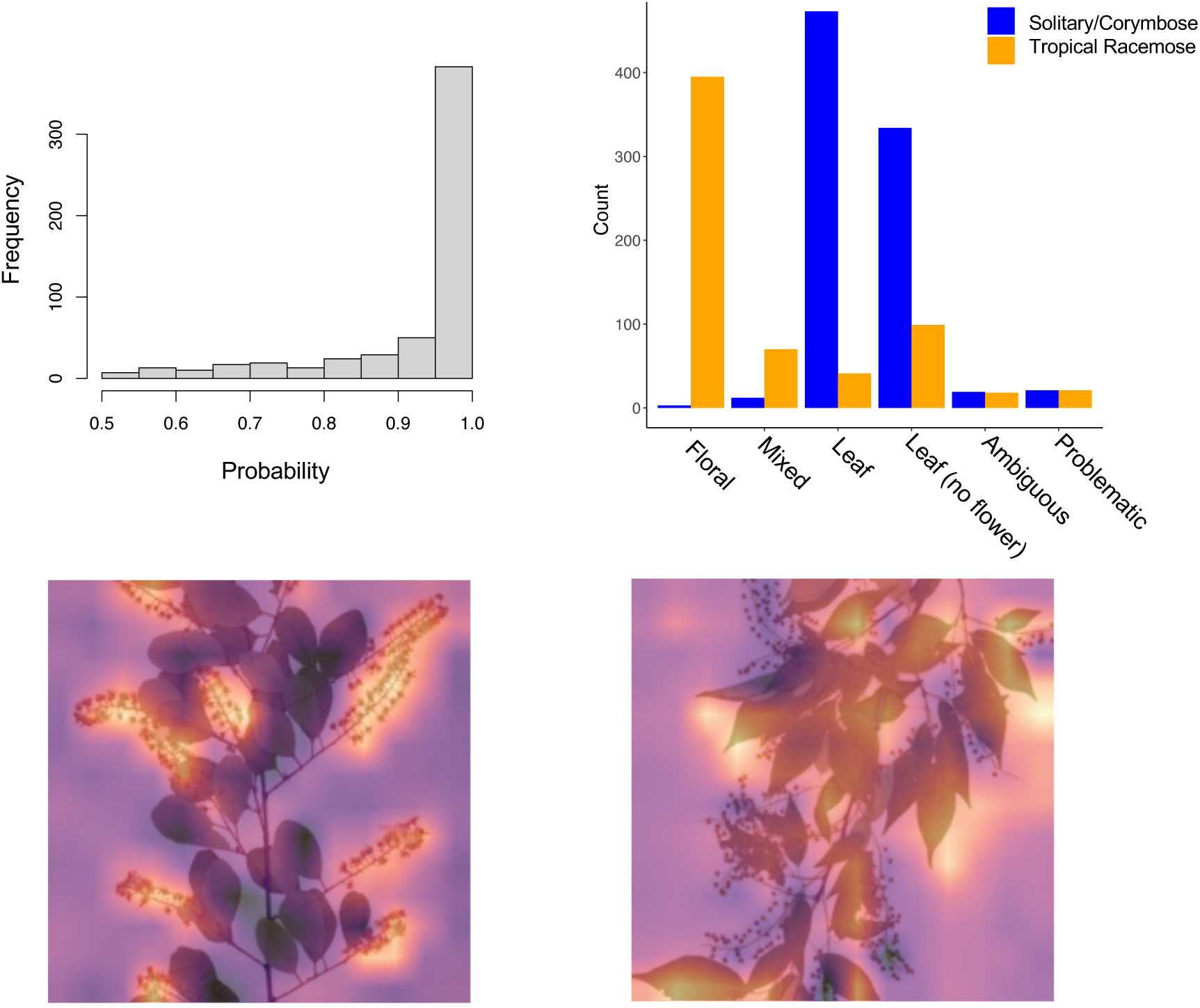
In the upper left, the distribution of probability scores when using the temperate racemose species as test data in the first machine learning model. These classifications include specimens that were classified as either ‘solitary/corymbose or tropical racemose’; regardless of classification, most specimens were assigned with high (i.e., >0.95) probability. The plot in the upper right shows the manual classifications we assigned when ground-truthing the gradient class activation maps for the 676 temperate racemose specimens for the second-to-last and third-to-last layer of the CNN. The bottom row shows the gradient class activation (Grad-CAM) maps for the second-to-last model layer for two representative temperate racemose specimens that were classified as ‘tropical racemose’ (left) and ‘solitary/corymbose’ (right), respectively.

In the second analysis, breadth of morphospace was measured using IVIS to quantify and compare the extent of morphological variation present in the following groups: temperate diploid (i.e., corymbose and solitary flower), temperate racemose, neotropical racemose, and paleotropical racemose. The IVIS analysis determined that the two tropical groups exhibited a greater range of morphological variation than the two temperate groups (Fig. 7). Within all tropical specimens, the paleotropical racemose species occupied a greater expanse of morphological variation than neotropical racemose (Fig. 7). The paleotropical species overlapped in morphospace to a small degree with the temperate diploid group (i.e., corymbose/solitary) and to a much greater degree with the neotropical racemose group. The neotropical species are positioned centrally in morphospace and overlap with all three other groups (Fig. 7).

**Figure 7.**
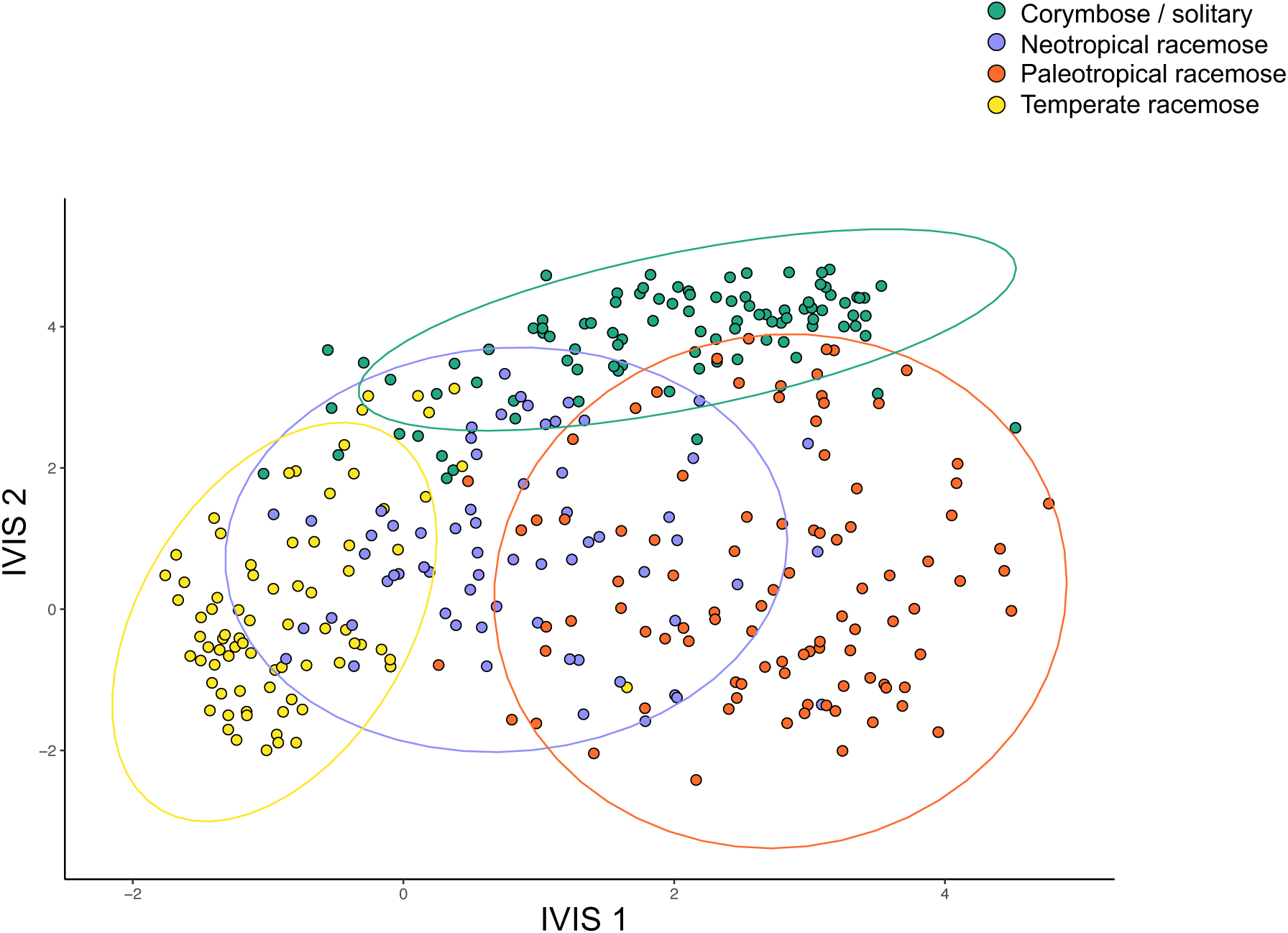
The representation of whole specimen phenotypes in IVIS space. The specimen images were assigned to four groups according to environmental preference/clade: neotropical racemose, paleotropical racemose, temperate racemose, corymbose/solitary.

## DISCUSSION

In this study, we provided increased resolution compared to previous studies of the phylogeny of the economically and ecologically important genus *Prunus*, pinpointed key genomic mechanisms promoting the diversification of this group, and improved our understanding of its biogeographic history. By sampling more densely the understudied racemose group and sequencing hundreds of nuclear loci and complete chloroplast genomes, we changed our understanding of how several groups within the genus diversified. Specifically, by combining analyses of hybridization, genome doubling, and cytonuclear and gene tree-species tree conflict, we inferred that the polyploid racemose group is paraphyletic, a result found in some previous studies using nuclear gene data (e.g., Lee & Wen 2001, Bortiri et al. 2002, 2006, Wen et al. 2008, Chin et al. 2014, Zhao et al. 2016), but typically not found when using plastid genes (e.g., Bortiri et al. 2001, Wen et al. 2008, Chin et al. 2014). The chloroplast phylogeny, with a monophyletic racemose group, detected in the present study is congruent with many previous studies using chloroplast markers (e.g., Bortiri et al. 2006, Wen et al. 2008, Chin et al. 2014, Zhao et al. 2016). Notably, our result conflicts with Su et al. (2023), which found a monophyletic racemose group in both nuclear and chloroplast datasets. However, the nuclear SNP markers used in Su et al. (2023) may have been unable to fully untangle the reticulate history of this genus. Previous studies have suggested that cytonuclear discord observed in *Prunus* phylogenies could be attributed to one (Chin et al. 2014) or multiple (Zhao et al. 2016) allopolyploid hybrid origins of the racemose group. Here, we isolated specific lineages as participants in likely hybridization and allopolyploidy events during the early Eocene ca. 50 Mya. These reticulation events early in the diversification of the genus help explain the high gene tree-species tree conflict and cytonuclear discord. Through biogeographic, diversification, and fossil-calibrated dating analyses, we traced the biogeographic history of the group and tied key climatic events to radiations of subgeneric lineages.

Using several novel applications of machine learning algorithms to classify and quantify morphological variation, we addressed how morphological evolution coincides with cytonuclear discord, identifying characters essential for survival in specific environments and how morphological variation in lineages impacts their biogeographic distribution. Phylogenomic data have revealed that gene tree-species tree conflict is widespread and may sometimes arise via ancient hybridization (e.g., McVay et al. 2017, Nie et al. 2023). Here, we demonstrate that a well-trained machine learning classification model struggles to classify specimens in a lineage that is the result of ancient hybridization, as these specimens that are the result of hybridization display morphological features common to both putative ancestors. Additionally, we applied a machine learning approach, IVIS, to quantify the morphological variation within major lineages, revealing that the distinct morphospaces of different lineages may be shaped by their biogeographic and evolutionary history. These morphological classifications, in combination with environmental data, change our concept of how lineages may have diversified in the tropics, and the conditions under which lineages transition from tropical to temperate biomes. Below, we discuss and contextualize our results.

### Improved understanding of Prunus phylogeny, diversification, and biogeography

Despite its economic and ecological importance, until now *Prunus* has not been thoroughly investigated in a phylogenomic context with sufficient sampling in the racemose group. Previous studies, based on chloroplast DNA and/or a few nuclear markers, inferred possible cytonuclear conflict—a telltale sign suggesting hybridization—on the backbone of the phylogeny. These analyses lacked the resolution to conclude whether the cytonuclear discord was real or due to unresolved nuclear relationships (Lee & Wen 2001, Bortiri et al. 2001, Wen et al. 2008). Based on cytonuclear discord (Chin et al. 2014), multiple copies of nuclear loci (Zhao et al. 2016), nuclear reduced representation sequencing (Su et al. 2023), and chromosome count data coupled with multiple gene tree topologies (Hodel et al. 2021), hypotheses of ancient allopolyploid hybridization have been proposed (Zhao et al. 2016). Here, multiple lines of evidence indicate a history of allopolyploidy and/or hybridization driving early diversification in *Prunus*. First, we demonstrate that the cytonuclear discord along the backbone of the genus, especially the differing phylogenetic placement of the temperate racemose group, provides a specific hypothesis of reticulation (i.e., the temperate racemose group is a product of hybridization and/or allopolyploidy). Phylonet analyses with 2- and 3-reticulations supported a likely hybrid origin of the temperate racemose group. When 1-reticulation networks were also considered, the reticulation analysis suggested possible hybrid and/or polyploid origins of the tropical racemose group, solitary-flower clade, and corymbose clade (Fig. 3, Supplemental Fig. S6). The GRAMPA analysis, which can detect and distinguish between different types of genome doubling, also identified reticulation deep in the tree. The temperate clade was inferred to have the best-supported allopolyploid origin deep in the phylogeny (Fig. 3, Supplemental Fig. S7). However, additional clades, including the tropical racemose clade, also were inferred to likely be the result of allopolyploidy (Fig. 3, Supplemental Fig. S7). Furthermore, mapping WGDs to nodes in the phylogeny identify several major clades with evidence of duplication (e.g., tropical racemose clade, excluding early-diverging species; Supplemental Fig. S8). Taken together, these analyses point to reticulation deep in the phylogeny, although it is not constrained to one instance of hybridization and/or allopolyploidy. These results are in line with previous hypotheses that predicted multiple rounds of allopolyploidy (Zhao et al. 2016). Probably the majority, if not all, racemose species, encompassing both temperate and tropical climates, are polyploid. These species likely arose from multiple rounds of ancient allopolyploidy rather than autopolyploidy. Only one of the GRAMPA analyses could possibly be autopolyploidy—the Australasian paleotropical group (Supplemental Fig. S7C). All other WGD events demonstrated that the duplicated clades were spread throughout the tree, necessarily implying allopolyploidy (Supplemental Fig S7).

The higher resolution, time-calibrated plum genus phylogeny enhances our understanding of the biogeographic history of the clade. The diversification and biogeographic history of *Prunus* can be contextualized by existing hypotheses describing biogeographic patterns, and it can shift our understanding of the range of possible outcomes for lineages that originated in the tropics. Our biogeographic and diversification results show pulses of diversification in several tropical and temperate *Prunus* lineages, with bursts of speciation in the early Miocene (Fig. 5, Supplemental Fig. S10). This contrasts with steadier diversification reported in previous studies (e.g., Chin et al. 2014). Our time-calibrated results also highlight the different patterns of speciation in different tropical *Prunus* groups: relatively steady speciation in the paleotropics, with punctuated bursts in the neotropics (Fig. 5). There were also key differences in greater temperate success in Asia versus North America: the temperate lineages in Asia speciated readily and rapidly while the species diversity in North America is depauperate. Although our sampling of species diversity and nuclear loci gives better resolution than previous studies, there are still limitations when interpreting our biogeographic results. Critically, biogeographic analyses depend on bifurcating phylogenies, and many of our results point to a reticulate topology for *Prunus*, especially deeper in the tree. It is likely that multiple lineages intermingled in the expansive Boreotropical region early in the evolutionary history of the genus. The biome-based biogeographic analysis is consistent with this explanation, where the deeper portion of the tree showed all nodes as either tropical or temperate/tropical (Supplemental Fig. S10). Furthermore, the biogeographic stochastic mapping analysis identified speciation in sympatry as the majority of biogeographic events in both the geography- and biome-based analyses (Supplemental Table S6). Many of the lineages implicated in reticulation, such as the early diverging tropical racemose lineages, may have overlapped during the Eocene, thus facilitating ancient hybridization and/or allopolyploidy. We must acknowledge that inferences of diversification may be affected by sampling bias. We present the densest sampling of the understudied tropical racemose clade to date, with accessions representing all major lineages. However, additional future work is needed to ensure that even unbiased sampling does not impact the results, as this genus is estimated to contain 250-400 species, depending on taxonomic treatments and sampling of tropical species, necessitating further sampling in future studies.

#### Using Prunus to inform broad biogeographic patterns

For over a century, biologists have observed a latitudinal gradient in species diversity in many clades across the Tree of Life, with greater species richness occurring near the equator. However, we lack scientific consensus about the causes of this biogeographic pattern and several hypotheses have been proposed. One explanation for the observed latitudinal gradient is the tropical conservatism hypothesis (TCH), which posits that the relatively massive biodiversity of the tropics can be primarily attributed to the geographic extent of tropical taxa over the past 55 million years and the subsequent evolutionary conservation of environmental niches (Wiens and Donoghue 2004). Many groups have diversified in the Eocene, facilitated by the vast expanse of the Boreotropics. Nevertheless, relatively few lineages have transitioned from tropical environments to temperate ones. This phenomenon may be explained by the challenge of acquiring the substantial adaptations necessary to tolerate the cooler conditions in temperate zones (Donoghue 2008).

Another explanation is provided by the ‘taxon pulse’ hypothesis (Erwin 1985), which is compatible with the TCH and posits that lineages that were able to transition from tropical to temperate regions originated in tropical environments and then migrated in waves to more temperate regions at higher altitudes and latitudes into increasingly harsh environments (Erwin 1985, Lü et al. 2020, Rapini et al. 2021). These radiations into temperate environments would also be accompanied by species loss in ancestral tropical lineages as older lineages went extinct (Liebherr and Porch 2015, Rapini et al. 2021, Nie et al. 2023). Some widespread plant clades, such as *Viburnum*, match the taxon pulse hypothesis to an extent (Spriggs et al. 2015). In this scenario, tropical lineages are posited to be ‘dying embers’ – lineages that diversified in the tropics tens of millions of years ago, and persist in place, but have ceased to continue diversifying (Spriggs et al. 2015).

Our results indicate *Prunus* does not precisely follow the predictions of either the TCH or the ‘taxon pulse’ hypothesis. In *Prunus*, the paleotropical lineages have diversified in place and do not appear to be ‘dying embers’, instead demonstrating steady diversification through time (Fig. 5, Supplemental Fig. S12). As described in the taxon pulse hypothesis, tropical lineages may migrate to higher temperate latitudes or to higher elevation regions in the tropics, which may have temperate-like conditions. Biogeographic stochastic mapping suggested that in *Prunus* there was substantial dispersal from the tropics to all other biome types (Supplemental Table S7). Meanwhile, the diversification of the neotropical lineages, instead of occurring steadily as observed in our paleotropical species, was characterized by bursts of speciation beginning in the late Oligocene-early Miocene (ca. 25-20 Mya). In the tropics, some *Prunus* species have moved to higher elevations, but there are also lineages that occupy truly tropical environments, as defined by some or all individuals occurring in environments with the minimum temperature of the coldest month greater than 18 °C. Among species in our study, these include *Prunus dolichobotrys* (Lauterb. & K.Schum.) Kalkman, *P. arborea* (Blume) Kalkman, *P. gazelle-peninsulae* (Kaneh. & Hatus.) Kalkman, *P. javanica* (Teijsm. & Binn.) Miq.*, P. undulata, P. fordiana* Dunn*, P. grisea* Kalkman*, P. costata* (Hemsl.) Kalkman*, Prunus maingayi* (Hook. f.) Wen*, P. oocarpa* (Stapf) Kalkman*, P. oligantha, P. schlechteri* (Koehne) Kalkman*, P. ceylanica* (Wight) Miq.*, and P. buergeriana* Miq. in the paleotropics, and *P. myrtifolia* (L.) Urb., *P. amplifolia* Pilg., *P. cornifolia, P. integrifolia* (C.Presl) Walp.*, P. occidentalis*, *P. skutchii*, and *P. reflexa* (Gardner) Walp. in the neotropics. We also note that, for the species we sampled genetically in this study, in both tropical groups, at least half of the species occurred at a median elevation of less than 1,000 m (Supplemental Fig. S8). The biome-based biogeographic analysis showed that some lineages and species in the tropics clear remained tropical over the past 20 million years (e.g., *P. andersonii*, *P. oleifolia*, *P. reflexa*, *P. amplifolia* in the neotropics, *P. dolichobotrys*, *P. gazelle-peninsulae*, *P. polystachya* in the paleotropics; Supplemental Fig. S10). At the same time, the analysis demonstrated partial or complete biome shifts in many tropical species, such as from tropical to temperate/tropical, tropical to dry/temperate, and tropical to cold/tropical (Supplemental Fig. S10).

The comparison of the neotropics and paleotropics demonstrates that diversification can occur differently in distinct tropical regions—even within a single genus. Our quantifications of the morphological breadth and environmental breadth revealed both to be greater in the paleotropics as compared to the neotropics. The larger environmental space of the paleotropics suggests there may have been greater ecological opportunity for lineages in the paleotropics relative to the neotropics due to increased niche availability (Wellborn & Langerhans 2015). For similar reasons, the greater morphological variation in the paleotropical *Prunus* shows these lineages were able to leverage advantageous morphological features to adapt to new environmental niches. This implies *Prunus* differs from many lineages with a tropical origin that successfully transitioned to temperate regions. Synthesis has shown that not all lineages could adapt to colder climates when tropical habitat retracted, and typically those lineages that tracked tropical habitats became more restricted (Donoghue 2008). However, *Prunus* bucks this pattern, exhibiting steady speciation in the paleotropics, and later bursts of speciation in the neotropics. It is possible that the neotropical racemose lineages have ancestors that occupied temperate zones in North America (Fig. 5). The small clade of four North American species (Fig. 2A; node ‘NNeo’) is a likely candidate—these species may be Boreotropical remnants in the New World, highlighting the importance of North America as a site of diversification early in the evolution of some tropical racemose species. Perhaps a history of surviving in the temperate zone enabled present-day neotropical lineages to develop and/or keep a suite of morphological characters that facilitated radiating into the neotropics. The substantial overlap in morphospace between neotropical and temperate lineages suggests this may be the case (Fig. 7).

#### Why was Prunus more successful than other lineages?

Because of our sampling of species diversity coupled with hundreds of nuclear loci, we identify allopolyploidy and/or hybridization in the backbone of the phylogeny, which may explain the genomic basis of the rapid diversification 55-45 Mya. This allopolyploidy event is supported by phylogenetic treatments of the Rosaceae, which found evidence of a WGD event at the base of *Prunus* (Xiang et al. 2017); this also is corroborated by the high duplication percentage at the base of the genus in our WGD mapping (Supplemental Fig. S8). This is an important insight for not only understanding the phylogeny of the group but also its biogeographic context. When numerous *Prunus* lineages were able to interact in the Boreotropics during the Eocene, ample opportunities for hybridization and allypolyploidy arose. This genetic reshuffling may explain why *Prunus* was more successful than other lineages at occupying both temperate and tropical regions. The Rosaceae family has diversified into many successful lineages (Potter et al. 2007), but even within this hyperdiverse group, *Prunus* stands out for its diversity of inflorescence types, which may have facilitated migration via a variety of different animal dispersers, as well as variation in leaf morphology supporting both evergreen and deciduous life histories. One key innovation in *Prunus* is the evolution of leaf glands, which appear to be associated with climatic conditions, with flat glands typical of species in tropical climates, and raised glands present in species occupying cooler climates (Chin et al. 2013). These glands may have had adaptive value due to their interactions with insects, with the flat glands found in tropical regions preventing herbivory, and may be a key feature driving the diversification of *Prunus*. Specifically, the variation in these morphological features may have enabled *Prunus* radiations into the temperate zone, while continuing to diversify in the tropics. Other Rosaceae, although successful, remained constrained in temperate regions (Xiang et al. 2017). Additionally, other groups with tropical origins that have experienced great success in temperate regions did not retain much species diversity in tropical regions, including *Quercus* L. (Hipp et al. 2020), *Viburnum* (Spriggs et al. 2015), Juglandaceae (Zhang et al. 2021), and Saxifragales (Folk et al. 2019).

#### Morphological variation informs biogeographic history

The morphological variation associated with recent hybridization in many cases can be readily tracked to inform the parental participants in hybridization, as well as to understand how and why hybridization occurred, and whether hybrids may persist based on their environment. However, in cases of ancient hybridization, it becomes difficult to tease apart the morphological characters associated with hybridization since subsequent diversification and evolution in response to environmental variation can obscure the circumstances surrounding the hybridization event. Here, we used machine learning to quantify morphological variation in clades resulting from reticulation, presenting a way to characterize the morphological legacy of hybridization/allopolyploidy. By using the ancient hybridization detected via cytonuclear discord and phylogenetic networks to guide our morphological analyses, we concluded that there were morphological signals of hybridization in the present species. Members of the temperate racemose group had morphological features of both the diploid clade and the tropical racemose clade—the ancestors of these two lineages were putatively the participants in hybridization. So, we would expect morphological features from both parents to persist in the lineage resulting from reticulation. However, lineages’ responses to environmental variation may also explain patterns of morphological variation not resulting, or minimally resulting, from the genomic results of hybridization (or allopolyploidy). In this case, we would expect that morphological variation would co-vary to some extent with environmental variation. This would lead to an expectation that the vast majority of test specimens from the temperate racemose group would be classified as solitary/corymbose, because the environment overlaps much more between these two groups than it does between temperate racemose and tropical racemose (Fig. 4). Contrary to this expectation, we observed a relatively even split of temperate racemose species between the two training categories.

Morphological shifts can result from transitioning to a novel environment, or existing morphological variation can enable lineages to adapt to changing environmental conditions. The key takeaway from our IVIS analysis is that both the neotropical and paleotropical groups exhibit substantially more morphological variation than do the temperate groups, including both the corymbose/solitary group, and the temperate racemose group. Moreover, the two temperate groups occupy peripheral portions of morphological space on both IVIS axes 1 and 2, suggesting that the morphological features in the temperate groups are specialized relative to the morphological features of the tropical groups. These features may represent morphological changes that promote transitions to temperate regions, diverging from morphological features typical for tropical zones, in response to a transition from tropical to temperate biomes. Broadly, if there is greater niche space available in the tropics—a proposed explanation for the increased species diversity in these regions—we would expect greater morphological variation to occupy more niches. In *Prunus*, the balance of morphological space we quantify is not surprising given greater tropical diversity. Notably, however, the morphological space occupied by the neotropical racemose species overlaps more with the two temperate groups than the paleotropical group does. This suggests an explanation for some of the biogeographic patterns detected—specifically that the more recent diversification of the neotropics was different from the initial diversification in the paleotropics. By the time the neotropical lineages began to diversify, lineages in the genus had already radiated into and adapted to temperate biomes. This may explain the rapid diversification of the neotropical *Prunus*—the lineages may have already acquired multiple morphological innovations for surviving in the New World’s warm temperate and tropical zones during the Oligocene-early Miocene.

#### The promise of machine learning to assess morphology via digitized herbarium specimens

Machine learning approaches such as IVIS offer the promise of using whole-specimen phenotyping with minimal preprocessing to classify species based on a biologically informed hypothesis. These hypotheses could be based on phylogeny, environmental gradients, ecological features, or geography. We demonstrate a method to use specimen data to investigate correspondence between environmental and morphological variation (Fig. 8). Approaches such as IVIS allow researchers to be agnostic regarding the morphological features studied, which can avoid researcher-meditated bias in selecting traits. By grouping together specimens in different and/or hierarchical groups, competing hypotheses can be tested. We demonstrate how specimens can be grouped together to test hypotheses of hybridization. It is common to use the species as the unit of classification with image-based machine learning algorithms. However, the species is not necessarily the only level of biological organization that can be used to train classifiers. Classifier algorithms that differentiate between species implicitly use a phenetic species concept (de Queiroz 2007), which is not ideal for all research questions. We argue that groups above and below species are useful, especially if they may be defined by shared phenotypic features; here we demonstrate an application at a broader systematic level. Taking a phylogenetic approach to grouping units to be classified, based on key phenetic differences, presents a way to mesh phylogenetic and phenotypic data. Although there are benefits to using whole-specimen phenotypes, there are also advantages to extracting individual traits, such as measurements of leaf area and perimeter, as well as floral traits. These can be extracted via machine learning methods such as segmentation in a high-throughput fashion.

**Figure 8.**
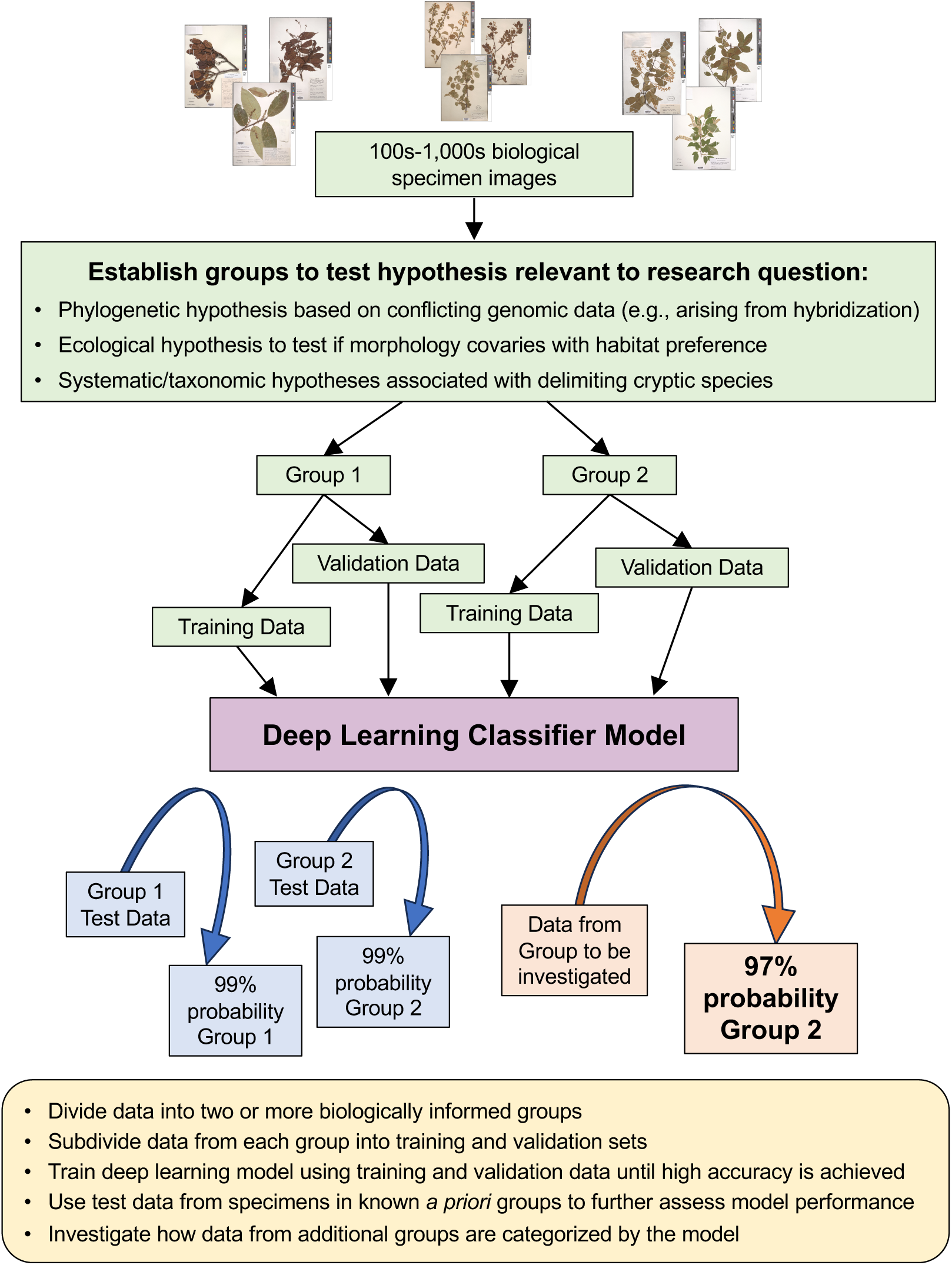
A flow chart for designing phylogenetically-informed hypotheses to test using machine learning applied to biological specimen image data.

#### Conclusions and future prospects

In this study, we employ phylogenomic, environmental, and morphological data to establish that the initial diversification of the plum genus *Prunus* was driven by ancient hybridization and/or allopolyploidy. Moreover, this complex history of reticulation may explain the success of the tropical lineages of *Prunus* compared to other groups that also moved out of the tropics and into temperate regions. To complement phylogenomic and environmental analyses, we present innovative applications of machine learning algorithms to analyze the variation in digitized herbarium sheets. We demonstrate inventive ways to harness computer vision-based machine learning approaches to help us understand biodiversity across the Tree of Life. Future studies in *Prunus* will require additional sampling, especially in the racemose group, to further examine biogeographic patterns. Subsequent research may also focus on developing segmentation masks to extract specific features from digitized image data. *Prunus* represents an excellent model for testing the capabilities of machine learning algorithms using museum data, given the wealth of publicly available digitized specimens in the genus.

## Data Availability

Data available from the Dryad Digital Repository: DOI: 10.5061/dryad.x95x69pwr; Reviewer URL: http://datadryad.org/share/L-QUcxrgnpTr6l0Td3MxdfJDCUT1iRbPmT4SzA2LwMA.

Supplementary material, image data, and DNA sequence matrices are available from the Dryad Digital Repository. Raw sequence data were submitted to NCBI GenBank (SUB13638423). Machine learning models are hosted on Hugging Face with temporary URLs (https://huggingface.co/richiehodel/Prunus_lineage_classifier; https://huggingface.co/richiehodel/Prunus_herbarium_sheet_classifier). Final Hugging Face URLs will be hosted by the Smithsonian Institution upon acceptance.

Jupyter notebooks for developing and processing machine learning models for image data are available on GitHub (https://github.com/richiehodel/machine_learning_Prunus_herbarium_sheets).

## Acknowledgements

We thank the following funding sources: Smithsonian Institution Scholarly Studies Grant to JW, EAZ, RGJH; Smithsonian Institution Peter Buck Fellowship to RGJH; National Museum of Natural History NHRE Award to SKW.

We thank the following people for valuable discussions and assistance with analyses: Lauren Ayers, Louis Cox, Greg Stull, Alicia Talavera.

All or portions of the laboratory and/or computer work were conducted in and with the support of the Laboratories of Analytical Biology facilities of the National Museum of Natural History or its partner labs.

