## Supplemental Figure S1 for "A phylogenomic approach, combined with morphological characters gleaned via machine learning, uncovers the hybrid origin and biogeographic diversification of the plum genus"

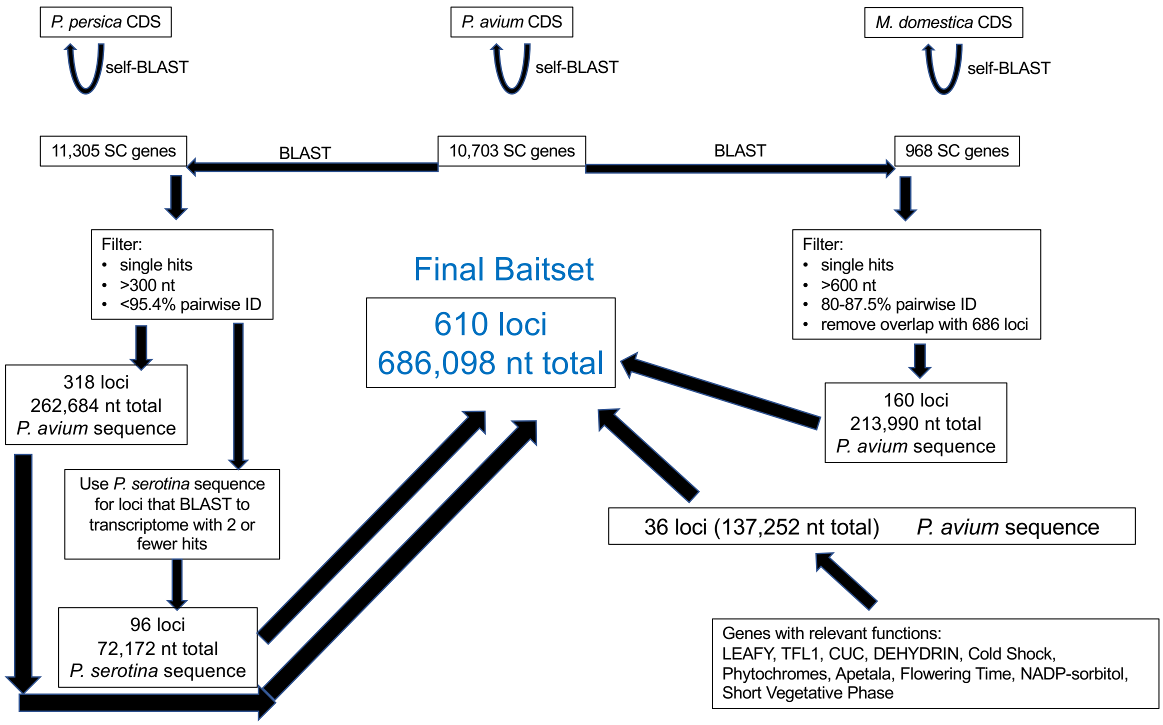


**Supplementary Figure S1**. Flowchart indicating the workflow to design a set of custom Hyb-Seq baits for use in *Prunus*. 610 loci were developed, with 587 retained for phylogenomic analyses after paralogy filtering.
