## Supplemental Figure S2 for "A phylogenomic approach, combined with morphological characters gleaned via machine learning, uncovers the hybrid origin and biogeographic diversification of the plum genus"

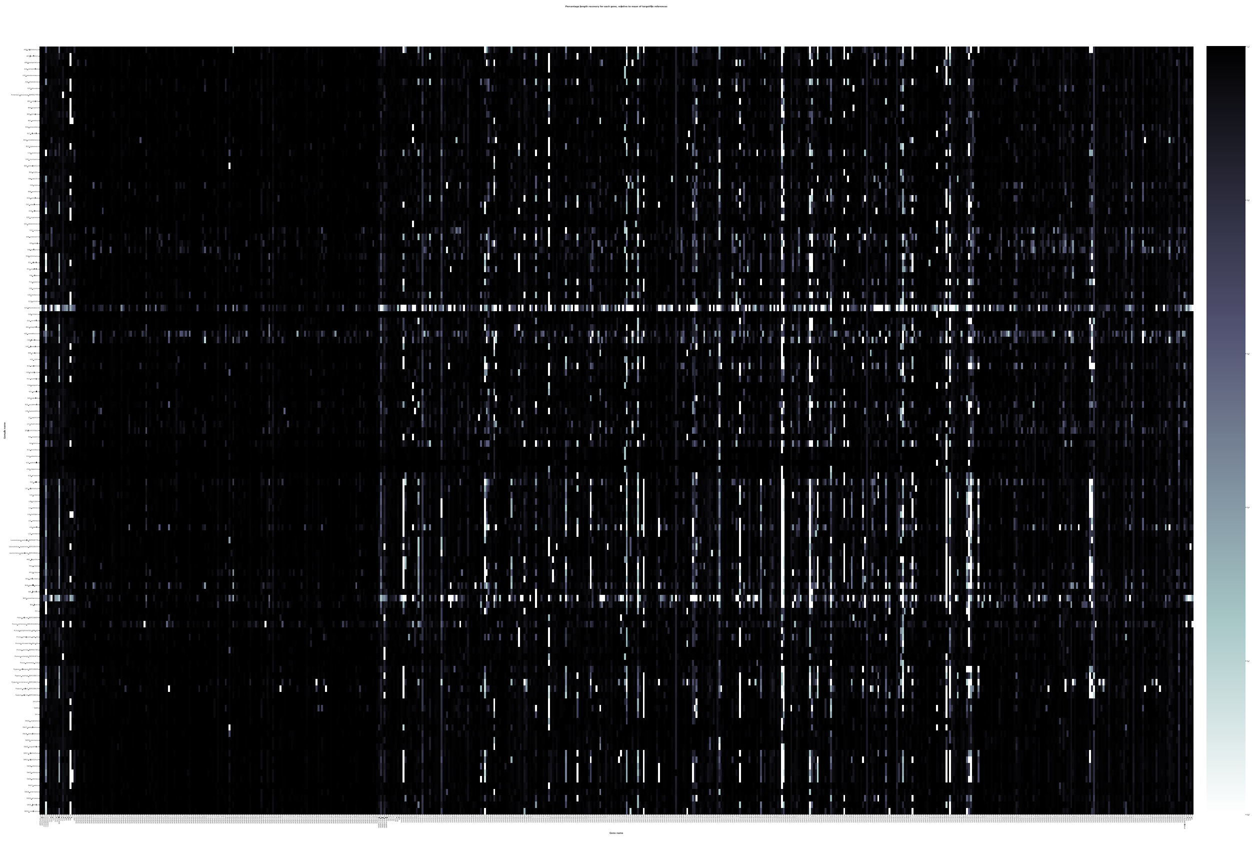

**Supplementary Figure S2**. Heatmap of the 610 genes (X-axis) and 119 accessions (Y-axis). Darker colors indicate that a higher percent of the reference protein length was recovered during the assembly.
