## Supplemental Figure S3 for "A phylogenomic approach, combined with morphological characters gleaned via machine learning, uncovers the hybrid origin and biogeographic diversification of the plum genus"

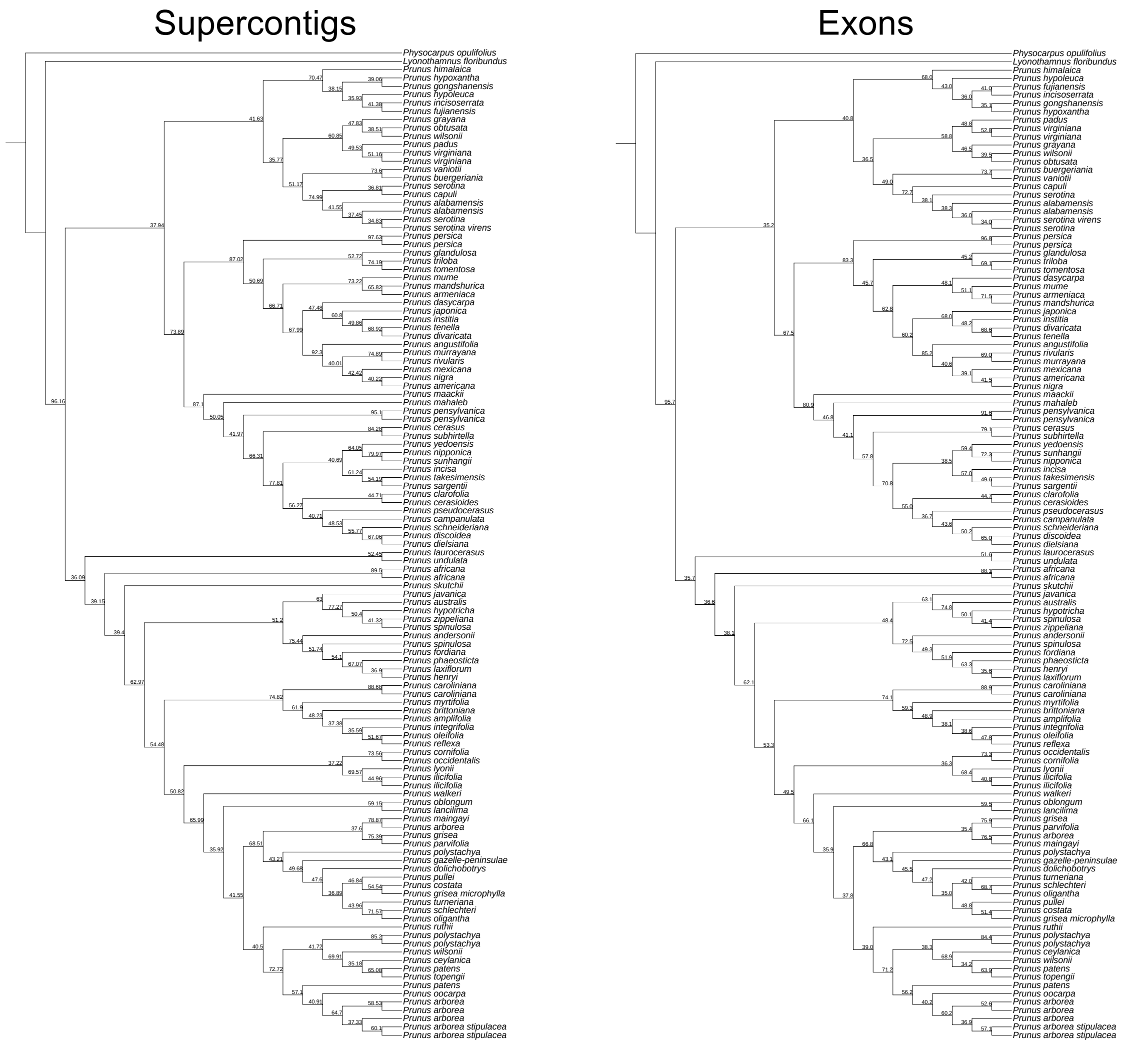


**Supplementary Figure S3**. Species trees generated using ASTRAL to summarize 587 supercontigs (left) and 587 exons (right). The topologies were largely the same and the supercontig dataset was used for subsequent analyses.
