## Supplemental Figure S4 for "A phylogenomic approach, combined with morphological characters gleaned via machine learning, uncovers the hybrid origin and biogeographic diversification of the plum genus"

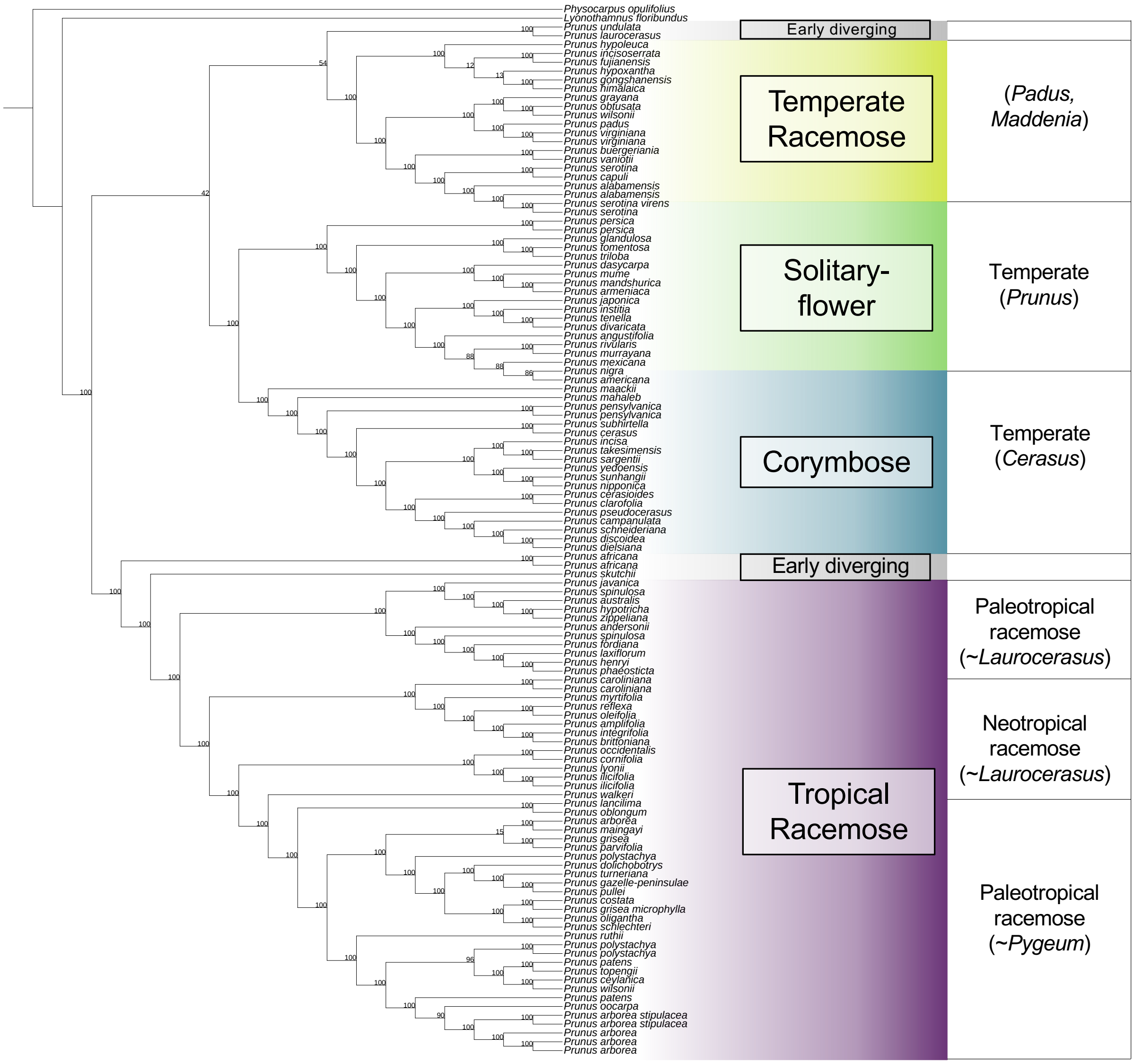


**Supplementary Figure S4**. The nuclear concatenation phylogeny inferred using RAxML on a concatenated supermatrix of the 587 supercontigs. The numbers at nodes indicate bootstrap support values.
