## Supplemental Figure S5 for "A phylogenomic approach, combined with morphological characters gleaned via machine learning, uncovers the hybrid origin and biogeographic diversification of the plum genus"

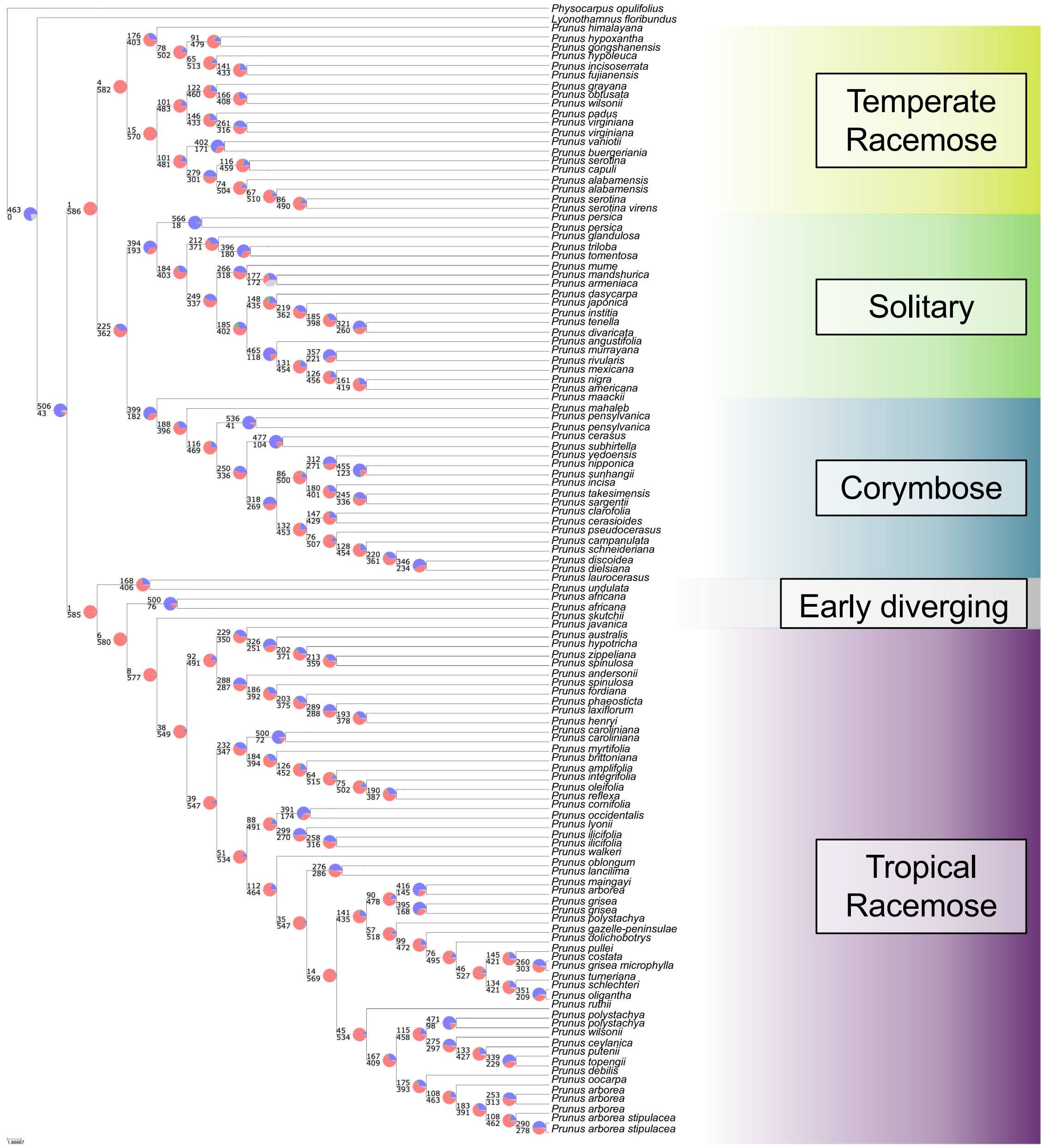


**Supplemental Figure S5.** The *phyparts* phylogeny indicating gene tree-species tree conflict at each node of the nuclear phylogeny. The pie charts at each node indicate the proportion of genes supporting the clade defined by node (blue), the proportion supporting the alternative topology (green), the proportion of other alternatives (red), and uninformative gene trees (defined as less than 50% bootstrap support; gray).
