## Supplemental Figure S6 for "A phylogenomic approach, combined with morphological characters gleaned via machine learning, uncovers the hybrid origin and biogeographic diversification of the plum genus"

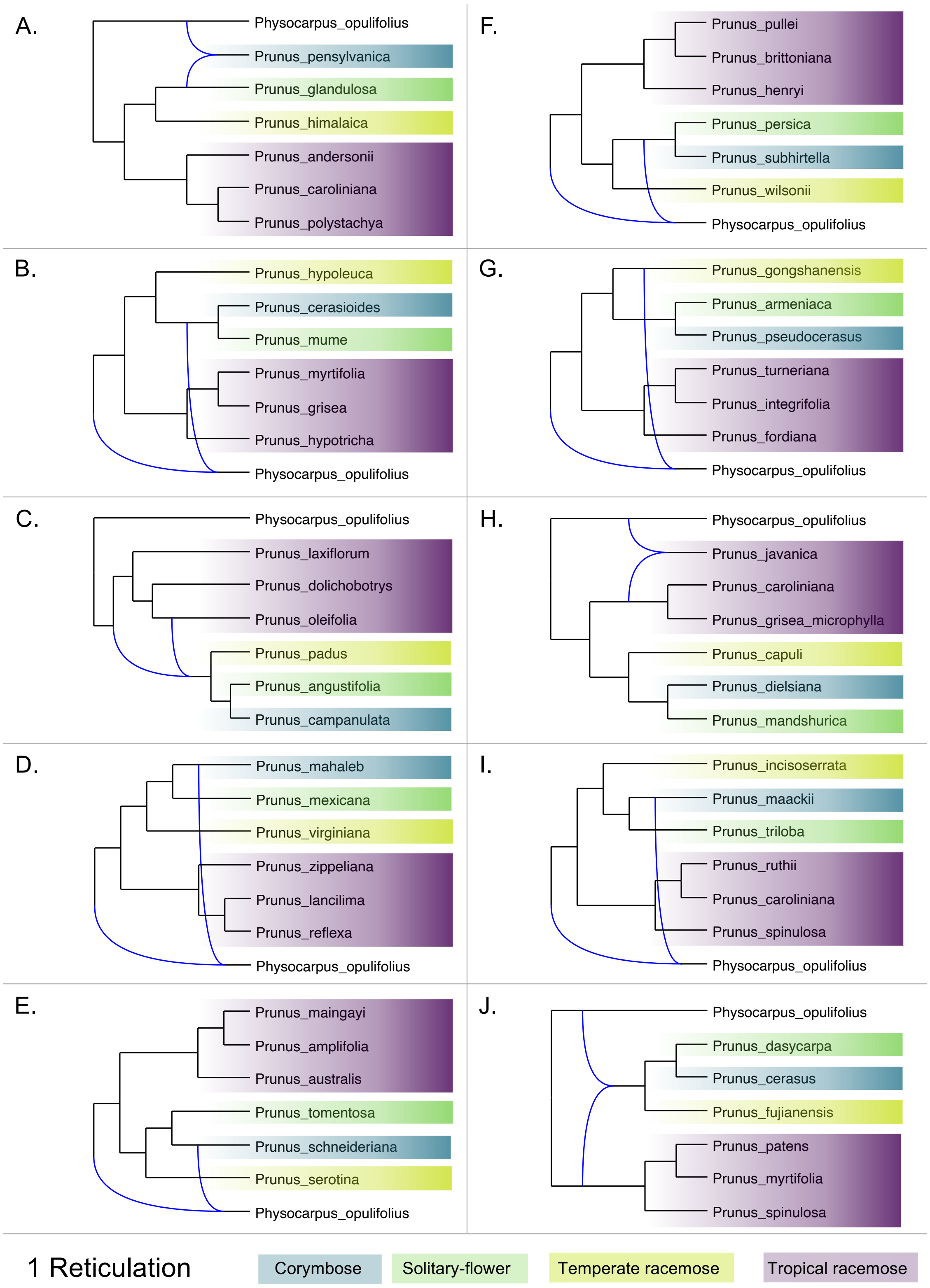

A.
Physocarpus_opulifolius
Prunus_pensylvanica
Prunus_glandulosa
Prunus_himalaica
Prunus_andersonii
Prunus_caroliniana
Prunus_polystachya
F.
Prunus_pullei
Prunus_brittoniana
Prunus_henryi
Prunus_persica
Prunus_subhirtella
Prunus_wilsonii
Physocarpus_opulifolius
B.
Prunus_hypoleuca
Prunus_cerasioides
Prunus_mume
Prunus_myrtifolia
Prunus_grisea
Prunus_hypotricha
Physocarpus_opulifolius
G.
Prunus_gongshanensis
Prunus_armeniaca
Prunus_pseudocerasus
Prunus_turneriana
Prunus_integrifolia
Prunus_fordiana
Physocarpus_opulifolius
C.
Physocarpus_opulifolius
Prunus_laxiflorum
Prunus_dolichobotrys
Prunus_oleifolia
Prunus_padus
Prunus_angustifolia
Prunus_campanulata
H.
Physocarpus_opulifolius
Prunus_javanica
Prunus_caroliniana
Prunus_grisea_microphylla
Prunus_capuli
Prunus_dielsiana
Prunus_mandshurica
D.
Prunus_mahaleb
Prunus_mexicana
Prunus_virginiana
Prunus_zippeliana
Prunus_lancilima
Prunus_reflexa
Physocarpus_opulifolius
I.
Prunus_incisoserrata
Prunus_maackii
Prunus_triloba
Prunus_ruthii
Prunus_caroliniana
Prunus_spinulosa
Physocarpus_opulifolius
E.
Prunus_maingayi
Prunus_amplifolia
Prunus_australis
Prunus_tomentosa
Prunus_schneideriana
Prunus_serotina
Physocarpus_opulifolius
J.
Physocarpus_opulifolius
Prunus_dasycarpa
Prunus_cerasus
Prunus_fujianensis
Prunus_patens
Prunus_myrtifolia
Prunus_spinulosa
1 Reticulation
Corymbose
Solitary-flower
Temperate racemose
Tropical racemose

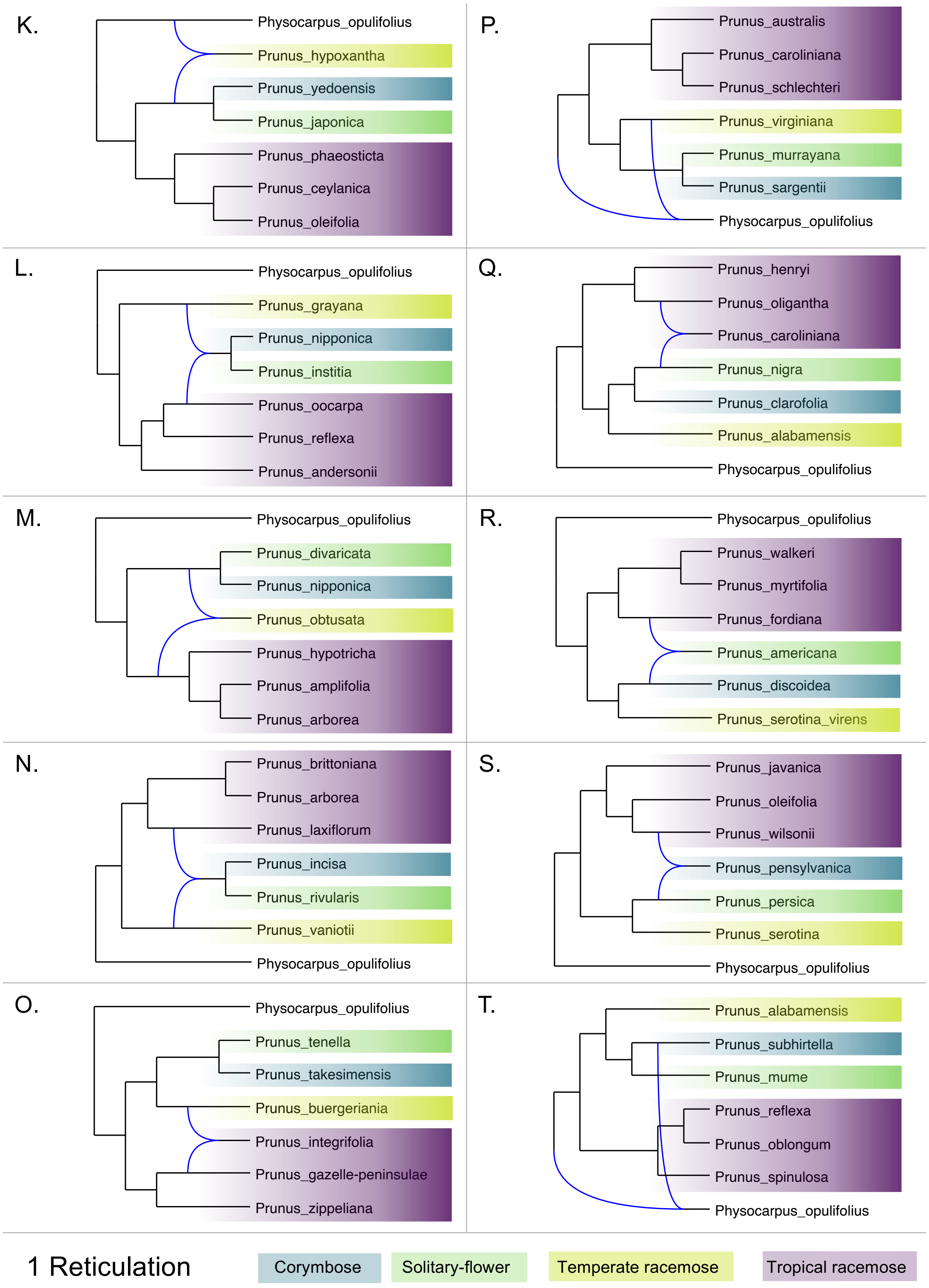
K.
Physocarpus_opulifolius
Prunus_hypoxantha
Prunus_yedoensis
Prunus_japonica
Prunus_phaeosticta
Prunus_ceylanica
Prunus_oleifolia
P.
Prunus_australis
Prunus_caroliniana
Prunus_schlechteri
Prunus_virginiana
Prunus_murrayana
Prunus_sargentii
Physocarpus_opulifolius
L.
Physocarpus_opulifolius
Prunus_grayana
Prunus_nipponica
Prunus_institia
Prunus_oocarpa
Prunus_reflexa
Prunus_andersonii
Q.
Prunus_henryi
Prunus_oligantha
Prunus_caroliniana
Prunus_nigra
Prunus_clarofolia
Prunus_alabamensis
Physocarpus_opulifolius
M.
Physocarpus_opulifolius
Prunus_divaricata
Prunus_nipponica
Prunus_obtusata
Prunus_hypotricha
Prunus_amplifolia
Prunus_arborea
R.
Physocarpus_opulifolius
Prunus_walkeri
Prunus_myrtifolia
Prunus_fordiana
Prunus_americana
Prunus_discoidea
Prunus_serotina_virens
N.
Prunus_brittoniana
Prunus_arborea
Prunus_laxiflorum
Prunus_incisa
Prunus_rivularis
Prunus_vaniotii
Physocarpus_opulifolius
S.
Prunus_javanica
Prunus_oleifolia
Prunus_wilsonii
Prunus_pensylvanica
Prunus_persica
Prunus_serotina
Physocarpus_opulifolius
O.
Physocarpus_opulifolius
Prunus_tenella
Prunus_takesimensis
Prunus_buergeriana
Prunus_integrifolia
Prunus_gazelle-peninsulae
Prunus_zippeliana
T.
Prunus_alabamensis
Prunus_subhirtella
Prunus_mume
Prunus_reflexa
Prunus_oblongum
Prunus_spinulosa
Physocarpus_opulifolius
1 Reticulation
Corymbose
Solitary-flower
Temperate racemose
Tropical racemose

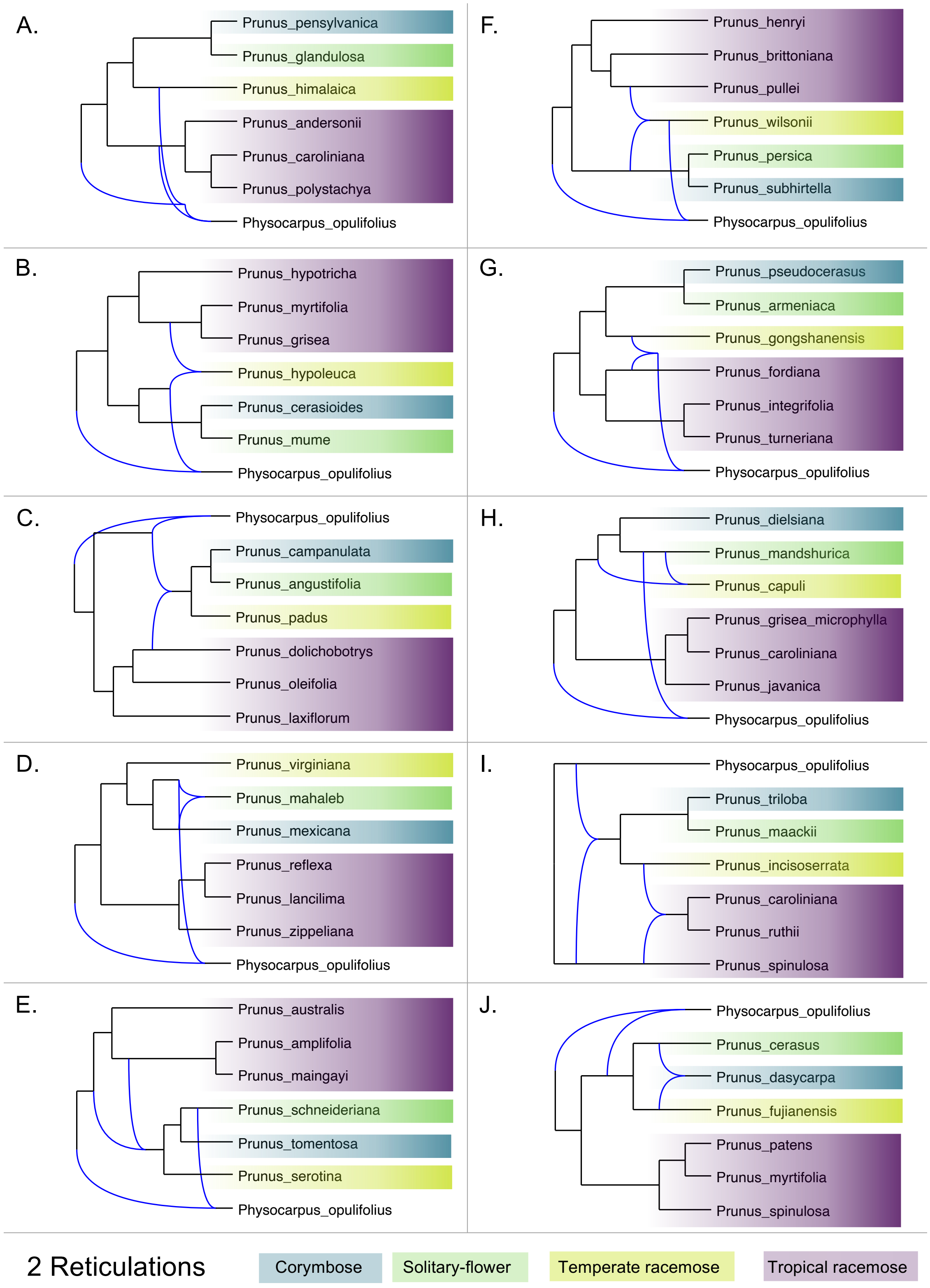
A.
Prunus_pensylvanica
Prunus_glandulosa
Prunus_himalaica
Prunus_andersonii
Prunus_caroliniana
Prunus_polystachya
Physocarpus_opulifolius
F.
Prunus_henryi
Prunus_brittoniana
Prunus_pullei
Prunus_wilsonii
Prunus_persica
Prunus_subhirtella
Physocarpus_opulifolius
B.
Prunus_hypotricha
Prunus_myrtifolia
Prunus_grisea
Prunus_hypoleuca
Prunus_cerasioides
Prunus_mume
Physocarpus_opulifolius
G.
Prunus_pseudocerasus
Prunus_armeniaca
Prunus_gongshanensis
Prunus_fordiana
Prunus_integrifolia
Prunus_turneriana
Physocarpus_opulifolius
C.
Physocarpus_opulifolius
Prunus_campanulata
Prunus_angustifolia
Prunus_padus
Prunus_dolichobotrys
Prunus_oleifolia
Prunus_laxiflorum
H.
Prunus_dielsiana
Prunus_mandshurica
Prunus_capuli
Prunus_grisea_microphylla
Prunus_caroliniana
Prunus_javanica
Physocarpus_opulifolius
D.
Prunus_virginiana
Prunus_mahaleb
Prunus_mexicana
Prunus_reflexa
Prunus_lancilima
Prunus_zippeliana
Physocarpus_opulifolius
I.
Physocarpus_opulifolius
Prunus_triloba
Prunus_maackii
Prunus_incisoserrata
Prunus_caroliniana
Prunus_ruthii
Prunus_spinulosa
E.
Prunus_australis
Prunus_amplifolia
Prunus_maingayi
Prunus_schneideriana
Prunus_tomentosa
Prunus_serotina
Physocarpus_opulifolius
J.
Physocarpus_opulifolius
Prunus_cerasus
Prunus_dasycarpa
Prunus_fujianensis
Prunus_patens
Prunus_myrtifolia
Prunus_spinulosa
2 Reticulations
Corymbose
Solitary-flower
Temperate racemose
Tropical racemose

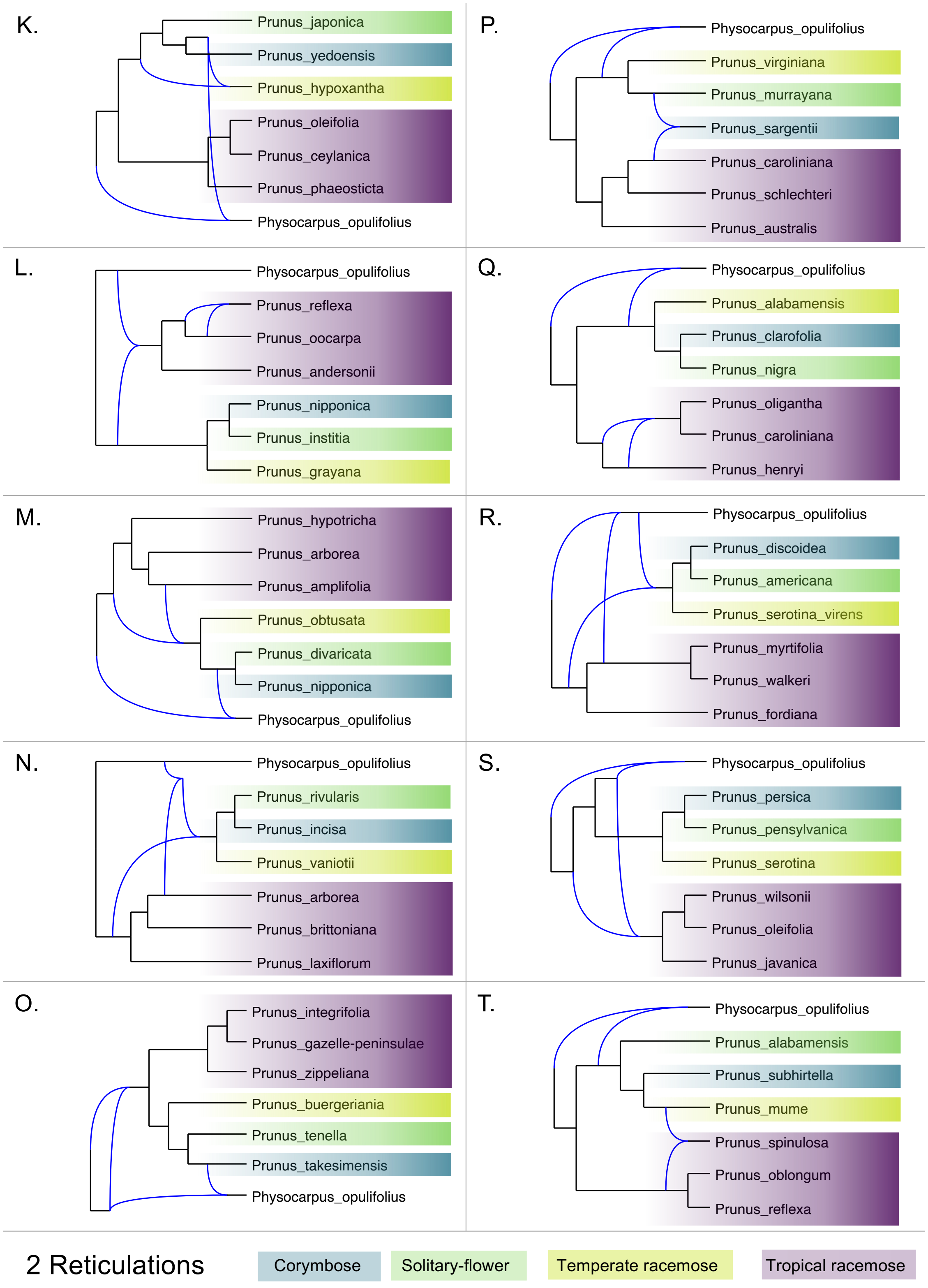
K.
Prunus_japonica
Prunus_yedoensis
Prunus_hypoxantha
Prunus_oleifolia
Prunus_ceylanica
Prunus_phaeosticta
Physocarpus_opulifolius
P.
Physocarpus_opulifolius
Prunus_virginiana
Prunus_murrayana
Prunus_sargentii
Prunus_caroliniana
Prunus_schlechteri
Prunus_australis
L.
Physocarpus_opulifolius
Prunus_reflexa
Prunus_oocarpa
Prunus_andersonii
Prunus_nipponica
Prunus_institia
Prunus_grayana
Q.
Physocarpus_opulifolius
Prunus_alabamensis
Prunus_clarofolia
Prunus_nigra
Prunus_oligantha
Prunus_caroliniana
Prunus_henryi
M.
Prunus_hypotricha
Prunus_arborea
Prunus_amplifolia
Prunus_obtusata
Prunus_divaricata
Prunus_nipponica
Physocarpus_opulifolius
R.
Physocarpus_opulifolius
Prunus_discoidea
Prunus_americana
Prunus_serotina_virens
Prunus_myrtifolia
Prunus_walkeri
Prunus_fordiana
N.
Physocarpus_opulifolius
Prunus_rivularis
Prunus_incisa
Prunus_vaniotii
Prunus_arborea
Prunus_brittoniana
Prunus_laxiflorum
S.
Physocarpus_opulifolius
Prunus_persica
Prunus_pensylvanica
Prunus_serotina
Prunus_wilsonii
Prunus_oleifolia
Prunus_javanica
O.
Prunus_integrifolia
Prunus_gazelle-peninsulae
Prunus_zippeliana
Prunus_buergeriania
Prunus_tenella
Prunus_takesimensis
Physocarpus_opulifolius
T.
Physocarpus_opulifolius
Prunus_alabamensis
Prunus_subhirtella
Prunus_mume
Prunus_spinulosa
Prunus_oblongum
Prunus_reflexa
2 Reticulations
Corymbose
Solitary-flower
Temperate racemose
Tropical racemose

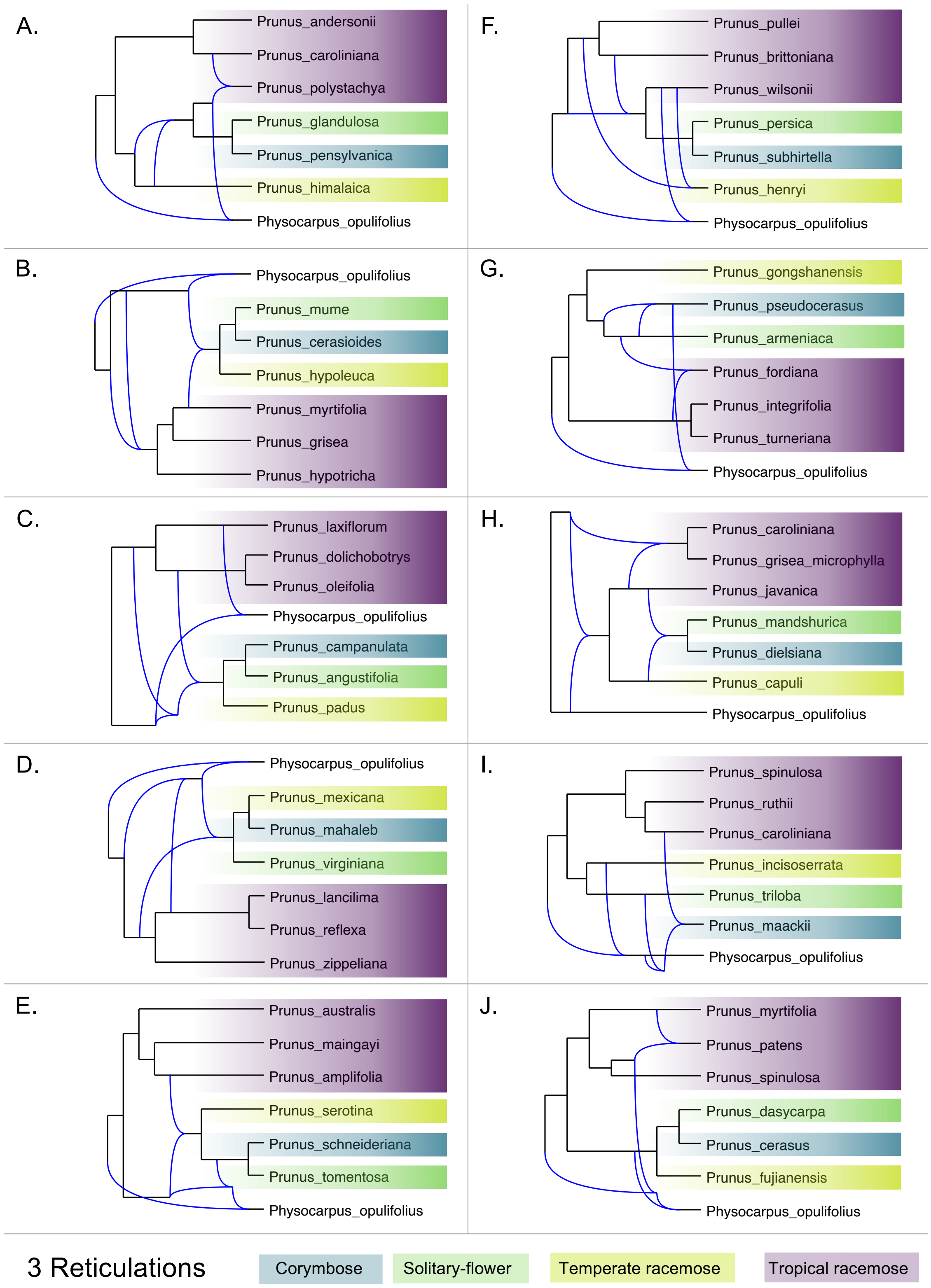

A.
Prunus_andersonii
Prunus_caroliniana
Prunus_polystachya
Prunus_glandulosa
Prunus_pensylvanica
Prunus_himalaica
Physocarpus_opulifolius
F.
Prunus_pullei
Prunus_brittoniana
Prunus_wilsonii
Prunus_persica
Prunus_subhirtella
Prunus_henryi
Physocarpus_opulifolius
B.
Physocarpus_opulifolius
Prunus_mume
Prunus_cerasioides
Prunus_hypoleuca
Prunus_myrtifolia
Prunus_grisea
Prunus_hypotricha
G.
Prunus_gongshanensis
Prunus_pseudocerasus
Prunus_armeniaca
Prunus_fordiana
Prunus_integrifolia
Prunus_turneriana
Physocarpus_opulifolius
C.
Prunus_laxiflorum
Prunus_dolichobotrys
Prunus_oleifolia
Physocarpus_opulifolius
Prunus_campanulata
Prunus_angustifolia
Prunus_padus
H.
Prunus_caroliniana
Prunus_grisea_microphylla
Prunus_javanica
Prunus_mandshurica
Prunus_dielsiana
Prunus_capuli
Physocarpus_opulifolius
D.
Physocarpus_opulifolius
Prunus_mexicana
Prunus_mahaleb
Prunus_virginiana
Prunus_lancilima
Prunus_reflexa
Prunus_zippeliana
I.
Prunus_spinulosa
Prunus_ruthii
Prunus_caroliniana
Prunus_incisoserrata
Prunus_triloba
Prunus_maackii
Physocarpus_opulifolius
E.
Prunus_australis
Prunus_maingayi
Prunus_amplifolia
Prunus_serotina
Prunus_schneideriana
Prunus_tomentosa
Physocarpus_opulifolius
J.
Prunus_myrtifolia
Prunus_patens
Prunus_spinulosa
Prunus_dasycarpa
Prunus_cerasus
Prunus_fujianensis
Physocarpus_opulifolius
3 Reticulations
Corymbose
Solitary-flower
Temperate racemose
Tropical racemose

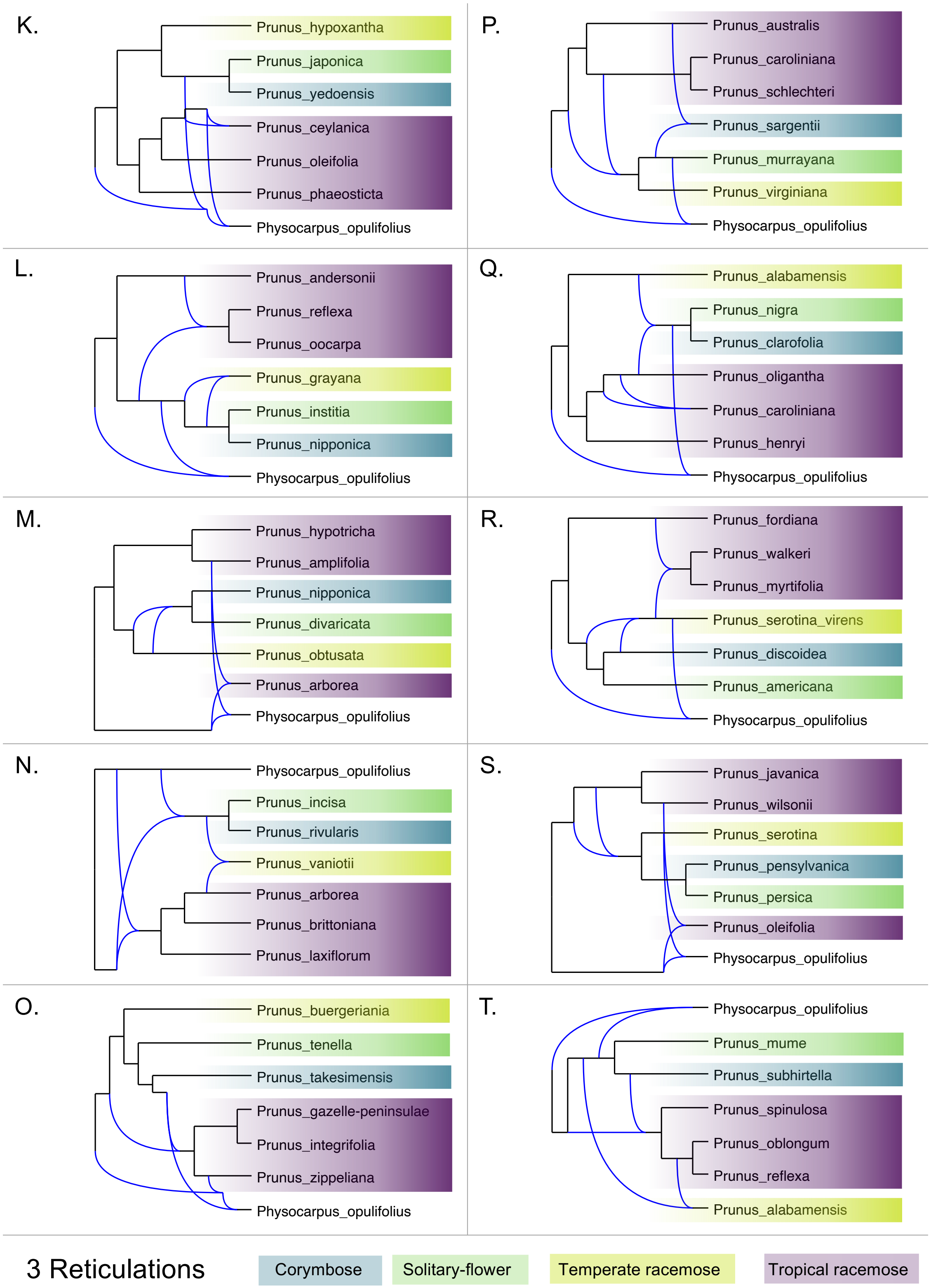
K.
Prunus_hypoxantha
Prunus_japonica
Prunus_yedoensis
Prunus_ceylanica
Prunus_oleifolia
Prunus_phaeosticta
Physocarpus_opulifolius
P.
Prunus_australis
Prunus_caroliniana
Prunus_schlechteri
Prunus_sargentii
Prunus_murrayana
Prunus_virginiana
Physocarpus_opulifolius
L.
Prunus_andersonii
Prunus_reflexa
Prunus_oocarpa
Prunus_grayana
Prunus_institia
Prunus_nipponica
Physocarpus_opulifolius
Q.
Prunus_alabamensis
Prunus_nigra
Prunus_clarofolia
Prunus_oligantha
Prunus_caroliniana
Prunus_henryi
Physocarpus_opulifolius
M.
Prunus_hypotricha
Prunus_amplifolia
Prunus_nipponica
Prunus_divaricata
Prunus_obtusata
Prunus_arborea
Physocarpus_opulifolius
R.
Prunus_fordiana
Prunus_walkeri
Prunus_myrtifolia
Prunus_serotina_virens
Prunus_discoidea
Prunus_americana
Physocarpus_opulifolius
N.
Physocarpus_opulifolius
Prunus_incisa
Prunus_rivularis
Prunus_vaniotii
Prunus_arborea
Prunus_brittoniana
Prunus_laxiflorum
S.
Prunus_javanica
Prunus_wilsonii
Prunus_serotina
Prunus_pensylvanica
Prunus_persica
Prunus_oleifolia
Physocarpus_opulifolius
O.
Prunus_buergeriania
Prunus_tenella
Prunus_takesimensis
Prunus_gazelle-peninsulae
Prunus_integrifolia
Prunus_zippeliana
Physocarpus_opulifolius
T.
Physocarpus_opulifolius
Prunus_mume
Prunus_subhirtella
Prunus_spinulosa
Prunus_oblongum
Prunus_reflexa
Prunus_alabamensis
3 Reticulations
Corymbose
Solitary-flower
Temperate racemose
Tropical racemose

**Supplemental Figure S6.** The phylonet networks using multiple samplings of seven species representing every major lineage in *Prunus*, including an outgroup (*Physocarpus opulifolius*). For 1-, 2-, and 3-reticulation networks, the network with the optimal pseudolikelihood score for each of the 20 taxon samplings is shown.
