## Supplemental Figure S7 for "A phylogenomic approach, combined with morphological characters gleaned via machine learning, uncovers the hybrid origin and biogeographic diversification of the plum genus"

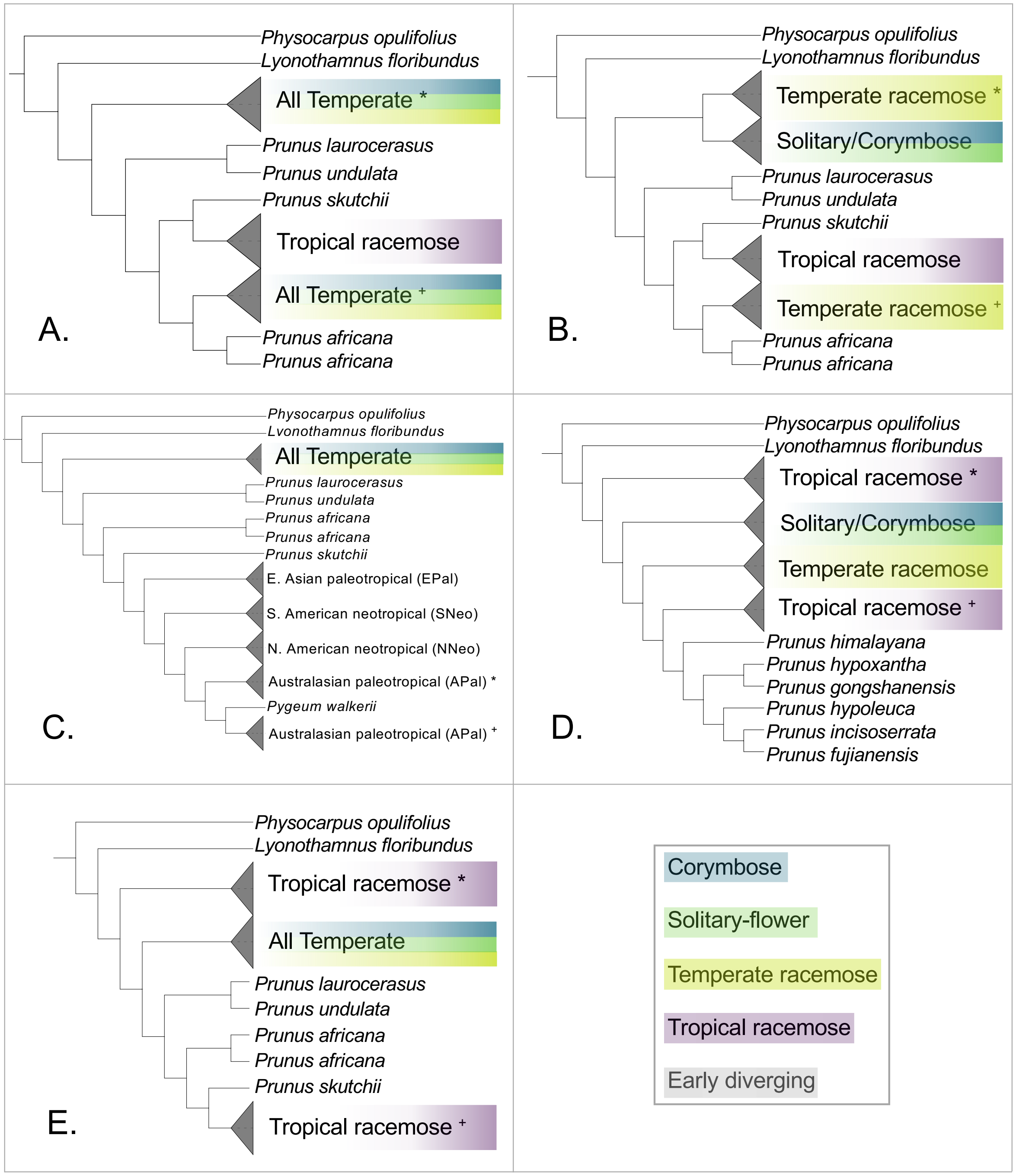


**Supplemental Figure S7.** For five key clades in the genus *Prunus*, the most parsimonious MUL-tree from the GRAMPA analysis, with duplicated clades noted by asterisks and plus symbols. A) the temperate clade was identified as having an ancient duplication event; B) the temperate racemose clade was the result of allopolyploidy; C) the Australasian clade is a product of gene duplication, although the two duplicated clades plus *Pygeum walkerii* form a clade; D) the tropical racemose clade, including early diverging species, is the result of genome duplication; E) the tropical racemose clade, excluding the early diverging species, is duplicated.
