## Supplemental Figure S8 for "A phylogenomic approach, combined with morphological characters gleaned via machine learning, uncovers the hybrid origin and biogeographic diversification of the plum genus"

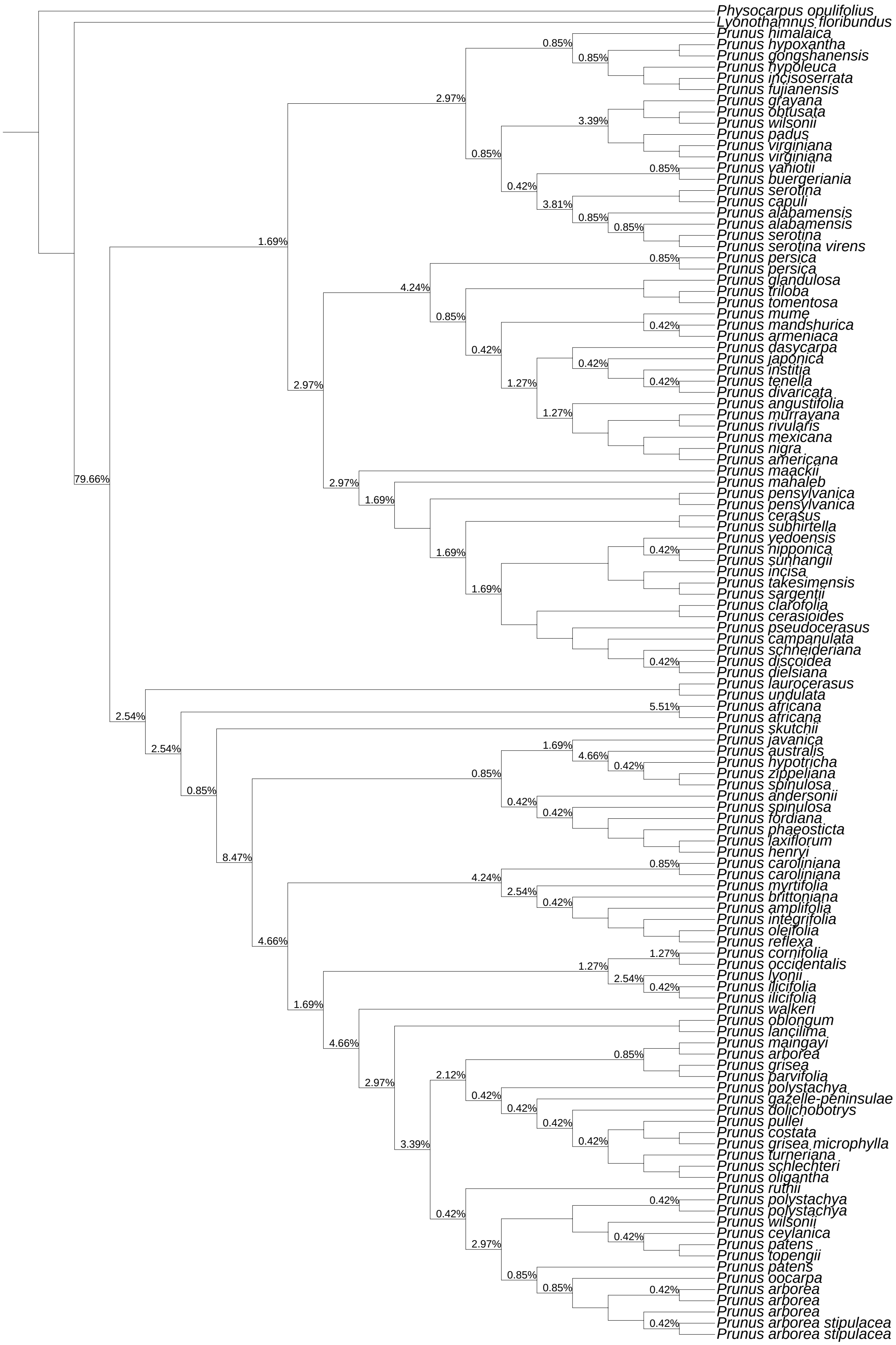

Physocarpus opulifolius
Lyonothamnus floribundus
Prunus himalaica
Prunus hypoxantha
Prunus gongshanensis
Prunus hypoleuca
Prunus incisoserrata
Prunus fujianensis
Prunus grayana
Prunus obtusata
Prunus wilsonii
Prunus padus
Prunus virginiana
Prunus virginiana
Prunus vaniotii
Prunus buergeriania
Prunus serotina
Prunus capuli
Prunus alabamensis
Prunus alabamensis
Prunus serotina
Prunus serotina virens
Prunus persica
Prunus persica
Prunus glandulosa
Prunus triloba
Prunus tomentosa
Prunus mume
Prunus mandshurica
Prunus armeniaca
Prunus dasycarpa
Prunus japonica
Prunus institia
Prunus tenella
Prunus divaricata
Prunus angustifolia
Prunus murrayana
Prunus rivularis
Prunus mexicana
Prunus nigra
Prunus americana
Prunus maackii
Prunus mahaleb
Prunus pensylvanica
Prunus pensylvanica
Prunus cerasus
Prunus subhirtella
Prunus yedoensis
Prunus nipponica
Prunus sunhangii
Prunus incisa
Prunus takesimensis
Prunus sargentii
Prunus clarofolia
Prunus cerasioides
Prunus pseudocerasus
Prunus campanulata
Prunus schneideriana
Prunus discoidea
Prunus dielsiana
Prunus laurocerasus
Prunus undulata
Prunus africana
Prunus africana
Prunus skutchii
Prunus javanica
Prunus australis
Prunus hypotricha
Prunus zippeliana
Prunus spinulosa
Prunus andersonii
Prunus spinulosa
Prunus fordiana
Prunus phaeosticta
Prunus laxiflorum
Prunus henryi
Prunus caroliniana
Prunus caroliniana
Prunus myrtifolia
Prunus brittoniana
Prunus amplifolia
Prunus integrifolia
Prunus oleifolia
Prunus reflexa
Prunus cornifolia
Prunus occidentalis
Prunus lyonii
Prunus ilicifolia
Prunus ilicifolia
Prunus walkeri
Prunus oblongum
Prunus lancilima
Prunus maingayi
Prunus arborea
Prunus grisea
Prunus parvifolia
Prunus polystachya
Prunus gazelle-peninsulae
Prunus dolichobotrys
Prunus pullei
Prunus costata
Prunus grisea microphylla
Prunus turneriana
Prunus schlechteri
Prunus oligantha
Prunus ruthii
Prunus polystachya
Prunus polystachya
Prunus wilsonii
Prunus ceylanica
Prunus patens
Prunus topengii
Prunus patens
Prunus oocarpa
Prunus arborea
Prunus arborea
Prunus arborea
Prunus arborea stipulacea
Prunus arborea stipulacea

**Supplemental Figure S8.** The ASTRAL species tree phylogeny with duplications mapped to each node of the species tree. The values at nodes indicate the percentages of subclades descended from that node with multiple tips per species.
