## Supplemental Figure S9 for "A phylogenomic approach, combined with morphological characters gleaned via machine learning, uncovers the hybrid origin and biogeographic diversification of the plum genus"

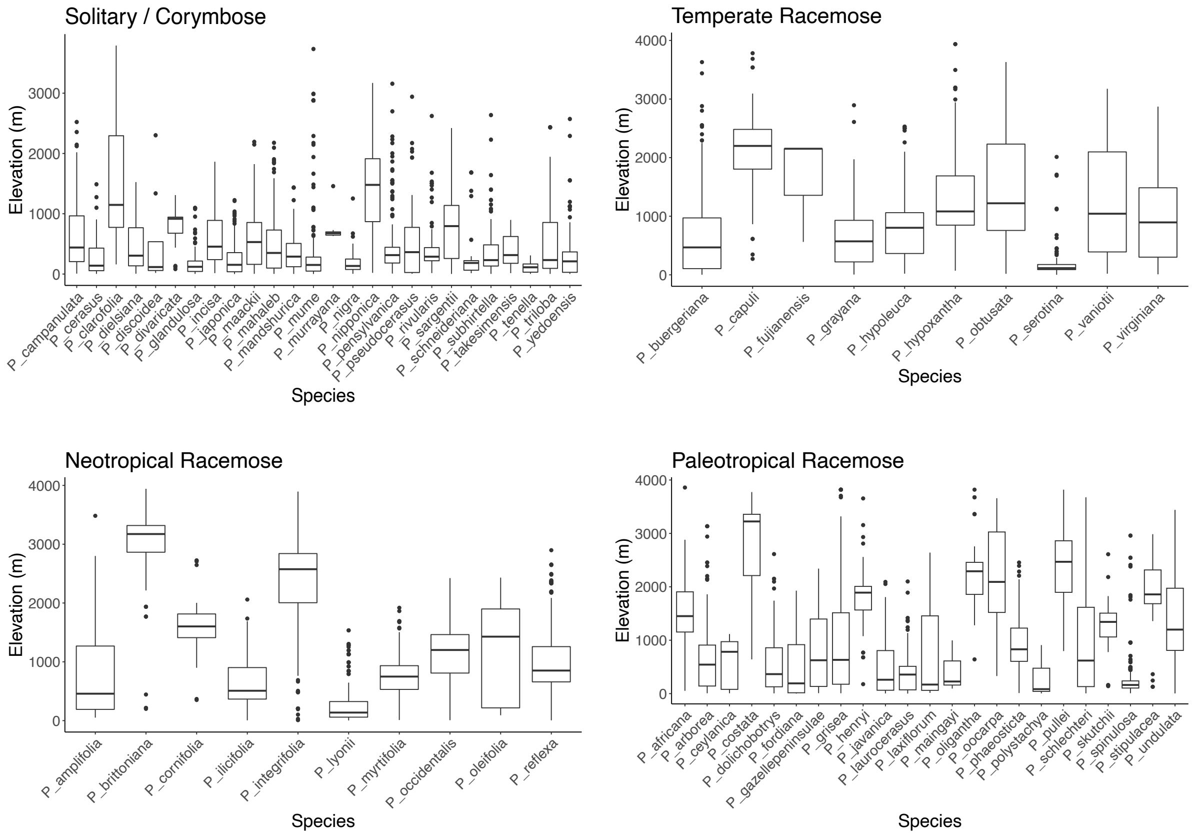


**Supplemental Figure S9.** The range of elevations across all specimens used in morphological analyses for each species.
