## Supplemental Figure S10 for "A phylogenomic approach, combined with morphological characters gleaned via machine learning, uncovers the hybrid origin and biogeographic diversification of the plum genus"

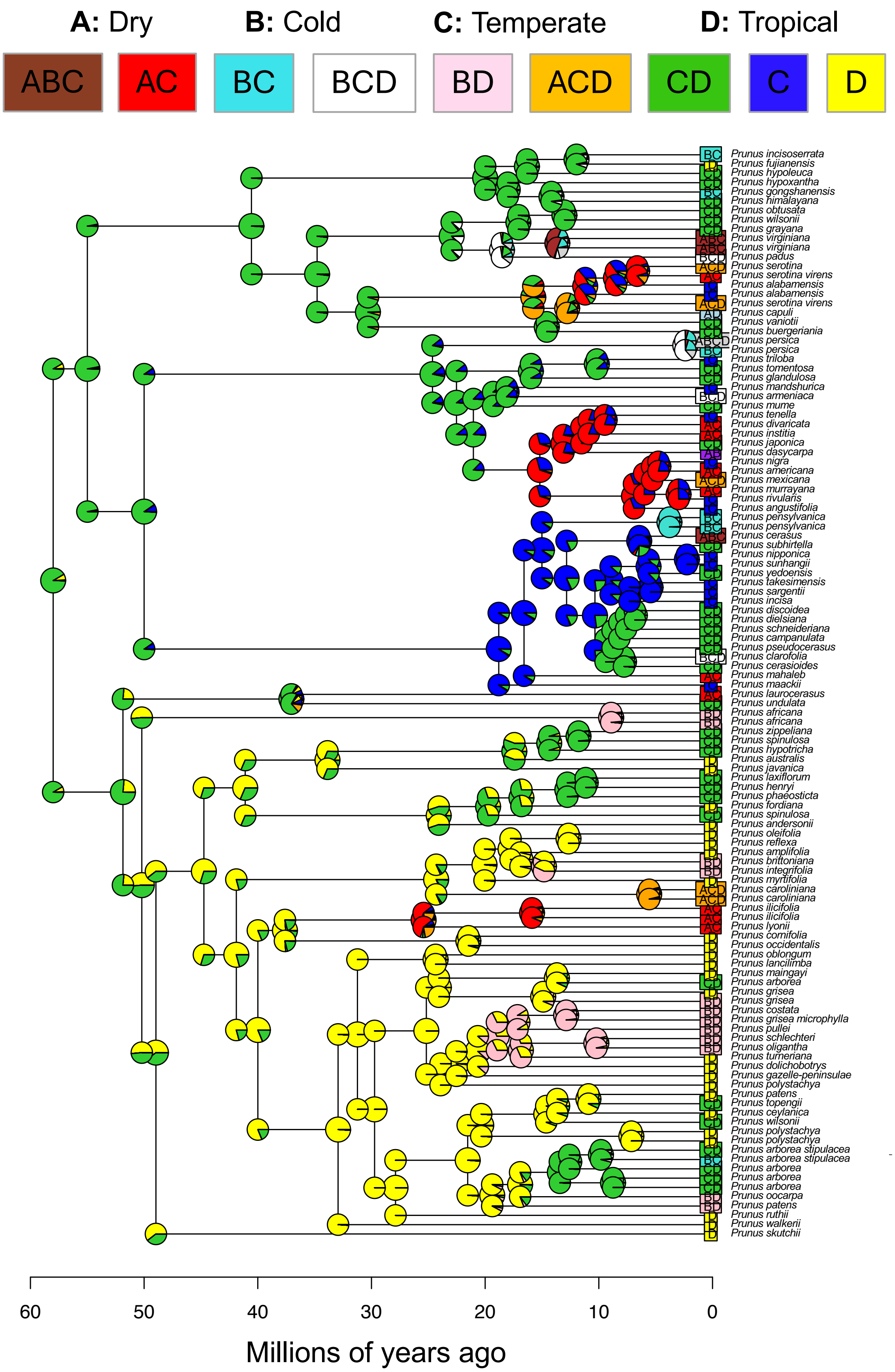

A: Dry
B: Cold
C: Temperate
D: Tropical
ABC
AC
BC
BCD
BD
ACD
CD
C
D
Prunus incisoserrata
Prunus fujianensis
Prunus hypoleuca
Prunus hypoxantha
Prunus gongshanensis
Prunus himalayana
Prunus obtusata
Prunus wilsonii
Prunus grayana
Prunus virginiana
Prunus virginiana
Prunus padus
Prunus serotina
Prunus serotina virens
Prunus alabamensis
Prunus alabamensis
Prunus serotina virens
Prunus capuli
Prunus vaniotii
Prunus buergeriania
Prunus persica
Prunus persica
Prunus triloba
Prunus tomentosa
Prunus glandulosa
Prunus mandshurica
Prunus armeniaca
Prunus mume
Prunus tenella
Prunus divaricata
Prunus institia
Prunus japonica
Prunus dasycarpa
Prunus nigra
Prunus americana
Prunus mexicana
Prunus murrayana
Prunus rivularis
Prunus angustifolia
Prunus pensylvanica
Prunus pensylvanica
Prunus cerasus
Prunus subhirtella
Prunus nipponica
Prunus sunhangii
Prunus yedoensis
Prunus takesimensis
Prunus sargentii
Prunus incisa
Prunus discoidea
Prunus dielsiana
Prunus schneideriana
Prunus campanulata
Prunus pseudocerasus
Prunus clarofolia
Prunus cerasioides
Prunus mahaleb
Prunus maackii
Prunus laurocerasus
Prunus undulata
Prunus africana
Prunus africana
Prunus zippeliana
Prunus spinulosa
Prunus hypotricha
Prunus australis
Prunus javanica
Prunus laxiflorum
Prunus henryi
Prunus phaeosticta
Prunus fordiana
Prunus spinulosa
Prunus andersonii
Prunus oleifolia
Prunus reflexa
Prunus amplifolia
Prunus brittoniana
Prunus integrifolia
Prunus myrtifolia
Prunus caroliniana
Prunus caroliniana
Prunus ilicifolia
Prunus ilicifolia
Prunus lyonii
Prunus cornifolia
Prunus occidentalis
Prunus oblongum
Prunus lancilimba
Prunus maingayi
Prunus arborea
Prunus grisea
Prunus grisea
Prunus costata
Prunus grisea microphylla
Prunus pullei
Prunus schlechteri
Prunus oligantha
Prunus turneriana
Prunus dolichobotrys
Prunus gazelle-peninsulae
Prunus polystachya
Prunus patens
Prunus topengii
Prunus ceylanica
Prunus wilsonii
Prunus polystachya
Prunus polystachya
Prunus arborea stipulacea
Prunus arborea stipulacea
Prunus arborea
Prunus arborea
Prunus arborea
Prunus oocarpa
Prunus patens
Prunus ruthii
Prunus walkerii
Prunus skutchii
60
50
40
30
20
10
0
Millions of years ago

**Supplemental Figure S10.** The BioGeoBears analysis of the biogeographic history of *Prunus* using the BAYAREALIKE model based on four biome-based color-coded biogeographic regions (top). Color-coded pie charts at each node indicate the probability of the ancestral state occurring in a given biogeographic region.
