## Supplemental Figure S11 for "A phylogenomic approach, combined with morphological characters gleaned via machine learning, uncovers the hybrid origin and biogeographic diversification of the plum genus"

**Supplemental Figure S11.** The 100 replicates of biogeographic stochastic mapping analysis in BioGeoBears. The first 50 are from the DEC+J geography-based biogeographic regions analysis. The next 50 correspond to the BAYAREALIKE analysis from the biome-based biogeographic regions analysis. The color changes along branches indicate anagenetic dispersal, whereas shifts at nodes would correspond to shifts at cladogenesis.

**DEC−J_M3_timestrat − Stochastic Map #1/50**
**ancstates: global optim, 2 areas max. d=0.0016; e=0; j=0.0148; LnL=−173.29**

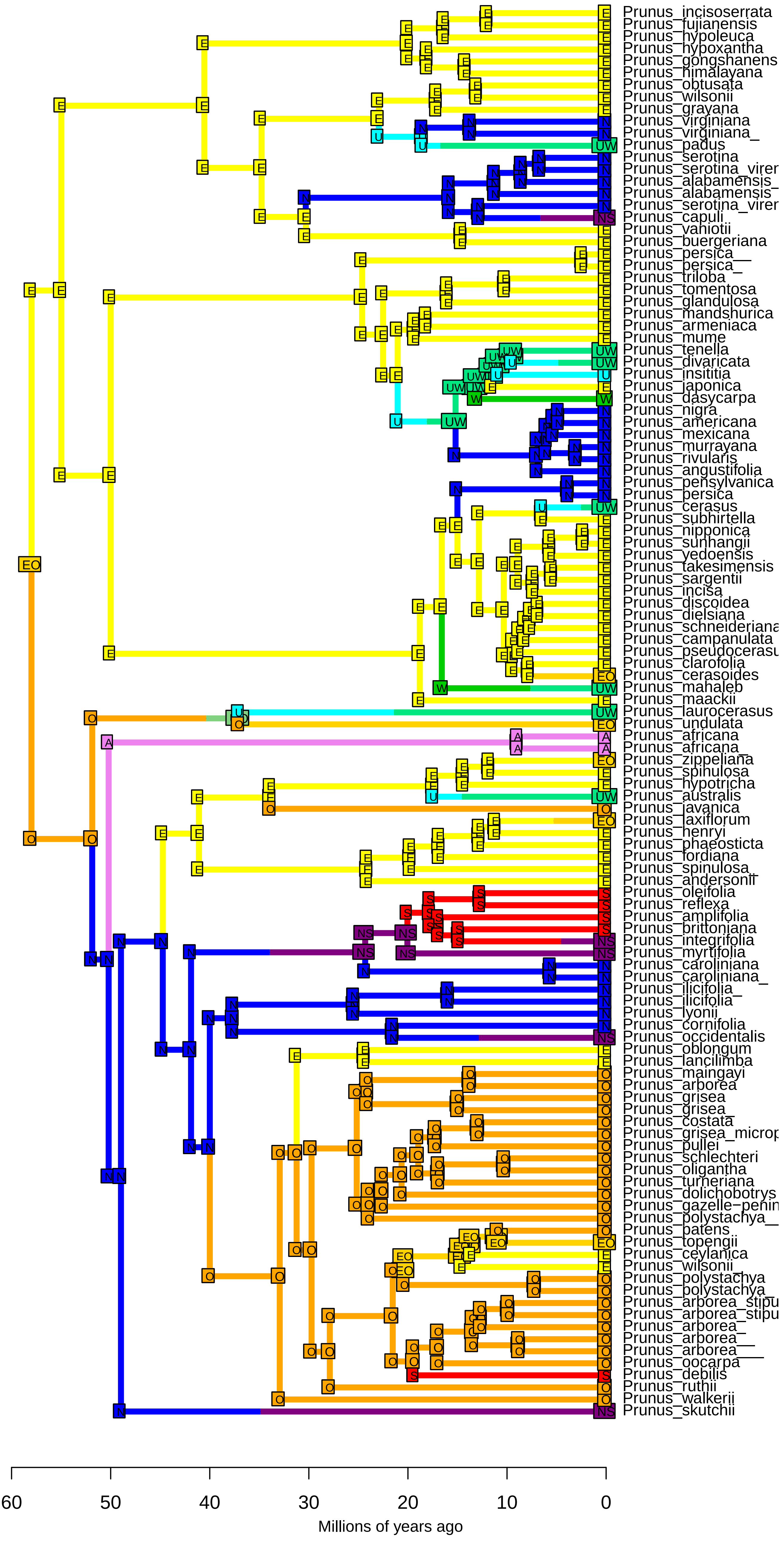

**DEC−J_M3_timestrat − Stochastic Map #2/50**
**ancstates: global optim, 2 areas max. d=0.0016; e=0; j=0.0148; LnL=−173.29**

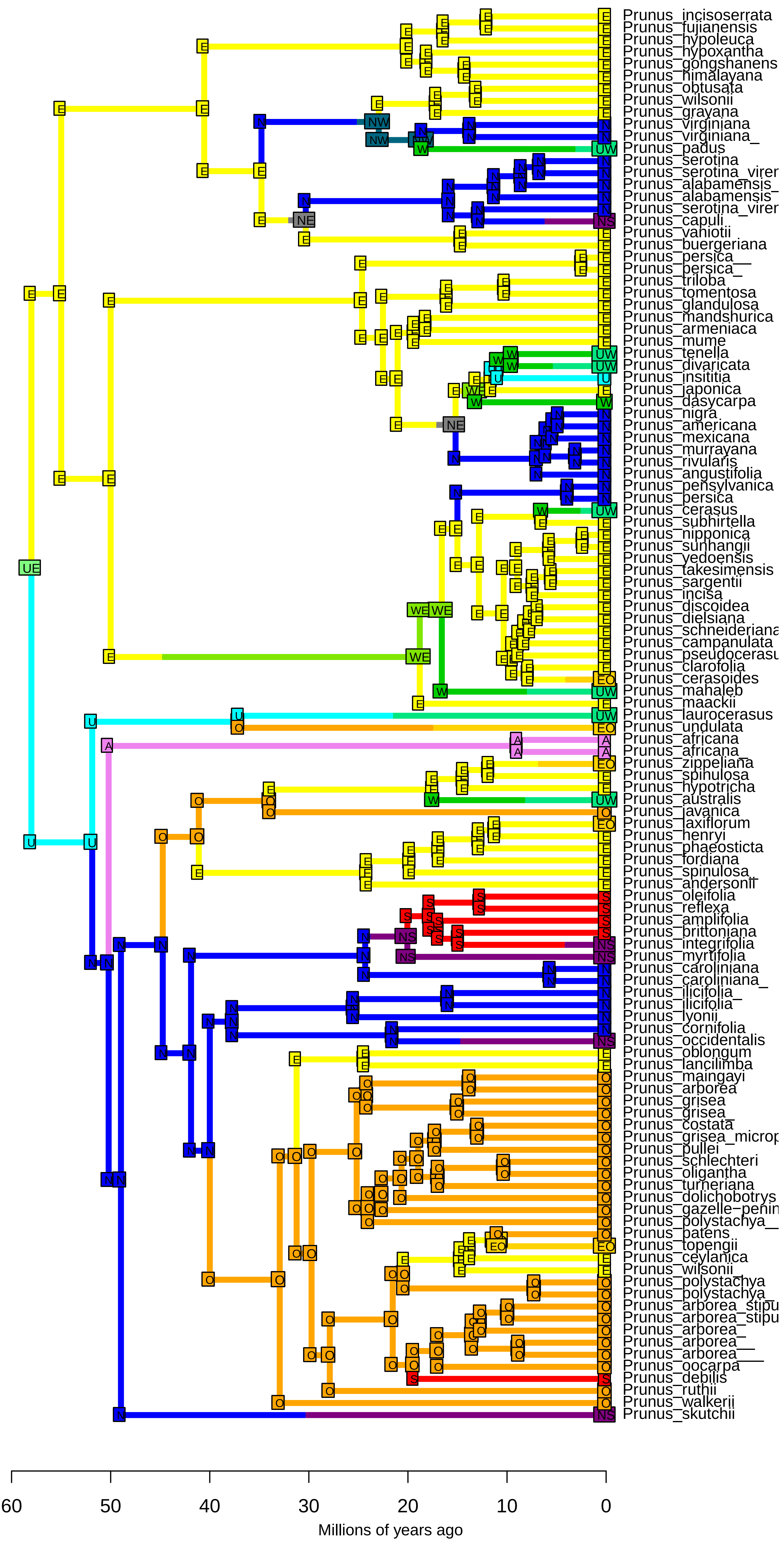

**DEC−J_M3_timestrat − Stochastic Map #3/50**
**ancstates: global optim, 2 areas max. d=0.0016; e=0; j=0.0148; LnL=−173.29**

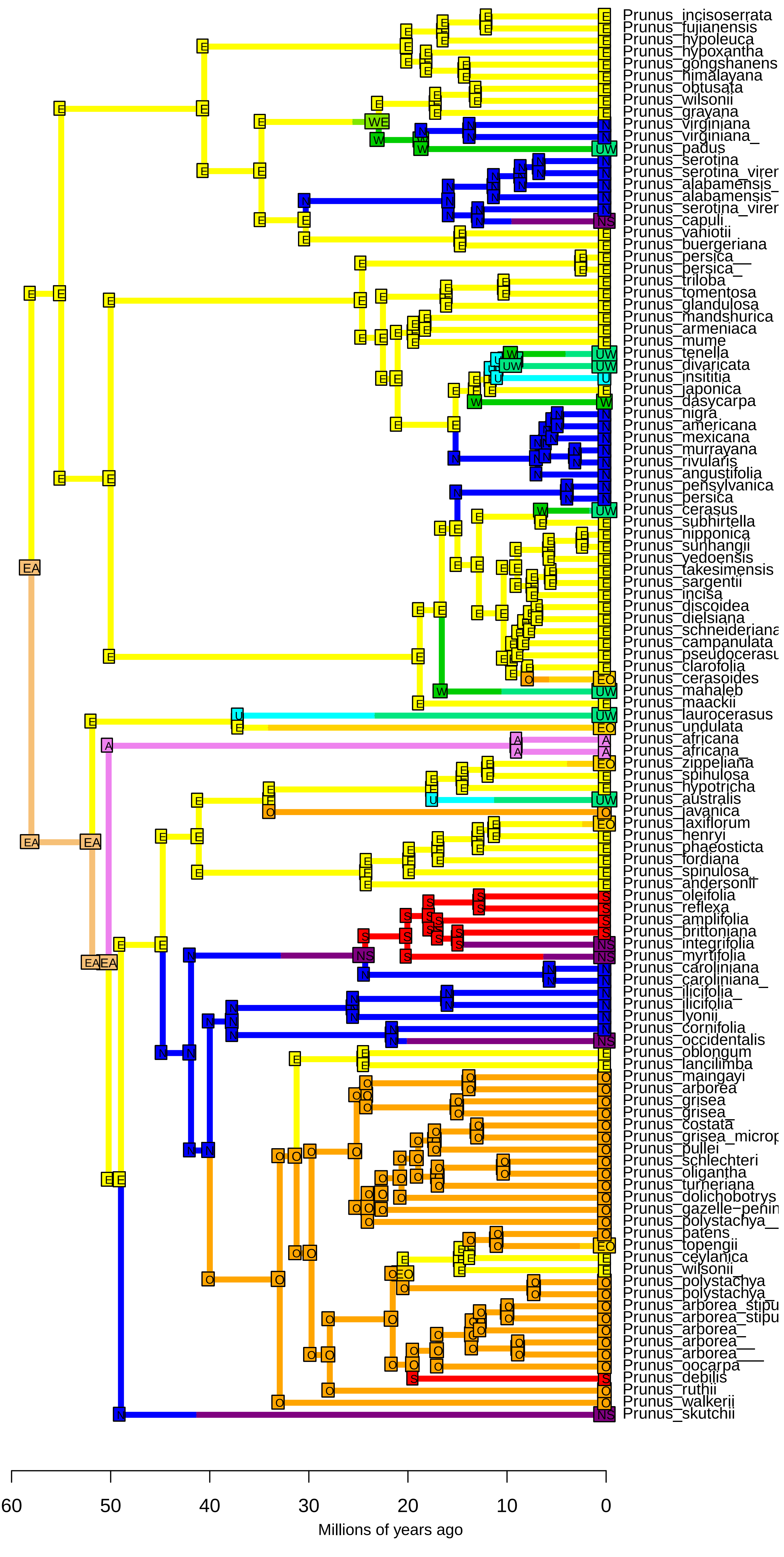

**DEC−J_M3_timestrat − Stochastic Map #4/50**
**ancstates: global optim, 2 areas max. d=0.0016; e=0; j=0.0148; LnL=−173.29**

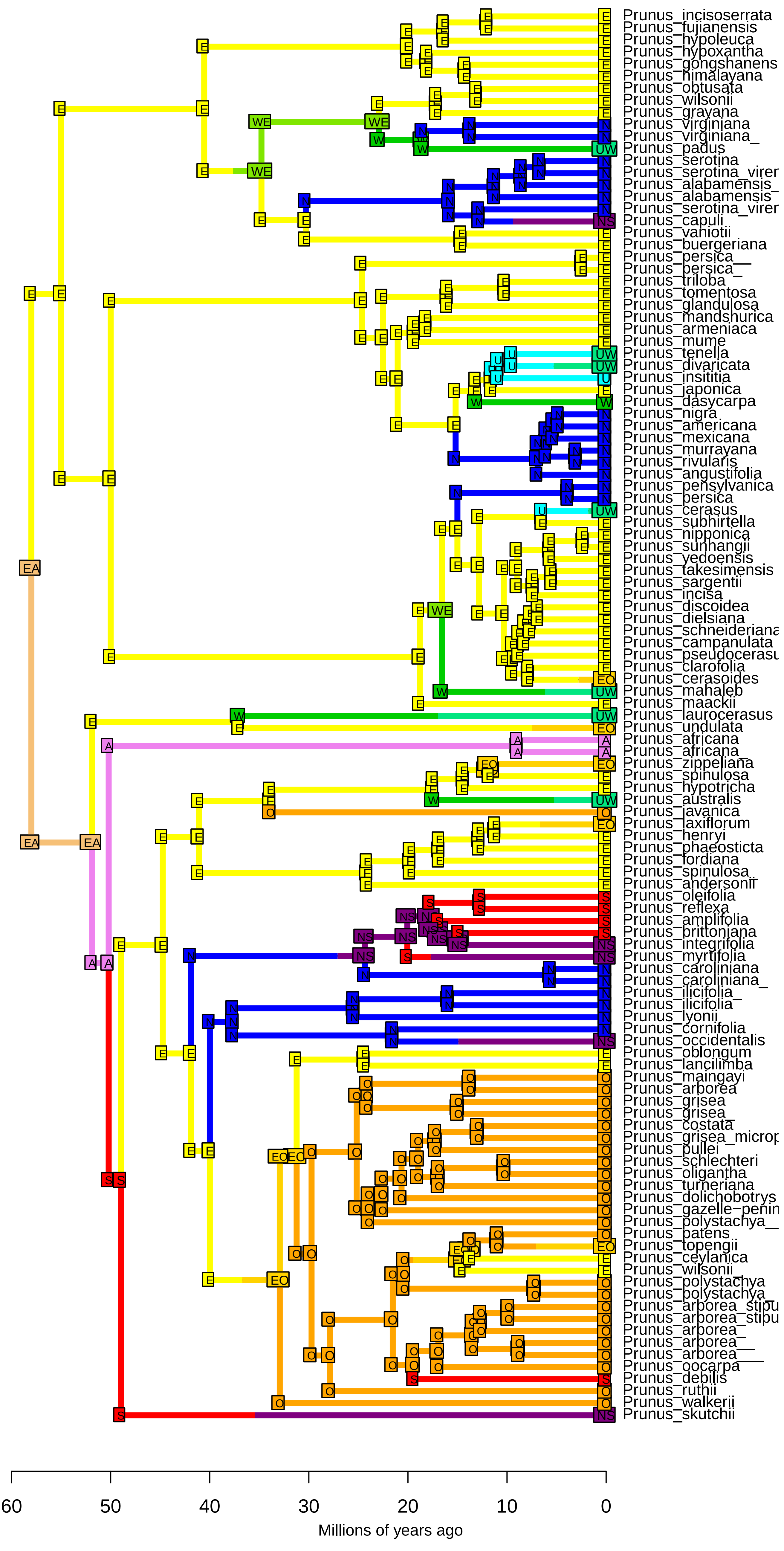

**DEC−J_M3_timestrat − Stochastic Map #5/50**
**ancstates: global optim, 2 areas max. d=0.0016; e=0; j=0.0148; LnL=−173.29**

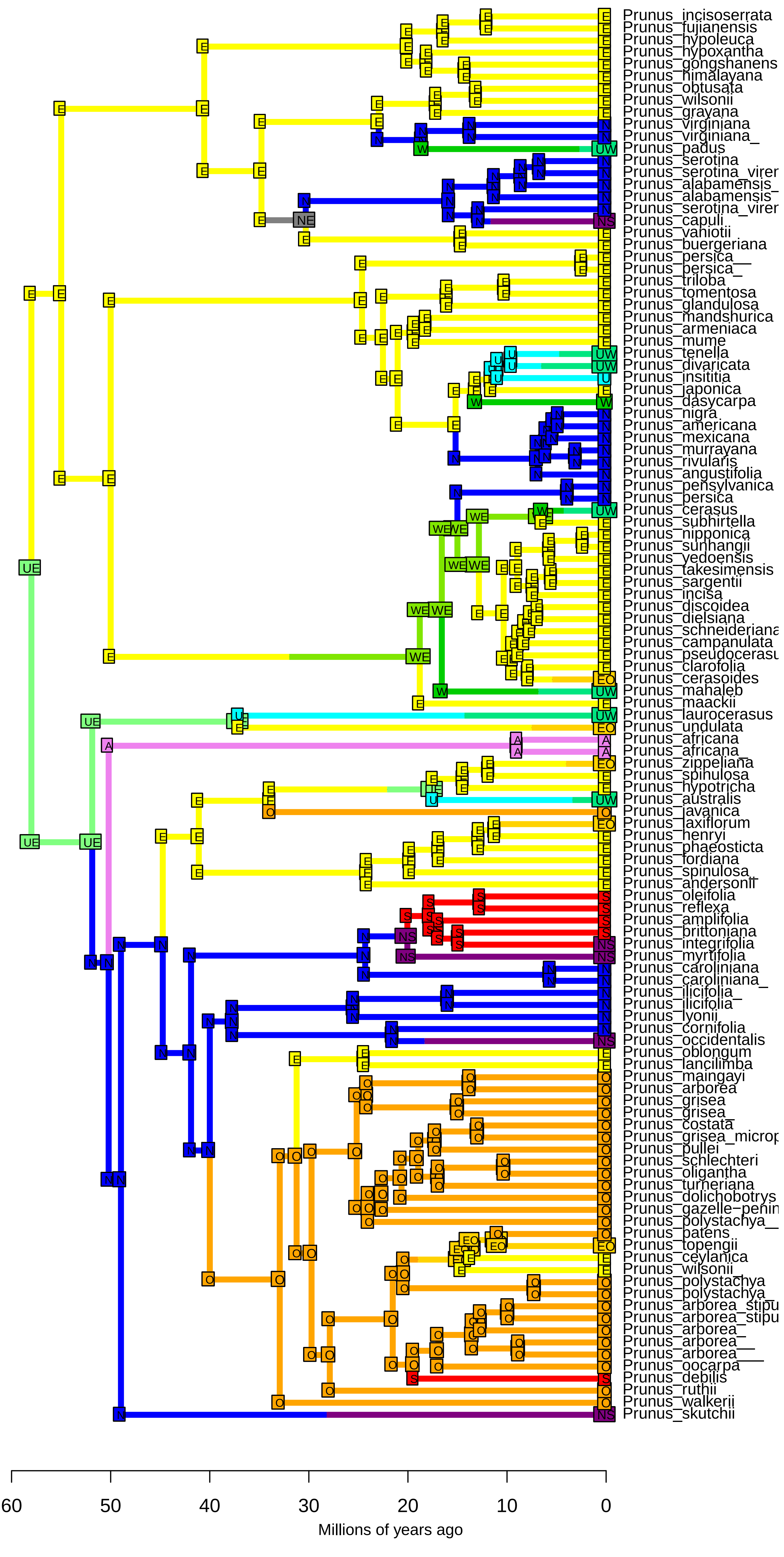

**DEC−J_M3_timestrat − Stochastic Map #6/50**
**ancstates: global optim, 2 areas max. d=0.0016; e=0; j=0.0148; LnL=−173.29**

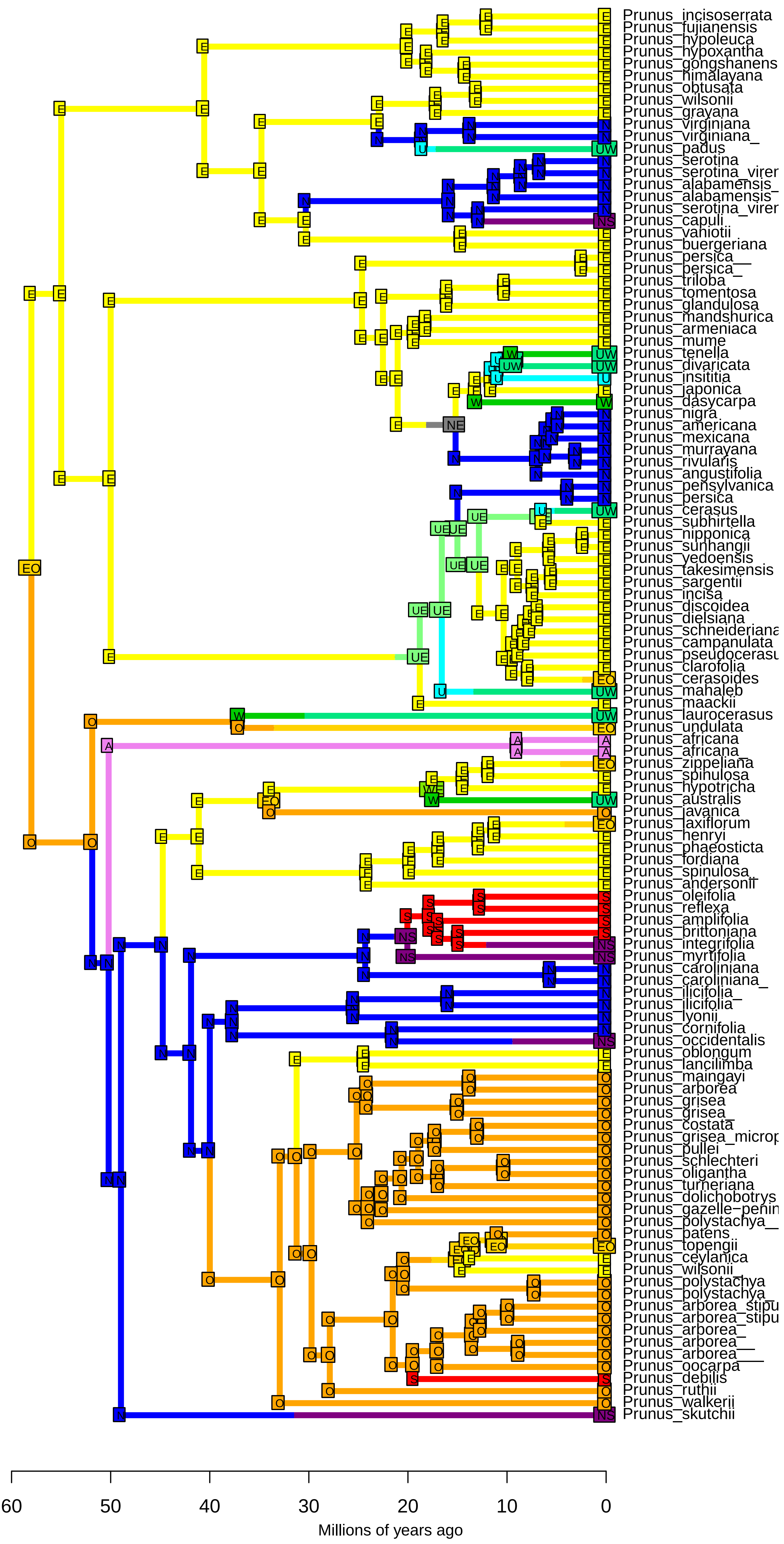

**DEC−J_M3_timestrat − Stochastic Map #7/50**
**ancstates: global optim, 2 areas max. d=0.0016; e=0; j=0.0148; LnL=−173.29**

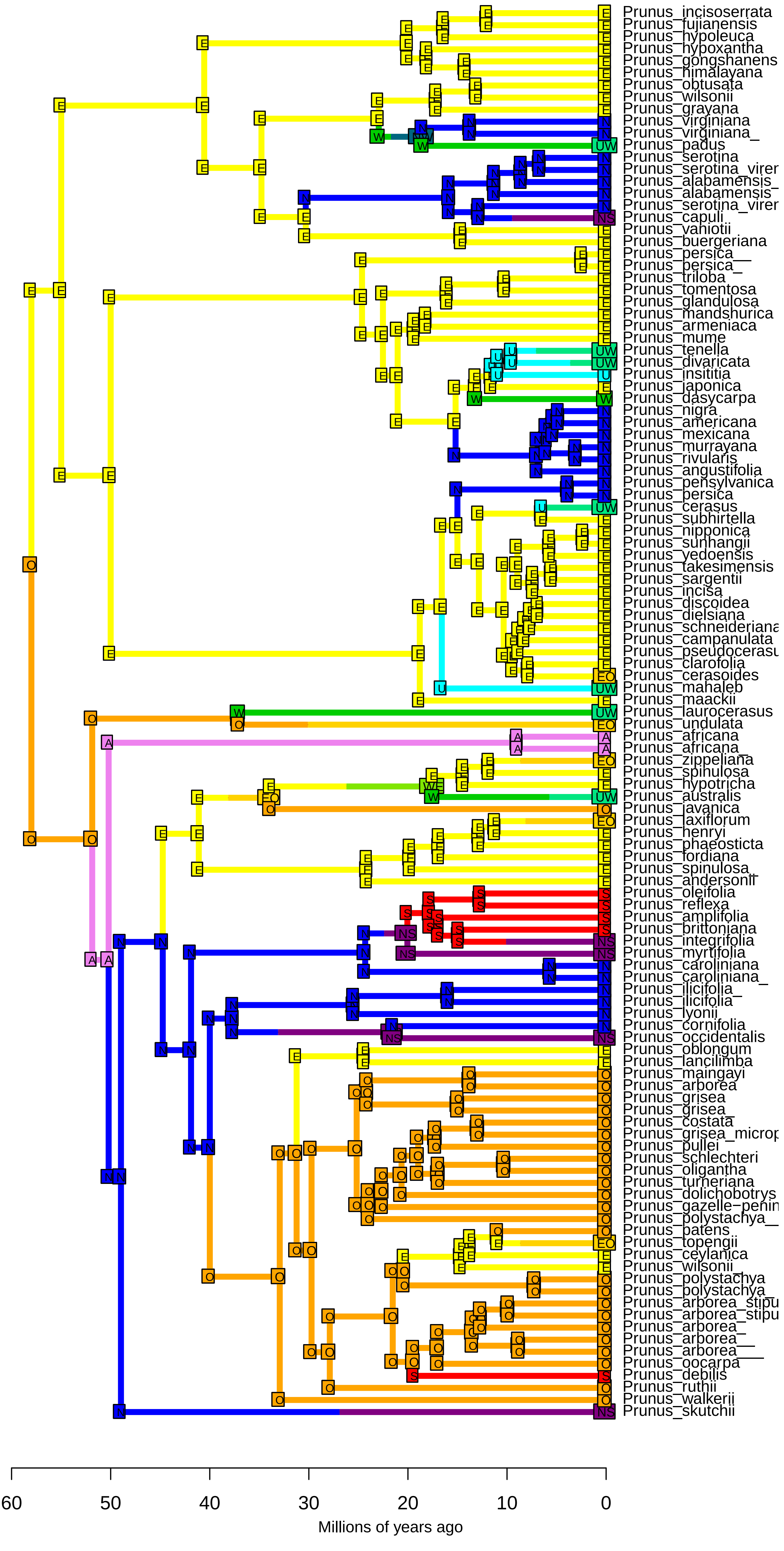

**DEC−J_M3_timestrat − Stochastic Map #8/50**
**ancstates: global optim, 2 areas max. d=0.0016; e=0; j=0.0148; LnL=−173.29**

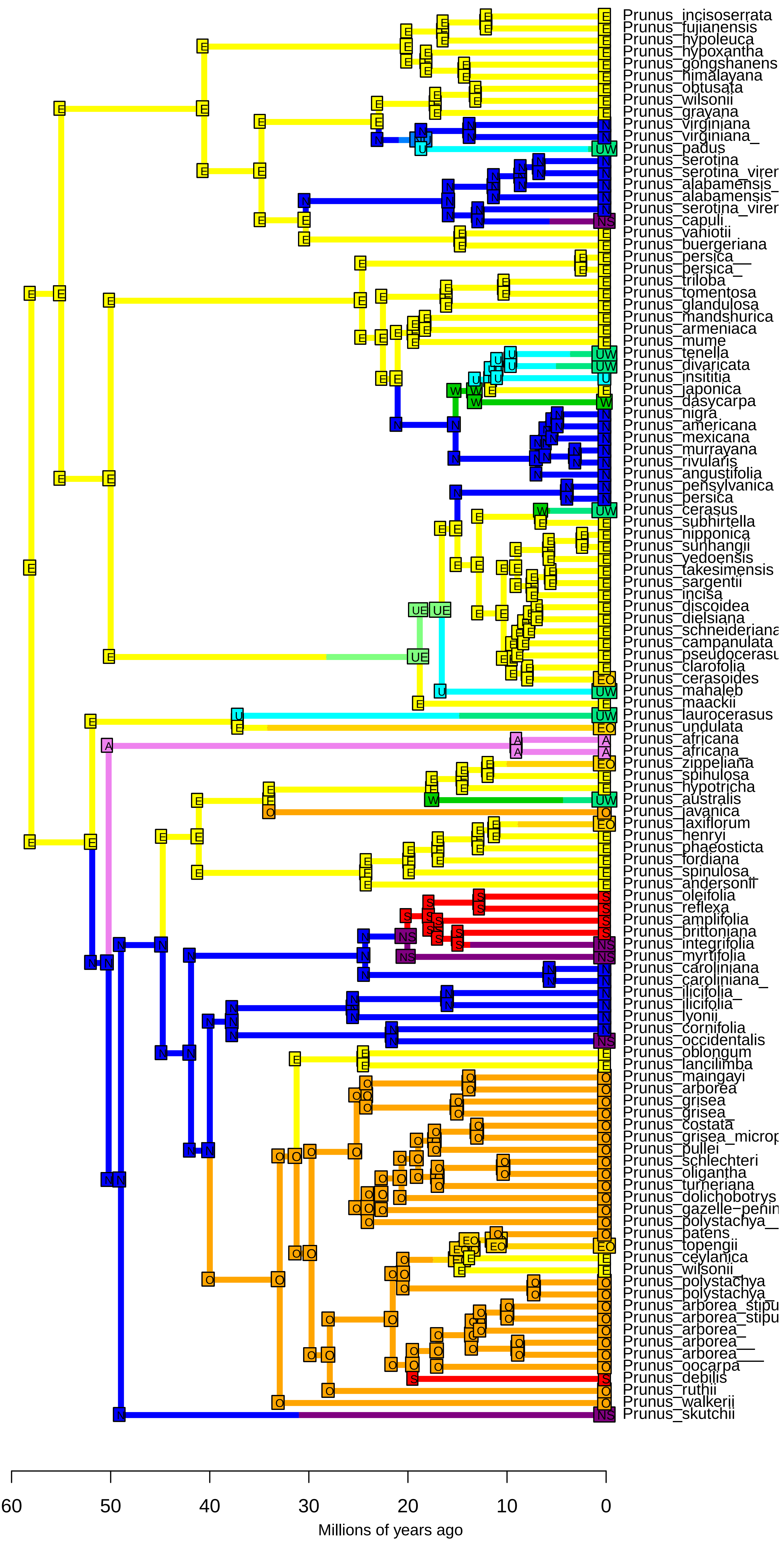

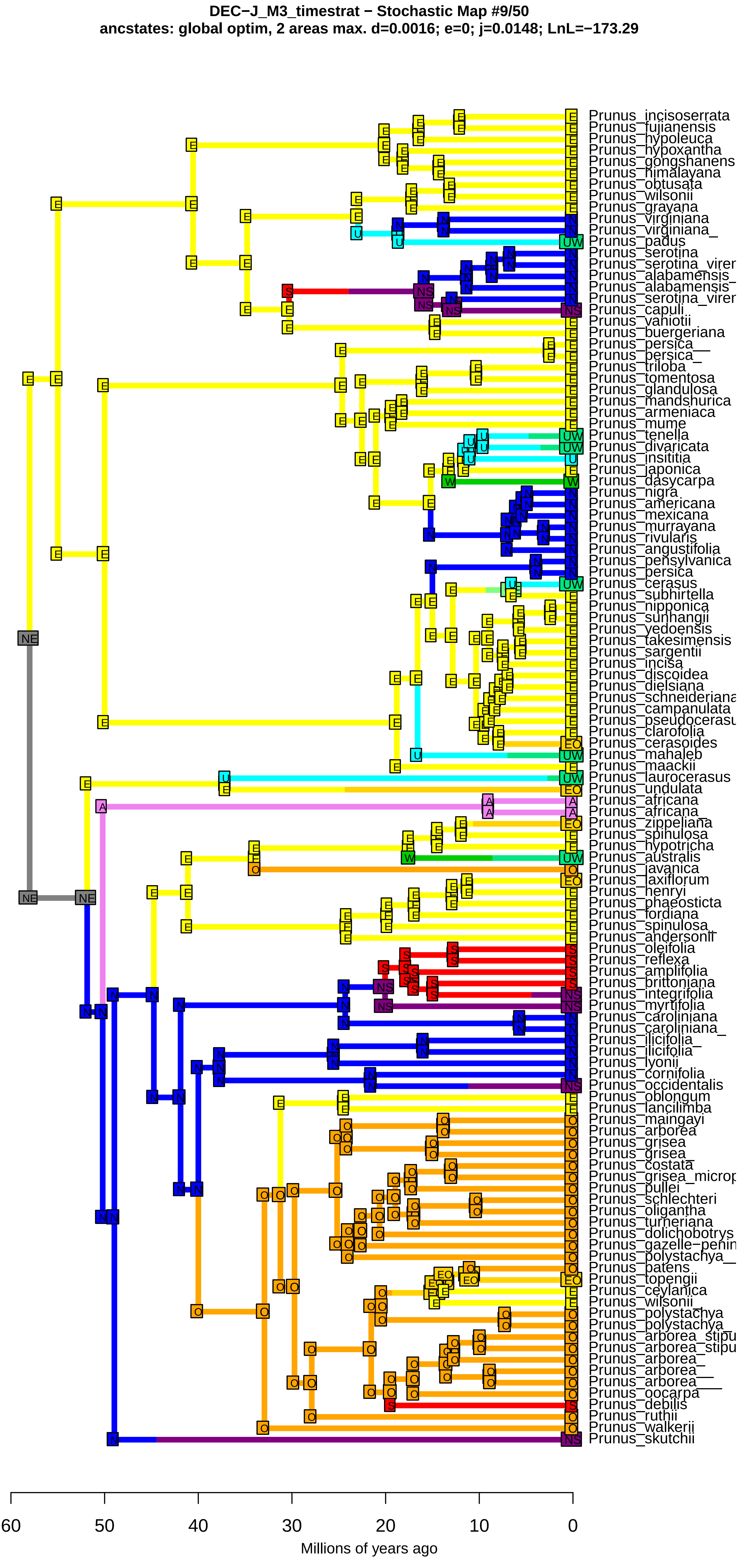
DEC−J_M3_timestrat − Stochastic Map #9/50
ancstates: global optim, 2 areas max. d=0.0016; e=0; j=0.0148; LnL=−173.29
Prunus_incisoserrata
Prunus_fujianensis
Prunus_hypoleuca
Prunus_hypoxantha
Prunus_gongshanens
Prunus_himalayana
Prunus_obtusata
Prunus_wilsonii
Prunus_grayana
Prunus_virginiana
Prunus_virginiana_
Prunus_padus
Prunus_serotina
Prunus_serotina_virer
Prunus_alabamensis_
Prunus_alabamensis
Prunus_serotina_virer
Prunus_capuli
Prunus_vaniotii
Prunus_buergeriana
Prunus_persica__
Prunus_persica_
Prunus_triloba
Prunus_tomentosa
Prunus_glandulosa
Prunus_mandshurica
Prunus_armeniaca
Prunus_mume
Prunus_tenella
Prunus_divaricata
Prunus_insititia
Prunus_japonica
Prunus_dasycarpa
Prunus_nigra
Prunus_americana
Prunus_mexicana
Prunus_murrayana
Prunus_rivularis
Prunus_angustifolia
Prunus_pensylvanica
Prunus_persica
Prunus_cerasus
Prunus_subhirtella
Prunus_nipponica
Prunus_sunhangii
Prunus_yedoensis
Prunus_takesimensis
Prunus_sargentii
Prunus_incisa
Prunus_discoidea
Prunus_dielsiana
Prunus_schneideriana
Prunus_campanulata
Prunus_pseudocerasu
Prunus_clarofolia
Prunus_cerasoides
Prunus_mahaleb
Prunus_maackii
Prunus_laurocerasus
Prunus_undulata
Prunus_africana
Prunus_africana_
Prunus_zippeliana
Prunus_spinulosa
Prunus_hypotricha
Prunus_australis
Prunus_javanica
Prunus_laxiflorum
Prunus_henryi
Prunus_phaeosticta
Prunus_fordiana
Prunus_spinulosa_
Prunus_andersonii
Prunus_oleifolia
Prunus_reflexa
Prunus_amplifolia
Prunus_brittoniana
Prunus_integrifolia
Prunus_myrtifolia
Prunus_caroliniana
Prunus_caroliniana_
Prunus_ilicifolia_
Prunus_ilicifolia
Prunus_lyonii
Prunus_cornifolia
Prunus_occidentalis
Prunus_oblongum
Prunus_lancilimba
Prunus_maingayi
Prunus_arborea
Prunus_grisea
Prunus_grisea_
Prunus_costata
Prunus_grisea_microp
Prunus_pullei
Prunus_schlechteri
Prunus_oligantha
Prunus_turneriana
Prunus_dolichobotrys
Prunus_gazelle−penin
Prunus_polystachya__
Prunus_patens
Prunus_topengii
Prunus_ceylanica
Prunus_wilsonii_
Prunus_polystachya
Prunus_polystachya_
Prunus_arborea_stipu
Prunus_arborea_stipu
Prunus_arborea_
Prunus_arborea__
Prunus_arborea___
Prunus_oocarpa
Prunus_debilis
Prunus_ruthii
Prunus_walkerii
Prunus_skutchii
60
50
40
30
20
10
0
Millions of years ago

**DEC−J_M3_timestrat − Stochastic Map #10/50**
**ancstates: global optim, 2 areas max. d=0.0016; e=0; j=0.0148; LnL=−173.29**

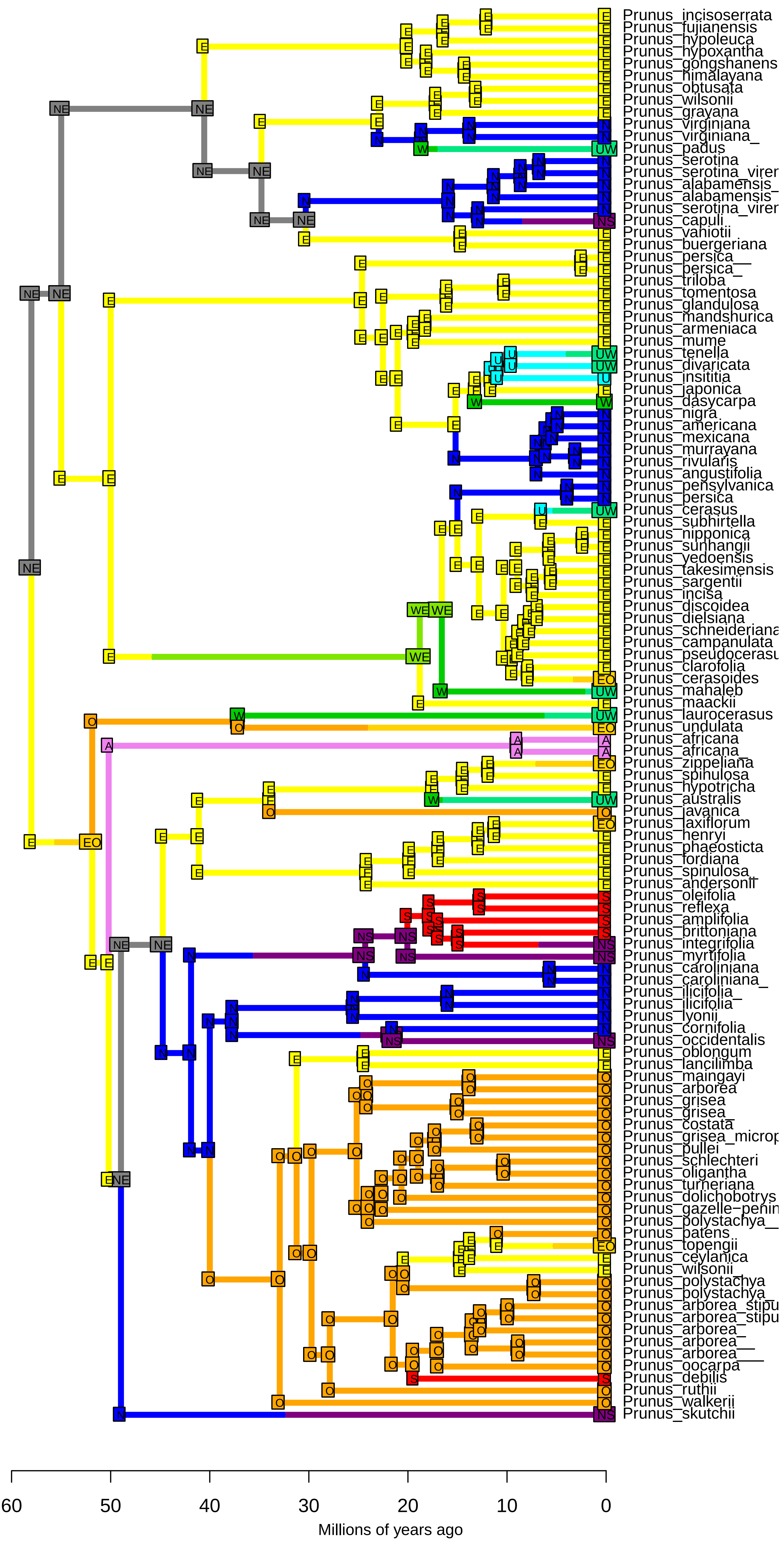

**DEC−J_M3_timestrat − Stochastic Map #11/50**
**ancstates: global optim, 2 areas max. d=0.0016; e=0; j=0.0148; LnL=−173.29**

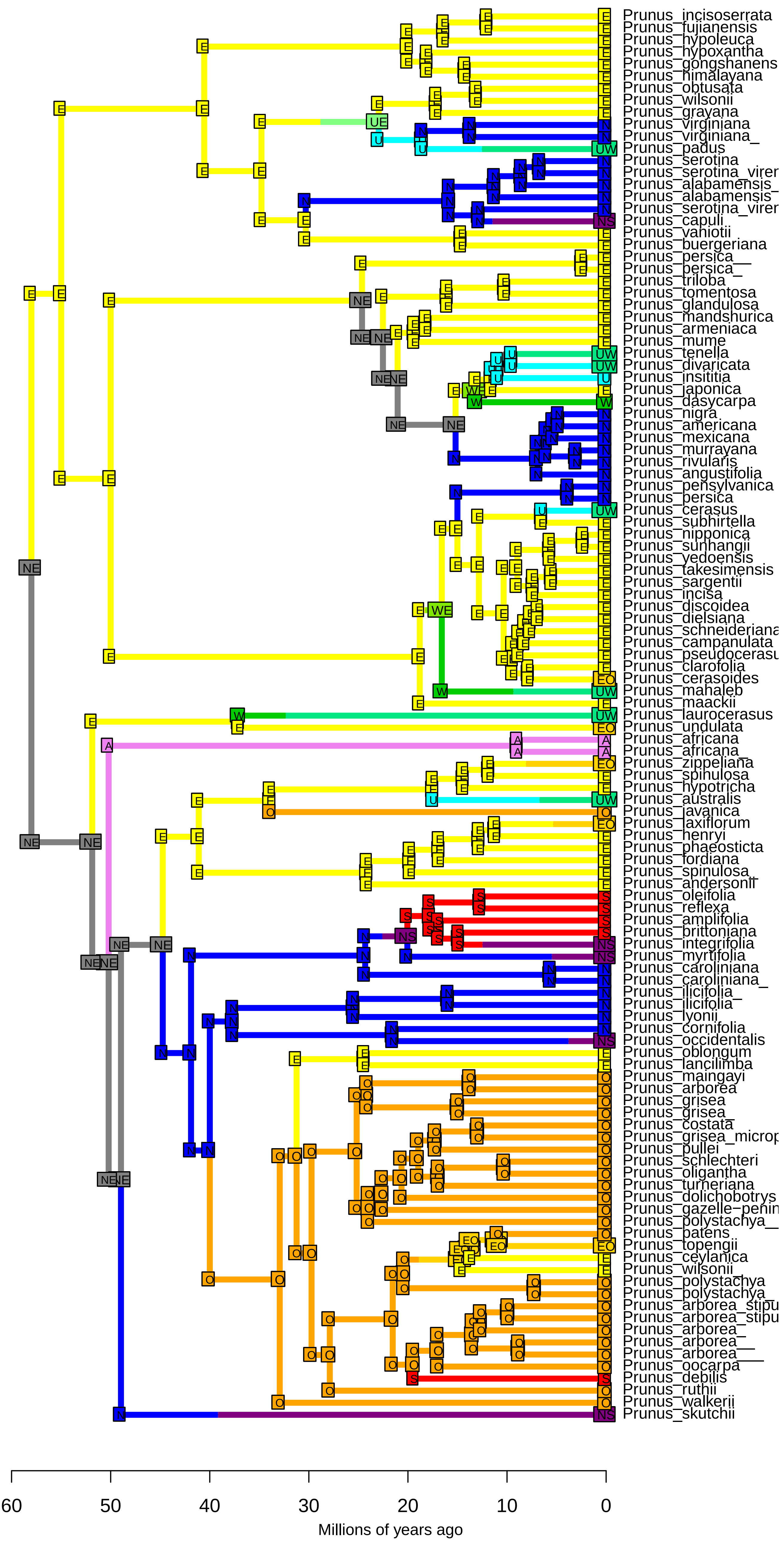

**DEC−J_M3_timestrat − Stochastic Map #12/50**
**ancstates: global optim, 2 areas max. d=0.0016; e=0; j=0.0148; LnL=−173.29**

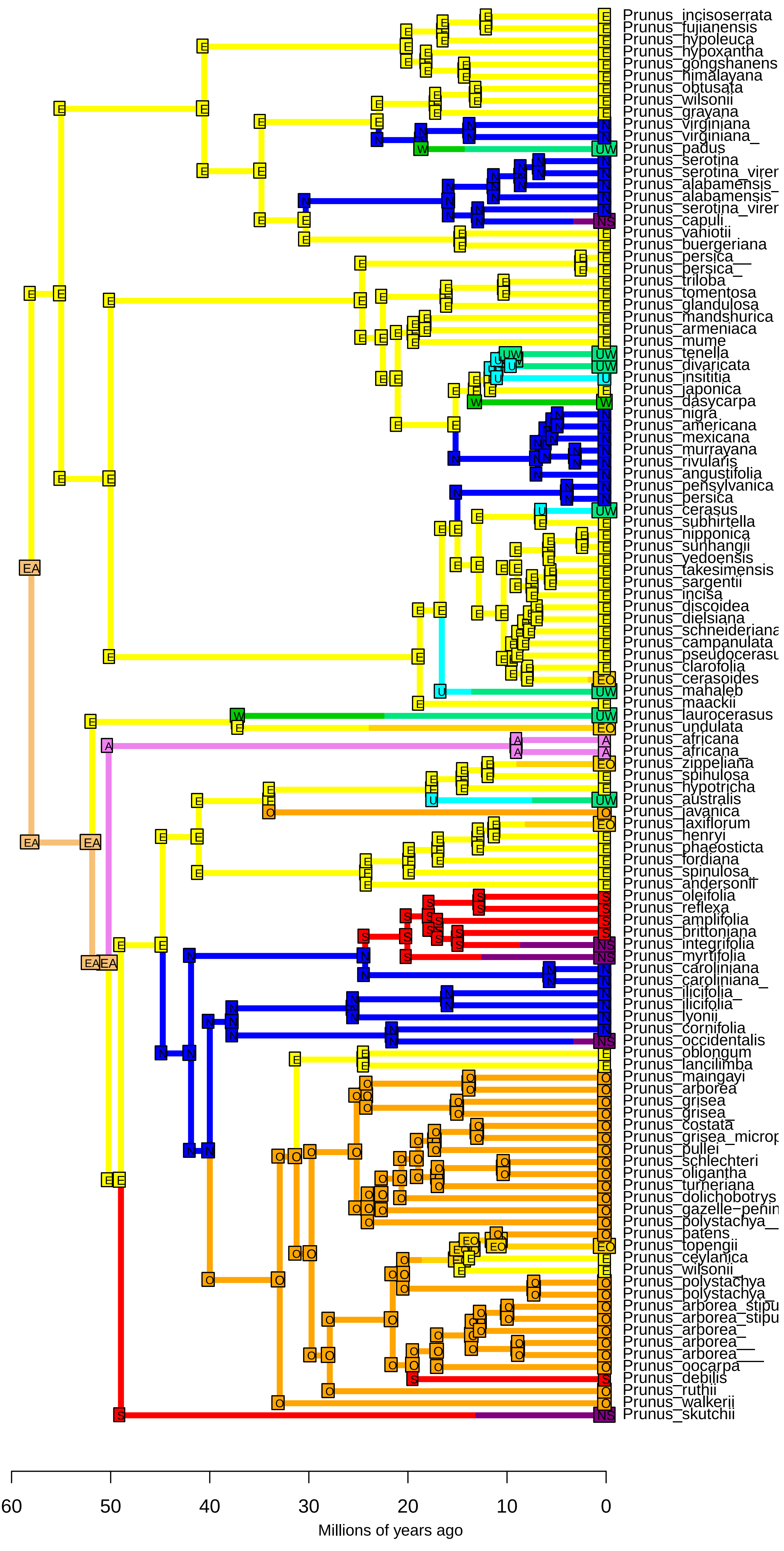

**DEC−J_M3_timestrat − Stochastic Map #13/50**
**ancstates: global optim, 2 areas max. d=0.0016; e=0; j=0.0148; LnL=−173.29**

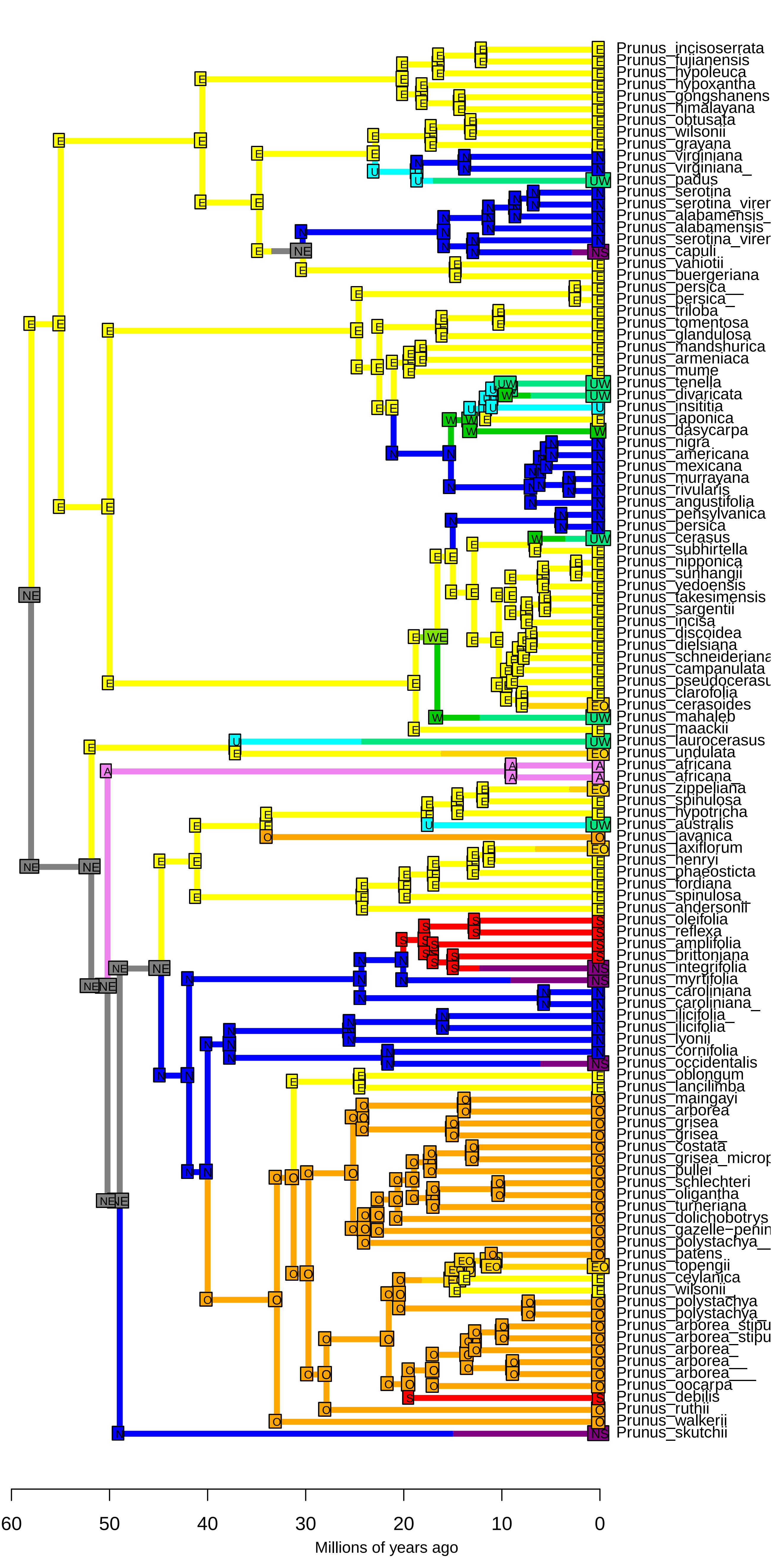

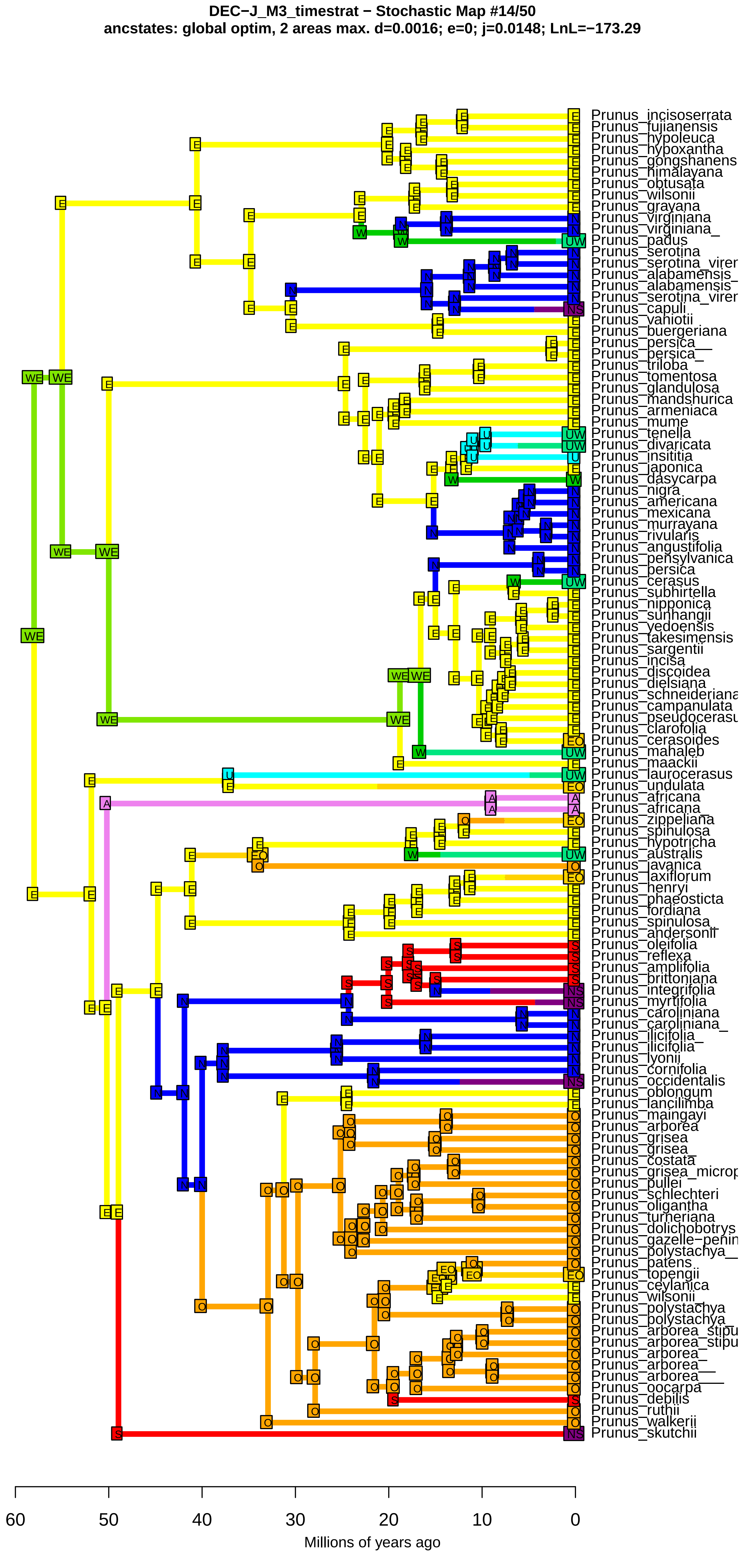
DEC−J_M3_timestrat − Stochastic Map #14/50
ancstates: global optim, 2 areas max. d=0.0016; e=0; j=0.0148; LnL=−173.29
Prunus_incisoserrata
Prunus_fujianensis
Prunus_hypoleuca
Prunus_hypoxantha
Prunus_gongshanens
Prunus_himalayana
Prunus_obtusata
Prunus_wilsonii
Prunus_grayana
Prunus_virginiana
Prunus_virginiana_
Prunus_padus
Prunus_serotina
Prunus_serotina_virer
Prunus_alabamensis_
Prunus_alabamensis
Prunus_serotina_virer
Prunus_capuli
Prunus_vaniotii
Prunus_buergeriana
Prunus_persica__
Prunus_persica_
Prunus_triloba
Prunus_tomentosa
Prunus_glandulosa
Prunus_mandshurica
Prunus_armeniaca
Prunus_mume
Prunus_tenella
Prunus_divaricata
Prunus_insititia
Prunus_japonica
Prunus_dasycarpa
Prunus_nigra
Prunus_americana
Prunus_mexicana
Prunus_murrayana
Prunus_rivularis
Prunus_angustifolia
Prunus_pensylvanica
Prunus_persica
Prunus_cerasus
Prunus_subhirtella
Prunus_nipponica
Prunus_sunhangii
Prunus_yedoensis
Prunus_takesimensis
Prunus_sargentii
Prunus_incisa
Prunus_discoidea
Prunus_dielsiana
Prunus_schneideriana
Prunus_campanulata
Prunus_pseudocerasu
Prunus_clarofolia
Prunus_cerasoides
Prunus_mahaleb
Prunus_maackii
Prunus_laurocerasus
Prunus_undulata
Prunus_africana
Prunus_africana_
Prunus_zippeliana
Prunus_spinulosa
Prunus_hypotricha
Prunus_australis
Prunus_javanica
Prunus_laxiflorum
Prunus_henryi
Prunus_phaeosticta
Prunus_fordiana
Prunus_spinulosa_
Prunus_andersonii
Prunus_oleifolia
Prunus_reflexa
Prunus_amplifolia
Prunus_brittoniana
Prunus_integrifolia
Prunus_myrtifolia
Prunus_caroliniana
Prunus_caroliniana_
Prunus_ilicifolia_
Prunus_ilicifolia
Prunus_lyonii
Prunus_cornifolia
Prunus_occidentalis
Prunus_oblongum
Prunus_lancilimba
Prunus_maingayi
Prunus_arborea
Prunus_grisea
Prunus_grisea_
Prunus_costata
Prunus_grisea_microp
Prunus_pullei
Prunus_schlechteri
Prunus_oligantha
Prunus_turneriana
Prunus_dolichobotrys
Prunus_gazelle−penin
Prunus_polystachya__
Prunus_patens
Prunus_topengii
Prunus_ceylanica
Prunus_wilsonii_
Prunus_polystachya
Prunus_polystachya_
Prunus_arborea_stipu
Prunus_arborea_stipu
Prunus_arborea_
Prunus_arborea__
Prunus_arborea___
Prunus_oocarpa
Prunus_debilis
Prunus_ruthii
Prunus_walkerii
Prunus_skutchii
60
50
40
30
20
10
0
Millions of years ago

**DEC−J_M3_timestrat − Stochastic Map #15/50**
**ancstates: global optim, 2 areas max. d=0.0016; e=0; j=0.0148; LnL=−173.29**

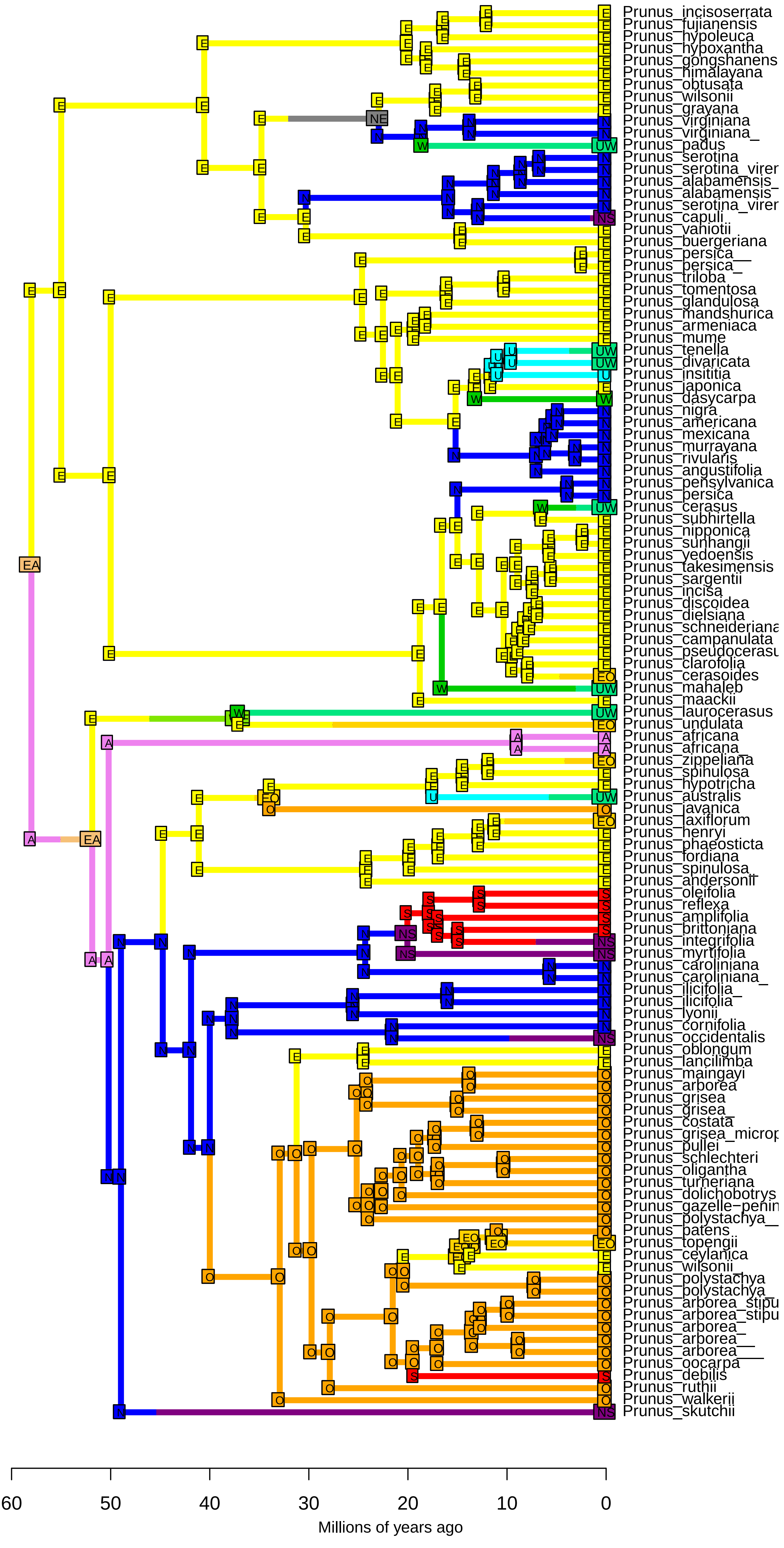

DEC−J_M3_timestrat − Stochastic Map #16/50
ancstates: global optim, 2 areas max. d=0.0016; e=0; j=0.0148; LnL=−173.29

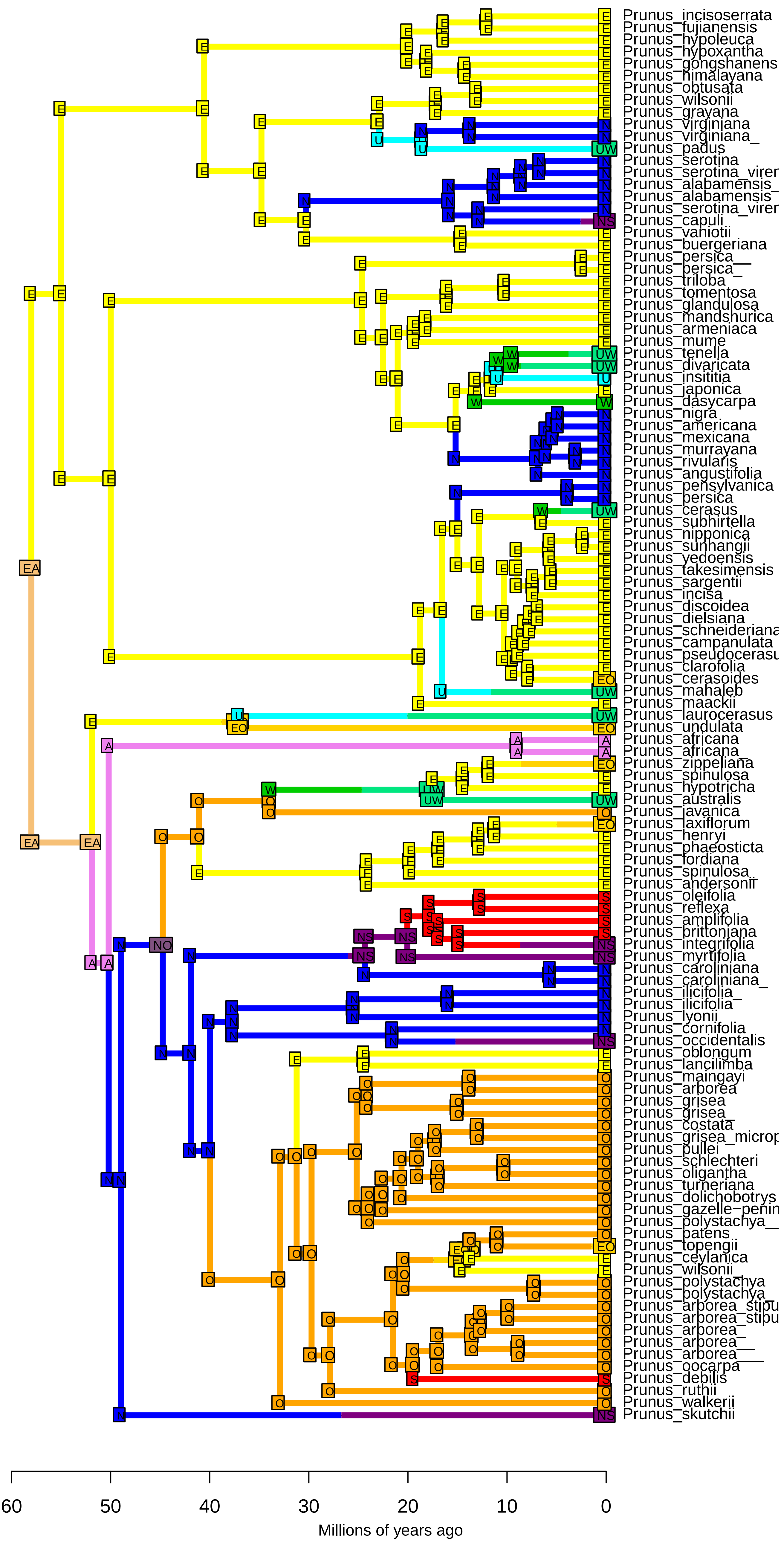

**DEC−J_M3_timestrat − Stochastic Map #17/50**
**ancstates: global optim, 2 areas max. d=0.0016; e=0; j=0.0148; LnL=−173.29**

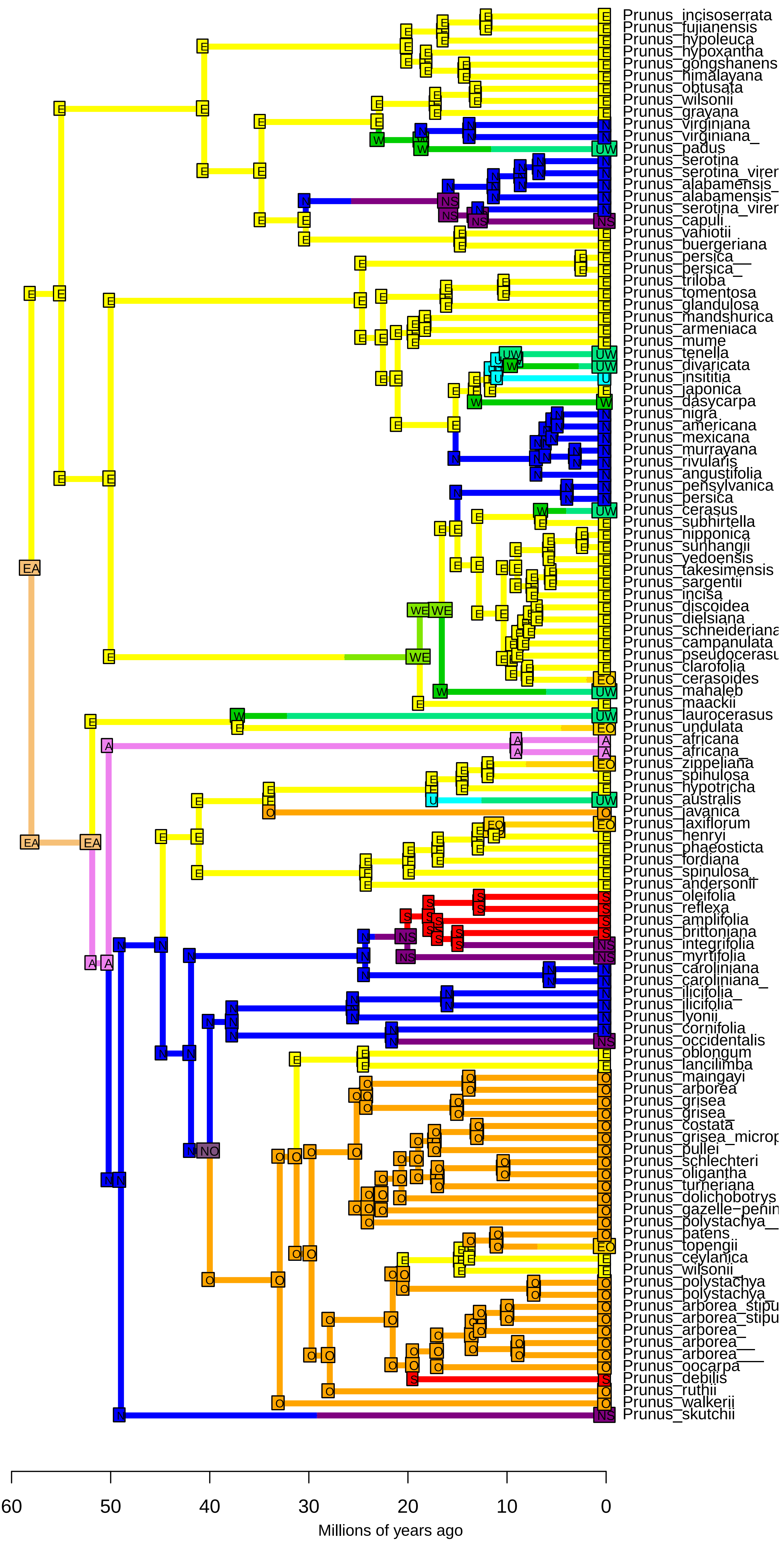

**DEC−J_M3_timestrat − Stochastic Map #18/50**
**ancstates: global optim, 2 areas max. d=0.0016; e=0; j=0.0148; LnL=−173.29**

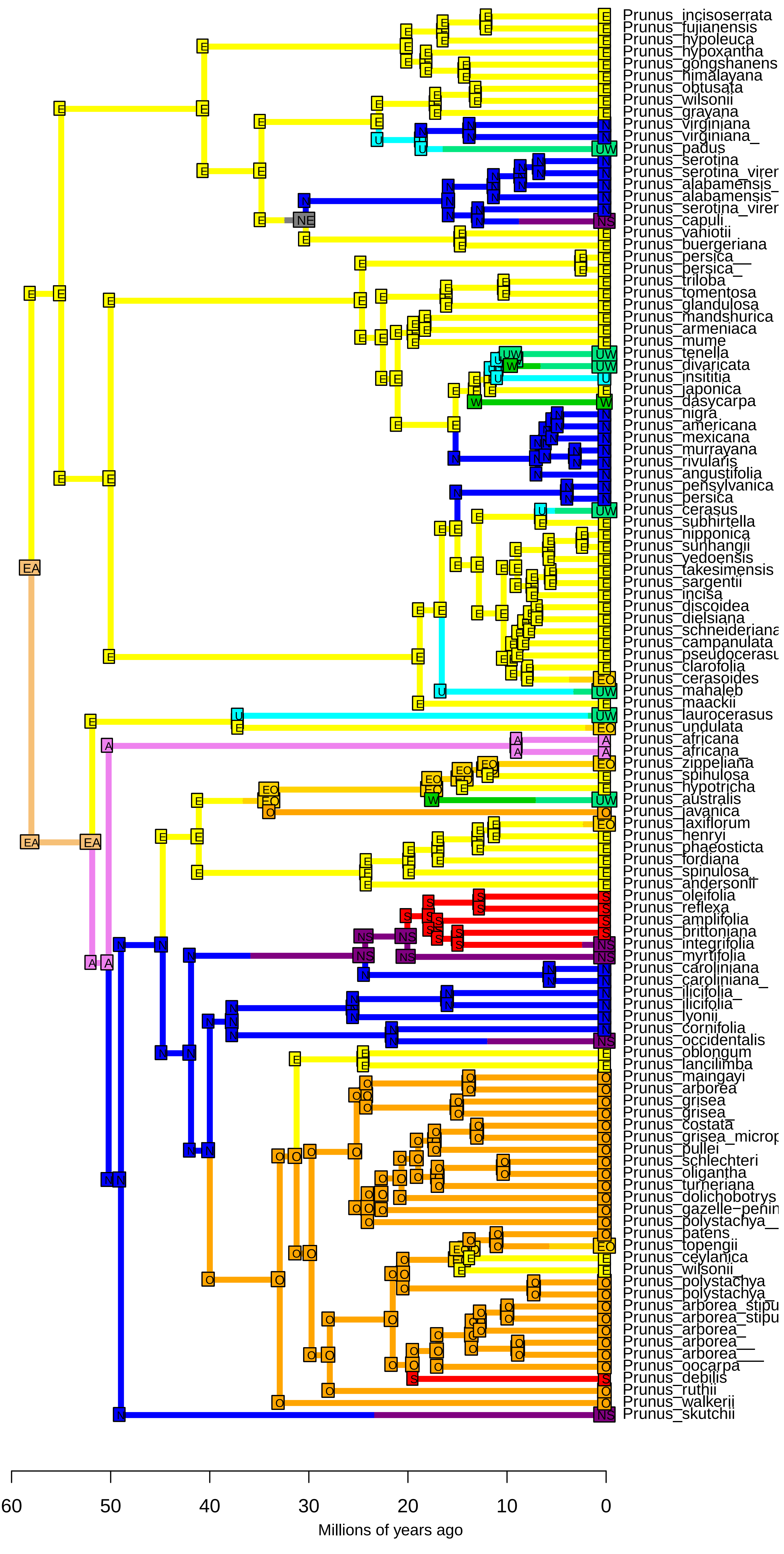

**DEC−J_M3_timestrat − Stochastic Map #19/50**
**ancstates: global optim, 2 areas max. d=0.0016; e=0; j=0.0148; LnL=−173.29**

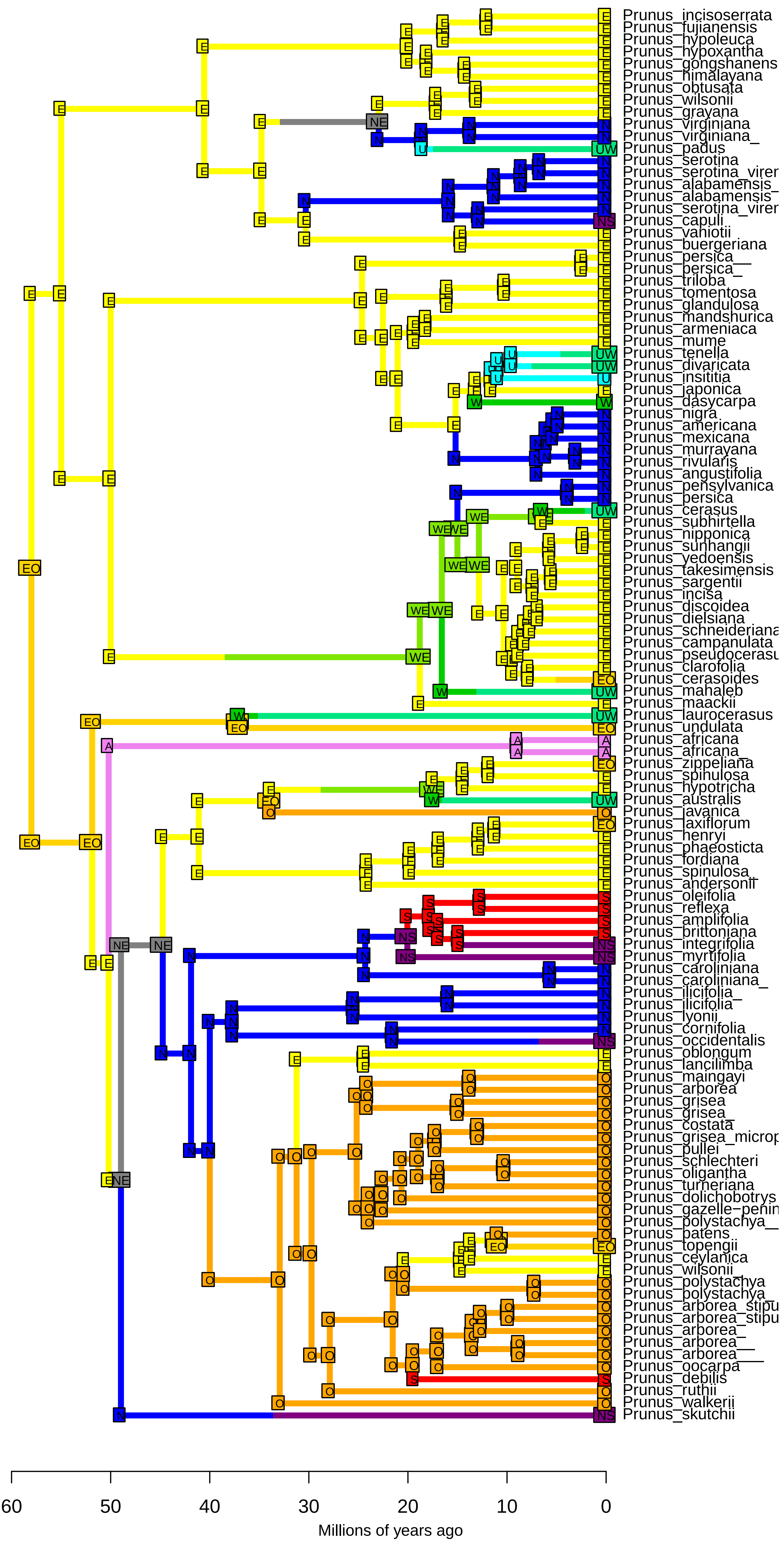

**DEC−J_M3_timestrat − Stochastic Map #20/50**
**ancstates: global optim, 2 areas max. d=0.0016; e=0; j=0.0148; LnL=−173.29**

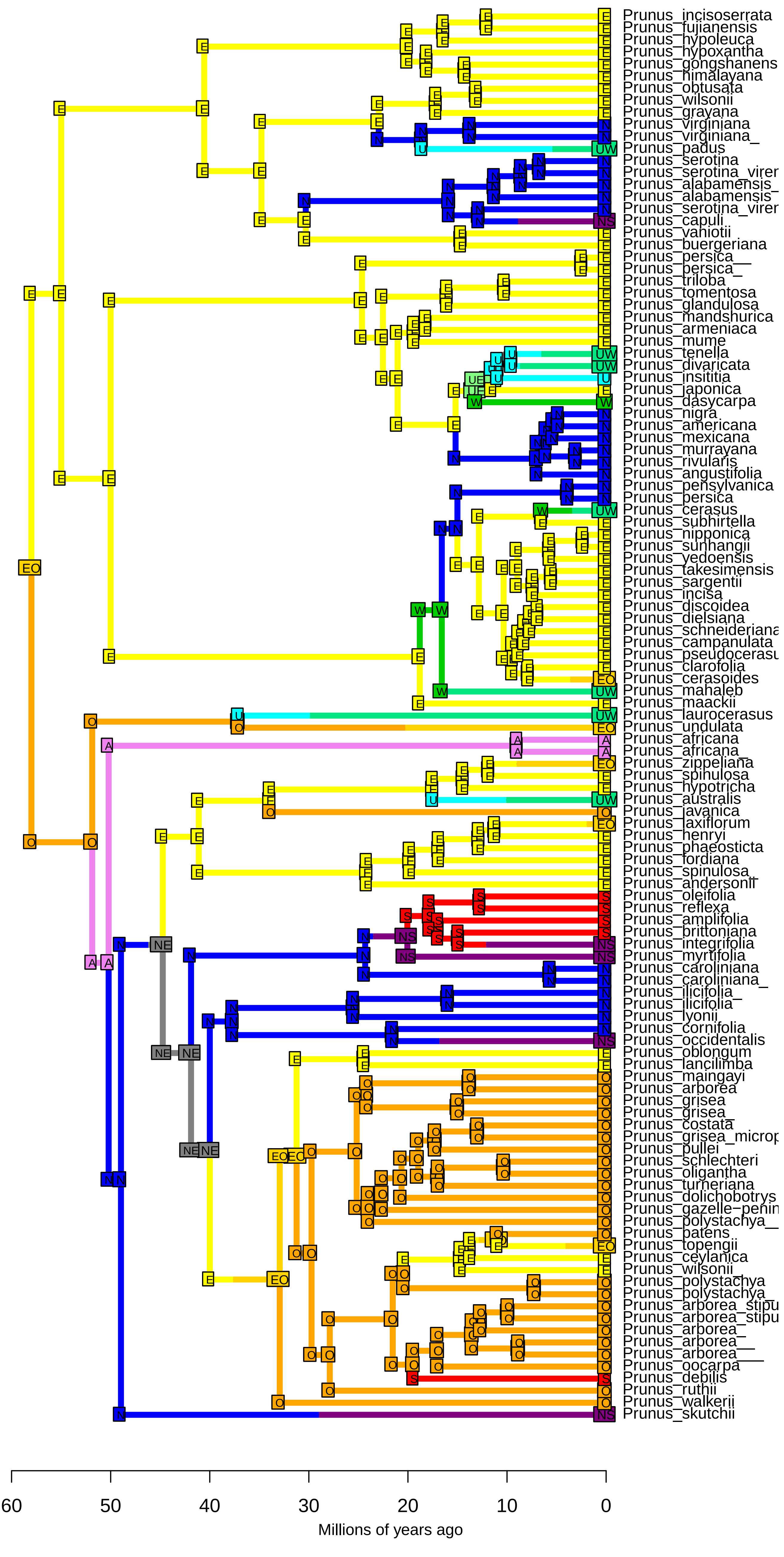

**DEC−J_M3_timestrat − Stochastic Map #21/50**
**ancstates: global optim, 2 areas max. d=0.0016; e=0; j=0.0148; LnL=−173.29**

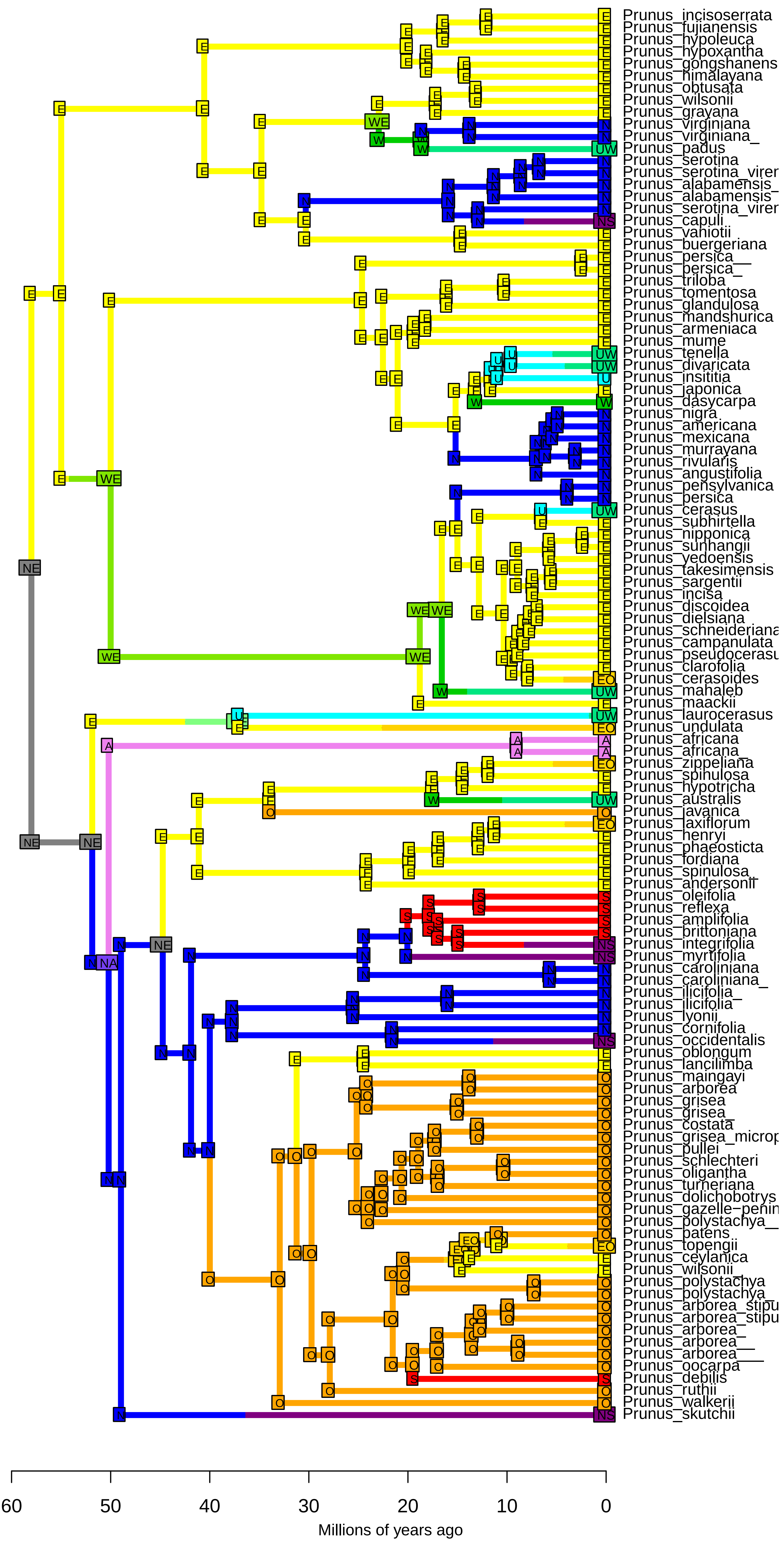

**DEC−J_M3_timestrat − Stochastic Map #22/50**
**ancstates: global optim, 2 areas max. d=0.0016; e=0; j=0.0148; LnL=−173.29**

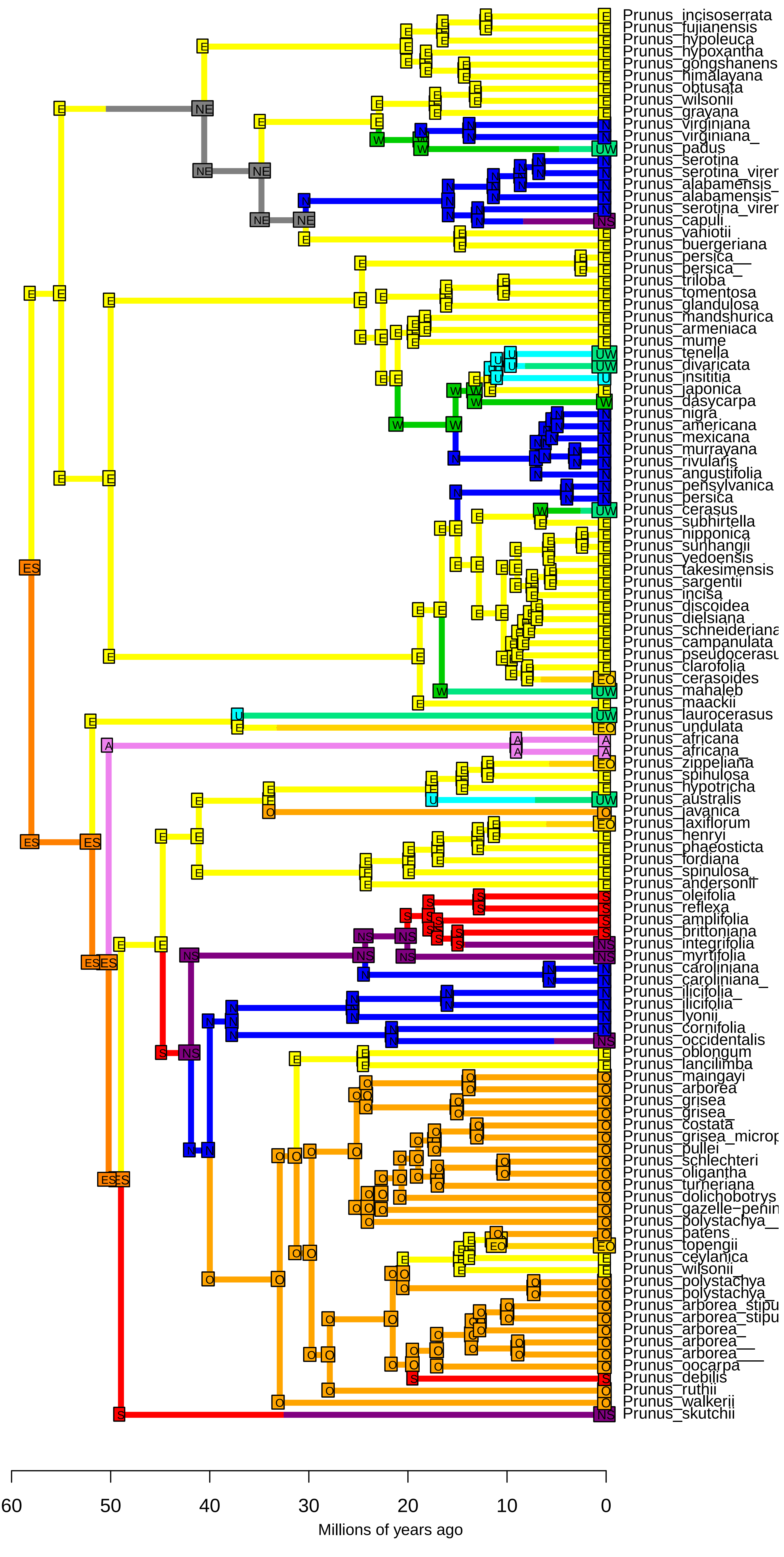

**DEC−J_M3_timestrat − Stochastic Map #23/50**
**ancstates: global optim, 2 areas max. d=0.0016; e=0; j=0.0148; LnL=−173.29**

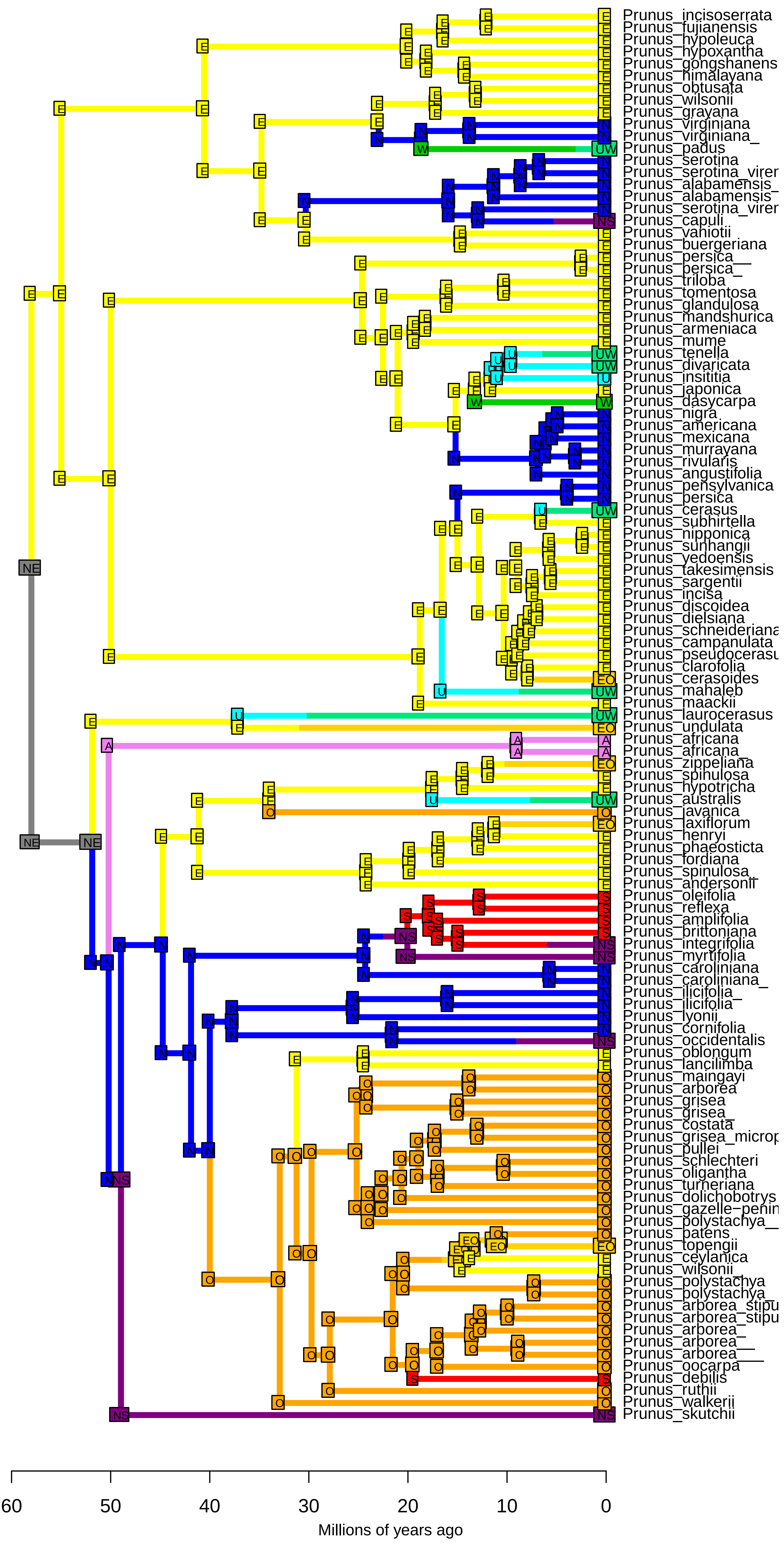

**DEC−J_M3_timestrat − Stochastic Map #24/50**
**ancstates: global optim, 2 areas max. d=0.0016; e=0; j=0.0148; LnL=−173.29**

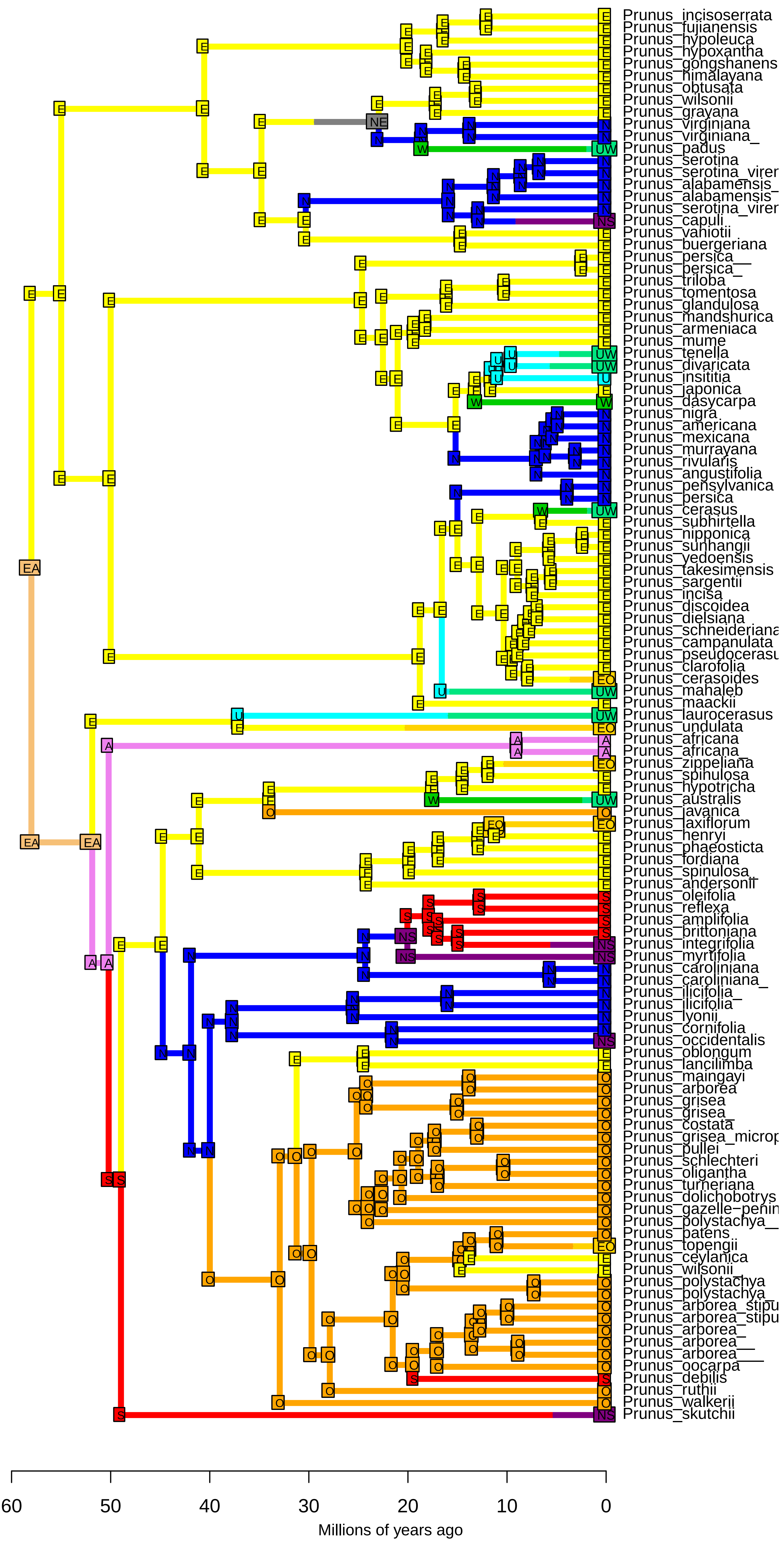

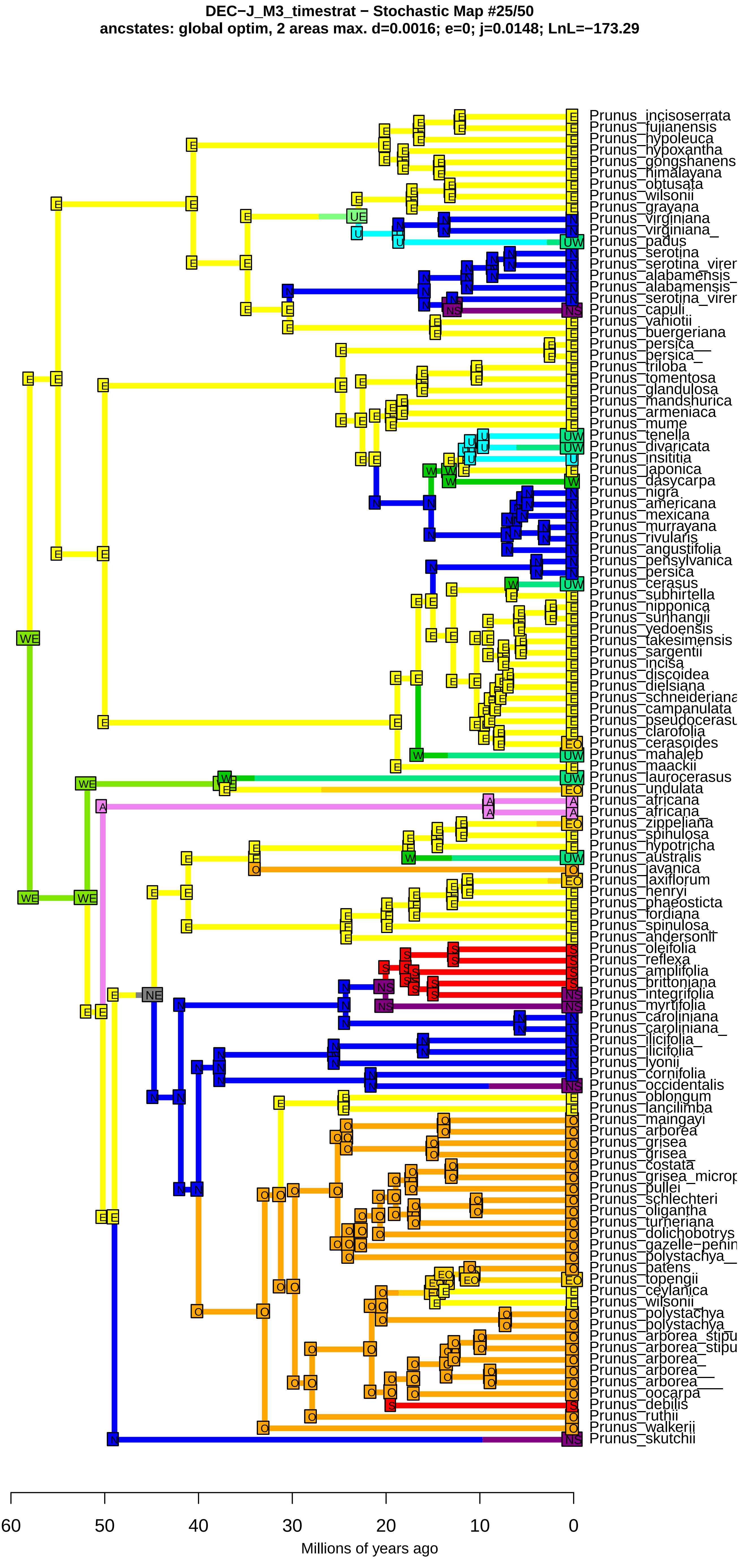
DEC−J_M3_timestrat − Stochastic Map #25/50
ancstates: global optim, 2 areas max. d=0.0016; e=0; j=0.0148; LnL=−173.29
Prunus_incisoserrata
Prunus_fujianensis
Prunus_hypoleuca
Prunus_hypoxantha
Prunus_gongshanens
Prunus_himalayana
Prunus_obtusata
Prunus_wilsonii
Prunus_grayana
Prunus_virginiana
Prunus_virginiana_
Prunus_padus
Prunus_serotina
Prunus_serotina_virer
Prunus_alabamensis_
Prunus_alabamensis
Prunus_serotina_virer
Prunus_capuli
Prunus_vaniotii
Prunus_buergeriana
Prunus_persica__
Prunus_persica_
Prunus_triloba
Prunus_tomentosa
Prunus_glandulosa
Prunus_mandshurica
Prunus_armeniaca
Prunus_mume
Prunus_tenella
Prunus_divaricata
Prunus_insititia
Prunus_japonica
Prunus_dasycarpa
Prunus_nigra
Prunus_americana
Prunus_mexicana
Prunus_murrayana
Prunus_rivularis
Prunus_angustifolia
Prunus_pensylvanica
Prunus_persica
Prunus_cerasus
Prunus_subhirtella
Prunus_nipponica
Prunus_sunhangii
Prunus_yedoensis
Prunus_takesimensis
Prunus_sargentii
Prunus_incisa
Prunus_discoidea
Prunus_dielsiana
Prunus_schneideriana
Prunus_campanulata
Prunus_pseudocerasu
Prunus_clarofolia
Prunus_cerasoides
Prunus_mahaleb
Prunus_maackii
Prunus_laurocerasus
Prunus_undulata
Prunus_africana
Prunus_africana_
Prunus_zippeliana
Prunus_spinulosa
Prunus_hypotricha
Prunus_australis
Prunus_javanica
Prunus_laxiflorum
Prunus_henryi
Prunus_phaeosticta
Prunus_fordiana
Prunus_spinulosa_
Prunus_andersonii
Prunus_oleifolia
Prunus_reflexa
Prunus_amplifolia
Prunus_brittoniana
Prunus_integrifolia
Prunus_myrtifolia
Prunus_caroliniana
Prunus_caroliniana_
Prunus_ilicifolia_
Prunus_ilicifolia
Prunus_lyonii
Prunus_cornifolia
Prunus_occidentalis
Prunus_oblongum
Prunus_lancilimba
Prunus_maingayi
Prunus_arborea
Prunus_grisea
Prunus_grisea_
Prunus_costata
Prunus_grisea_microp
Prunus_pullei
Prunus_schlechteri
Prunus_oligantha
Prunus_turneriana
Prunus_dolichobotrys
Prunus_gazelle−penin
Prunus_polystachya__
Prunus_patens
Prunus_topengii
Prunus_ceylanica
Prunus_wilsonii_
Prunus_polystachya
Prunus_polystachya_
Prunus_arborea_stipu
Prunus_arborea_stipu
Prunus_arborea_
Prunus_arborea__
Prunus_arborea___
Prunus_oocarpa
Prunus_debilis
Prunus_ruthii
Prunus_walkerii
Prunus_skutchii
60
50
40
30
20
10
0
Millions of years ago

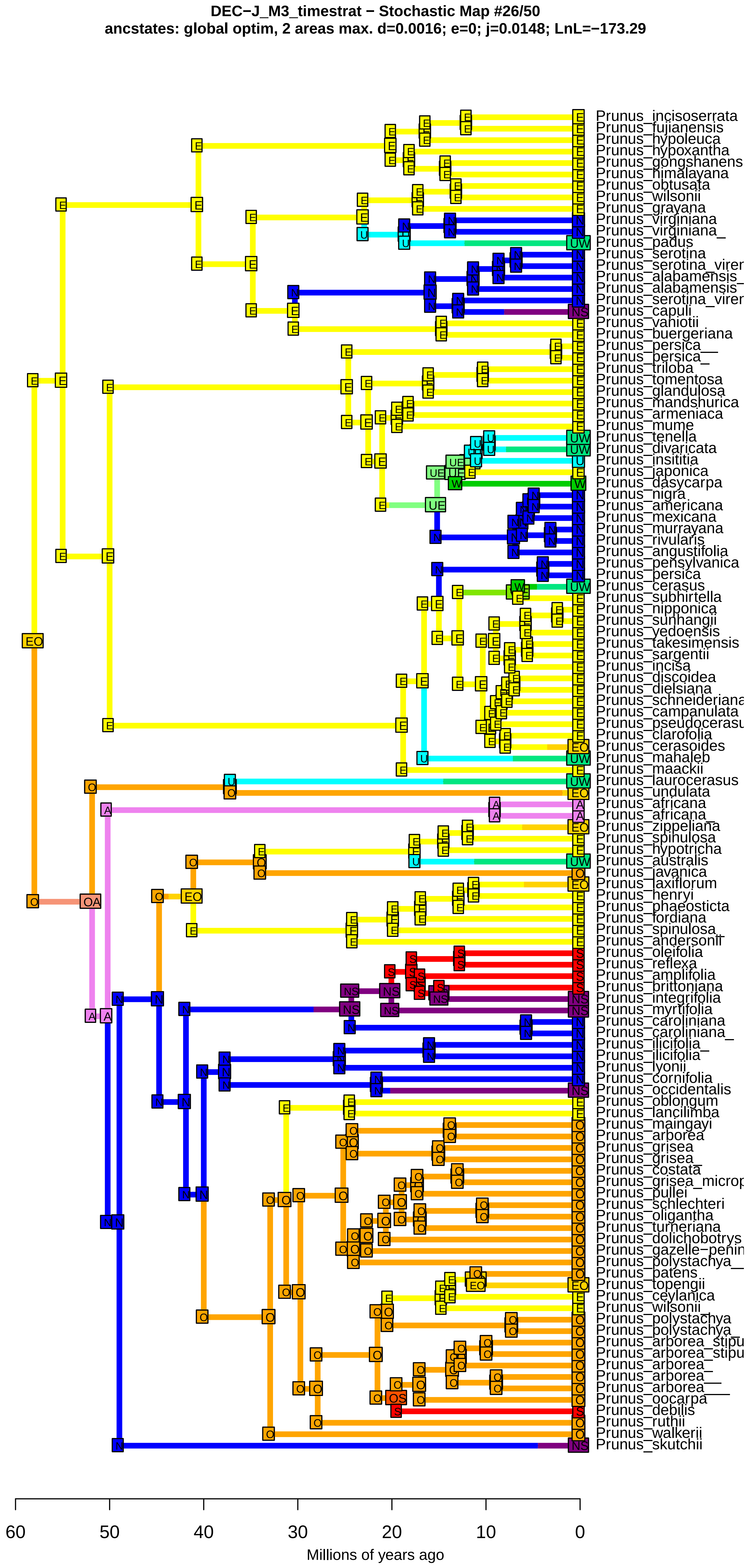
DEC−J_M3_timestrat − Stochastic Map #26/50
ancstates: global optim, 2 areas max. d=0.0016; e=0; j=0.0148; LnL=−173.29
Prunus_incisoserrata
Prunus_fujianensis
Prunus_hypoleuca
Prunus_hypoxantha
Prunus_gongshanens
Prunus_himalayana
Prunus_obtusata
Prunus_wilsonii
Prunus_grayana
Prunus_virginiana
Prunus_virginiana_
Prunus_padus
Prunus_serotina
Prunus_serotina_virer
Prunus_alabamensis_
Prunus_alabamensis
Prunus_serotina_virer
Prunus_capuli
Prunus_vaniotii
Prunus_buergeriana
Prunus_persica__
Prunus_persica_
Prunus_triloba
Prunus_tomentosa
Prunus_glandulosa
Prunus_mandshurica
Prunus_armeniaca
Prunus_mume
Prunus_tenella
Prunus_divaricata
Prunus_insititia
Prunus_japonica
Prunus_dasycarpa
Prunus_nigra
Prunus_americana
Prunus_mexicana
Prunus_murrayana
Prunus_rivularis
Prunus_angustifolia
Prunus_pensylvanica
Prunus_persica
Prunus_cerasus
Prunus_subhirtella
Prunus_nipponica
Prunus_sunhangii
Prunus_yedoensis
Prunus_takesimensis
Prunus_sargentii
Prunus_incisa
Prunus_discoidea
Prunus_dielsiana
Prunus_schneideriana
Prunus_campanulata
Prunus_pseudocerasu
Prunus_clarofolia
Prunus_cerasoides
Prunus_mahaleb
Prunus_maackii
Prunus_laurocerasus
Prunus_undulata
Prunus_africana
Prunus_africana_
Prunus_zippeliana
Prunus_spinulosa
Prunus_hypotricha
Prunus_australis
Prunus_javanica
Prunus_laxiflorum
Prunus_henryi
Prunus_phaeosticta
Prunus_fordiana
Prunus_spinulosa_
Prunus_andersonii
Prunus_oleifolia
Prunus_reflexa
Prunus_amplifolia
Prunus_brittoniana
Prunus_integrifolia
Prunus_myrtifolia
Prunus_caroliniana
Prunus_caroliniana_
Prunus_ilicifolia_
Prunus_ilicifolia
Prunus_lyonii
Prunus_cornifolia
Prunus_occidentalis
Prunus_oblongum
Prunus_lancilimba
Prunus_maingayi
Prunus_arborea
Prunus_grisea
Prunus_grisea_
Prunus_costata
Prunus_grisea_microp
Prunus_pullei
Prunus_schlechteri
Prunus_oligantha
Prunus_turneriana
Prunus_dolichobotrys
Prunus_gazelle−penin
Prunus_polystachya__
Prunus_patens
Prunus_topengii
Prunus_ceylanica
Prunus_wilsonii_
Prunus_polystachya
Prunus_polystachya_
Prunus_arborea_stipu
Prunus_arborea_stipu
Prunus_arborea_
Prunus_arborea__
Prunus_arborea___
Prunus_oocarpa
Prunus_debilis
Prunus_ruthii
Prunus_walkerii
Prunus_skutchii
60
50
40
30
20
10
0
Millions of years ago

**DEC−J_M3_timestrat − Stochastic Map #27/50**
**ancstates: global optim, 2 areas max. d=0.0016; e=0; j=0.0148; LnL=−173.29**

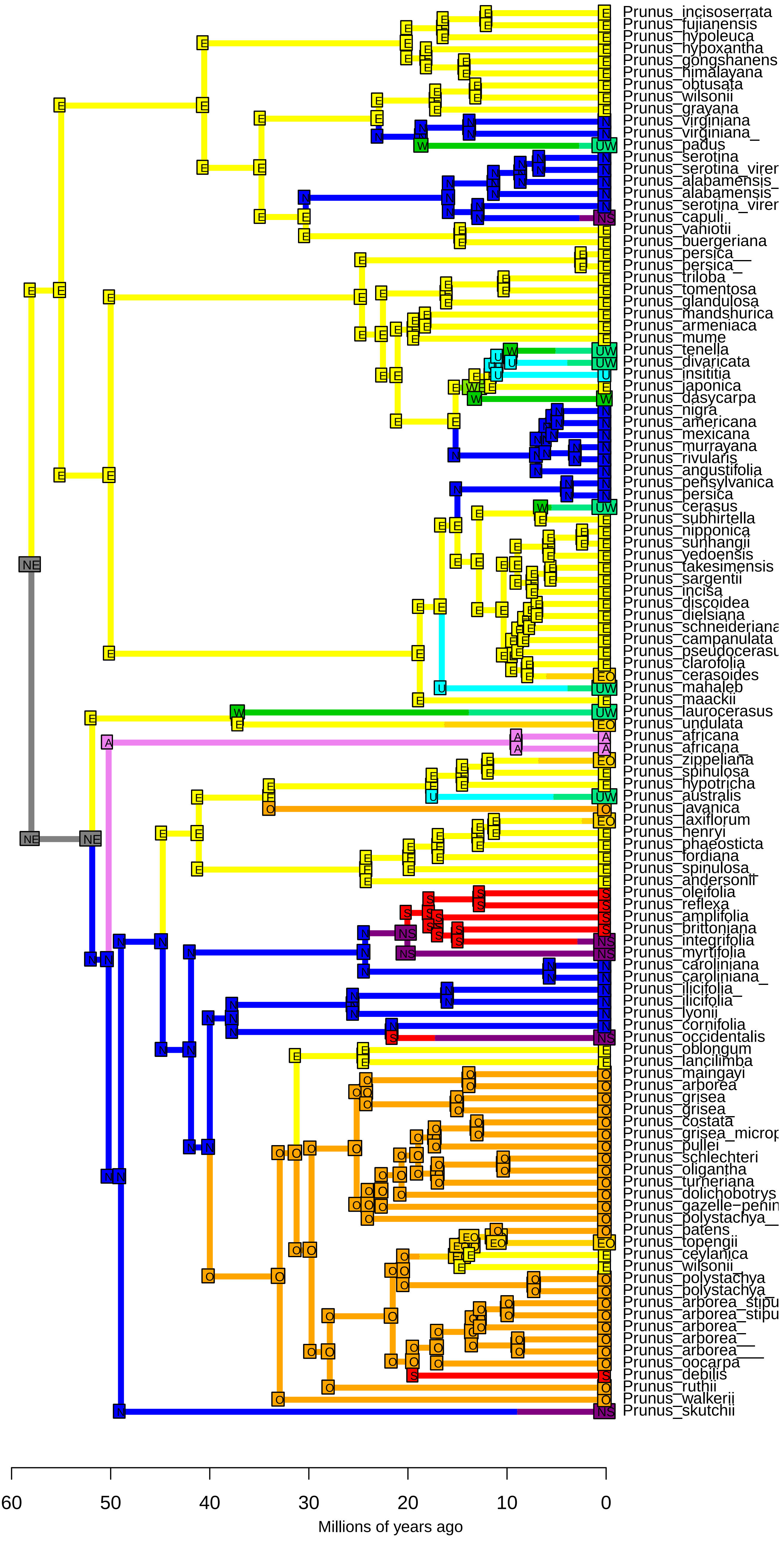

**DEC−J_M3_timestrat − Stochastic Map #28/50**
**ancstates: global optim, 2 areas max. d=0.0016; e=0; j=0.0148; LnL=−173.29**

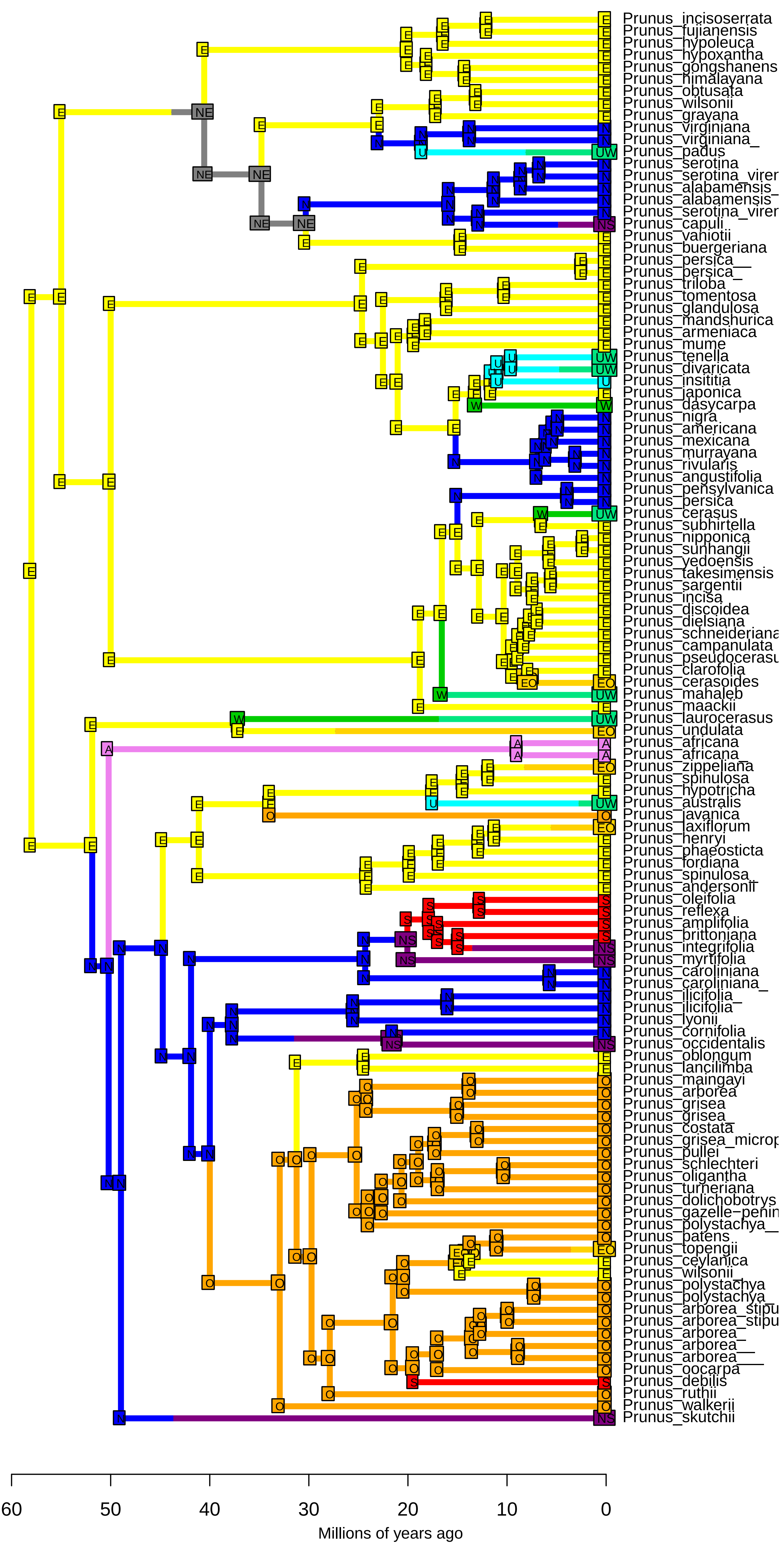

DEC−J_M3_timestrat − Stochastic Map #29/50
ancstates: global optim, 2 areas max. d=0.0016; e=0; j=0.0148; LnL=−173.29

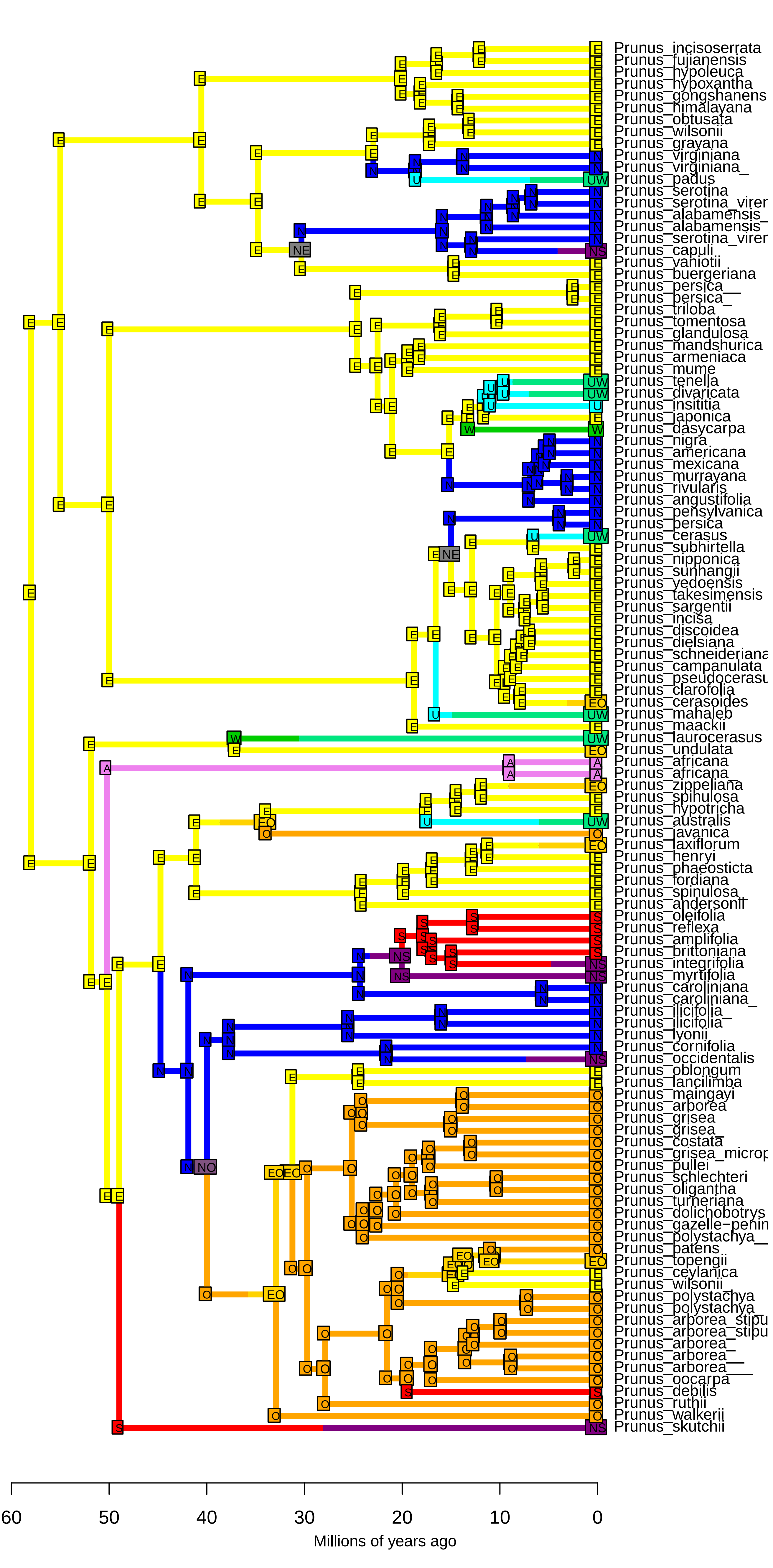

DEC−J_M3_timestrat − Stochastic Map #30/50
ancstates: global optim, 2 areas max. d=0.0016; e=0; j=0.0148; LnL=−173.29

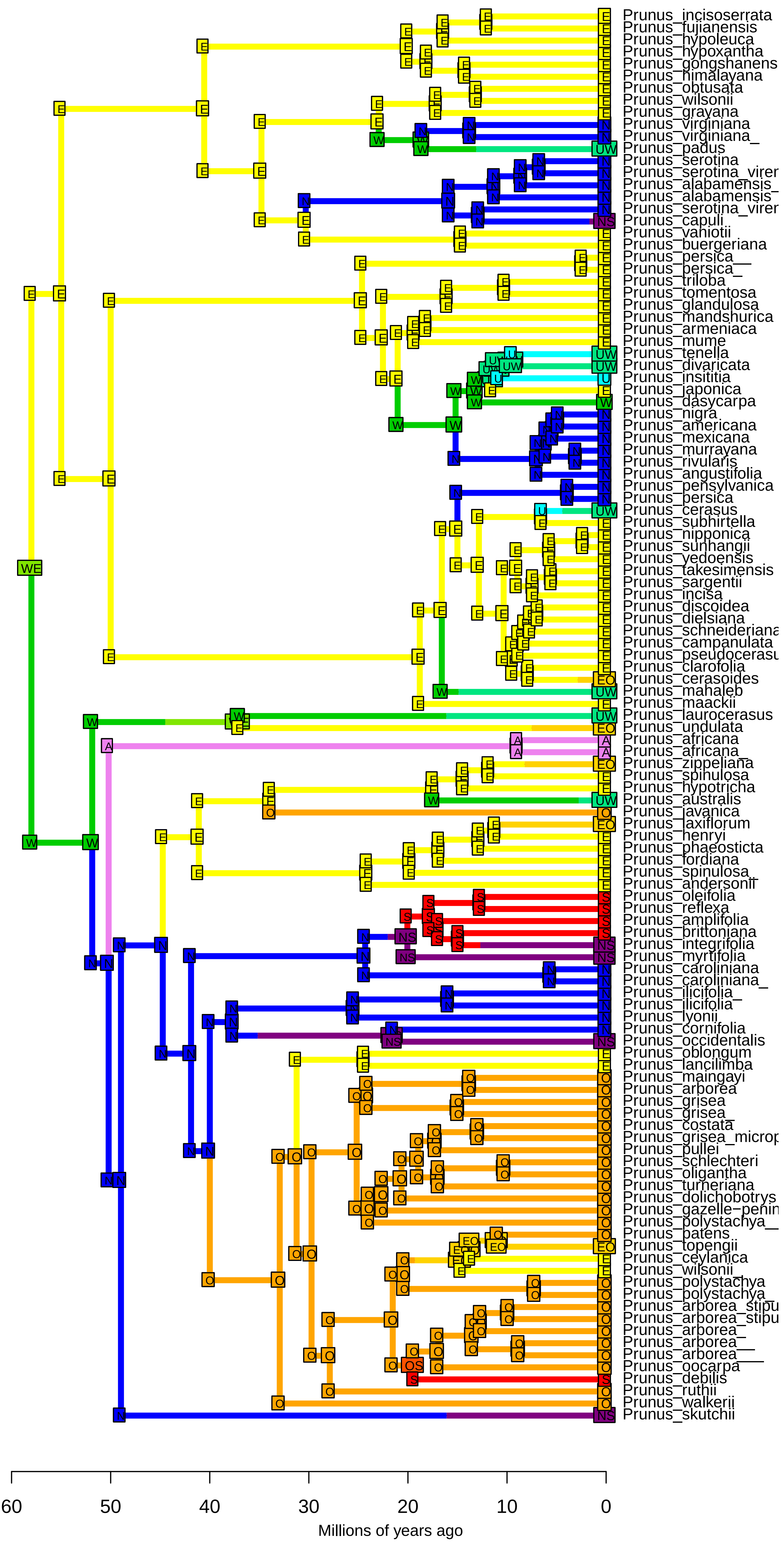

**DEC−J_M3_timestrat − Stochastic Map #31/50**
**ancstates: global optim, 2 areas max. d=0.0016; e=0; j=0.0148; LnL=−173.29**

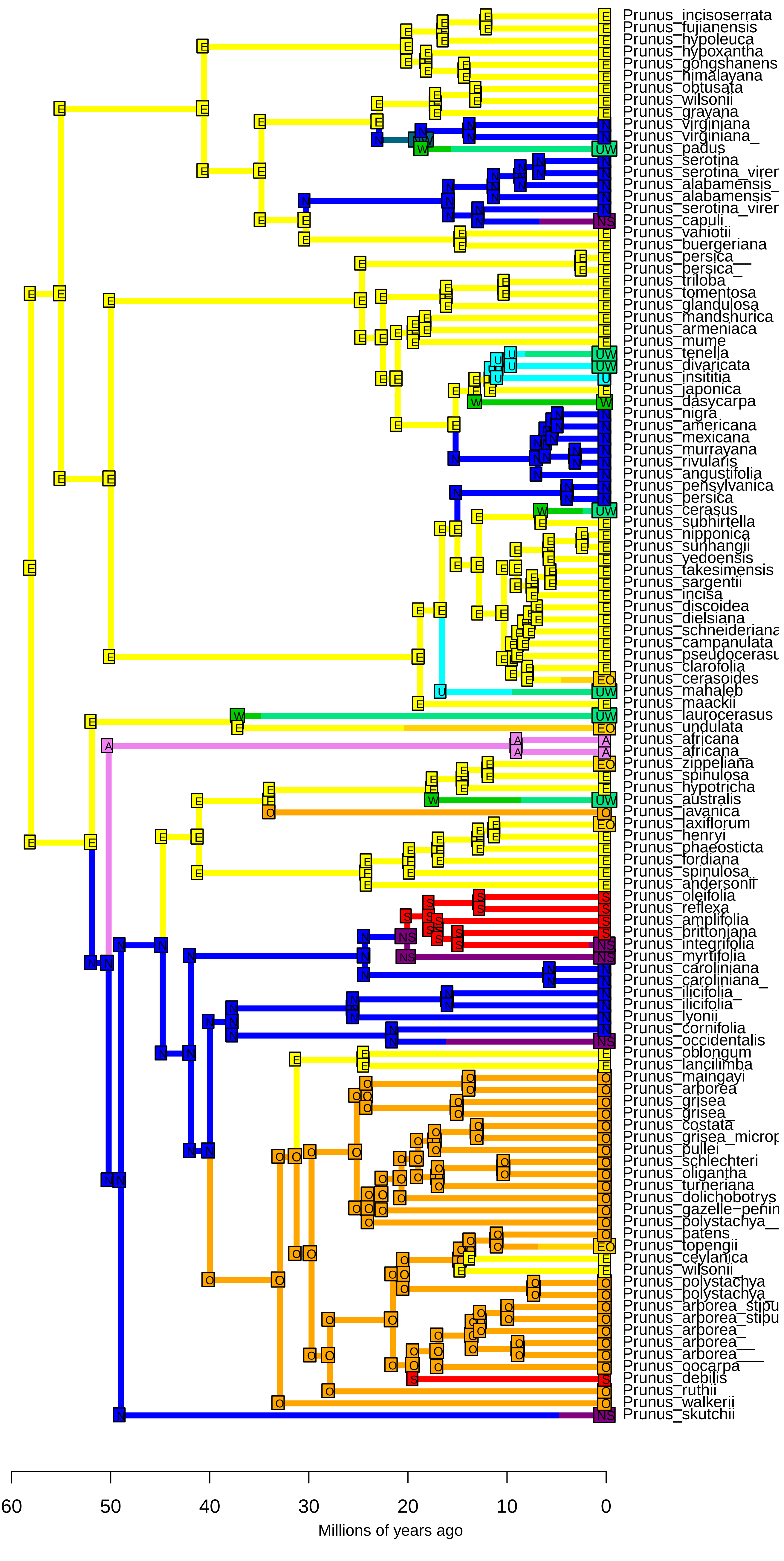

DEC−J_M3_timestrat − Stochastic Map #32/50
ancstates: global optim, 2 areas max. d=0.0016; e=0; j=0.0148; LnL=−173.29

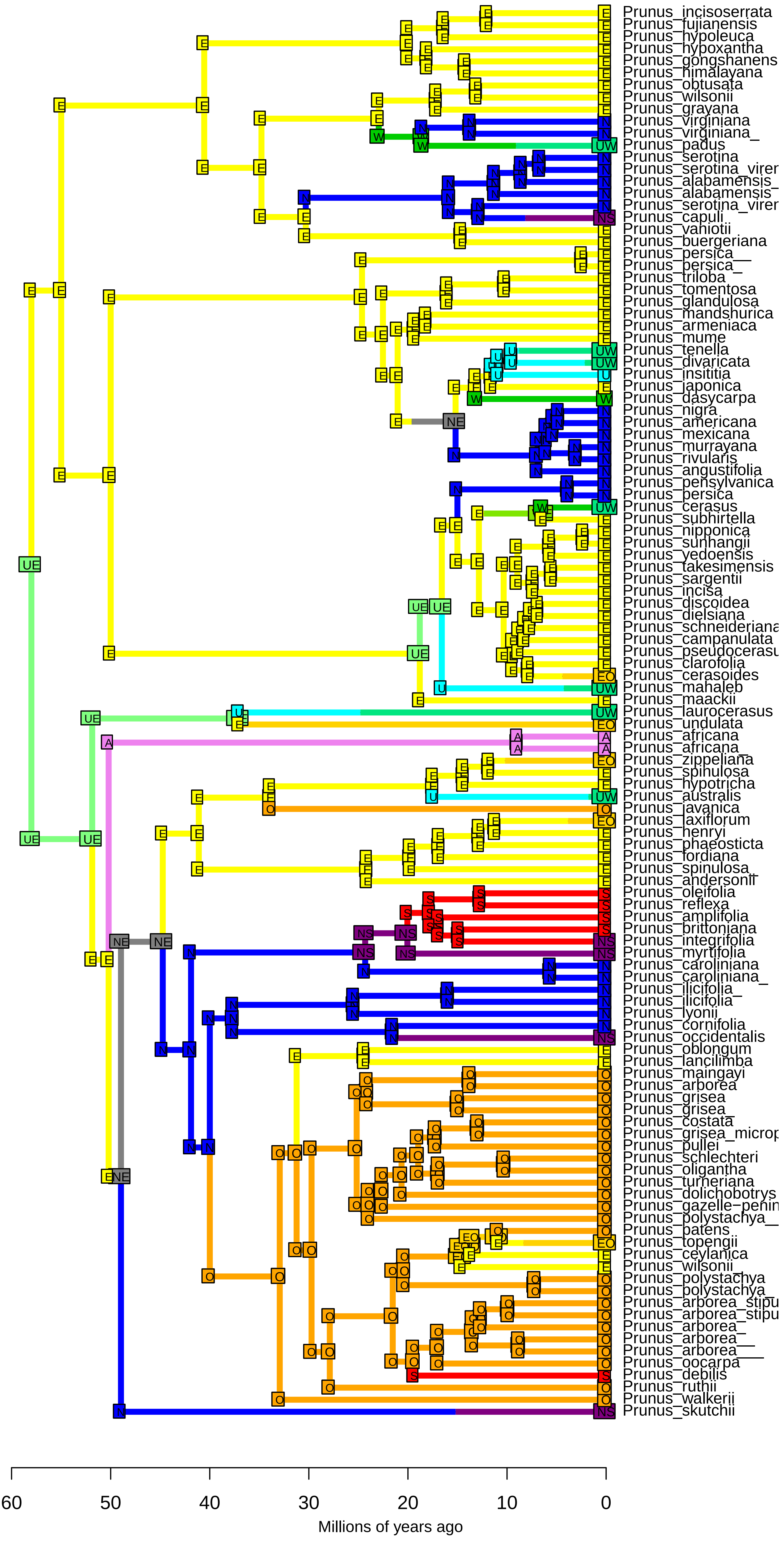

**DEC−J_M3_timestrat − Stochastic Map #33/50**
**ancstates: global optim, 2 areas max. d=0.0016; e=0; j=0.0148; LnL=−173.29**

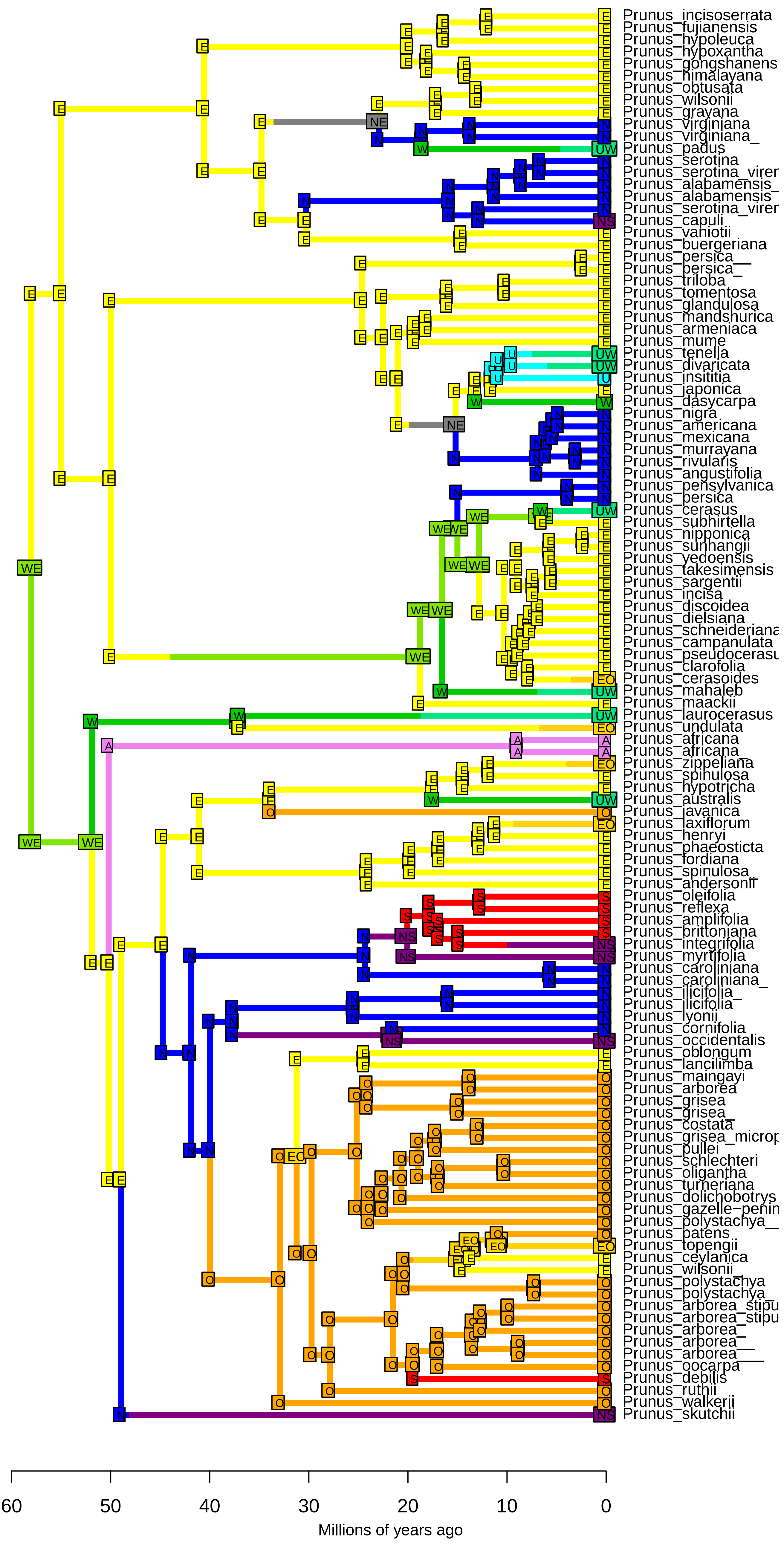

**DEC−J_M3_timestrat − Stochastic Map #34/50**
**ancstates: global optim, 2 areas max. d=0.0016; e=0; j=0.0148; LnL=−173.29**

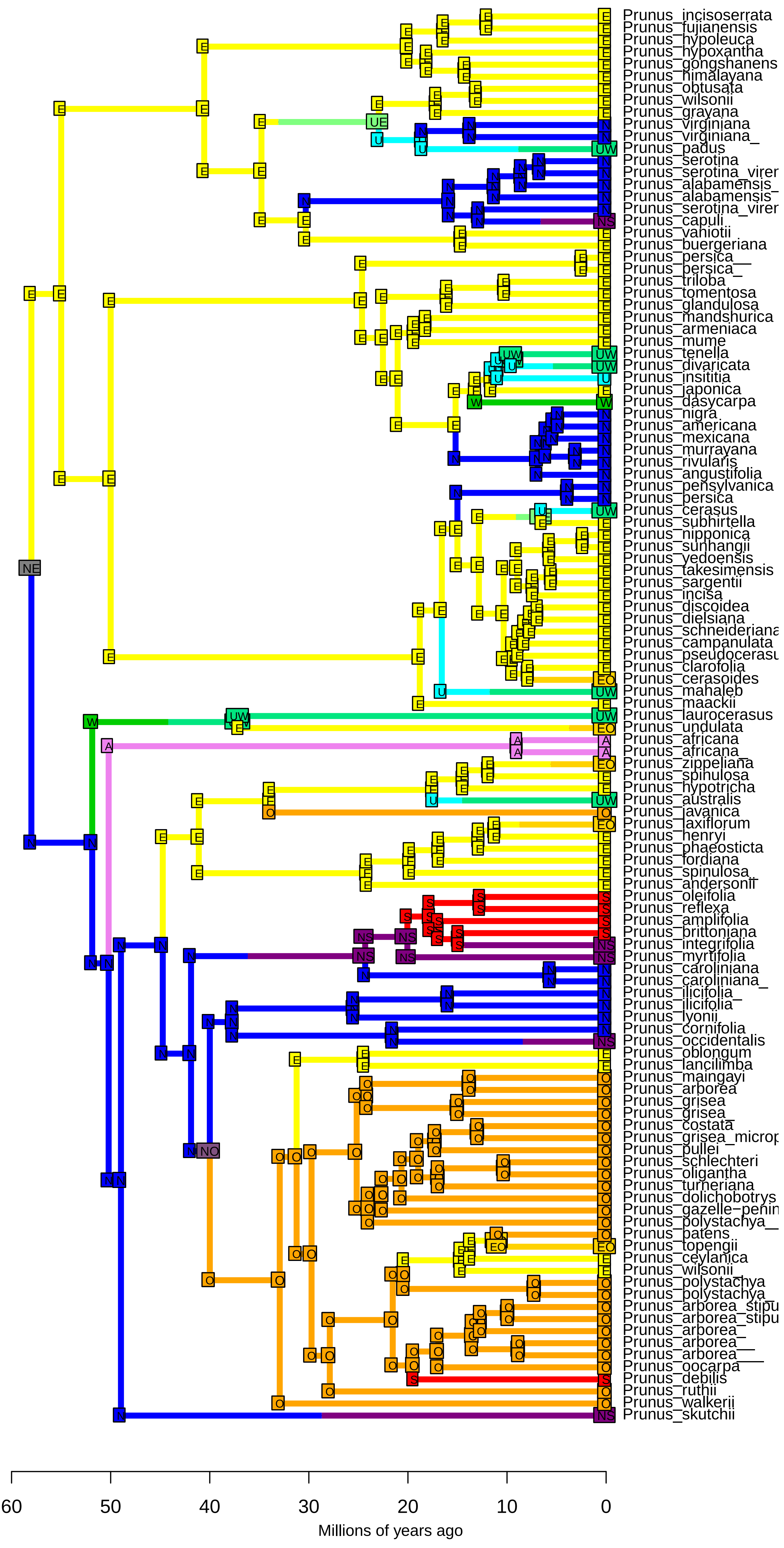

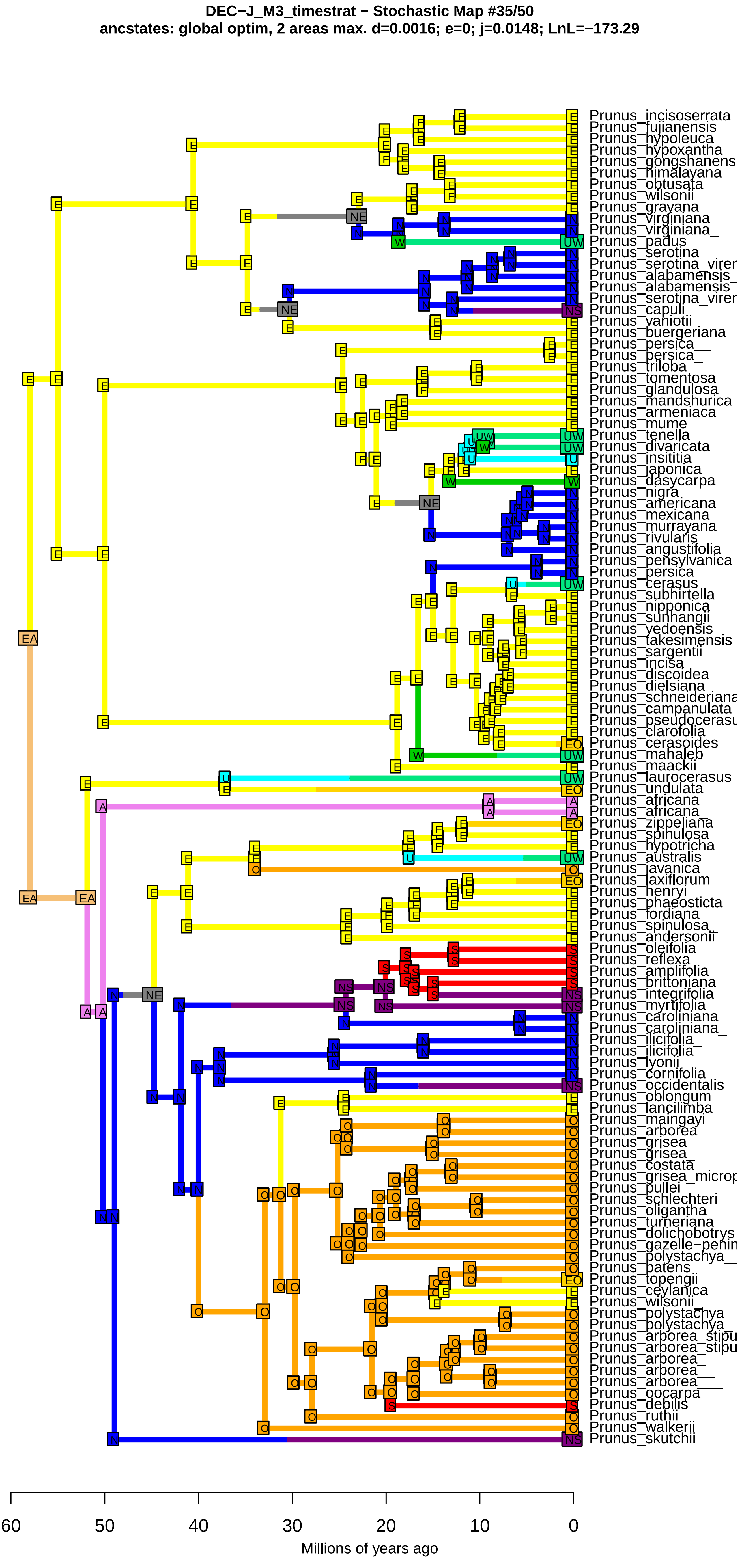
DEC−J_M3_timestrat − Stochastic Map #35/50
ancstates: global optim, 2 areas max. d=0.0016; e=0; j=0.0148; LnL=−173.29
Prunus_incisoserrata
Prunus_fujianensis
Prunus_hypoleuca
Prunus_hypoxantha
Prunus_gongshanens
Prunus_himalayana
Prunus_obtusata
Prunus_wilsonii
Prunus_grayana
Prunus_virginiana
Prunus_virginiana_
Prunus_padus
Prunus_serotina
Prunus_serotina_virer
Prunus_alabamensis_
Prunus_alabamensis
Prunus_serotina_virer
Prunus_capuli
Prunus_vaniotii
Prunus_buergeriana
Prunus_persica__
Prunus_persica_
Prunus_triloba
Prunus_tomentosa
Prunus_glandulosa
Prunus_mandshurica
Prunus_armeniaca
Prunus_mume
Prunus_tenella
Prunus_divaricata
Prunus_insititia
Prunus_japonica
Prunus_dasycarpa
Prunus_nigra
Prunus_americana
Prunus_mexicana
Prunus_murrayana
Prunus_rivularis
Prunus_angustifolia
Prunus_pensylvanica
Prunus_persica
Prunus_cerasus
Prunus_subhirtella
Prunus_nipponica
Prunus_sunhangii
Prunus_yedoensis
Prunus_takesimensis
Prunus_sargentii
Prunus_incisa
Prunus_discoidea
Prunus_dielsiana
Prunus_schneideriana
Prunus_campanulata
Prunus_pseudocerasu
Prunus_clarofolia
Prunus_cerasoides
Prunus_mahaleb
Prunus_maackii
Prunus_laurocerasus
Prunus_undulata
Prunus_africana
Prunus_africana_
Prunus_zippeliana
Prunus_spinulosa
Prunus_hypotricha
Prunus_australis
Prunus_javanica
Prunus_laxiflorum
Prunus_henryi
Prunus_phaeosticta
Prunus_fordiana
Prunus_spinulosa_
Prunus_andersonii
Prunus_oleifolia
Prunus_reflexa
Prunus_amplifolia
Prunus_brittoniana
Prunus_integrifolia
Prunus_myrtifolia
Prunus_caroliniana
Prunus_caroliniana_
Prunus_ilicifolia_
Prunus_ilicifolia
Prunus_lyonii
Prunus_cornifolia
Prunus_occidentalis
Prunus_oblongum
Prunus_lancilimba
Prunus_maingayi
Prunus_arborea
Prunus_grisea
Prunus_grisea_
Prunus_costata
Prunus_grisea_microp
Prunus_pullei
Prunus_schlechteri
Prunus_oligantha
Prunus_turneriana
Prunus_dolichobotrys
Prunus_gazelle−penin
Prunus_polystachya__
Prunus_patens
Prunus_topengii
Prunus_ceylanica
Prunus_wilsonii_
Prunus_polystachya
Prunus_polystachya_
Prunus_arborea_stipu
Prunus_arborea_stipu
Prunus_arborea_
Prunus_arborea__
Prunus_arborea___
Prunus_oocarpa
Prunus_debilis
Prunus_ruthii
Prunus_walkerii
Prunus_skutchii
60
50
40
30
20
10
0
Millions of years ago

**DEC−J_M3_timestrat − Stochastic Map #36/50**
**ancstates: global optim, 2 areas max. d=0.0016; e=0; j=0.0148; LnL=−173.29**

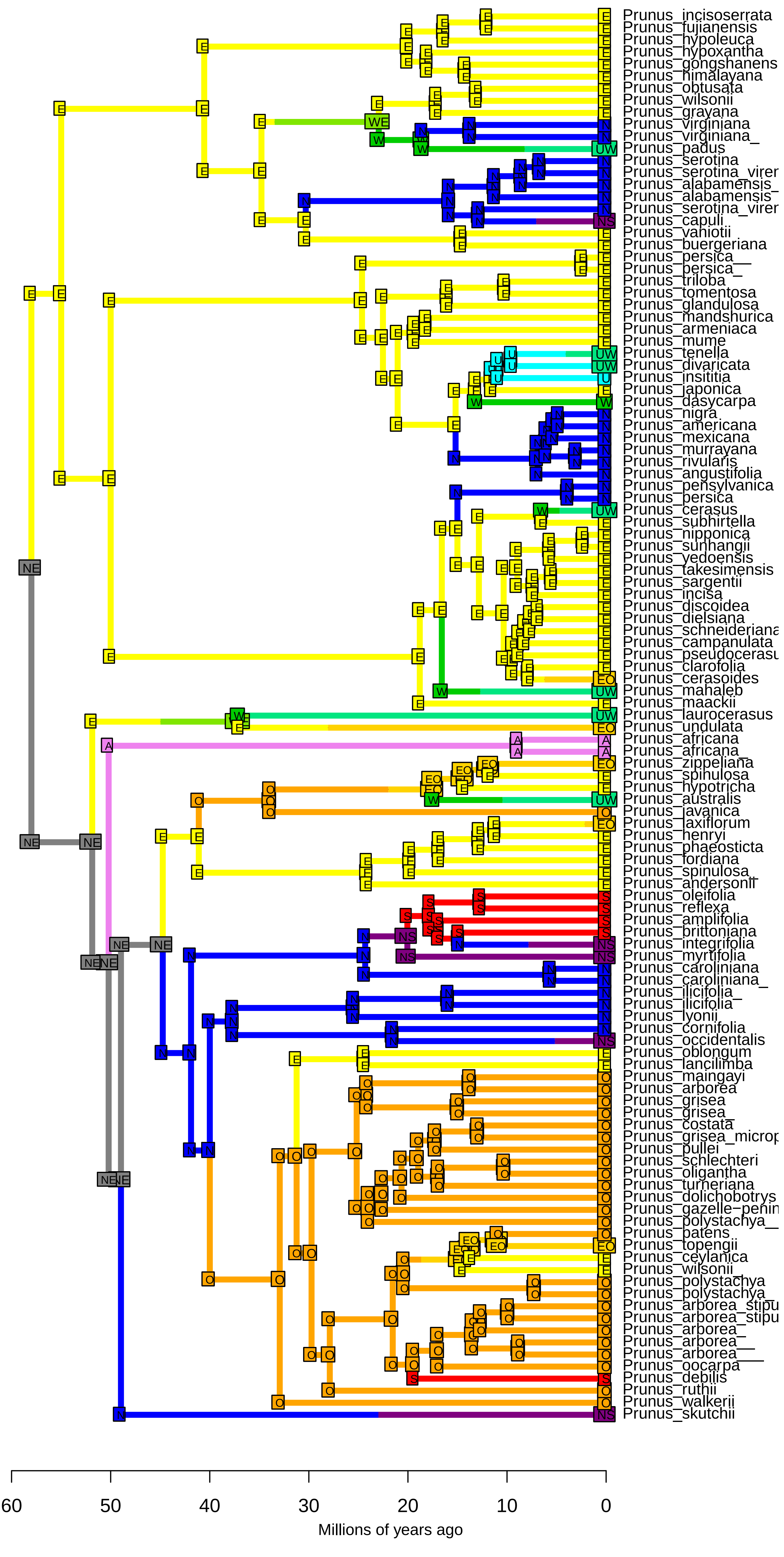

**DEC−J_M3_timestrat − Stochastic Map #37/50**
**ancstates: global optim, 2 areas max. d=0.0016; e=0; j=0.0148; LnL=−173.29**

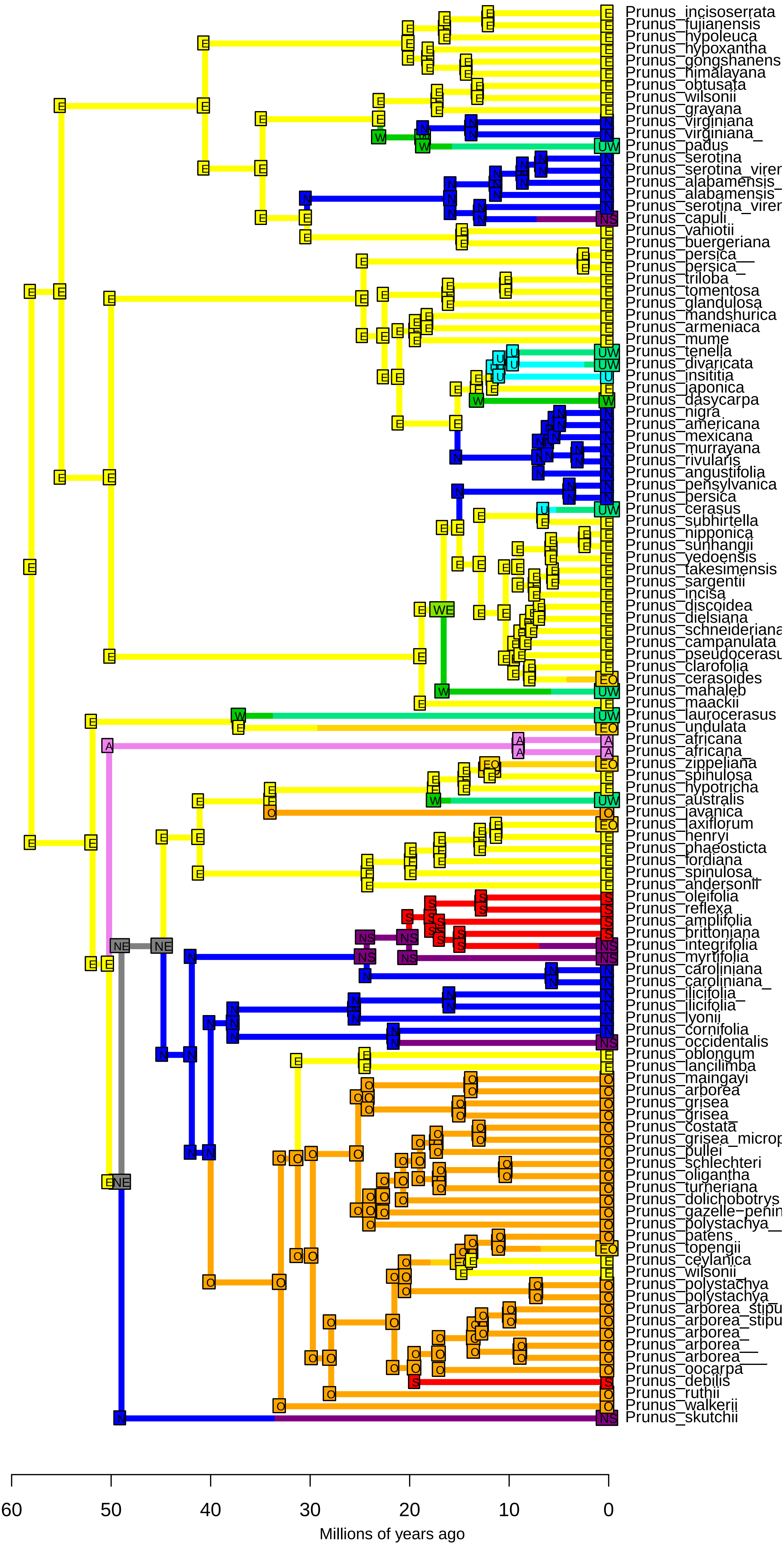

**DEC−J_M3_timestrat − Stochastic Map #38/50**
**ancstates: global optim, 2 areas max. d=0.0016; e=0; j=0.0148; LnL=−173.29**

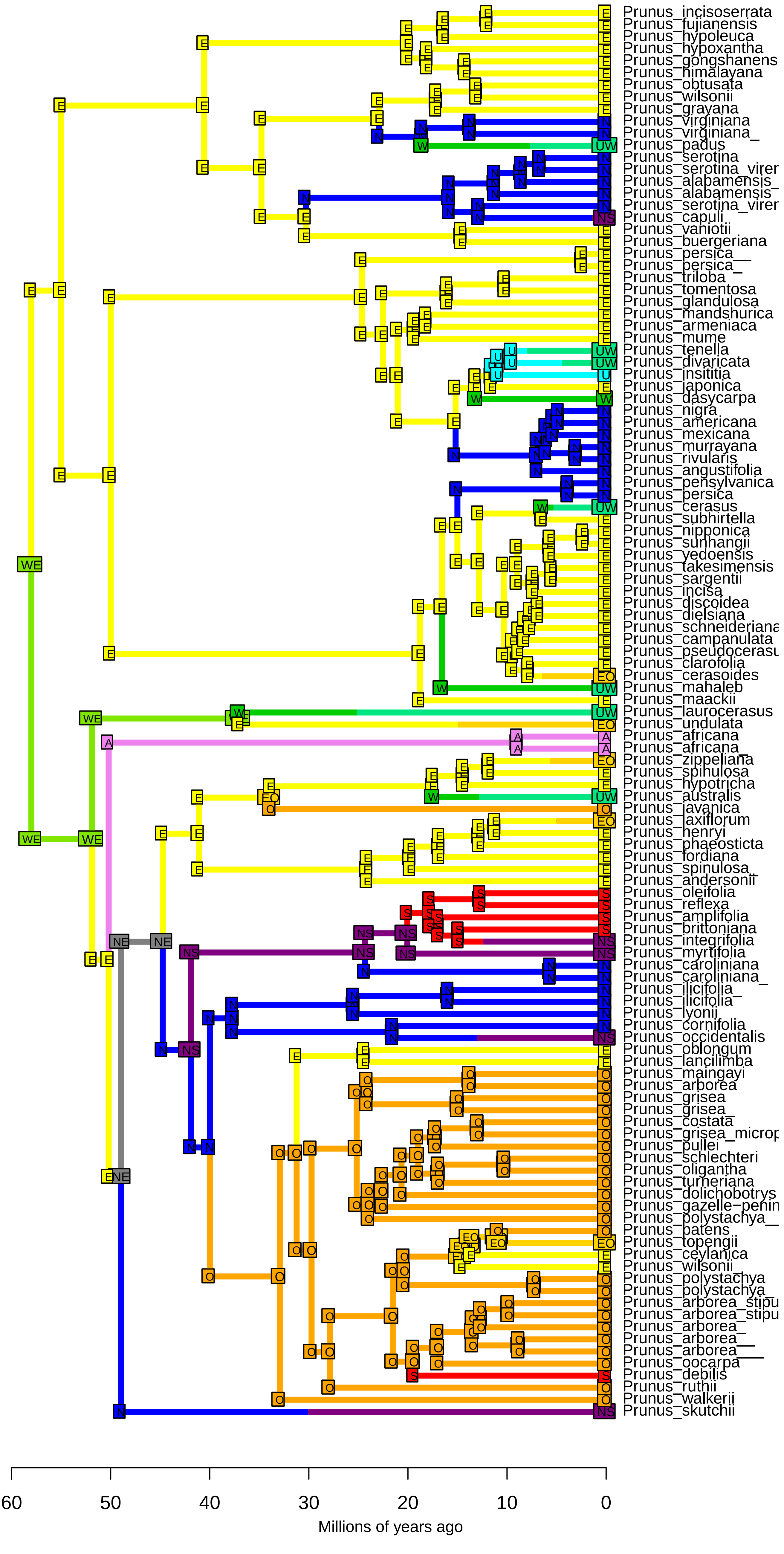

**DEC−J_M3_timestrat − Stochastic Map #39/50**
**ancstates: global optim, 2 areas max. d=0.0016; e=0; j=0.0148; LnL=−173.29**

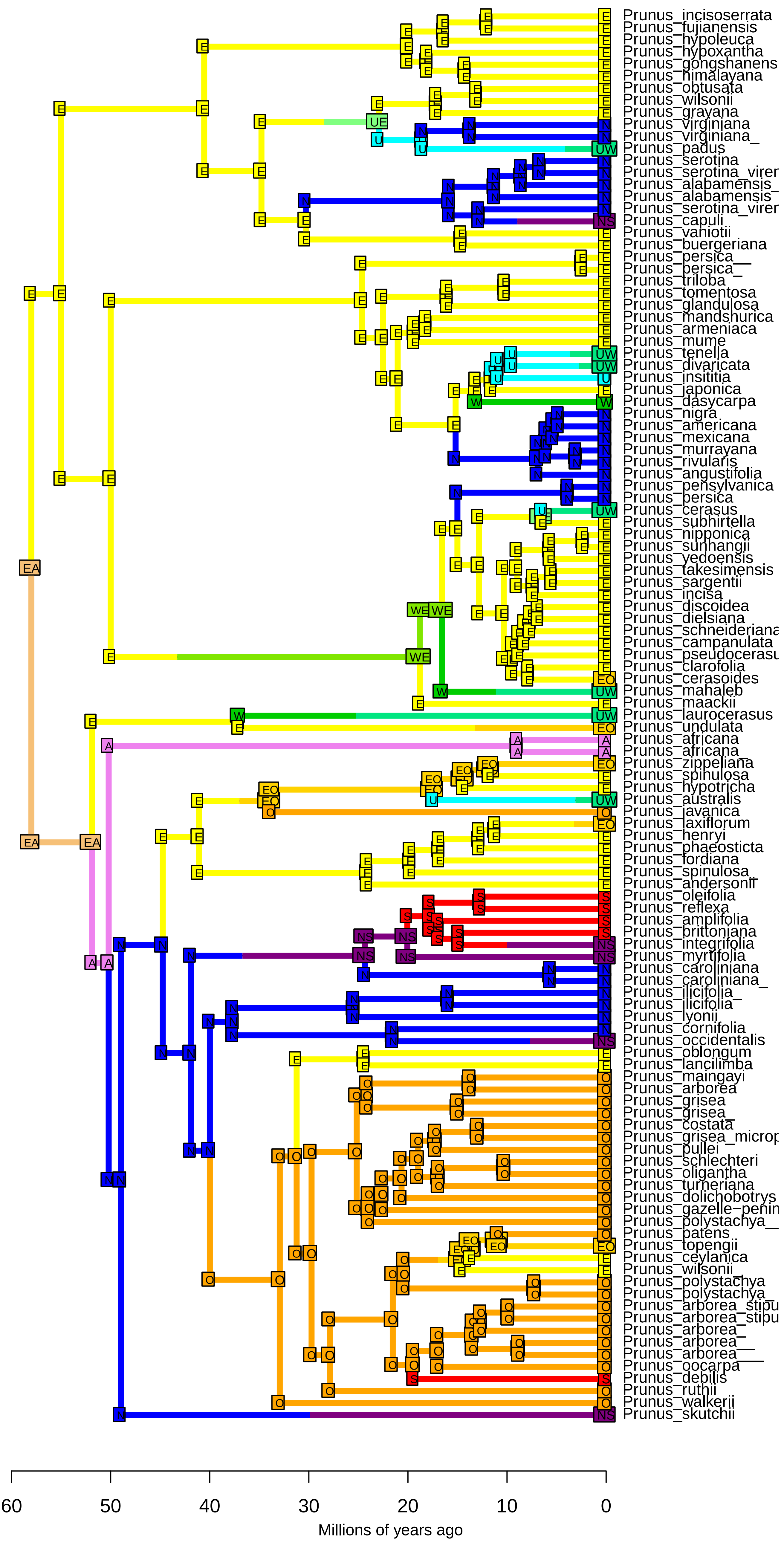

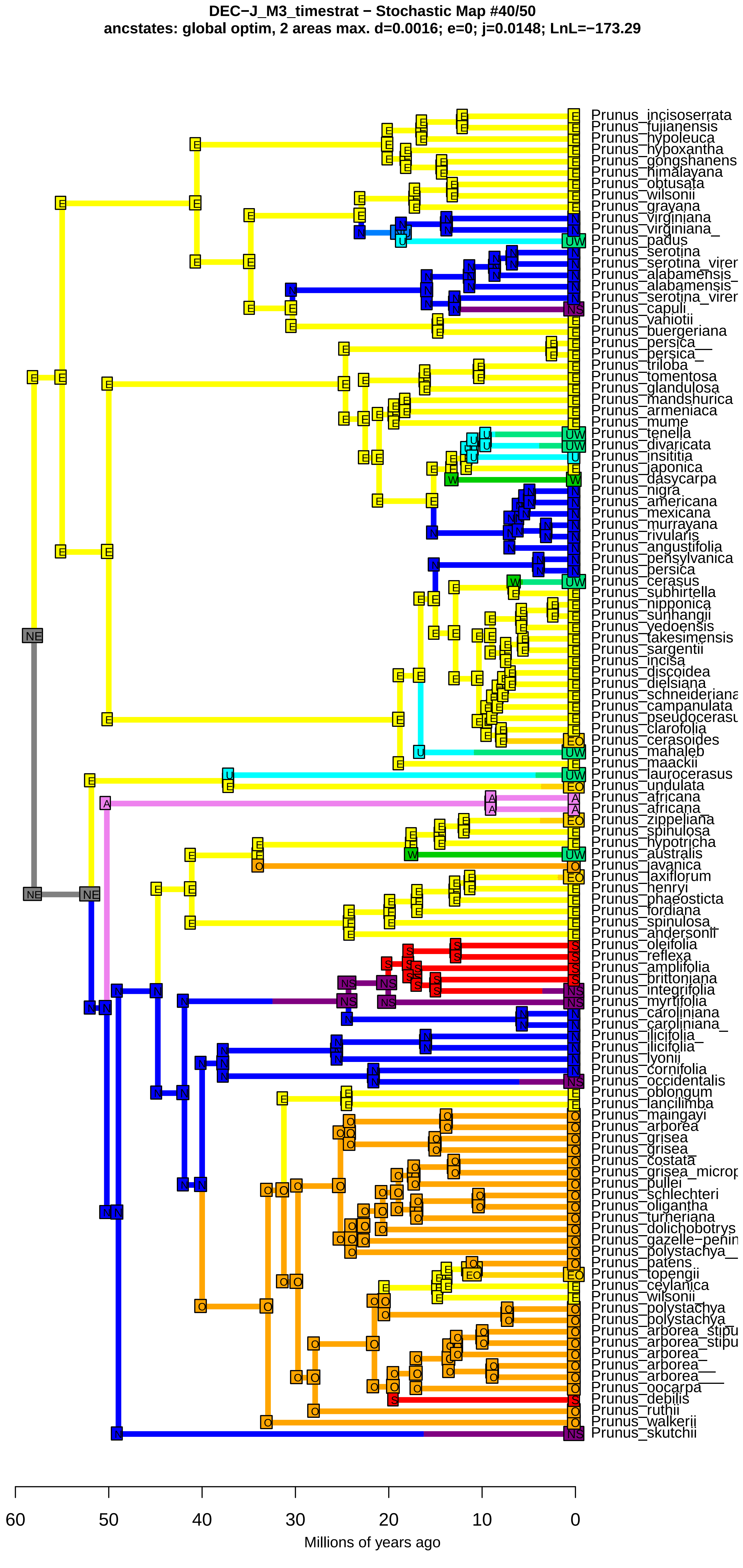
DEC−J_M3_timestrat − Stochastic Map #40/50
ancstates: global optim, 2 areas max. d=0.0016; e=0; j=0.0148; LnL=−173.29
Prunus_incisoserrata
Prunus_fujianensis
Prunus_hypoleuca
Prunus_hypoxantha
Prunus_gongshanens
Prunus_himalayana
Prunus_obtusata
Prunus_wilsonii
Prunus_grayana
Prunus_virginiana
Prunus_virginiana_
Prunus_padus
Prunus_serotina
Prunus_serotina_virer
Prunus_alabamensis_
Prunus_alabamensis
Prunus_serotina_virer
Prunus_capuli
Prunus_vaniotii
Prunus_buergeriana
Prunus_persica__
Prunus_persica_
Prunus_triloba
Prunus_tomentosa
Prunus_glandulosa
Prunus_mandshurica
Prunus_armeniaca
Prunus_mume
Prunus_tenella
Prunus_divaricata
Prunus_insititia
Prunus_japonica
Prunus_dasycarpa
Prunus_nigra
Prunus_americana
Prunus_mexicana
Prunus_murrayana
Prunus_rivularis
Prunus_angustifolia
Prunus_pensylvanica
Prunus_persica
Prunus_cerasus
Prunus_subhirtella
Prunus_nipponica
Prunus_sunhangii
Prunus_yedoensis
Prunus_takesimensis
Prunus_sargentii
Prunus_incisa
Prunus_discoidea
Prunus_dielsiana
Prunus_schneideriana
Prunus_campanulata
Prunus_pseudocerasu
Prunus_clarofolia
Prunus_cerasoides
Prunus_mahaleb
Prunus_maackii
Prunus_laurocerasus
Prunus_undulata
Prunus_africana
Prunus_africana_
Prunus_zippeliana
Prunus_spinulosa
Prunus_hypotricha
Prunus_australis
Prunus_javanica
Prunus_laxiflorum
Prunus_henryi
Prunus_phaeosticta
Prunus_fordiana
Prunus_spinulosa_
Prunus_andersonii
Prunus_oleifolia
Prunus_reflexa
Prunus_amplifolia
Prunus_brittoniana
Prunus_integrifolia
Prunus_myrtifolia
Prunus_caroliniana
Prunus_caroliniana_
Prunus_ilicifolia_
Prunus_ilicifolia
Prunus_lyonii
Prunus_cornifolia
Prunus_occidentalis
Prunus_oblongum
Prunus_lancilimba
Prunus_maingayi
Prunus_arborea
Prunus_grisea
Prunus_grisea_
Prunus_costata
Prunus_grisea_microp
Prunus_pullei
Prunus_schlechteri
Prunus_oligantha
Prunus_turneriana
Prunus_dolichobotrys
Prunus_gazelle−penin
Prunus_polystachya__
Prunus_patens
Prunus_topengii
Prunus_ceylanica
Prunus_wilsonii_
Prunus_polystachya
Prunus_polystachya_
Prunus_arborea_stipu
Prunus_arborea_stipu
Prunus_arborea_
Prunus_arborea__
Prunus_arborea___
Prunus_oocarpa
Prunus_debilis
Prunus_ruthii
Prunus_walkerii
Prunus_skutchii
60
50
40
30
20
10
0
Millions of years ago

**DEC−J_M3_timestrat − Stochastic Map #41/50**
**ancstates: global optim, 2 areas max. d=0.0016; e=0; j=0.0148; LnL=−173.29**

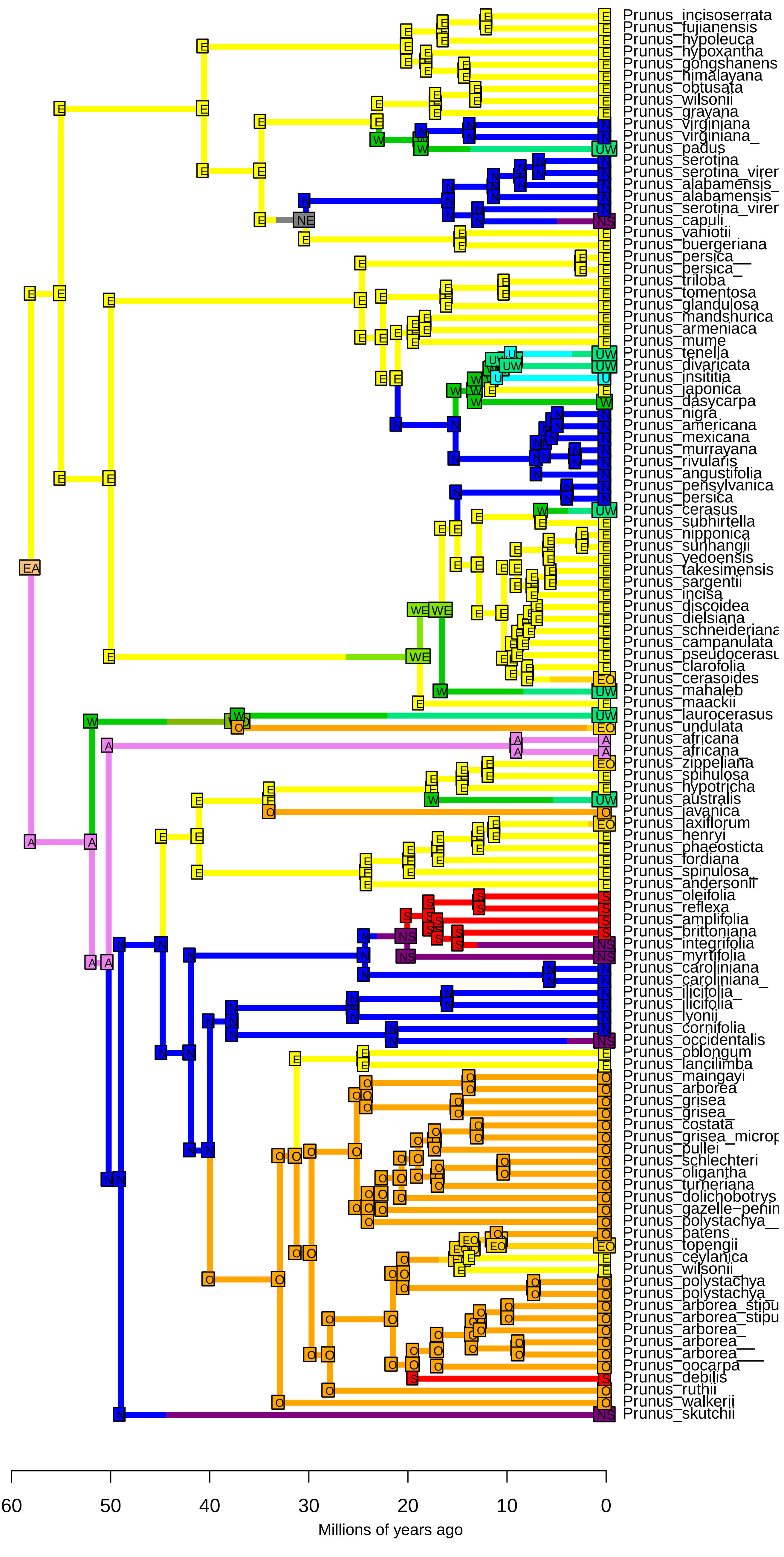

**DEC−J_M3_timestrat − Stochastic Map #42/50**
**ancstates: global optim, 2 areas max. d=0.0016; e=0; j=0.0148; LnL=−173.29**

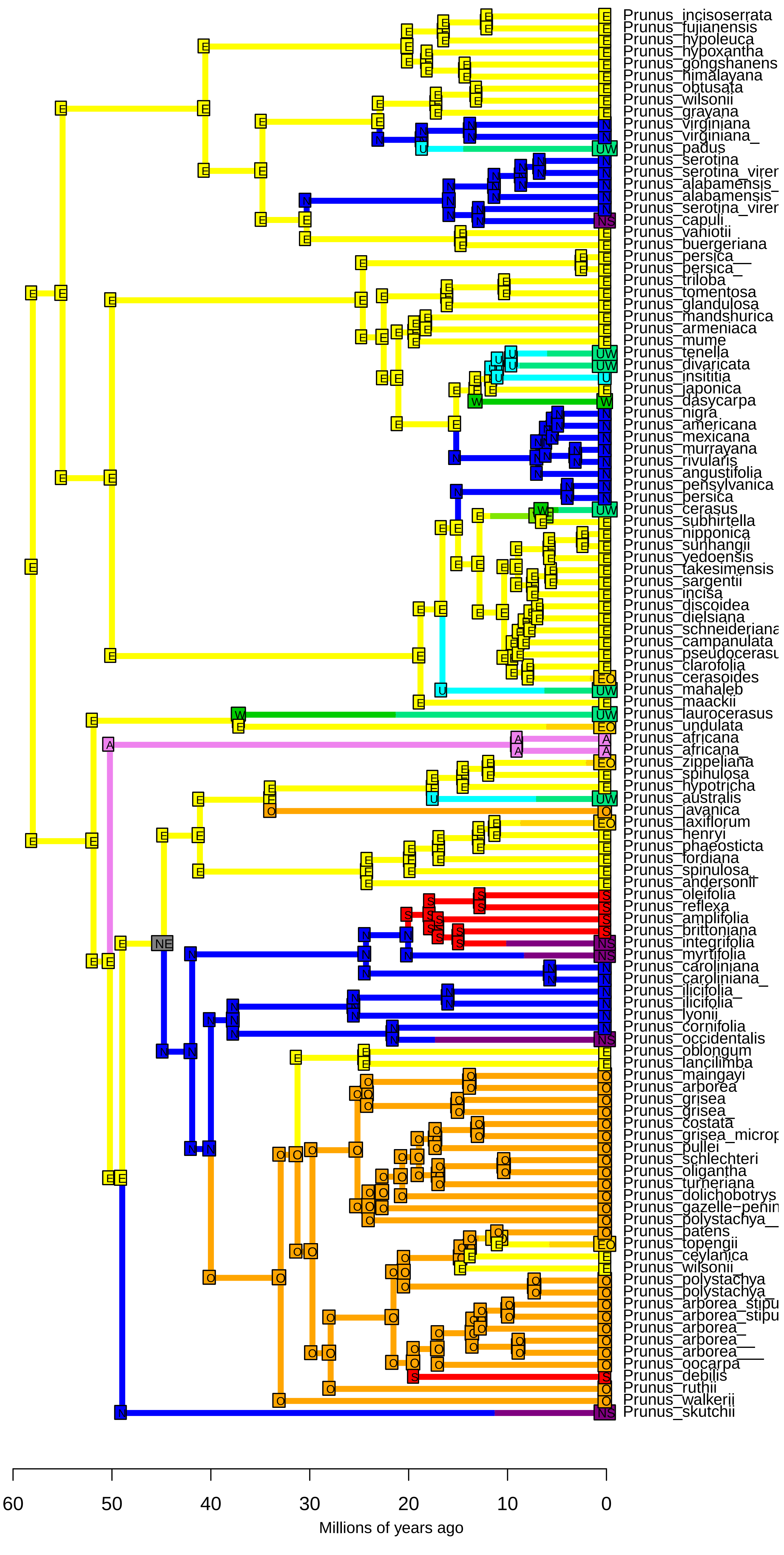

**DEC−J_M3_timestrat − Stochastic Map #43/50**
**ancstates: global optim, 2 areas max. d=0.0016; e=0; j=0.0148; LnL=−173.29**

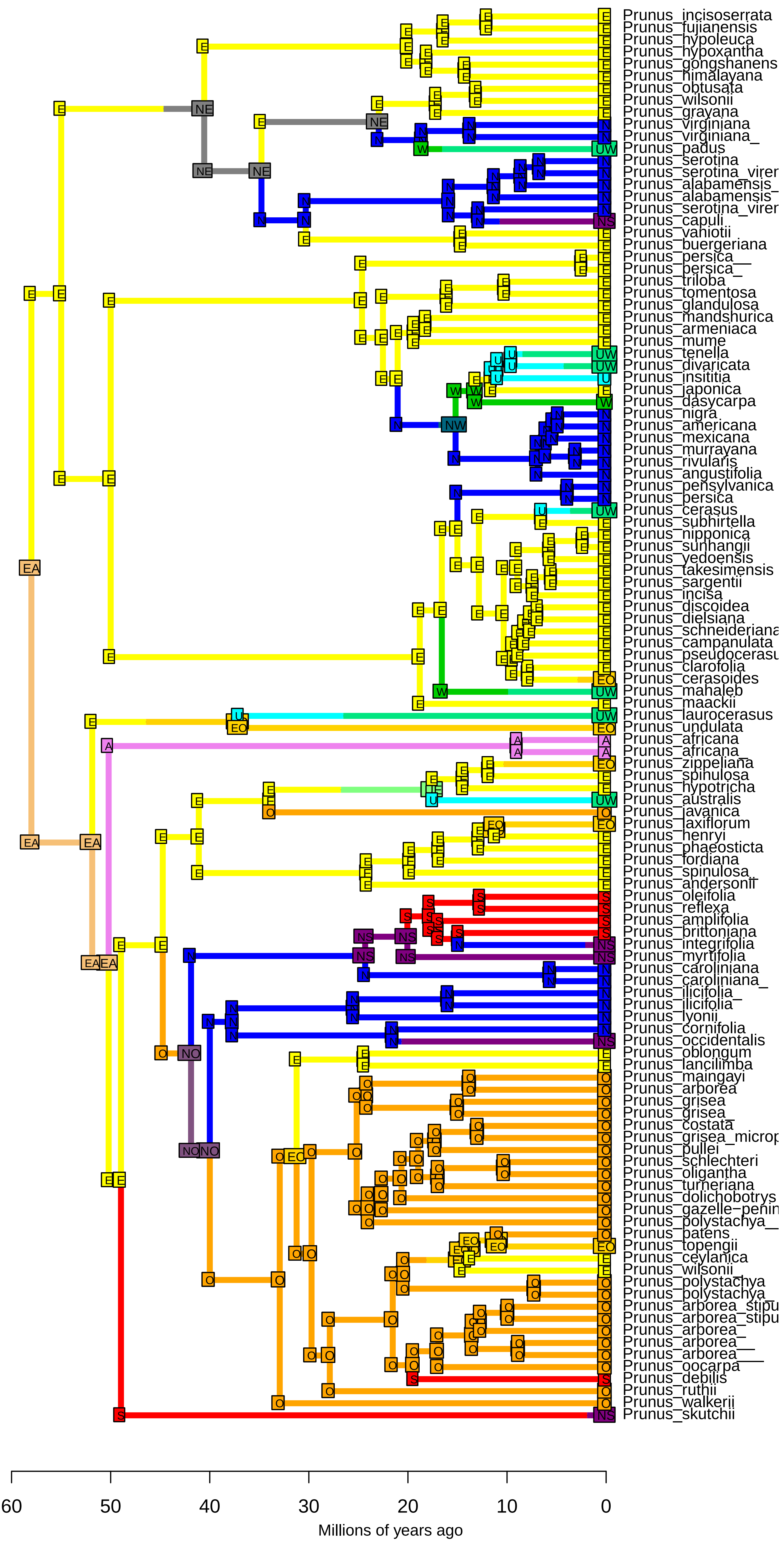

**DEC−J_M3_timestrat − Stochastic Map #44/50**
**ancstates: global optim, 2 areas max. d=0.0016; e=0; j=0.0148; LnL=−173.29**

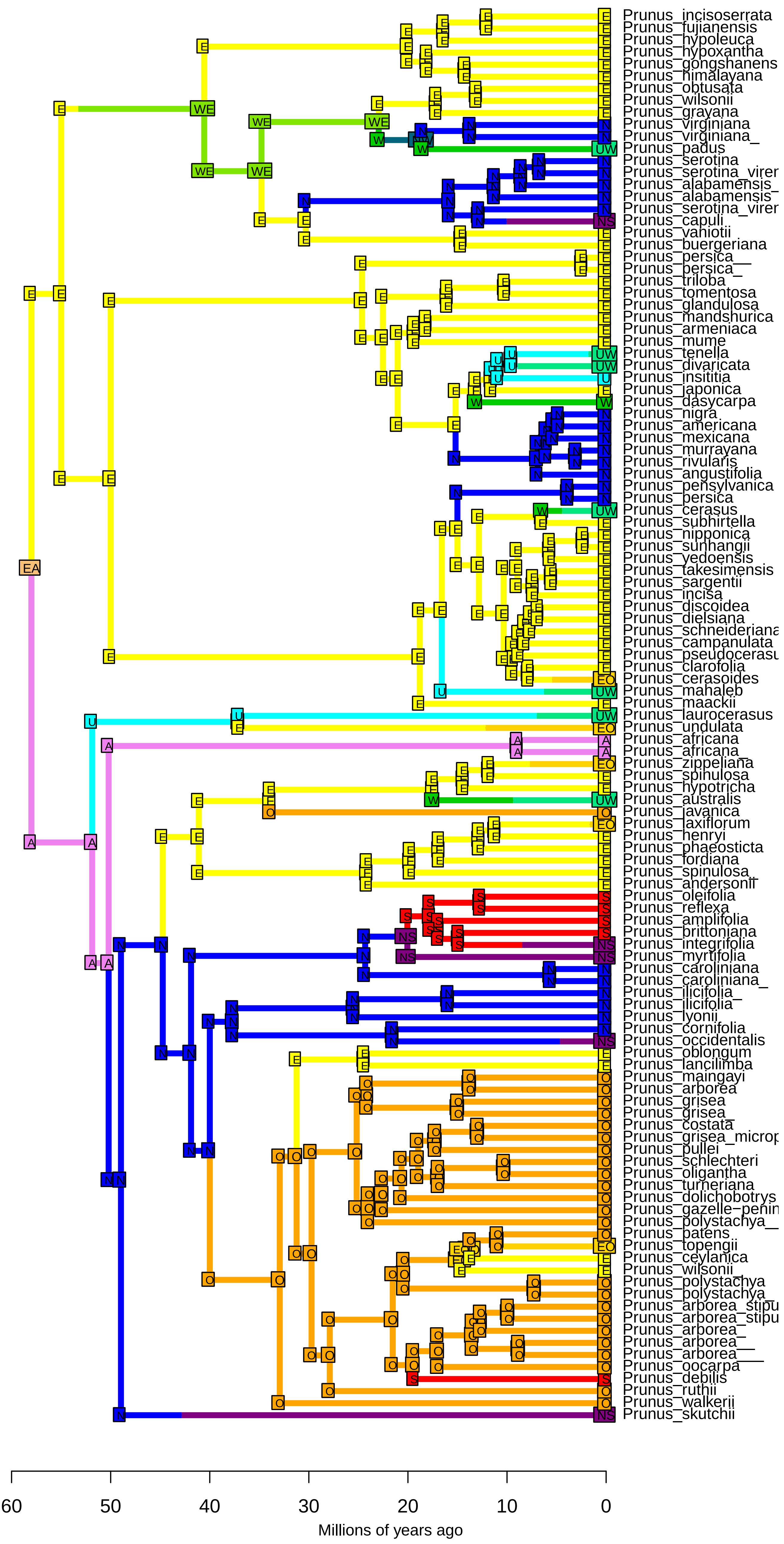

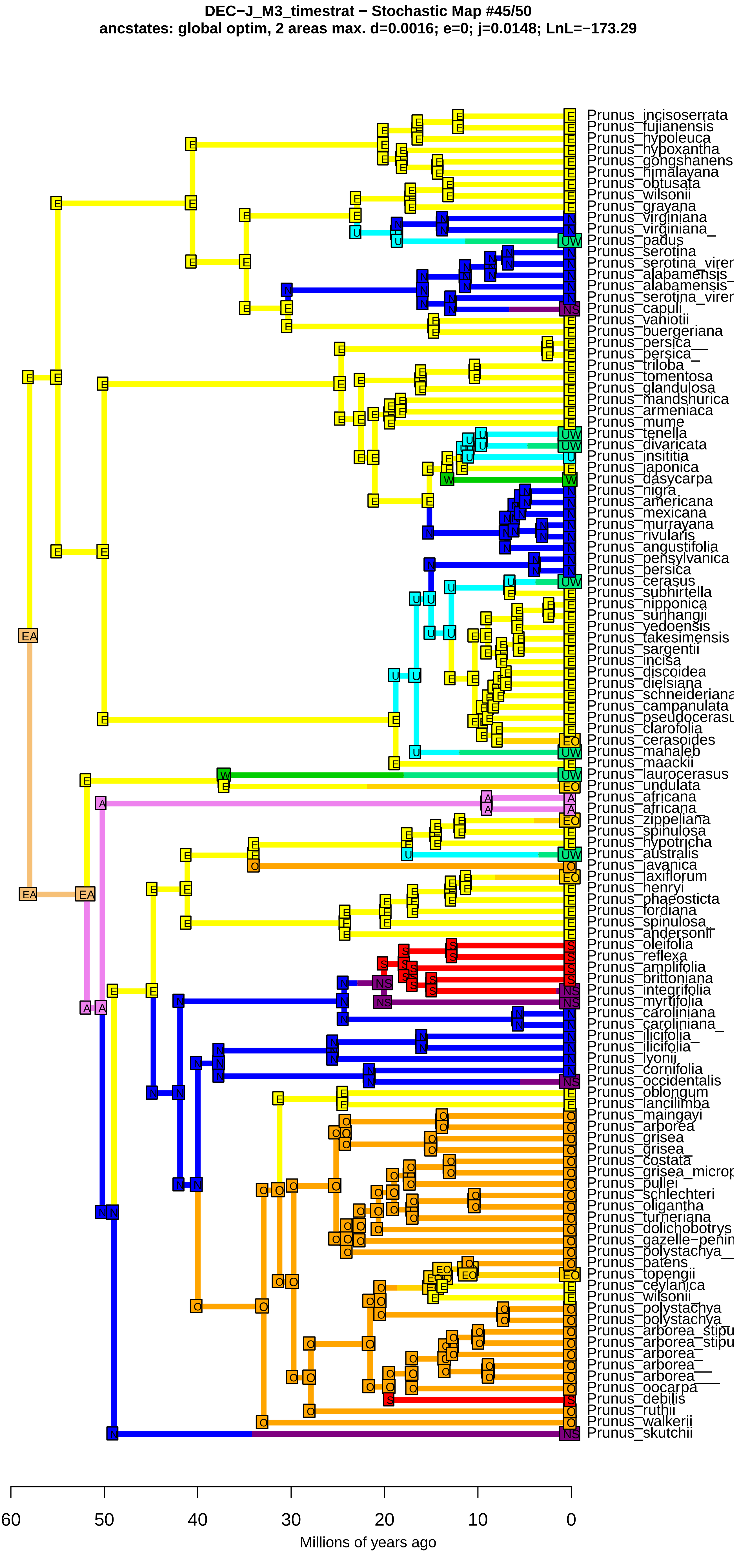
DEC−J_M3_timestrat − Stochastic Map #45/50
ancstates: global optim, 2 areas max. d=0.0016; e=0; j=0.0148; LnL=−173.29
Prunus_incisoserrata
Prunus_fujianensis
Prunus_hypoleuca
Prunus_hypoxantha
Prunus_gongshanens
Prunus_himalayana
Prunus_obtusata
Prunus_wilsonii
Prunus_grayana
Prunus_virginiana
Prunus_virginiana_
Prunus_padus
Prunus_serotina
Prunus_serotina_virer
Prunus_alabamensis_
Prunus_alabamensis
Prunus_serotina_virer
Prunus_capuli
Prunus_vaniotii
Prunus_buergeriana
Prunus_persica__
Prunus_persica_
Prunus_triloba
Prunus_tomentosa
Prunus_glandulosa
Prunus_mandshurica
Prunus_armeniaca
Prunus_mume
Prunus_tenella
Prunus_divaricata
Prunus_insititia
Prunus_japonica
Prunus_dasycarpa
Prunus_nigra
Prunus_americana
Prunus_mexicana
Prunus_murrayana
Prunus_rivularis
Prunus_angustifolia
Prunus_pensylvanica
Prunus_persica
Prunus_cerasus
Prunus_subhirtella
Prunus_nipponica
Prunus_sunhangii
Prunus_yedoensis
Prunus_takesimensis
Prunus_sargentii
Prunus_incisa
Prunus_discoidea
Prunus_dielsiana
Prunus_schneideriana
Prunus_campanulata
Prunus_pseudocerasu
Prunus_clarofolia
Prunus_cerasoides
Prunus_mahaleb
Prunus_maackii
Prunus_laurocerasus
Prunus_undulata
Prunus_africana
Prunus_africana_
Prunus_zippeliana
Prunus_spinulosa
Prunus_hypotricha
Prunus_australis
Prunus_javanica
Prunus_laxiflorum
Prunus_henryi
Prunus_phaeosticta
Prunus_fordiana
Prunus_spinulosa_
Prunus_andersonii
Prunus_oleifolia
Prunus_reflexa
Prunus_amplifolia
Prunus_brittoniana
Prunus_integrifolia
Prunus_myrtifolia
Prunus_caroliniana
Prunus_caroliniana_
Prunus_ilicifolia_
Prunus_ilicifolia
Prunus_lyonii
Prunus_cornifolia
Prunus_occidentalis
Prunus_oblongum
Prunus_lancilimba
Prunus_maingayi
Prunus_arborea
Prunus_grisea
Prunus_grisea_
Prunus_costata
Prunus_grisea_microp
Prunus_pullei
Prunus_schlechteri
Prunus_oligantha
Prunus_turneriana
Prunus_dolichobotrys
Prunus_gazelle−penin
Prunus_polystachya__
Prunus_patens
Prunus_topengii
Prunus_ceylanica
Prunus_wilsonii_
Prunus_polystachya
Prunus_polystachya_
Prunus_arborea_stipu
Prunus_arborea_stipu
Prunus_arborea_
Prunus_arborea__
Prunus_arborea___
Prunus_oocarpa
Prunus_debilis
Prunus_ruthii
Prunus_walkerii
Prunus_skutchii
60
50
40
30
20
10
0
Millions of years ago

**DEC−J_M3_timestrat − Stochastic Map #46/50**
**ancstates: global optim, 2 areas max. d=0.0016; e=0; j=0.0148; LnL=−173.29**

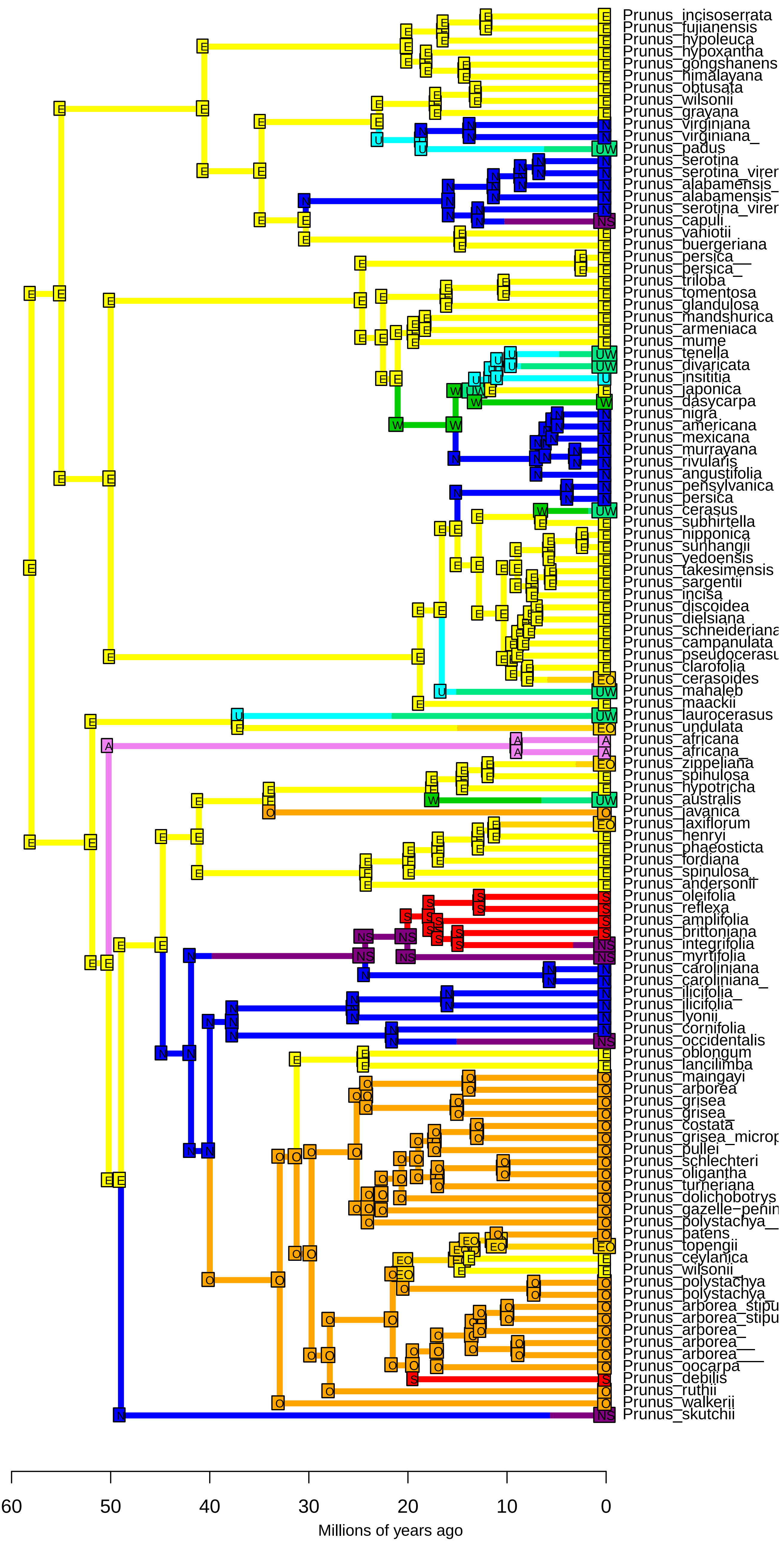

**DEC−J_M3_timestrat − Stochastic Map #47/50**
**ancstates: global optim, 2 areas max. d=0.0016; e=0; j=0.0148; LnL=−173.29**

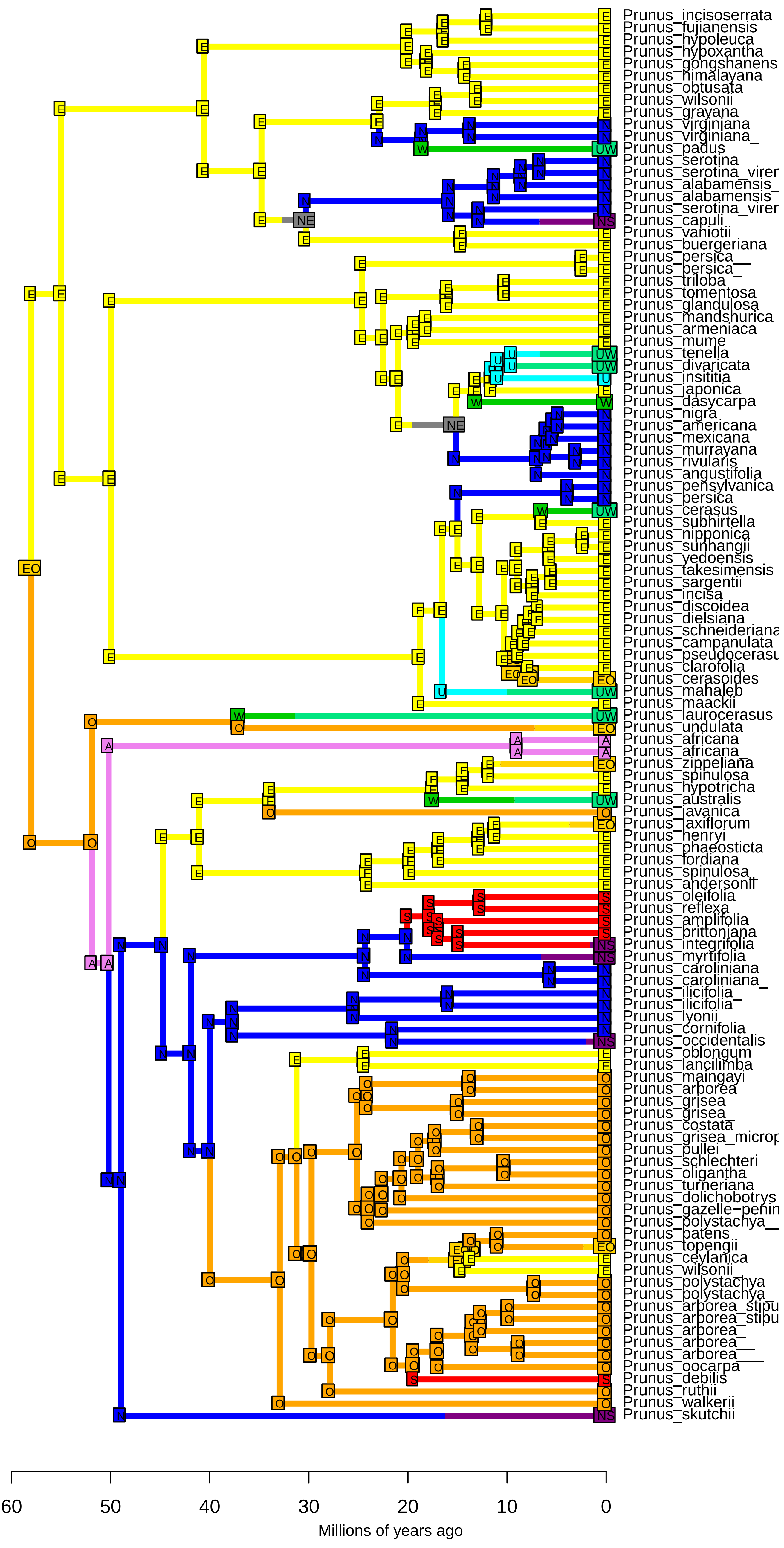

**DEC−J_M3_timestrat − Stochastic Map #48/50**
**ancstates: global optim, 2 areas max. d=0.0016; e=0; j=0.0148; LnL=−173.29**

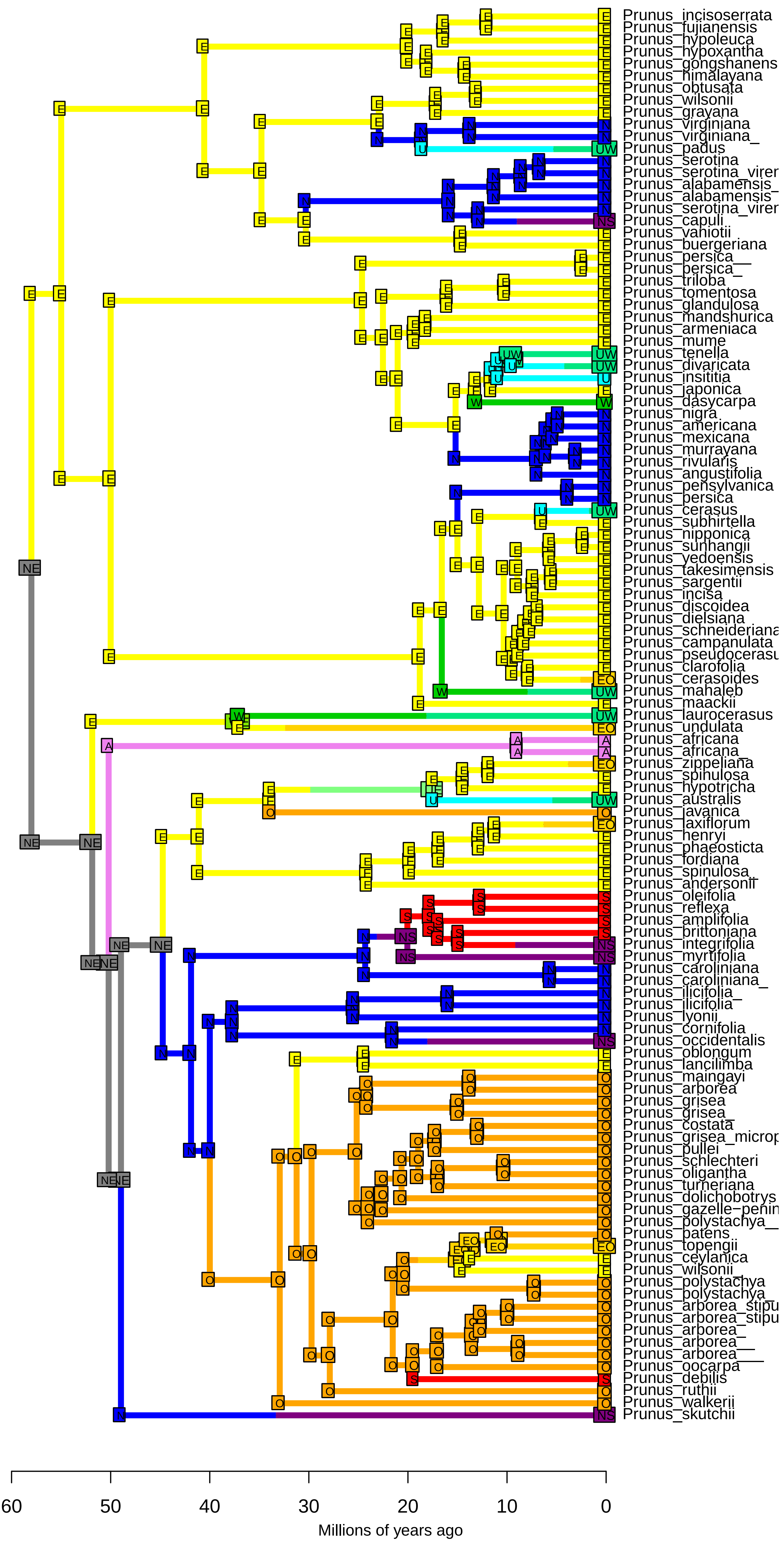

**DEC−J_M3_timestrat − Stochastic Map #49/50**
**ancstates: global optim, 2 areas max. d=0.0016; e=0; j=0.0148; LnL=−173.29**

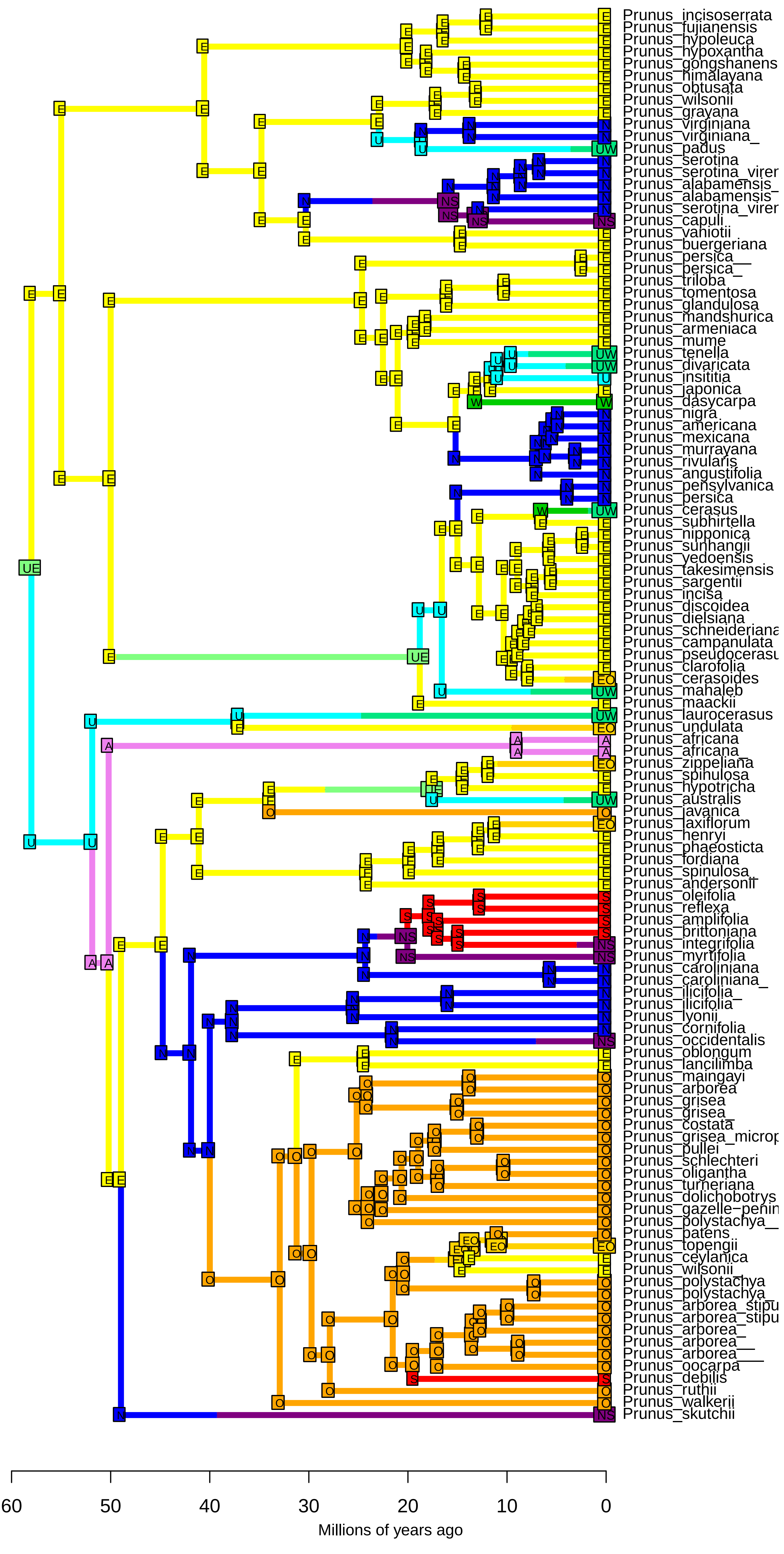

**DEC−J_M3_timestrat − Stochastic Map #50/50**
**ancstates: global optim, 2 areas max. d=0.0016; e=0; j=0.0148; LnL=−173.29**

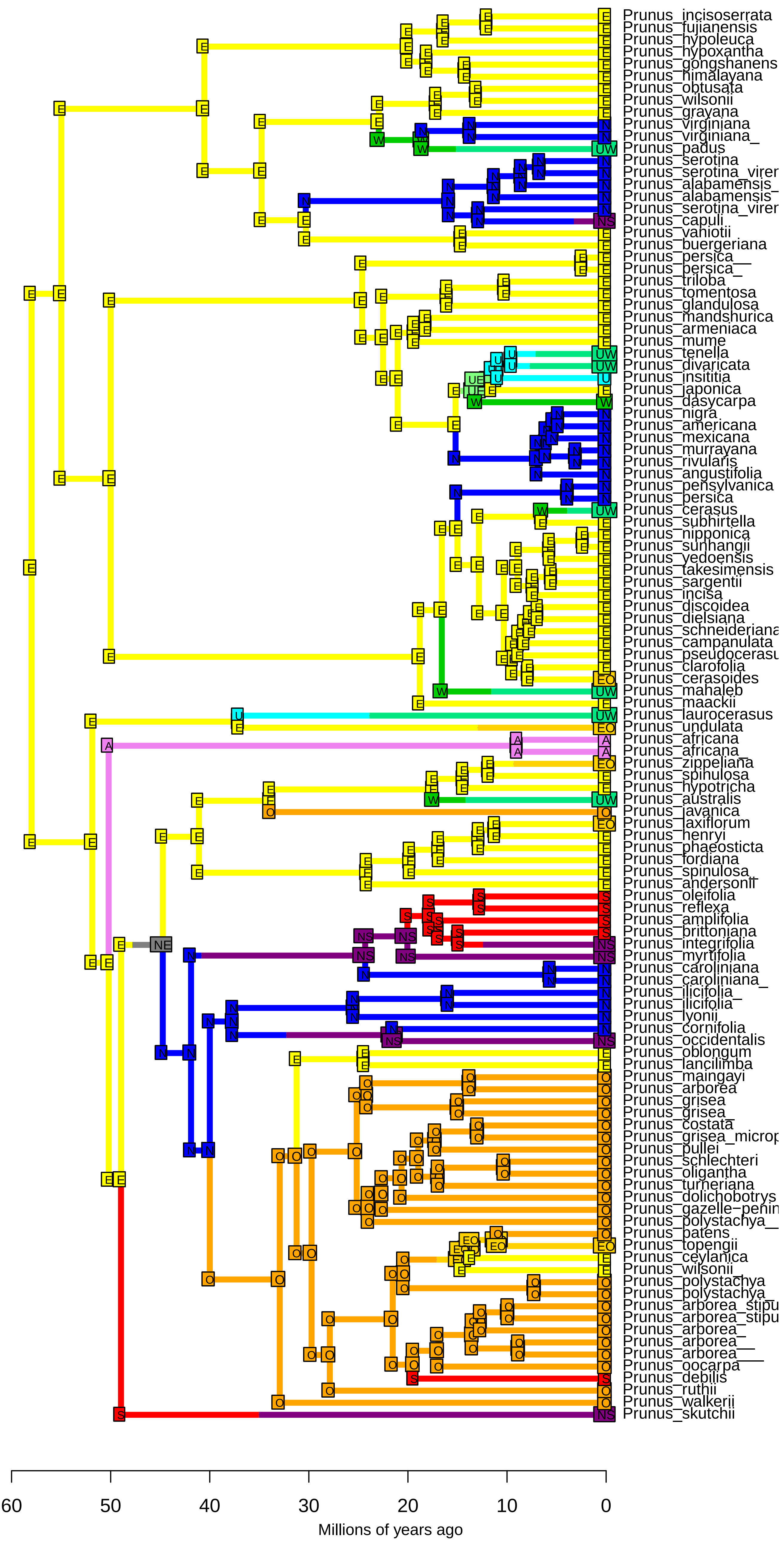

**BAYAL−J_M3_timestrat − Stochastic Map #1/50**
**ancstates: global optim, 4 areas max. d=0.0061; e=0.009; j=0; LnL=−231.76**

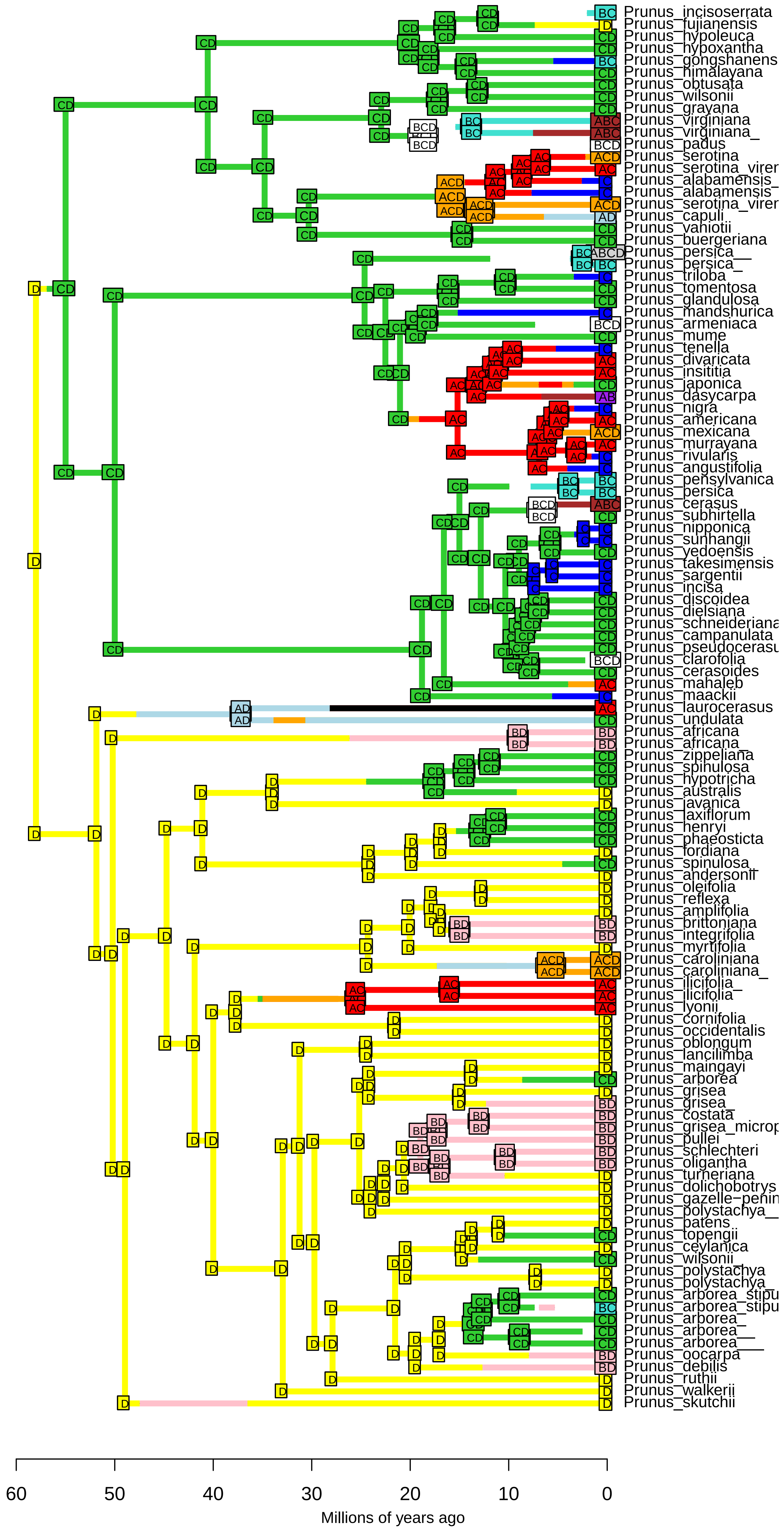

**BAYAL−J_M3_timestrat − Stochastic Map #2/50**
**ancstates: global optim, 4 areas max. d=0.0061; e=0.009; j=0; LnL=−231.76**

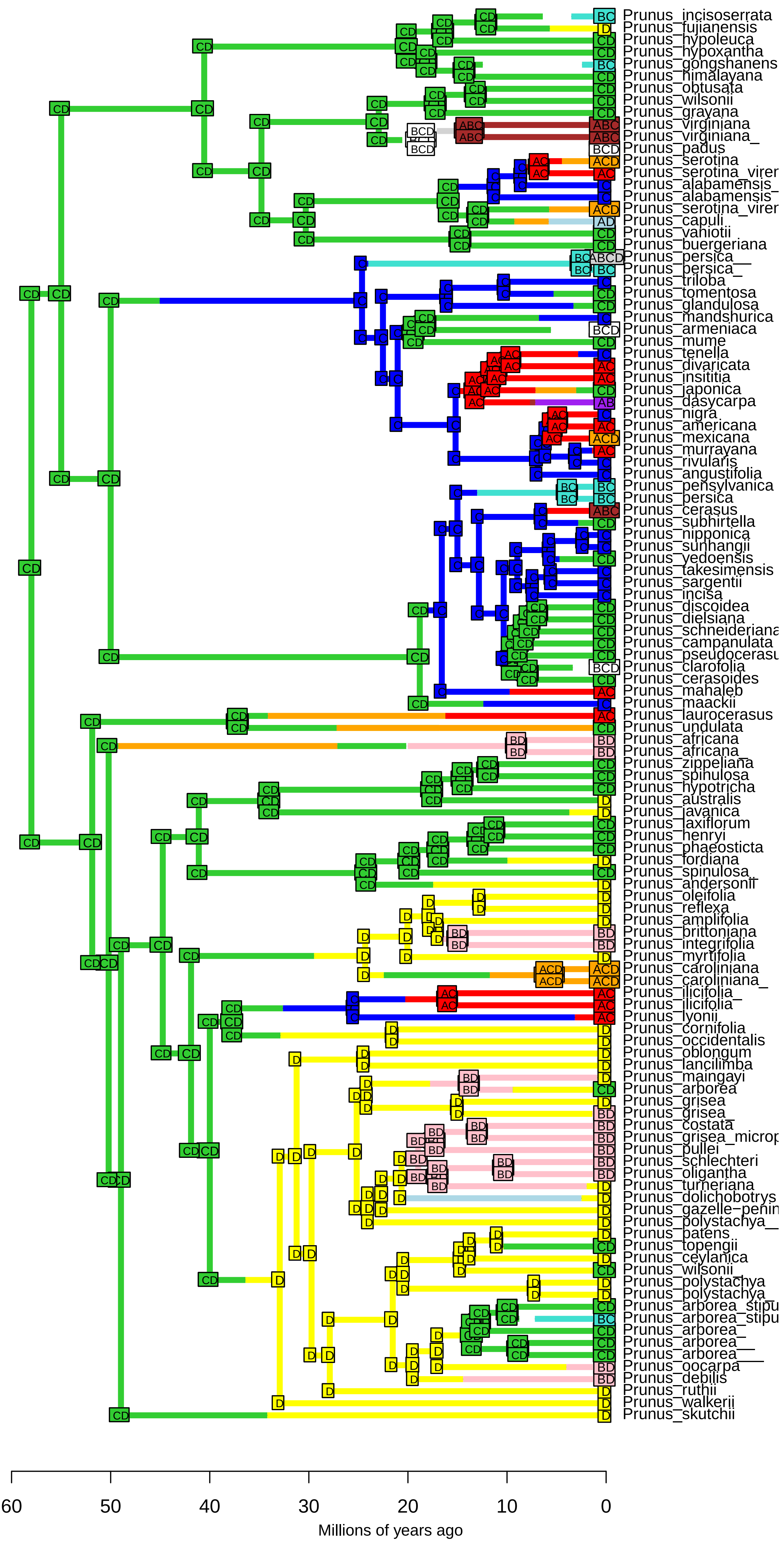

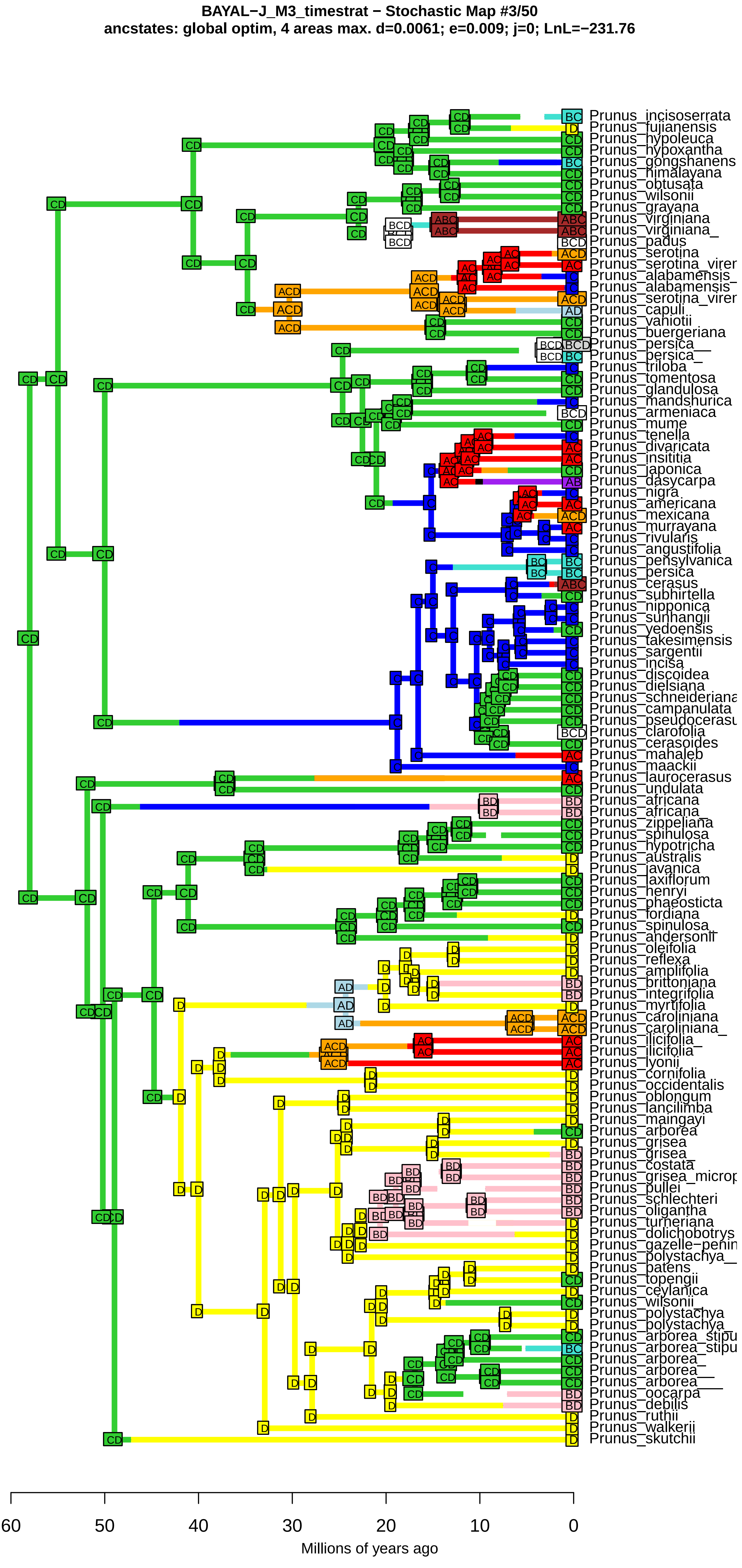

BAYAL−J_M3_timestrat − Stochastic Map #3/50
ancstates: global optim, 4 areas max. d=0.0061; e=0.009; j=0; LnL=−231.76
Prunus_incisoserrata
Prunus_fujianensis
Prunus_hypoleuca
Prunus_hypoxantha
Prunus_gongshanens
Prunus_himalayana
Prunus_obtusata
Prunus_wilsonii
Prunus_grayana
Prunus_virginiana
Prunus_virginiana_
Prunus_padus
Prunus_serotina
Prunus_serotina_virer
Prunus_alabamensis_
Prunus_alabamensis
Prunus_serotina_virer
Prunus_capuli
Prunus_vaniotii
Prunus_buergeriana
Prunus_persica__
Prunus_persica_
Prunus_triloba
Prunus_tomentosa
Prunus_glandulosa
Prunus_mandshurica
Prunus_armeniaca
Prunus_mume
Prunus_tenella
Prunus_divaricata
Prunus_insititia
Prunus_japonica
Prunus_dasycarpa
Prunus_nigra
Prunus_americana
Prunus_mexicana
Prunus_murrayana
Prunus_rivularis
Prunus_angustifolia
Prunus_pensylvanica
Prunus_persica
Prunus_cerasus
Prunus_subhirtella
Prunus_nipponica
Prunus_sunhangii
Prunus_yedoensis
Prunus_takesimensis
Prunus_sargentii
Prunus_incisa
Prunus_discoidea
Prunus_dielsiana
Prunus_schneideriana
Prunus_campanulata
Prunus_pseudocerasu
Prunus_clarofolia
Prunus_cerasoides
Prunus_mahaleb
Prunus_maackii
Prunus_laurocerasus
Prunus_undulata
Prunus_africana
Prunus_africana_
Prunus_zippeliana
Prunus_spinulosa
Prunus_hypotricha
Prunus_australis
Prunus_javanica
Prunus_laxiflorum
Prunus_henryi
Prunus_phaeosticta
Prunus_fordiana
Prunus_spinulosa_
Prunus_andersonii
Prunus_oleifolia
Prunus_reflexa
Prunus_amplifolia
Prunus_brittoniana
Prunus_integrifolia
Prunus_myrtifolia
Prunus_caroliniana
Prunus_caroliniana_
Prunus_ilicifolia_
Prunus_ilicifolia
Prunus_lyonii
Prunus_cornifolia
Prunus_occidentalis
Prunus_oblongum
Prunus_lancilimba
Prunus_maingayi
Prunus_arborea
Prunus_grisea
Prunus_grisea
Prunus_costata
Prunus_grisea_microp
Prunus_pullei
Prunus_schlechteri
Prunus_oligantha
Prunus_turneriana
Prunus_dolichobotrys
Prunus_gazelle−penin
Prunus_polystachya__
Prunus_patens
Prunus_topengii
Prunus_ceylanica
Prunus_wilsonii_
Prunus_polystachya
Prunus_polystachya_
Prunus_arborea_stipu
Prunus_arborea_stipu
Prunus_arborea_
Prunus_arborea__
Prunus_arborea___
Prunus_oocarpa
Prunus_debilis
Prunus_ruthii
Prunus_walkerii
Prunus_skutchii
60
50
40
30
20
10
0
Millions of years ago

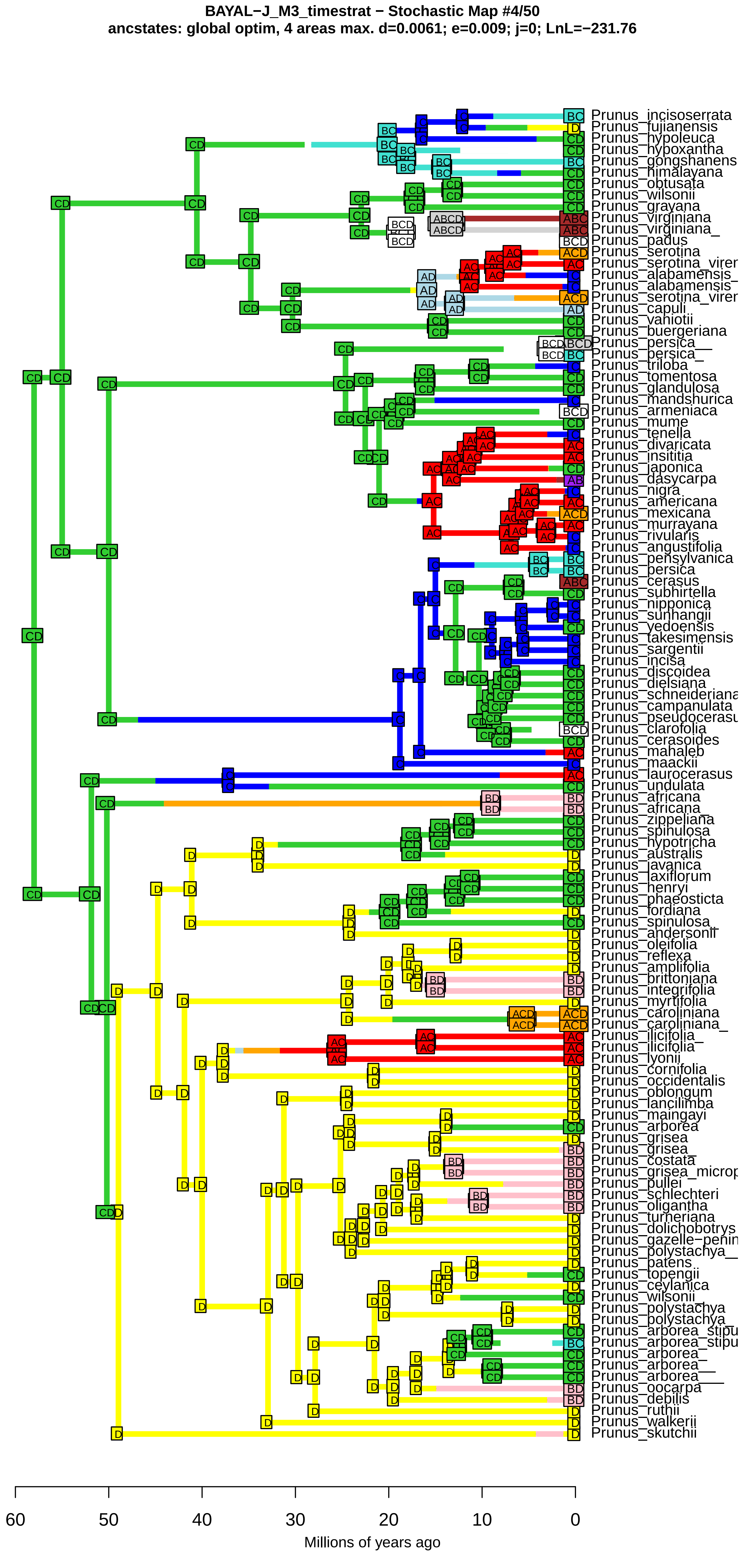
BAYAL−J_M3_timestrat − Stochastic Map #4/50
ancstates: global optim, 4 areas max. d=0.0061; e=0.009; j=0; LnL=−231.76
Prunus_incisoserrata
Prunus_fujianensis
Prunus_hypoleuca
Prunus_hypoxantha
Prunus_gongshanens
Prunus_himalayana
Prunus_obtusata
Prunus_wilsonii
Prunus_grayana
Prunus_virginiana
Prunus_virginiana_
Prunus_padus
Prunus_serotina
Prunus_serotina_virer
Prunus_alabamensis_
Prunus_alabamensis
Prunus_serotina_virer
Prunus_capuli
Prunus_vaniotii
Prunus_buergeriana
Prunus_persica__
Prunus_persica_
Prunus_triloba
Prunus_tomentosa
Prunus_glandulosa
Prunus_mandshurica
Prunus_armeniaca
Prunus_mume
Prunus_tenella
Prunus_divaricata
Prunus_insititia
Prunus_japonica
Prunus_dasycarpa
Prunus_nigra
Prunus_americana
Prunus_mexicana
Prunus_murrayana
Prunus_rivularis
Prunus_angustifolia
Prunus_pensylvanica
Prunus_persica
Prunus_cerasus
Prunus_subhirtella
Prunus_nipponica
Prunus_sunhangii
Prunus_yedoensis
Prunus_takesimensis
Prunus_sargentii
Prunus_incisa
Prunus_discoidea
Prunus_dielsiana
Prunus_schneideriana
Prunus_campanulata
Prunus_pseudocerasu
Prunus_clarofolia
Prunus_cerasoides
Prunus_mahaleb
Prunus_maackii
Prunus_laurocerasus
Prunus_undulata
Prunus_africana
Prunus_africana_
Prunus_zippeliana
Prunus_spinulosa
Prunus_hypotricha
Prunus_australis
Prunus_javanica
Prunus_laxiflorum
Prunus_henryi
Prunus_phaeosticta
Prunus_fordiana
Prunus_spinulosa_
Prunus_andersonii
Prunus_oleifolia
Prunus_reflexa
Prunus_amplifolia
Prunus_brittoniana
Prunus_integrifolia
Prunus_myrtifolia
Prunus_caroliniana
Prunus_caroliniana_
Prunus_ilicifolia_
Prunus_ilicifolia
Prunus_lyonii
Prunus_cornifolia
Prunus_occidentalis
Prunus_oblongum
Prunus_lancilimba
Prunus_maingayi
Prunus_arborea
Prunus_grisea
Prunus_grisea_
Prunus_costata
Prunus_grisea_microp
Prunus_pullei
Prunus_schlechteri
Prunus_oligantha
Prunus_turneriana
Prunus_dolichobotrys
Prunus_gazelle−penin
Prunus_polystachya__
Prunus_patens
Prunus_topengii
Prunus_ceylanica
Prunus_wilsonii_
Prunus_polystachya
Prunus_polystachya_
Prunus_arborea_stipu
Prunus_arborea_stipu
Prunus_arborea_
Prunus_arborea__
Prunus_arborea___
Prunus_oocarpa
Prunus_debilis
Prunus_ruthii
Prunus_walkerii
Prunus_skutchii
60
50
40
30
20
10
0
Millions of years ago

BAYAL−J_M3_timestrat − Stochastic Map #5/50
ancstates: global optim, 4 areas max. d=0.0061; e=0.009; j=0; LnL=−231.76

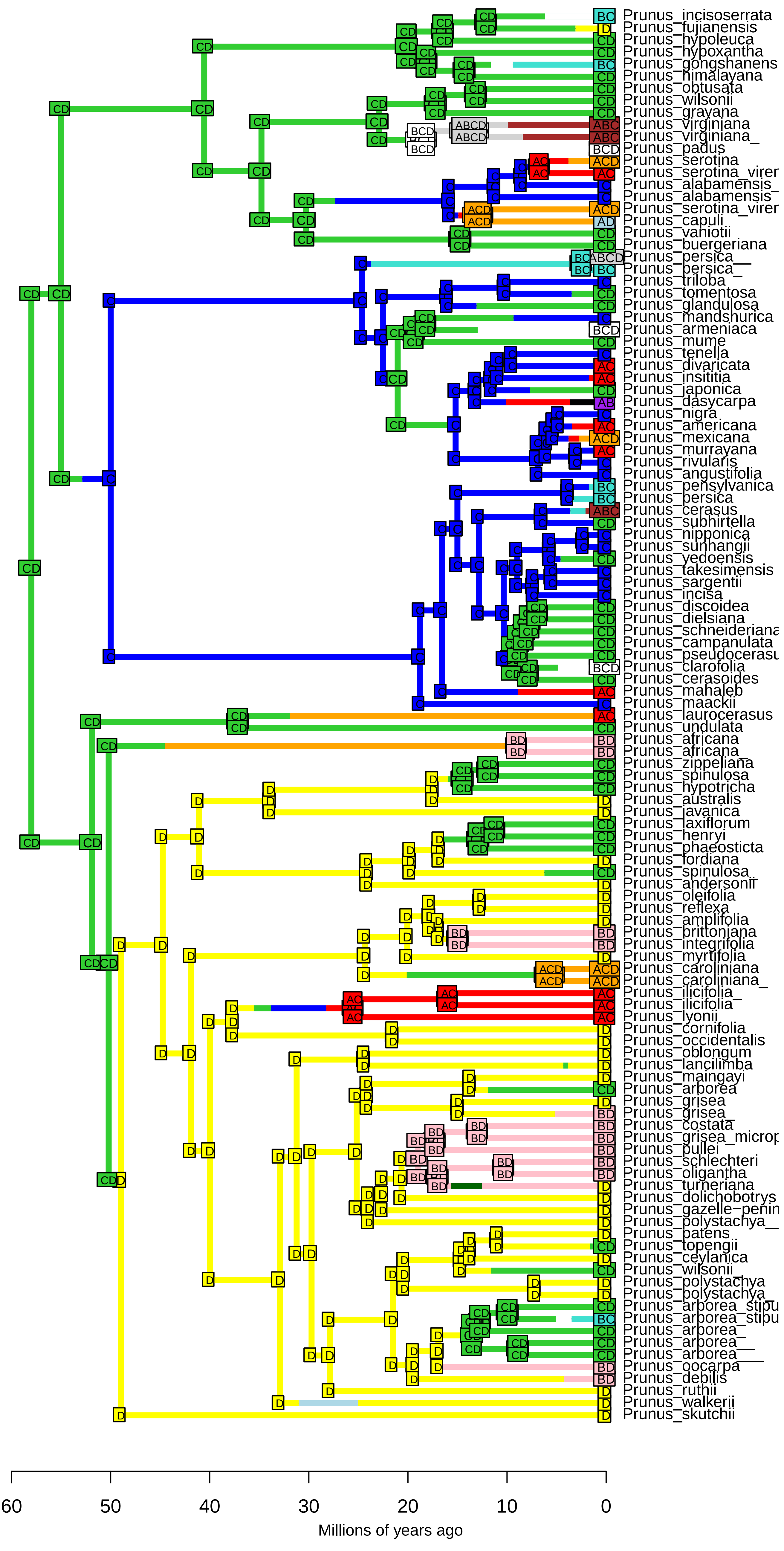

**BAYAL−J_M3_timestrat − Stochastic Map #6/50**
**ancstates: global optim, 4 areas max. d=0.0061; e=0.009; j=0; LnL=−231.76**

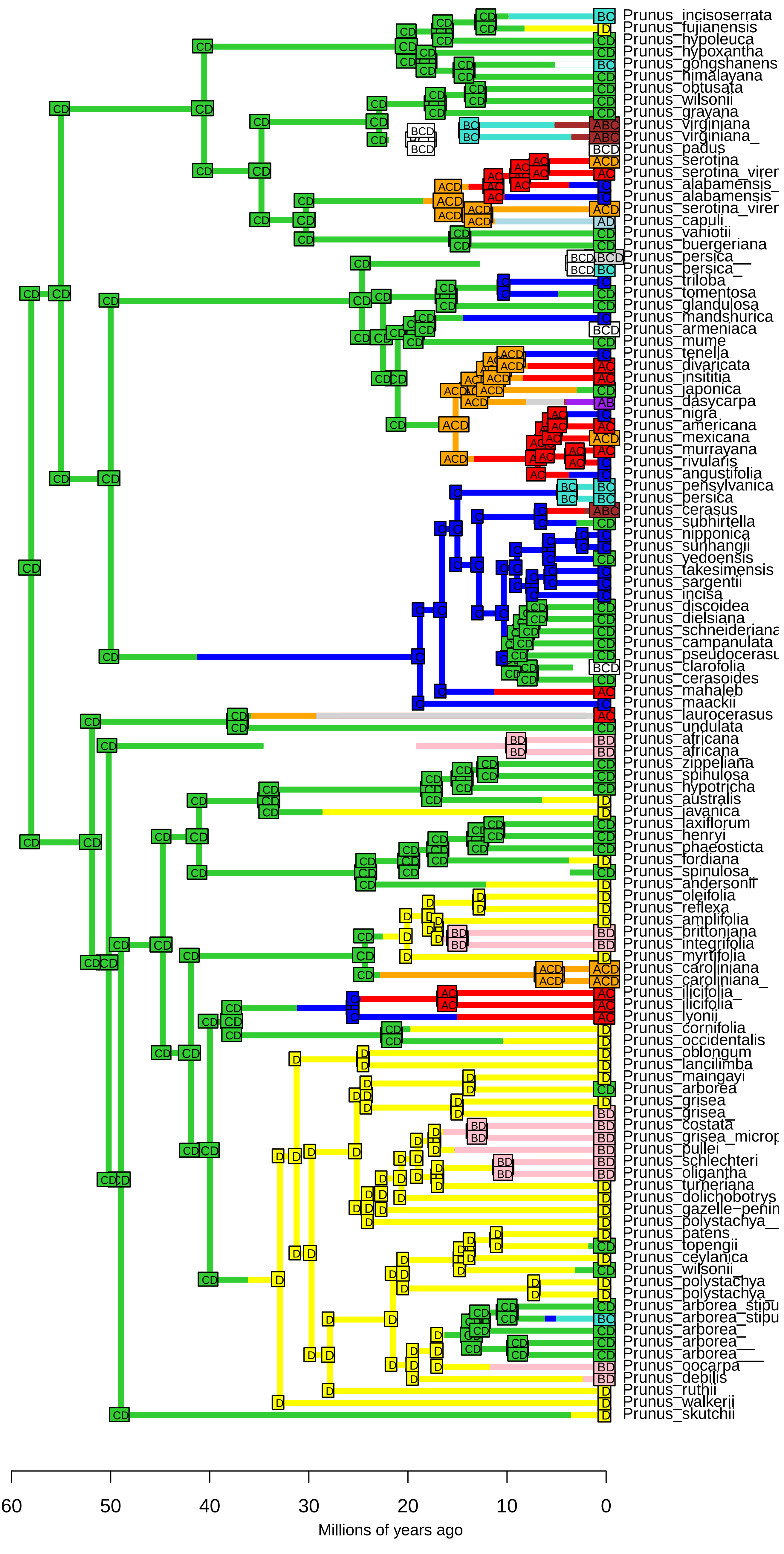

**BAYAL−J_M3_timestrat − Stochastic Map #7/50**
**ancstates: global optim, 4 areas max. d=0.0061; e=0.009; j=0; LnL=−231.76**

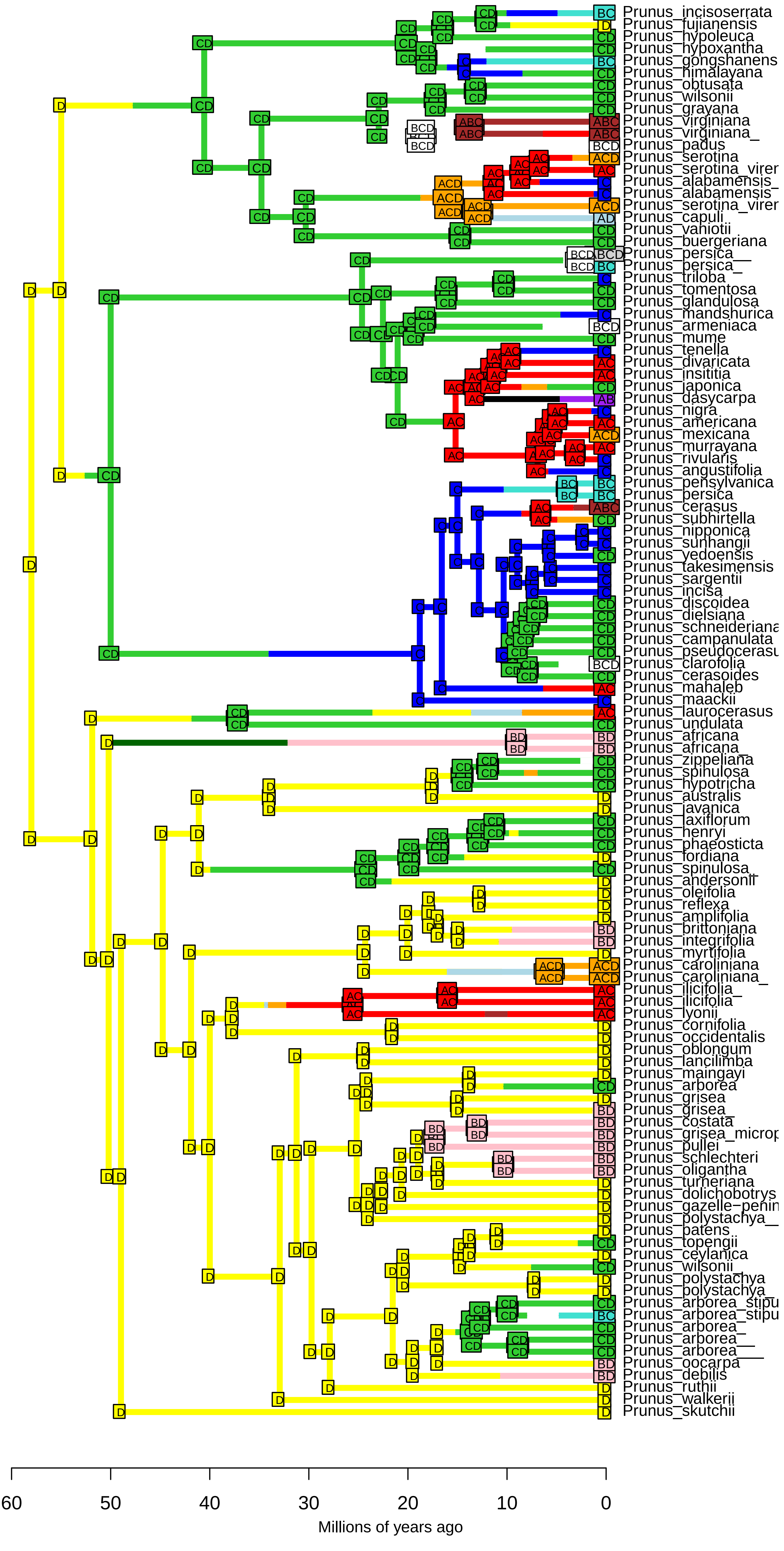

**BAYAL−J_M3_timestrat − Stochastic Map #8/50**
**ancstates: global optim, 4 areas max. d=0.0061; e=0.009; j=0; LnL=−231.76**

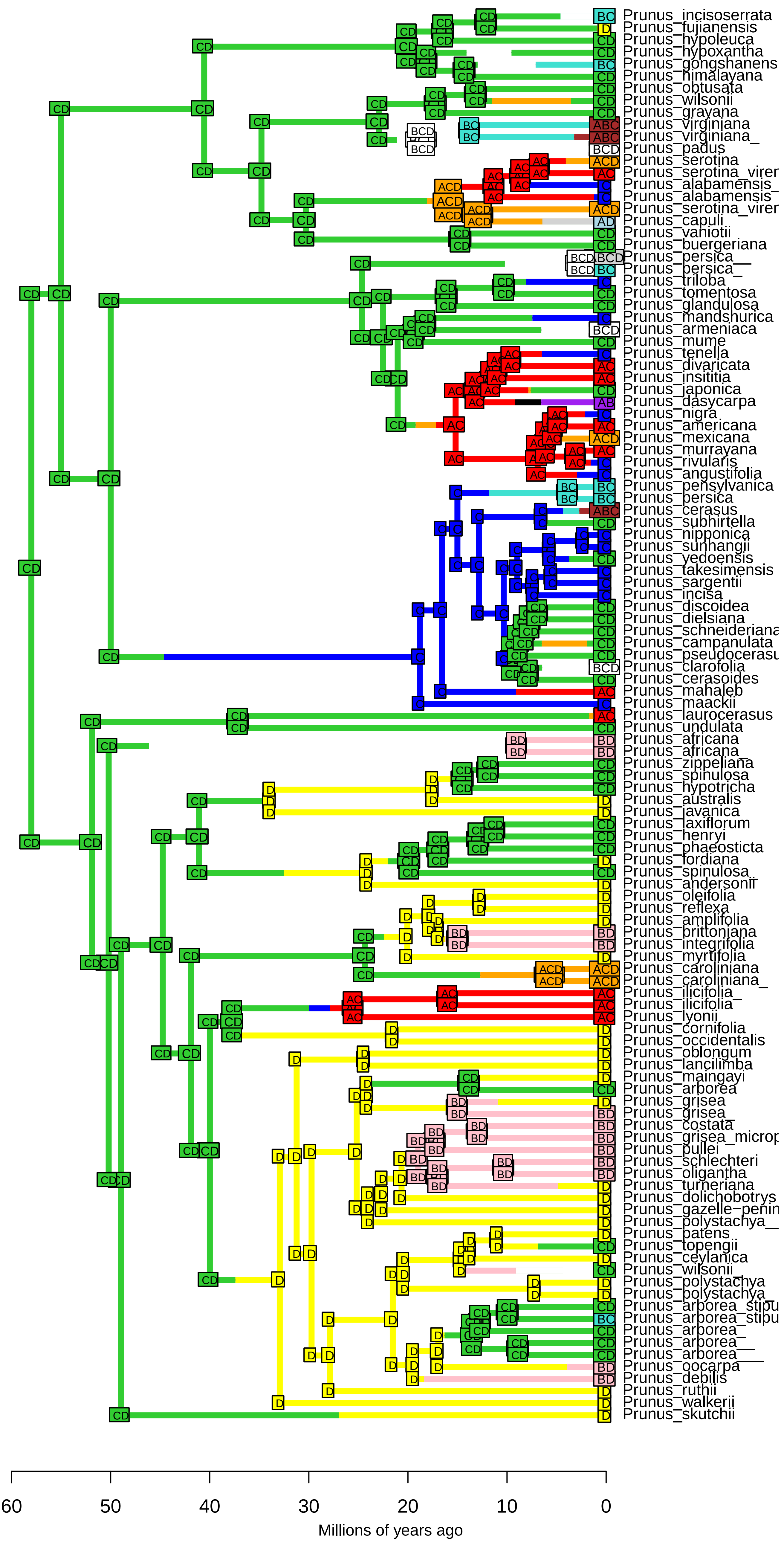

BAYAL−J_M3_timestrat − Stochastic Map #9/50
ancstates: global optim, 4 areas max. d=0.0061; e=0.009; j=0; LnL=−231.76

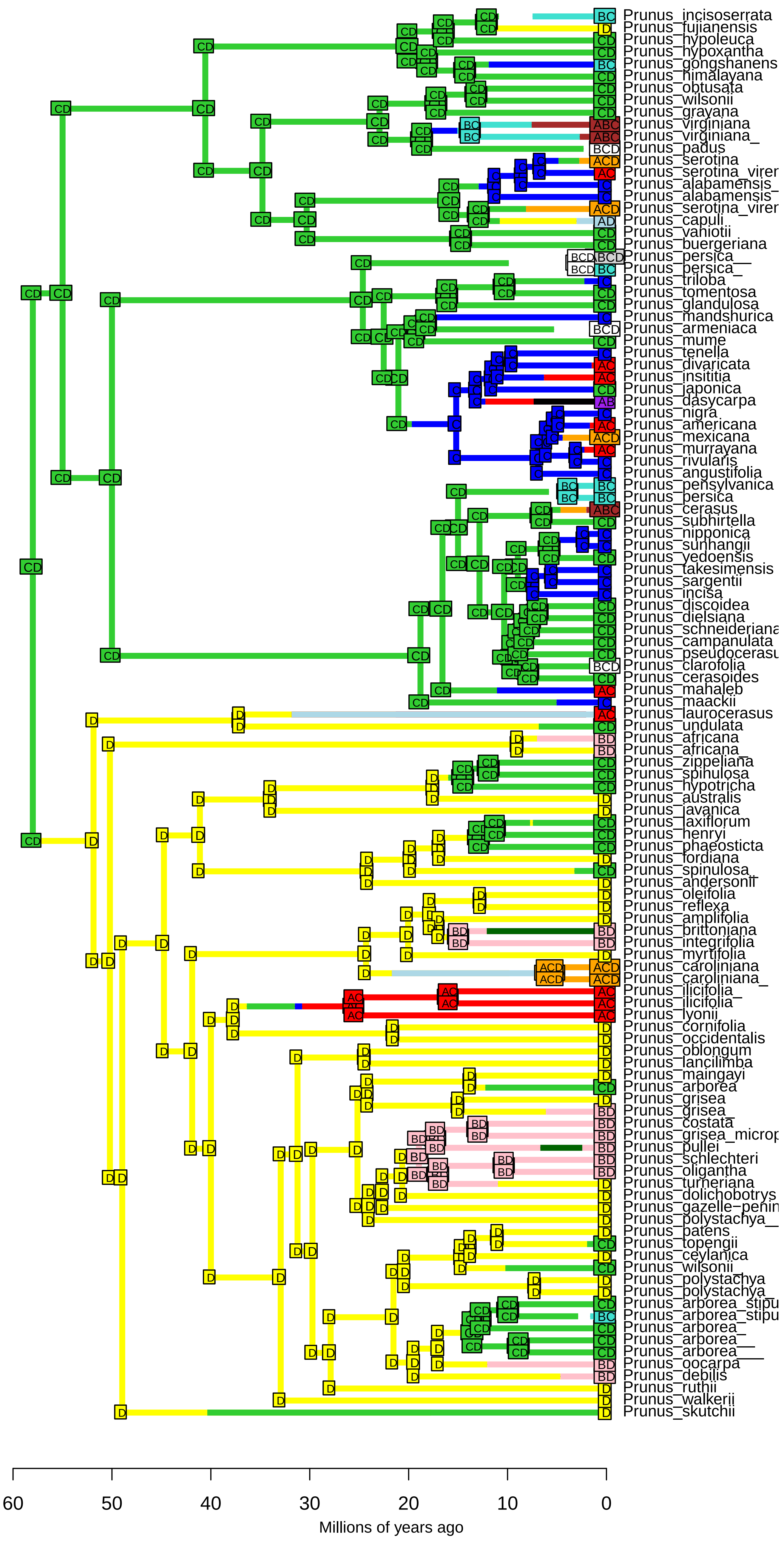

**BAYAL−J_M3_timestrat − Stochastic Map #10/50**
**ancstates: global optim, 4 areas max. d=0.0061; e=0.009; j=0; LnL=−231.76**

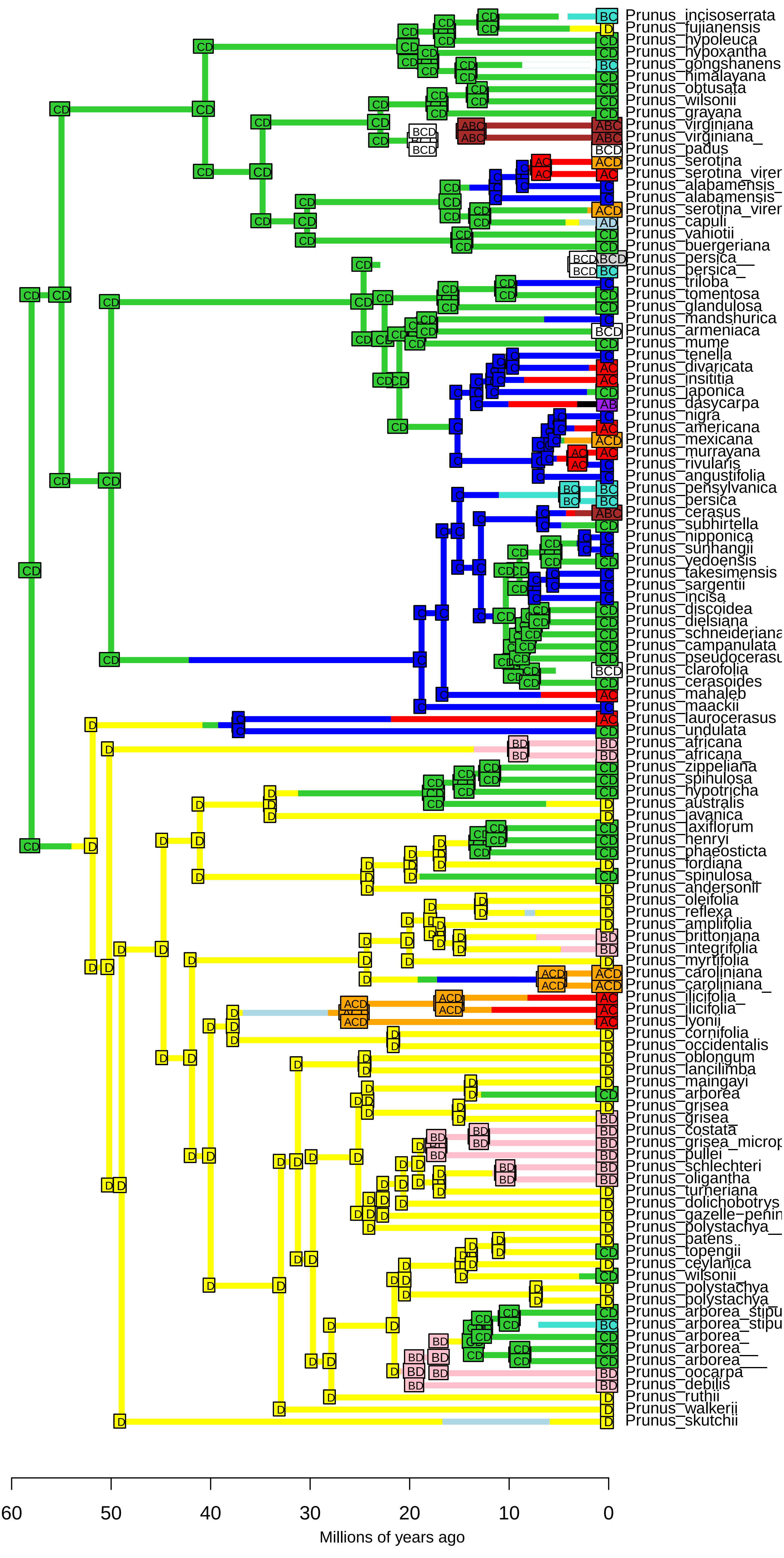

**BAYAL−J_M3_timestrat − Stochastic Map #11/50**
**ancstates: global optim, 4 areas max. d=0.0061; e=0.009; j=0; LnL=−231.76**

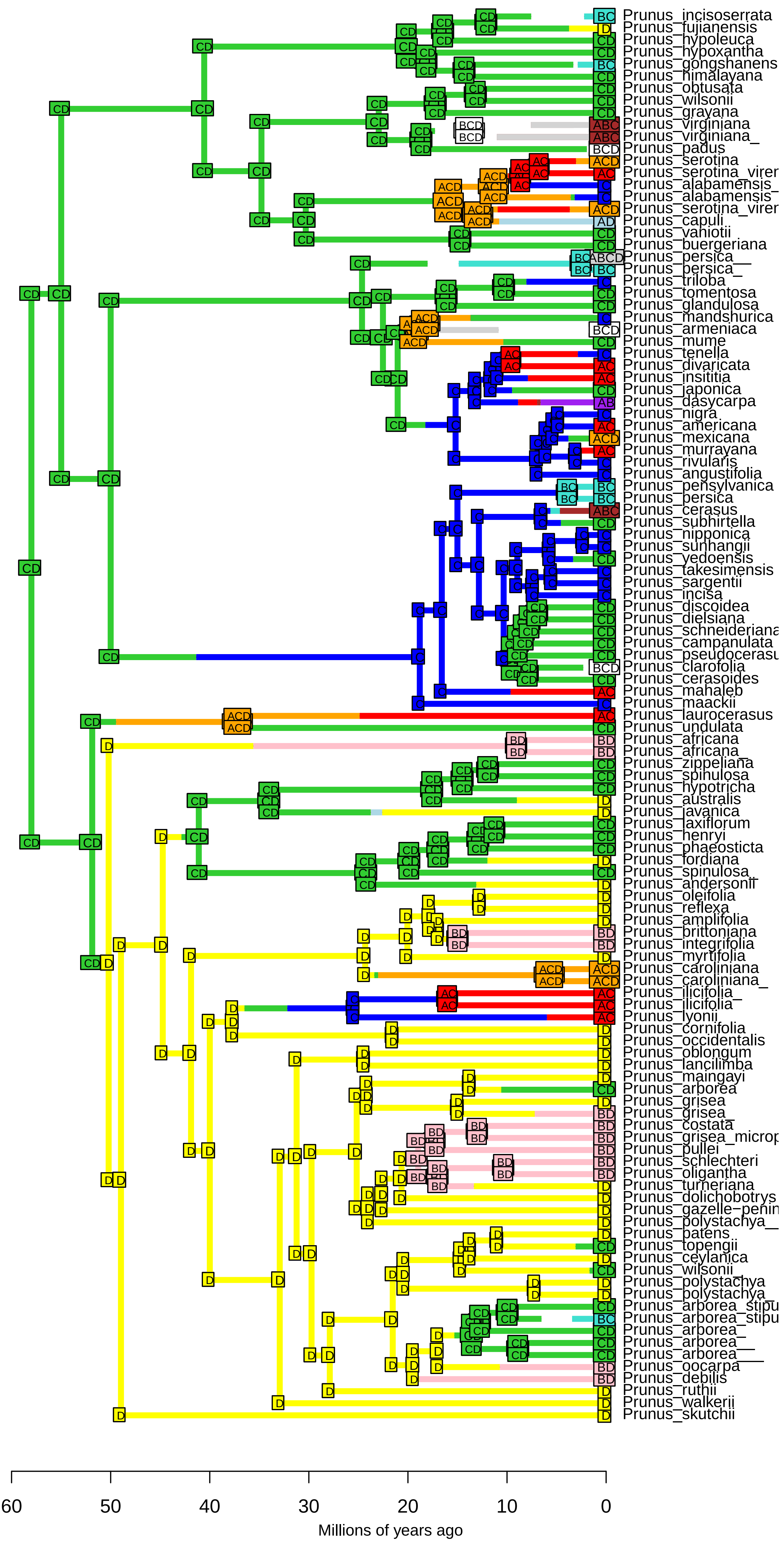

**BAYAL−J_M3_timestrat − Stochastic Map #12/50**
**ancstates: global optim, 4 areas max. d=0.0061; e=0.009; j=0; LnL=−231.76**

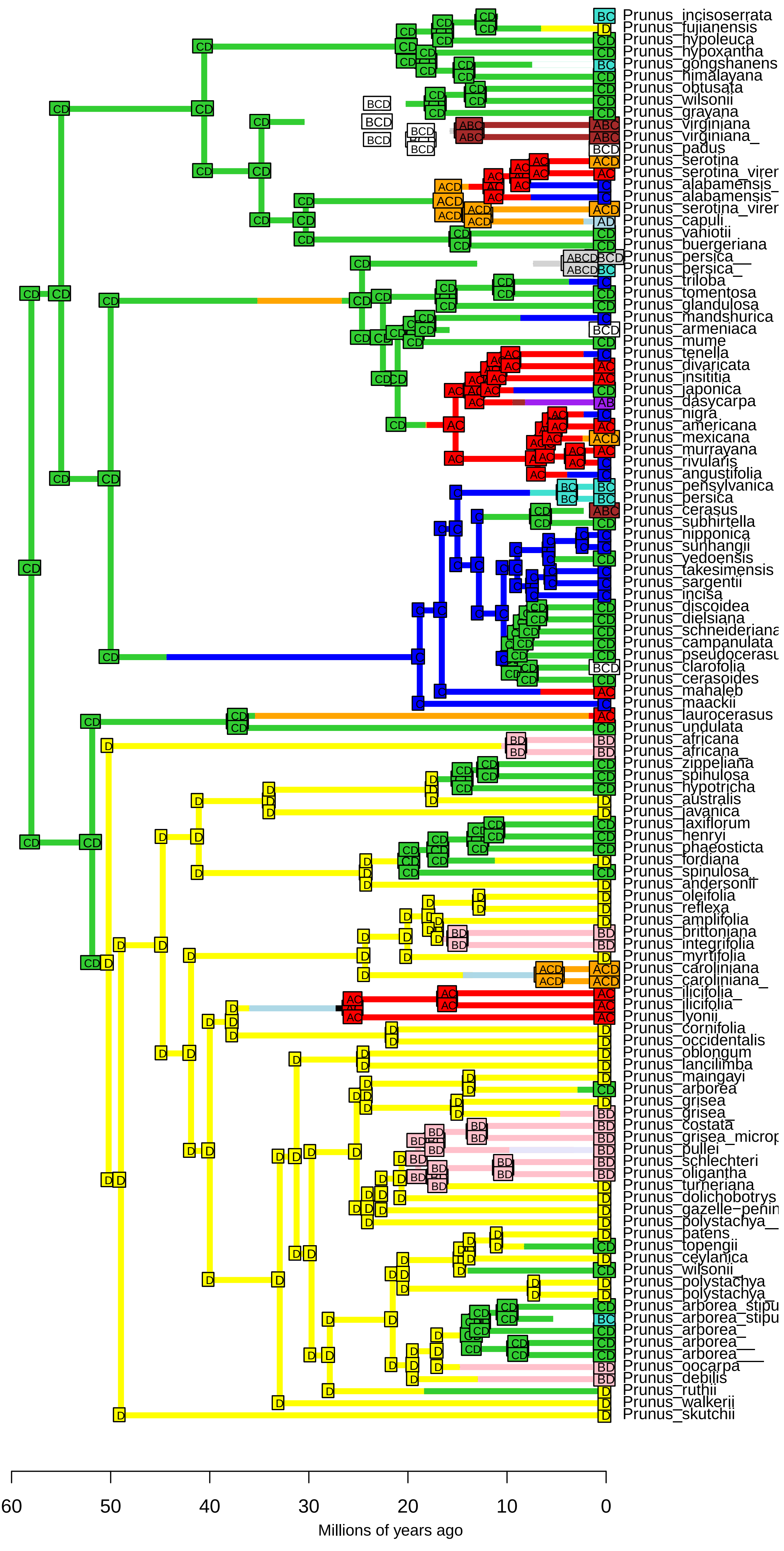

**BAYAL−J_M3_timestrat − Stochastic Map #13/50**
**ancstates: global optim, 4 areas max. d=0.0061; e=0.009; j=0; LnL=−231.76**

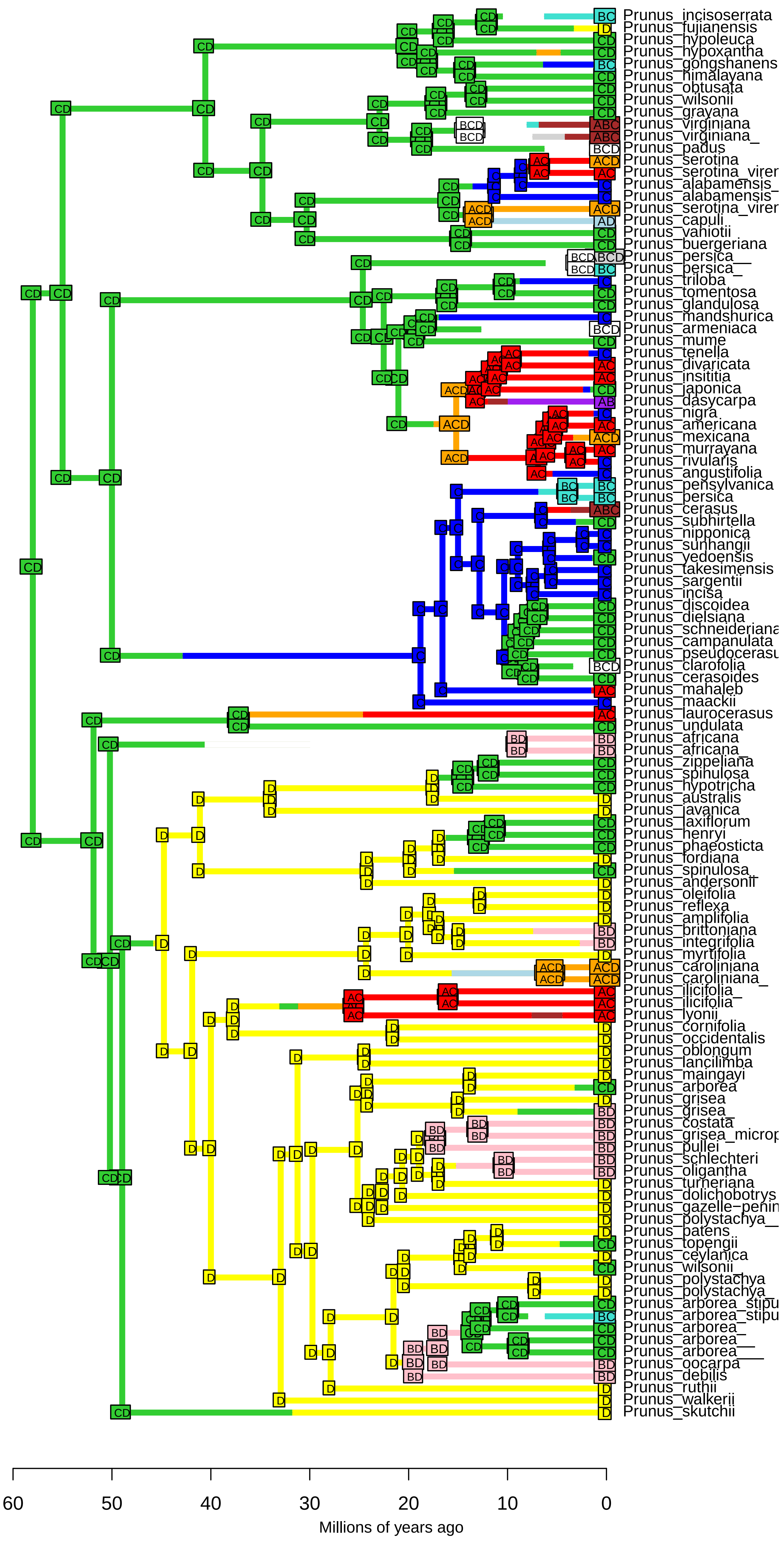

**BAYAL−J_M3_timestrat − Stochastic Map #14/50**
**ancstates: global optim, 4 areas max. d=0.0061; e=0.009; j=0; LnL=−231.76**

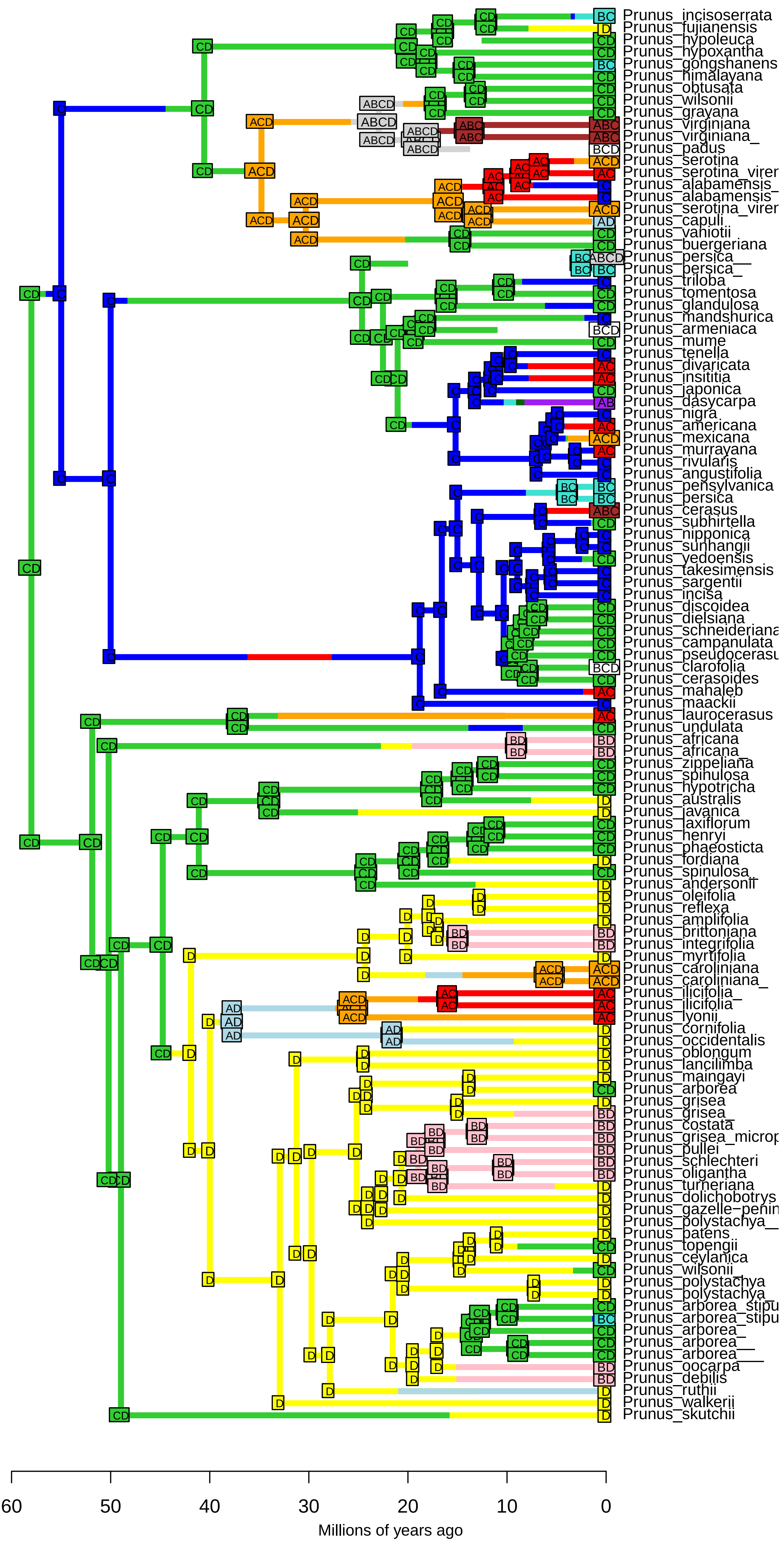

**BAYAL−J_M3_timestrat − Stochastic Map #15/50**
**ancstates: global optim, 4 areas max. d=0.0061; e=0.009; j=0; LnL=−231.76**

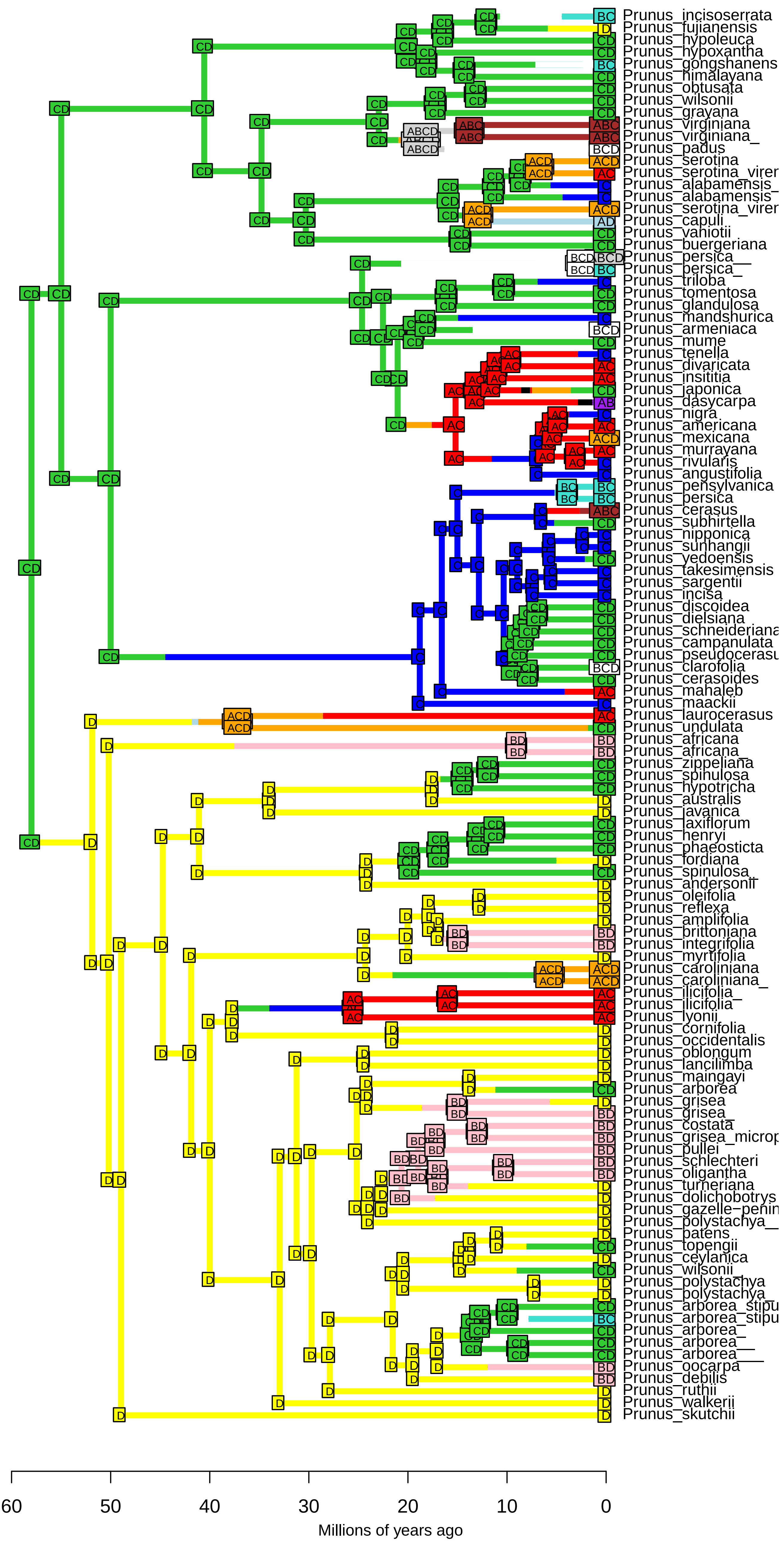

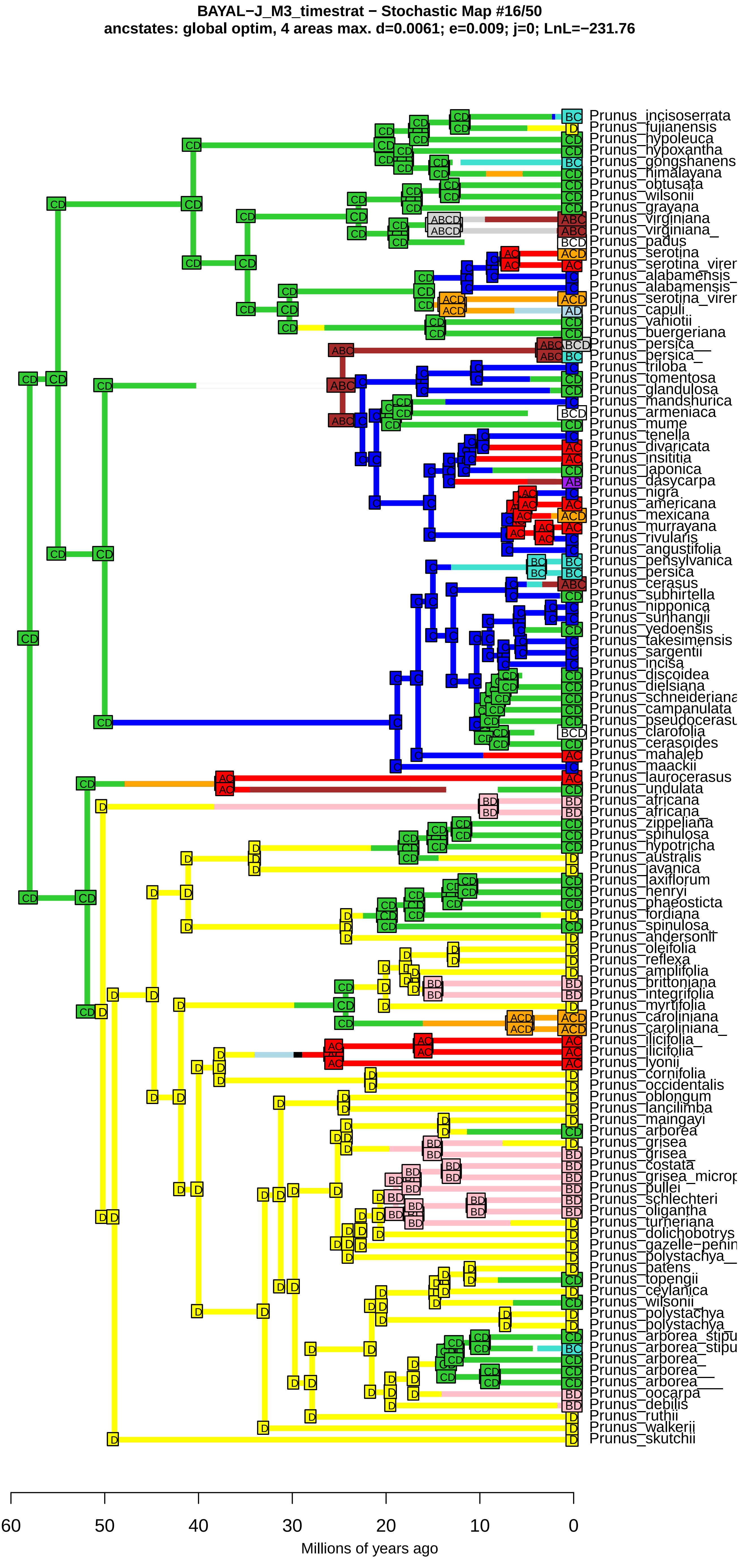
BAYAL−J_M3_timestrat − Stochastic Map #16/50
ancstates: global optim, 4 areas max. d=0.0061; e=0.009; j=0; LnL=−231.76
Prunus_incisoserrata
Prunus_fujianensis
Prunus_hypoleuca
Prunus_hypoxantha
Prunus_gongshanens
Prunus_himalayana
Prunus_obtusata
Prunus_wilsonii
Prunus_grayana
Prunus_virginiana
Prunus_virginiana_
Prunus_padus
Prunus_serotina
Prunus_serotina_virer
Prunus_alabamensis_
Prunus_alabamensis
Prunus_serotina_virer
Prunus_capuli
Prunus_vaniotii
Prunus_buergeriana
Prunus_persica__
Prunus_persica_
Prunus_triloba
Prunus_tomentosa
Prunus_glandulosa
Prunus_mandshurica
Prunus_armeniaca
Prunus_mume
Prunus_tenella
Prunus_divaricata
Prunus_insititia
Prunus_japonica
Prunus_dasycarpa
Prunus_nigra
Prunus_americana
Prunus_mexicana
Prunus_murrayana
Prunus_rivularis
Prunus_angustifolia
Prunus_pensylvanica
Prunus_persica
Prunus_cerasus
Prunus_subhirtella
Prunus_nipponica
Prunus_sunhangii
Prunus_yedoensis
Prunus_takesimensis
Prunus_sargentii
Prunus_incisa
Prunus_discoidea
Prunus_dielsiana
Prunus_schneideriana
Prunus_campanulata
Prunus_pseudocerasu
Prunus_clarofolia
Prunus_cerasoides
Prunus_mahaleb
Prunus_maackii
Prunus_laurocerasus
Prunus_undulata
Prunus_africana
Prunus_africana_
Prunus_zippeliana
Prunus_spinulosa
Prunus_hypotricha
Prunus_australis
Prunus_javanica
Prunus_laxiflorum
Prunus_henryi
Prunus_phaeosticta
Prunus_fordiana
Prunus_spinulosa_
Prunus_andersonii
Prunus_oleifolia
Prunus_reflexa
Prunus_amplifolia
Prunus_brittoniana
Prunus_integrifolia
Prunus_myrtifolia
Prunus_caroliniana
Prunus_caroliniana_
Prunus_ilicifolia_
Prunus_ilicifolia
Prunus_lyonii
Prunus_cornifolia
Prunus_occidentalis
Prunus_oblongum
Prunus_lancilimba
Prunus_maingayi
Prunus_arborea
Prunus_grisea
Prunus_grisea_
Prunus_costata
Prunus_grisea_microp
Prunus_pullei
Prunus_schlechteri
Prunus_oligantha
Prunus_turneriana
Prunus_dolichobotrys
Prunus_gazelle−penin
Prunus_polystachya__
Prunus_patens
Prunus_topengii
Prunus_ceylanica
Prunus_wilsonii_
Prunus_polystachya
Prunus_polystachya_
Prunus_arborea_stipu
Prunus_arborea_stipu
Prunus_arborea_
Prunus_arborea__
Prunus_arborea___
Prunus_oocarpa
Prunus_debilis
Prunus_ruthii
Prunus_walkerii
Prunus_skutchii
60
50
40
30
20
10
0
Millions of years ago

**BAYAL−J_M3_timestrat − Stochastic Map #17/50**
**ancstates: global optim, 4 areas max. d=0.0061; e=0.009; j=0; LnL=−231.76**

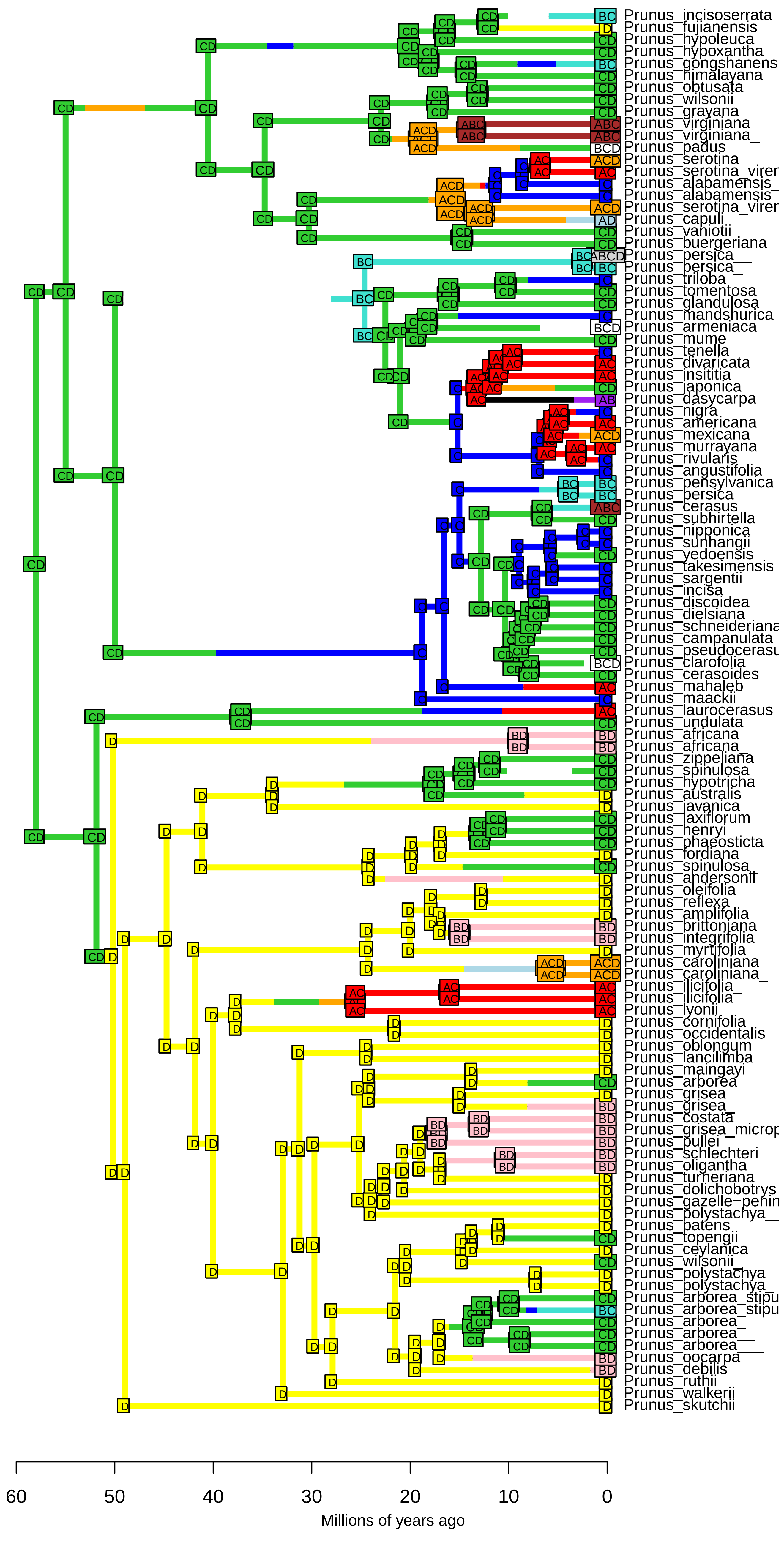

BAYAL−J_M3_timestrat − Stochastic Map #18/50
ancstates: global optim, 4 areas max. d=0.0061; e=0.009; j=0; LnL=−231.76

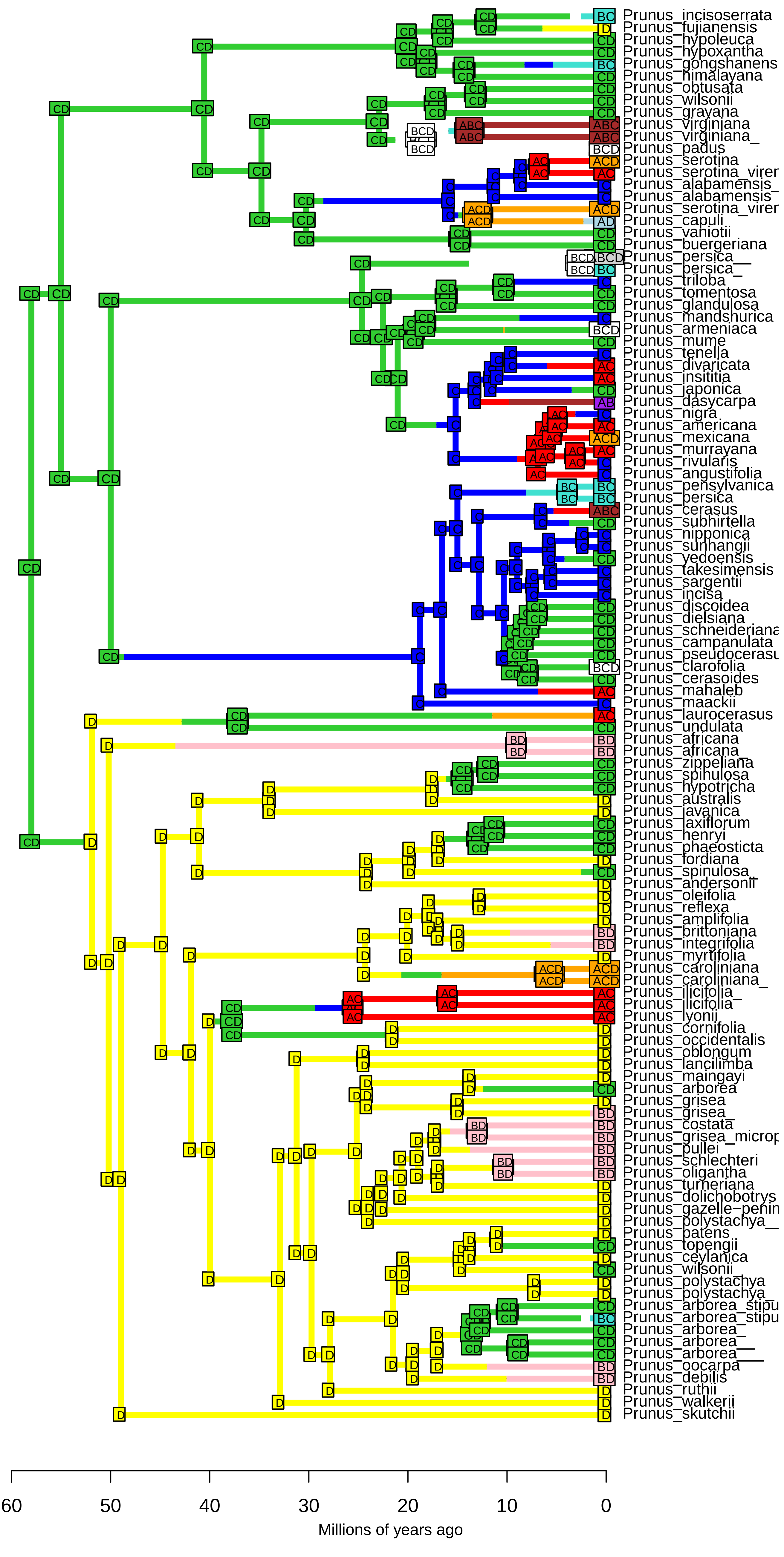

**BAYAL−J_M3_timestrat − Stochastic Map #19/50**
**ancstates: global optim, 4 areas max. d=0.0061; e=0.009; j=0; LnL=−231.76**

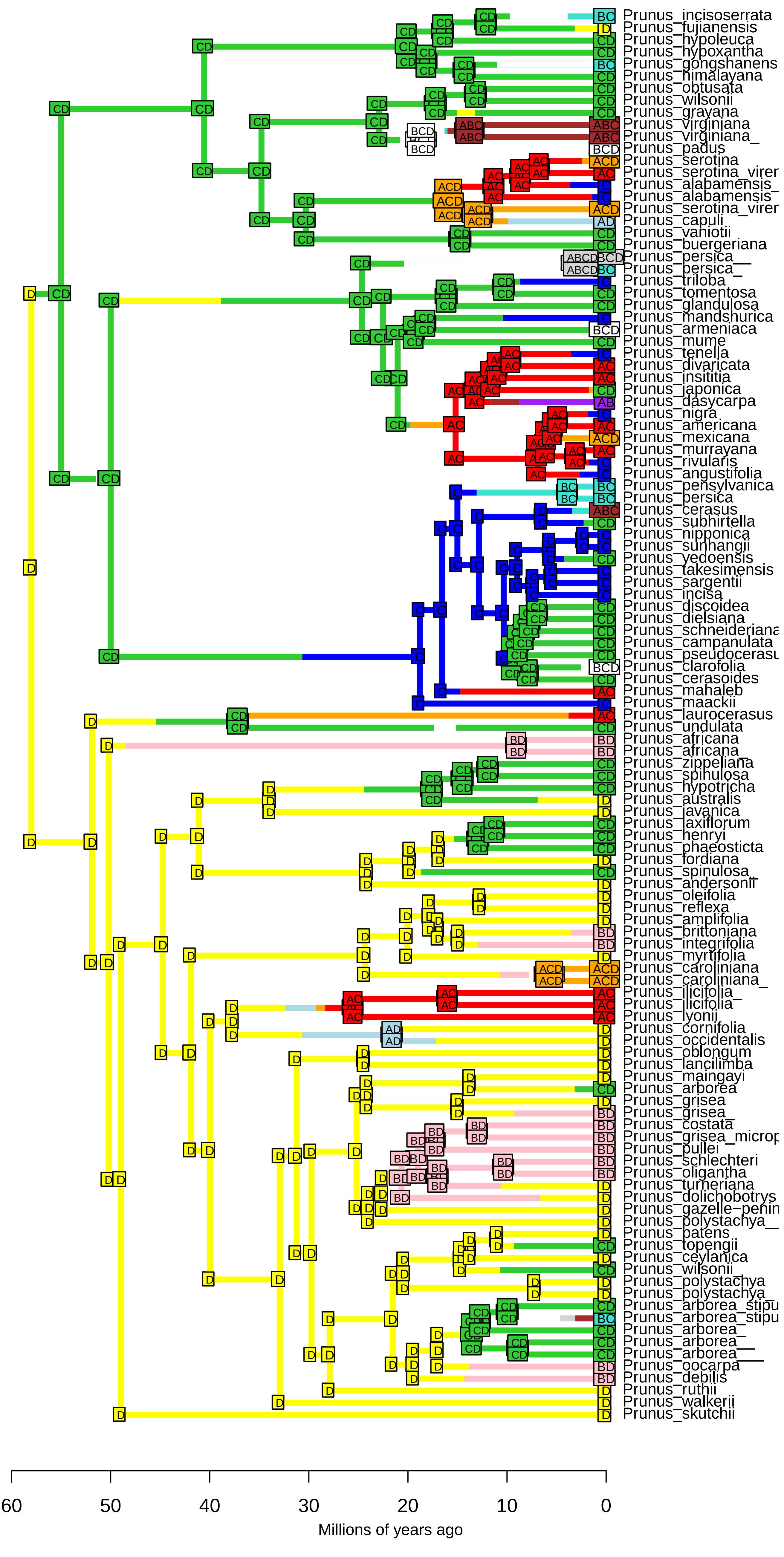

**BAYAL−J_M3_timestrat − Stochastic Map #20/50**
**ancstates: global optim, 4 areas max. d=0.0061; e=0.009; j=0; LnL=−231.76**

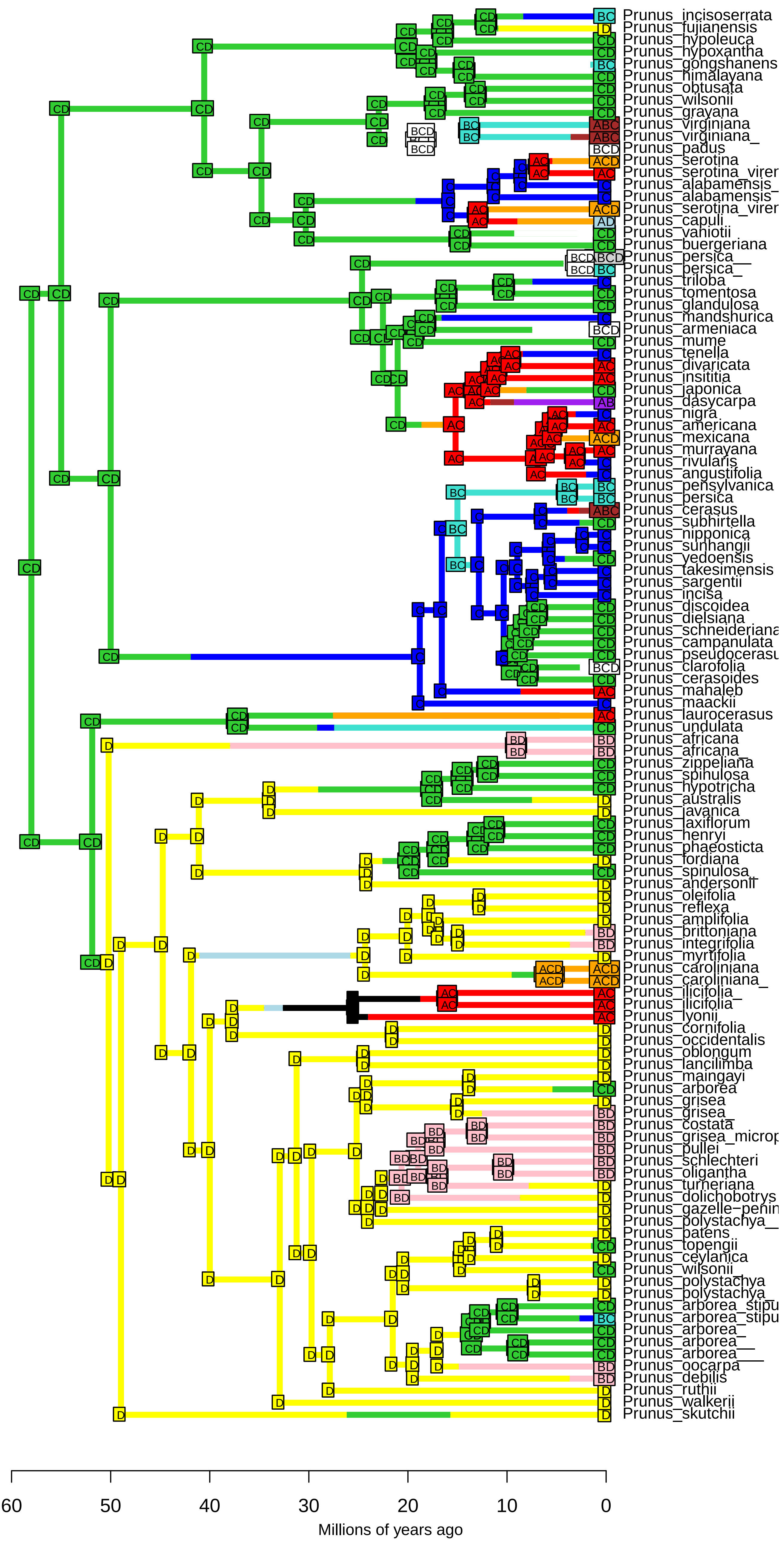

**BAYAL−J_M3_timestrat − Stochastic Map #21/50**
**ancstates: global optim, 4 areas max. d=0.0061; e=0.009; j=0; LnL=−231.76**

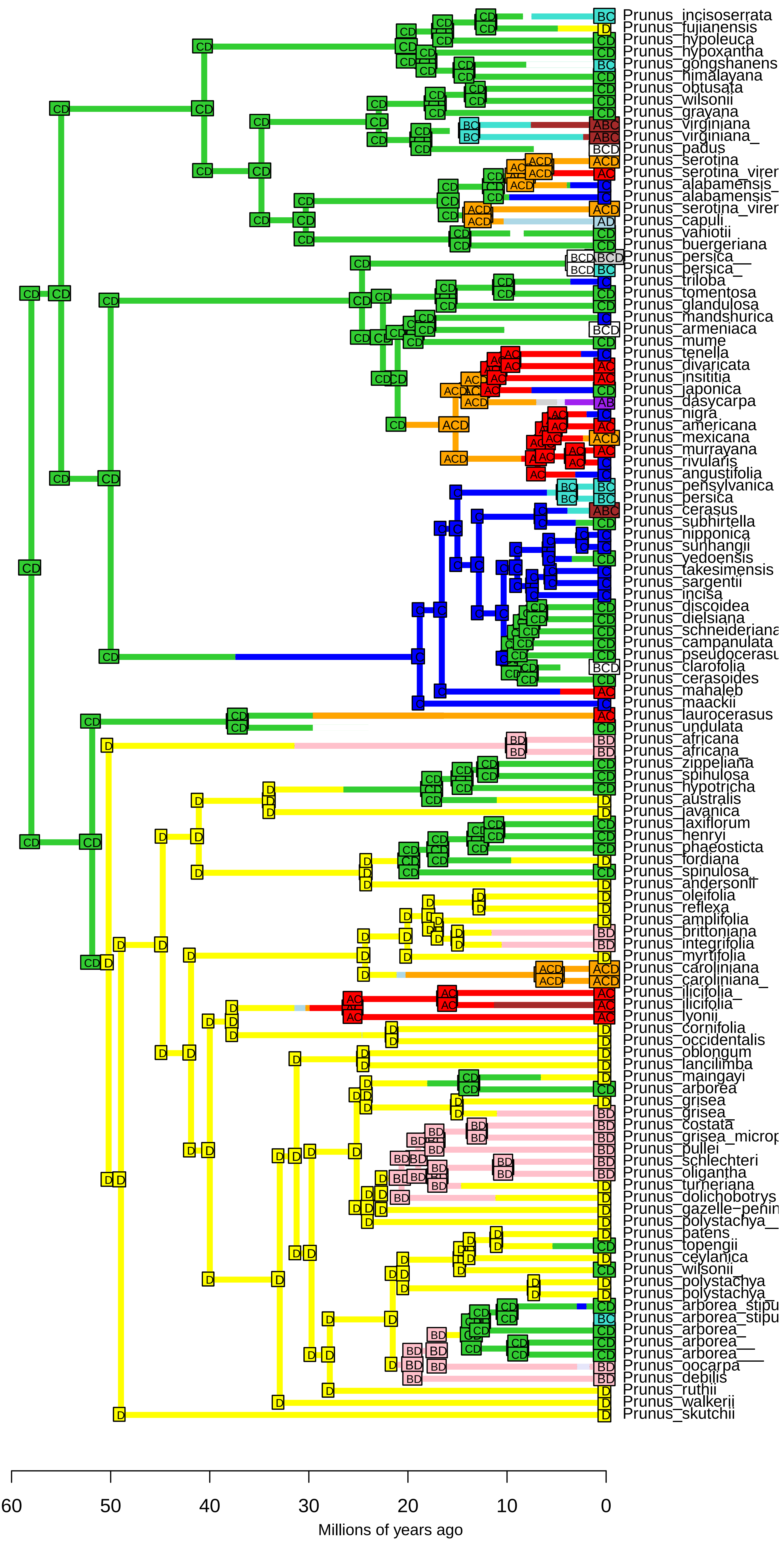

**BAYAL−J_M3_timestrat − Stochastic Map #22/50**
**ancstates: global optim, 4 areas max. d=0.0061; e=0.009; j=0; LnL=−231.76**

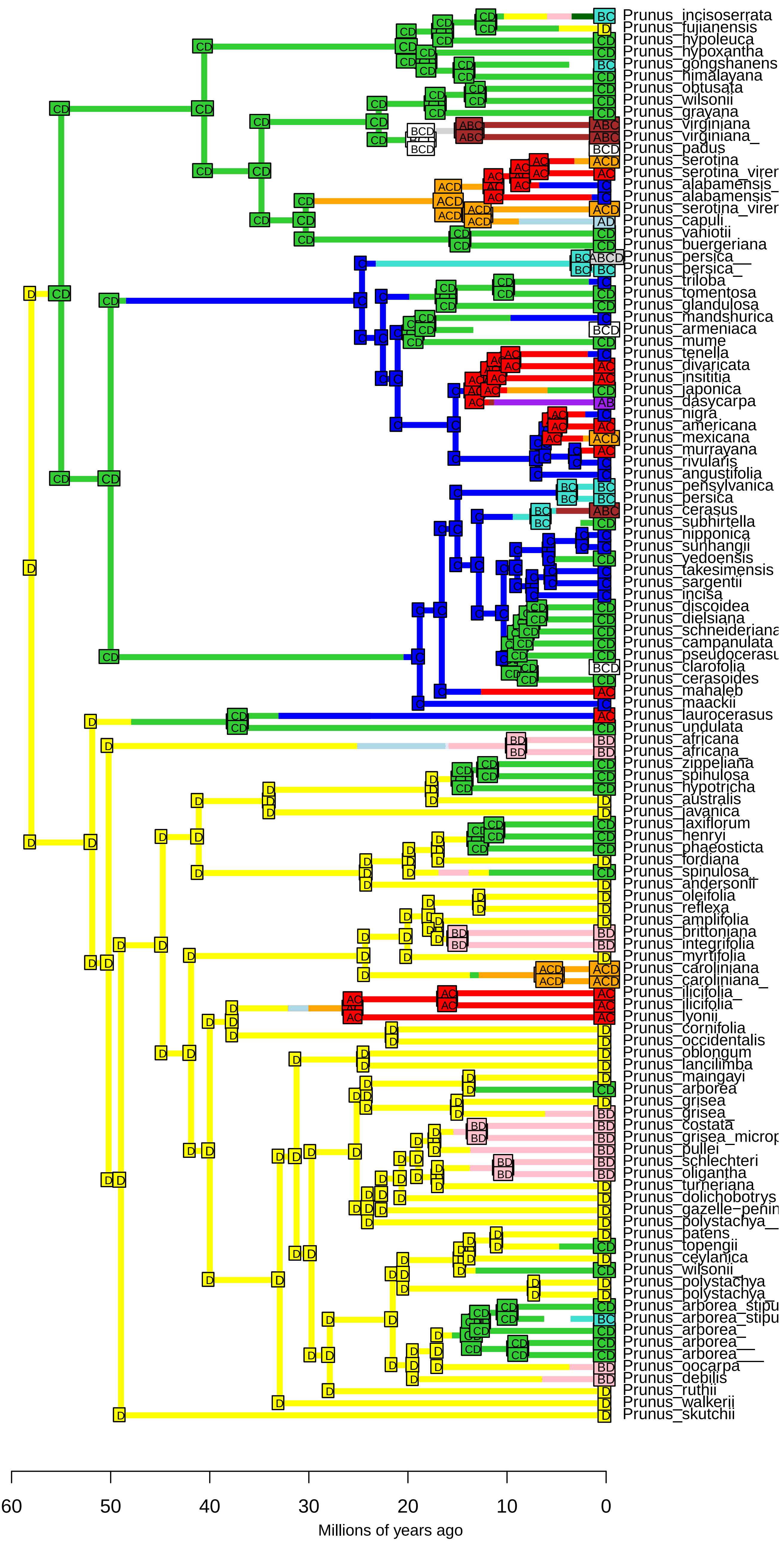

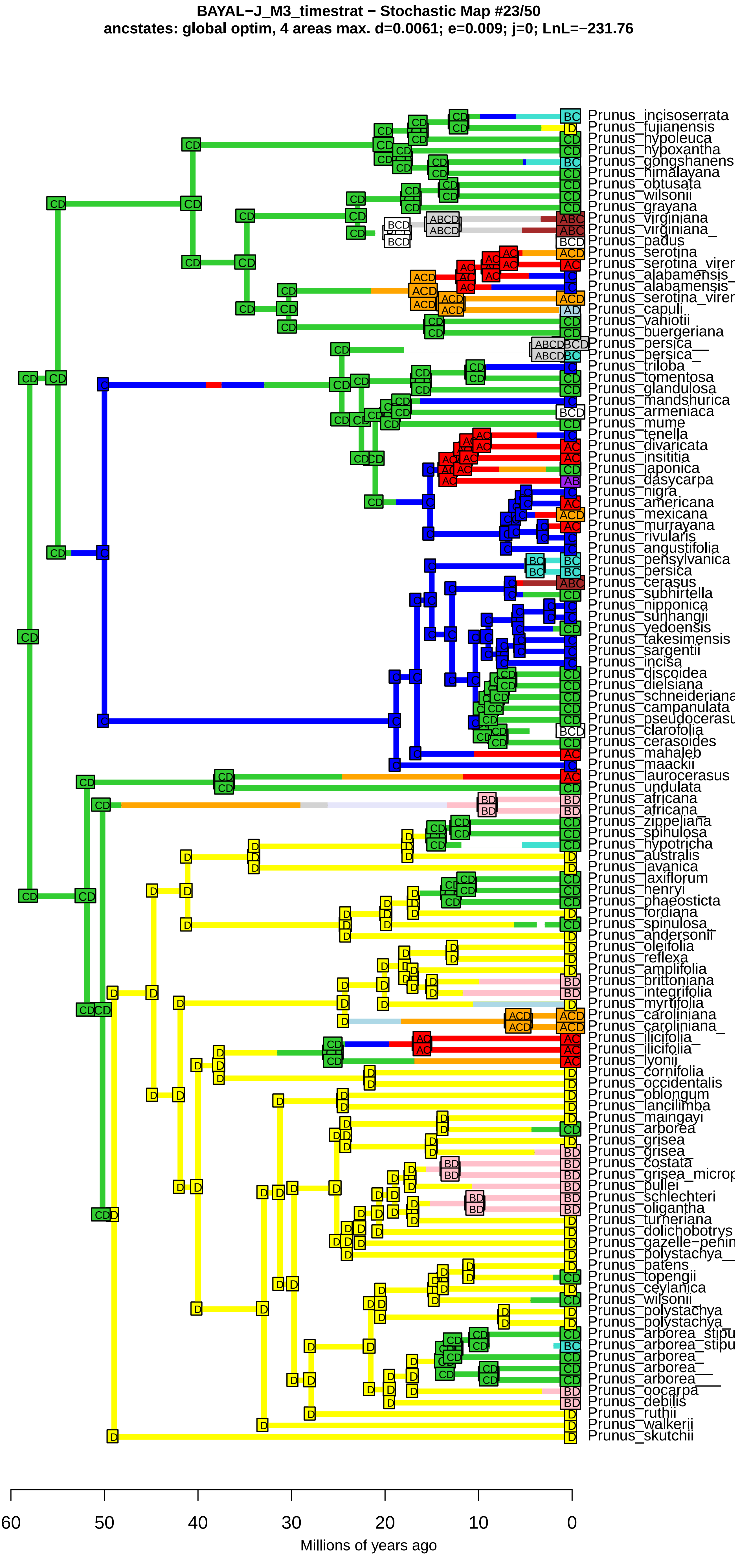
BAYAL−J_M3_timestrat − Stochastic Map #23/50
ancstates: global optim, 4 areas max. d=0.0061; e=0.009; j=0; LnL=−231.76
Prunus_incisoserrata
Prunus_fujianensis
Prunus_hypoleuca
Prunus_hypoxantha
Prunus_gongshanens
Prunus_himalayana
Prunus_obtusata
Prunus_wilsonii
Prunus_grayana
Prunus_virginiana
Prunus_virginiana_
Prunus_padus
Prunus_serotina
Prunus_serotina_virer
Prunus_alabamensis_
Prunus_alabamensis
Prunus_serotina_virer
Prunus_capuli
Prunus_vaniotii
Prunus_buergeriana
Prunus_persica__
Prunus_persica_
Prunus_triloba
Prunus_tomentosa
Prunus_glandulosa
Prunus_mandshurica
Prunus_armeniaca
Prunus_mume
Prunus_tenella
Prunus_divaricata
Prunus_insititia
Prunus_japonica
Prunus_dasycarpa
Prunus_nigra
Prunus_americana
Prunus_mexicana
Prunus_murrayana
Prunus_rivularis
Prunus_angustifolia
Prunus_pensylvanica
Prunus_persica
Prunus_cerasus
Prunus_subhirtella
Prunus_nipponica
Prunus_sunhangii
Prunus_yedoensis
Prunus_takesimensis
Prunus_sargentii
Prunus_incisa
Prunus_discoidea
Prunus_dielsiana
Prunus_schneideriana
Prunus_campanulata
Prunus_pseudocerasu
Prunus_clarofolia
Prunus_cerasoides
Prunus_mahaleb
Prunus_maackii
Prunus_laurocerasus
Prunus_undulata
Prunus_africana
Prunus_africana_
Prunus_zippeliana
Prunus_spinulosa
Prunus_hypotricha
Prunus_australis
Prunus_javanica
Prunus_laxiflorum
Prunus_henryi
Prunus_phaeosticta
Prunus_fordiana
Prunus_spinulosa_
Prunus_andersonii
Prunus_oleifolia
Prunus_reflexa
Prunus_amplifolia
Prunus_brittoniana
Prunus_integrifolia
Prunus_myrtifolia
Prunus_caroliniana
Prunus_caroliniana_
Prunus_ilicifolia_
Prunus_ilicifolia
Prunus_lyonii
Prunus_cornifolia
Prunus_occidentalis
Prunus_oblongum
Prunus_lancilimba
Prunus_maingayi
Prunus_arborea
Prunus_grisea
Prunus_grisea_
Prunus_costata
Prunus_grisea_microp
Prunus_pullei
Prunus_schlechteri
Prunus_oligantha
Prunus_turneriana
Prunus_dolichobotrys
Prunus_gazelle−penin
Prunus_polystachya__
Prunus_patens
Prunus_topengii
Prunus_ceylanica
Prunus_wilsonii_
Prunus_polystachya
Prunus_polystachya_
Prunus_arborea_stipu
Prunus_arborea_stipu
Prunus_arborea_
Prunus_arborea__
Prunus_arborea___
Prunus_oocarpa
Prunus_debilis
Prunus_ruthii
Prunus_walkerii
Prunus_skutchii
60
50
40
30
20
10
0
Millions of years ago

**BAYAL−J_M3_timestrat − Stochastic Map #24/50**
**ancstates: global optim, 4 areas max. d=0.0061; e=0.009; j=0; LnL=−231.76**

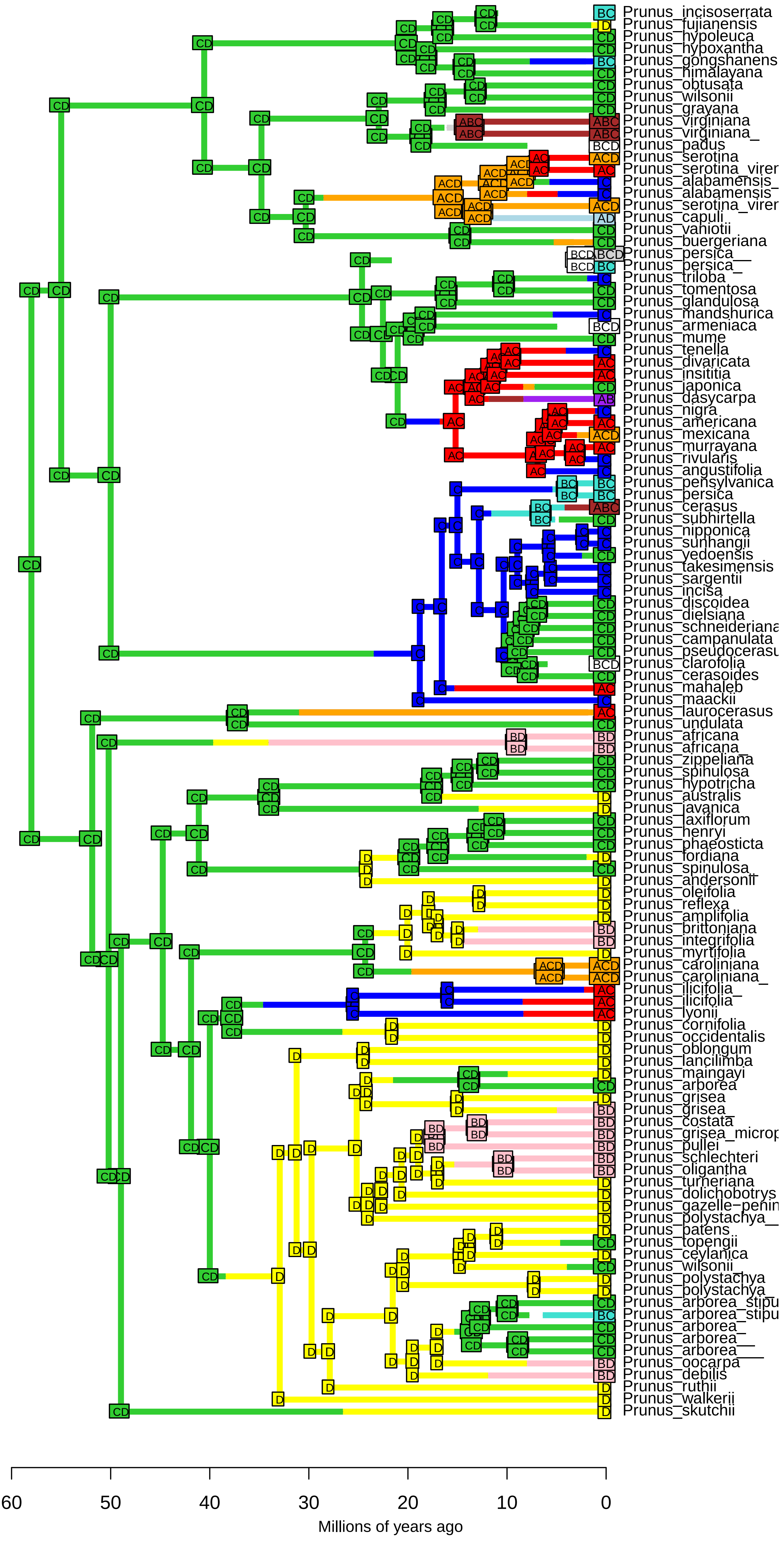

BAYAL−J_M3_timestrat − Stochastic Map #25/50
ancstates: global optim, 4 areas max. d=0.0061; e=0.009; j=0; LnL=−231.76

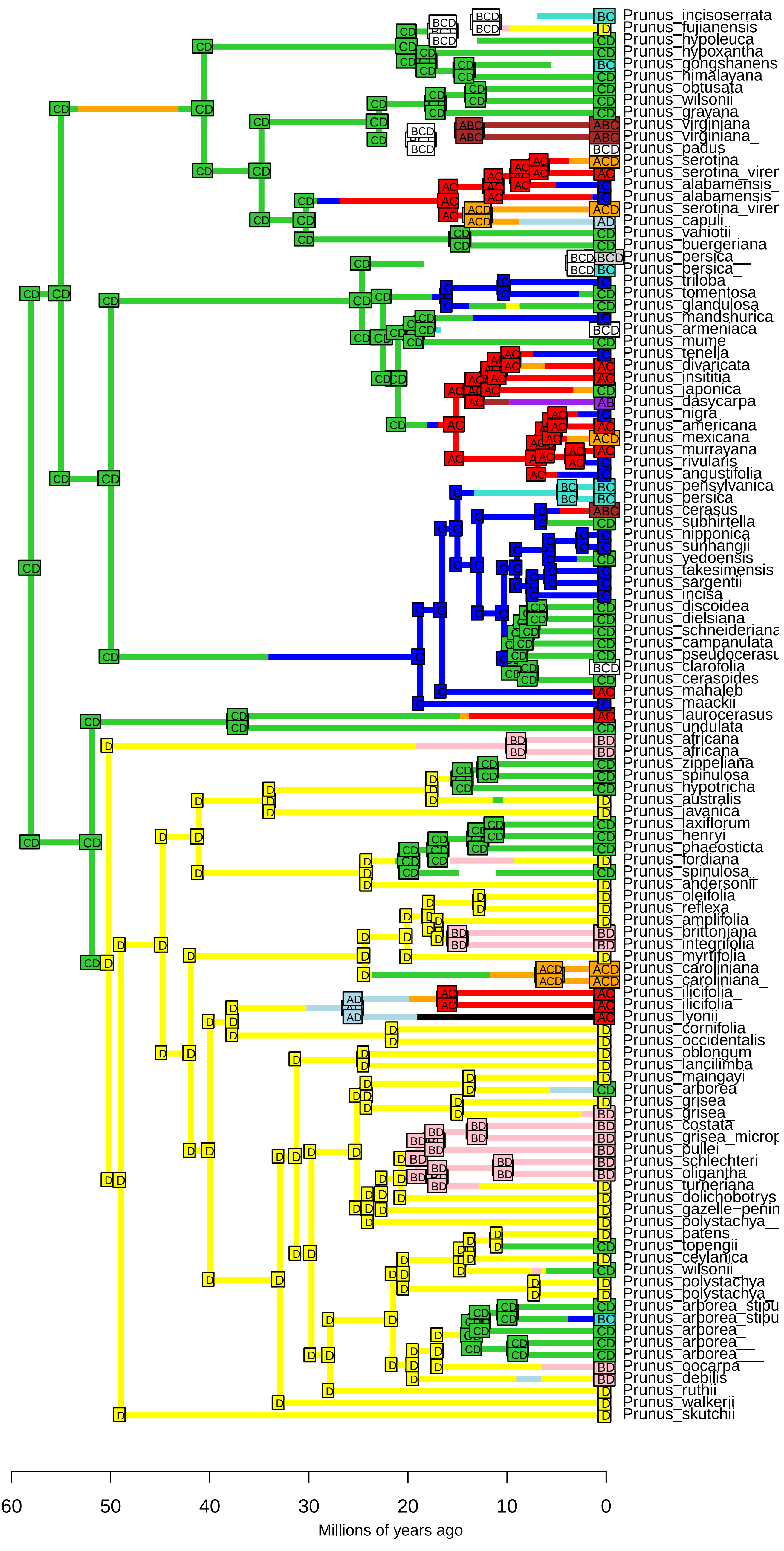

**BAYAL−J_M3_timestrat − Stochastic Map #26/50**
**ancstates: global optim, 4 areas max. d=0.0061; e=0.009; j=0; LnL=−231.76**

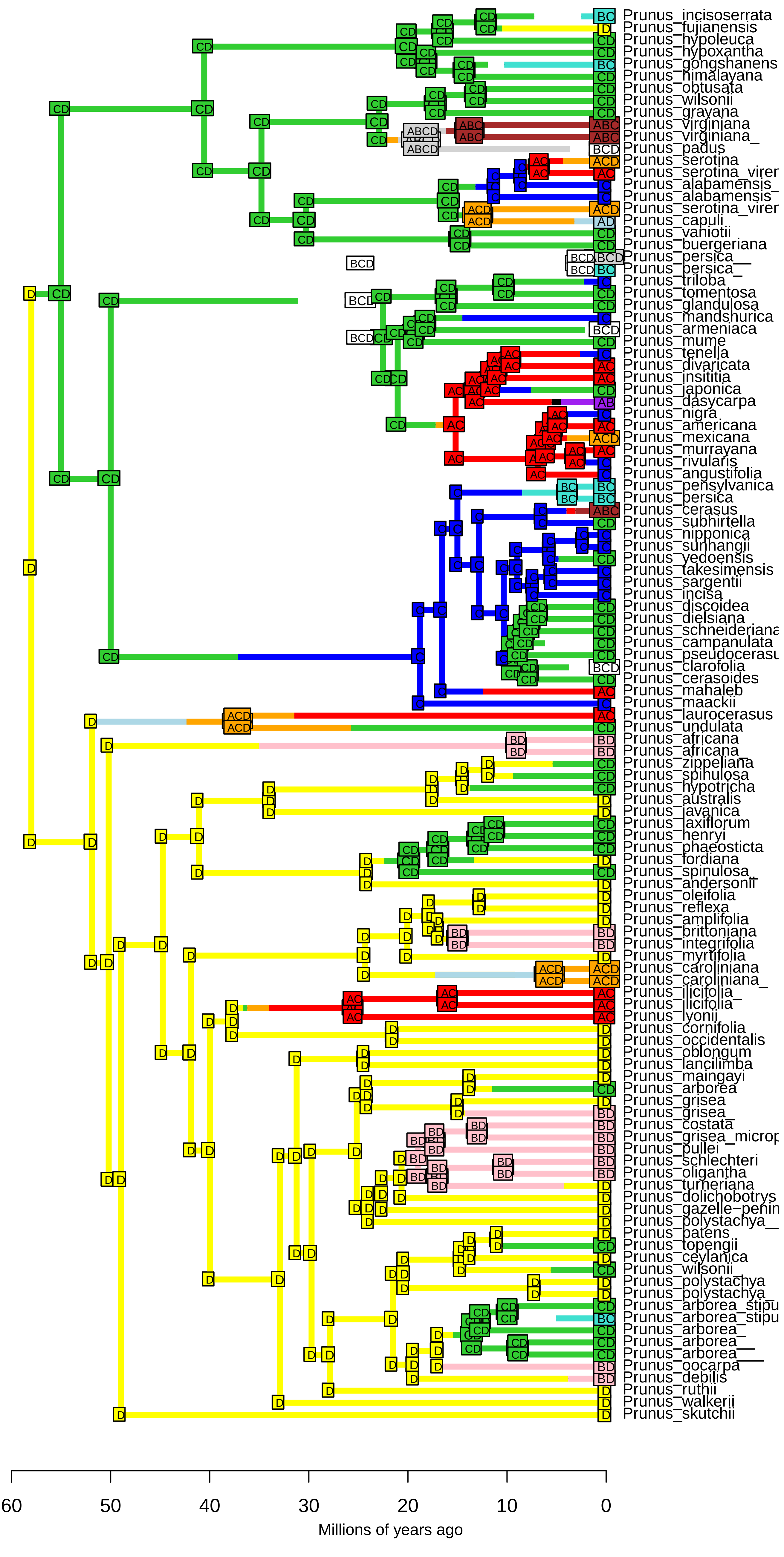

BAYAL−J_M3_timestrat − Stochastic Map #27/50
ancstates: global optim, 4 areas max. d=0.0061; e=0.009; j=0; LnL=−231.76

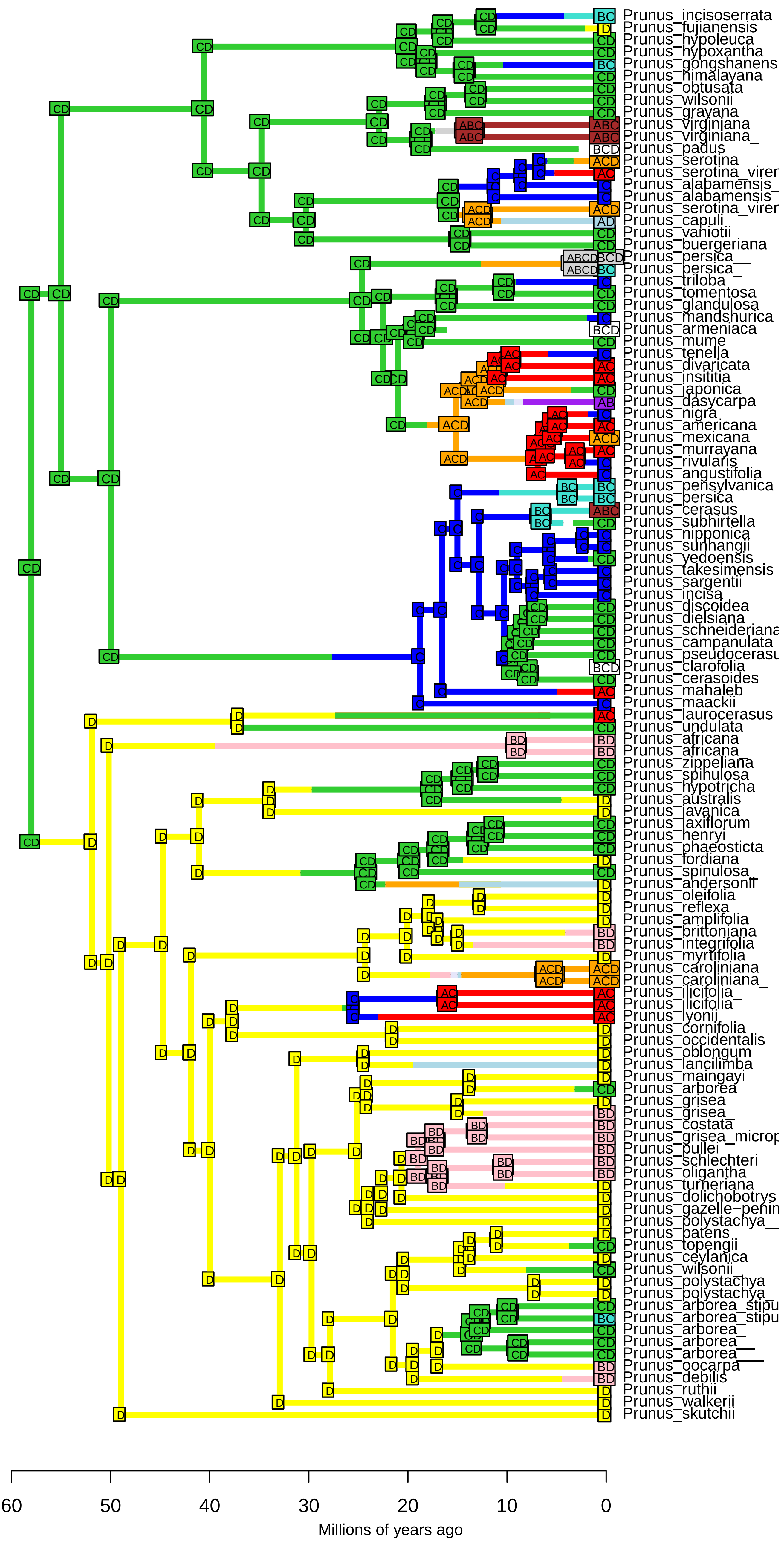

BAYAL−J_M3_timestrat − Stochastic Map #28/50
ancstates: global optim, 4 areas max. d=0.0061; e=0.009; j=0; LnL=−231.76

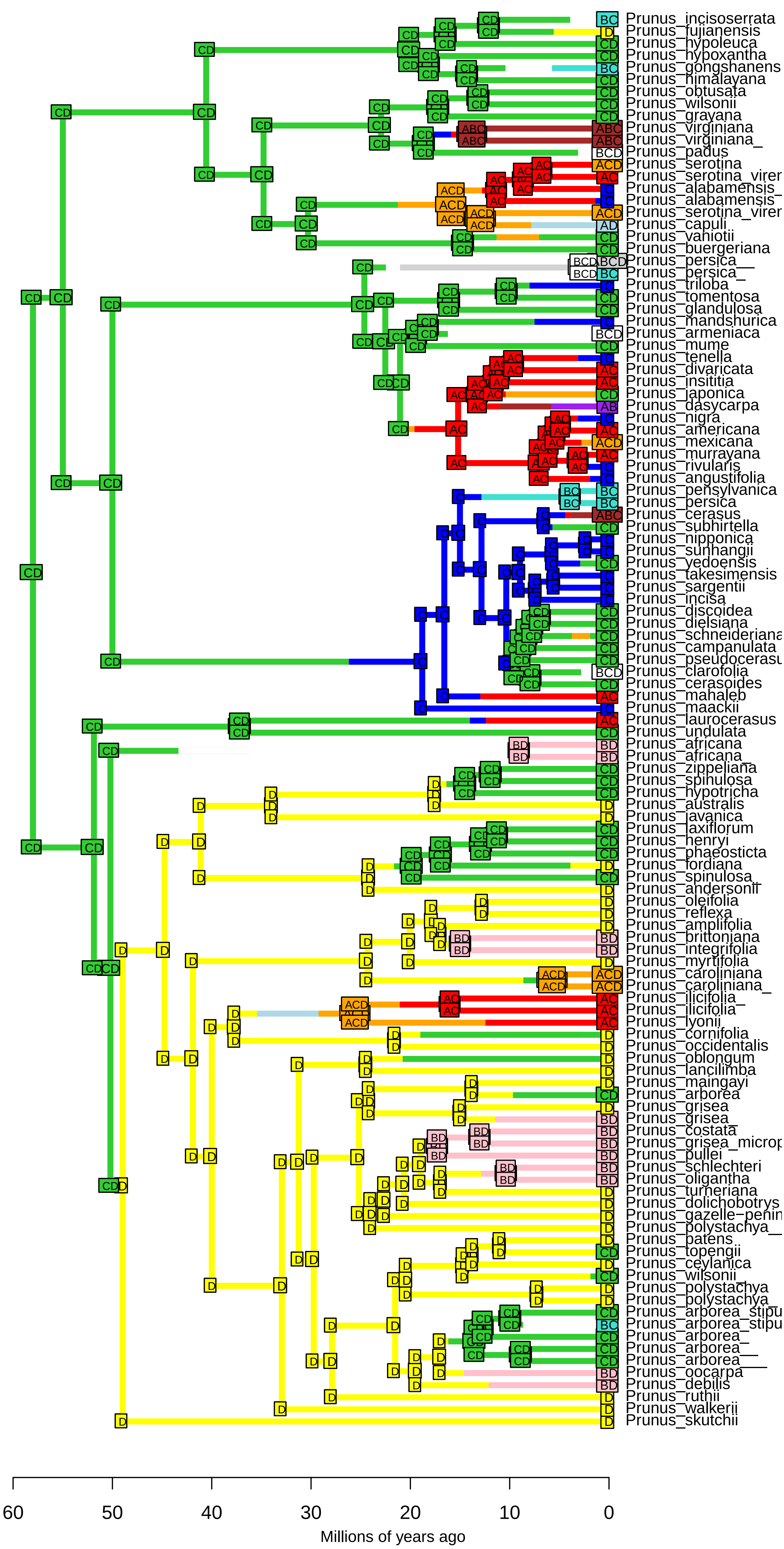

**BAYAL−J_M3_timestrat − Stochastic Map #29/50**
**ancstates: global optim, 4 areas max. d=0.0061; e=0.009; j=0; LnL=−231.76**

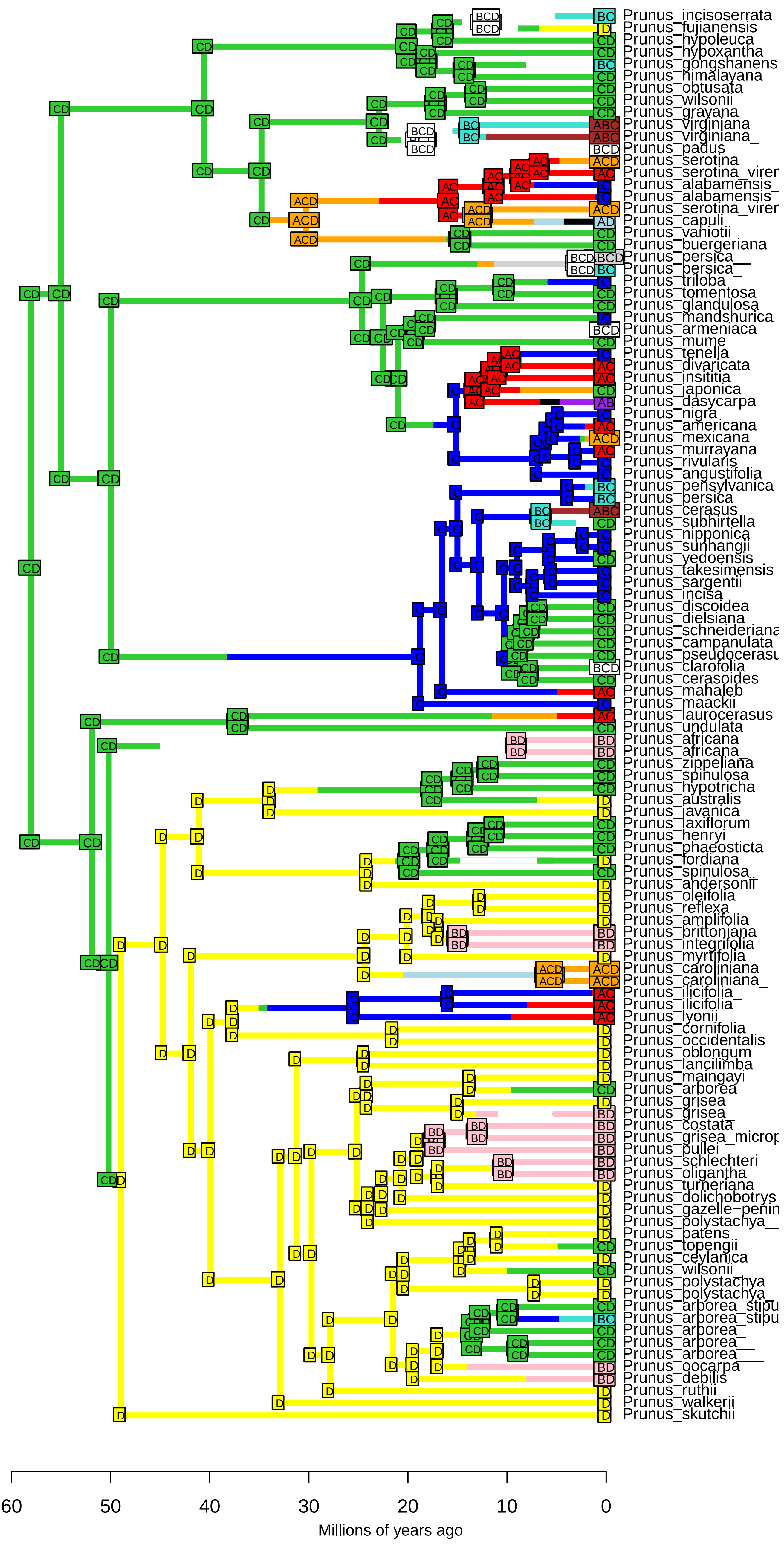

**BAYAL−J_M3_timestrat − Stochastic Map #30/50**
**ancstates: global optim, 4 areas max. d=0.0061; e=0.009; j=0; LnL=−231.76**

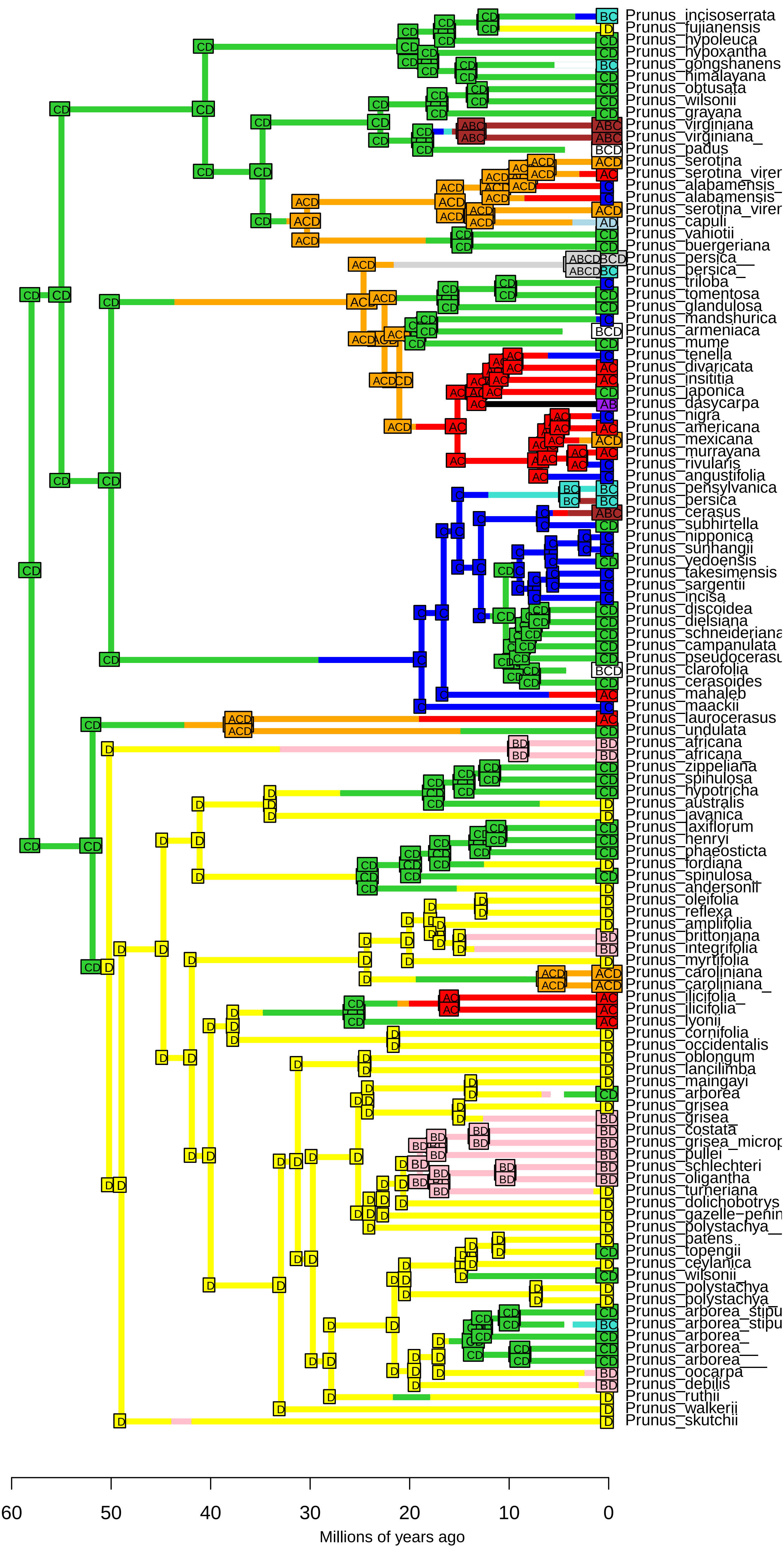

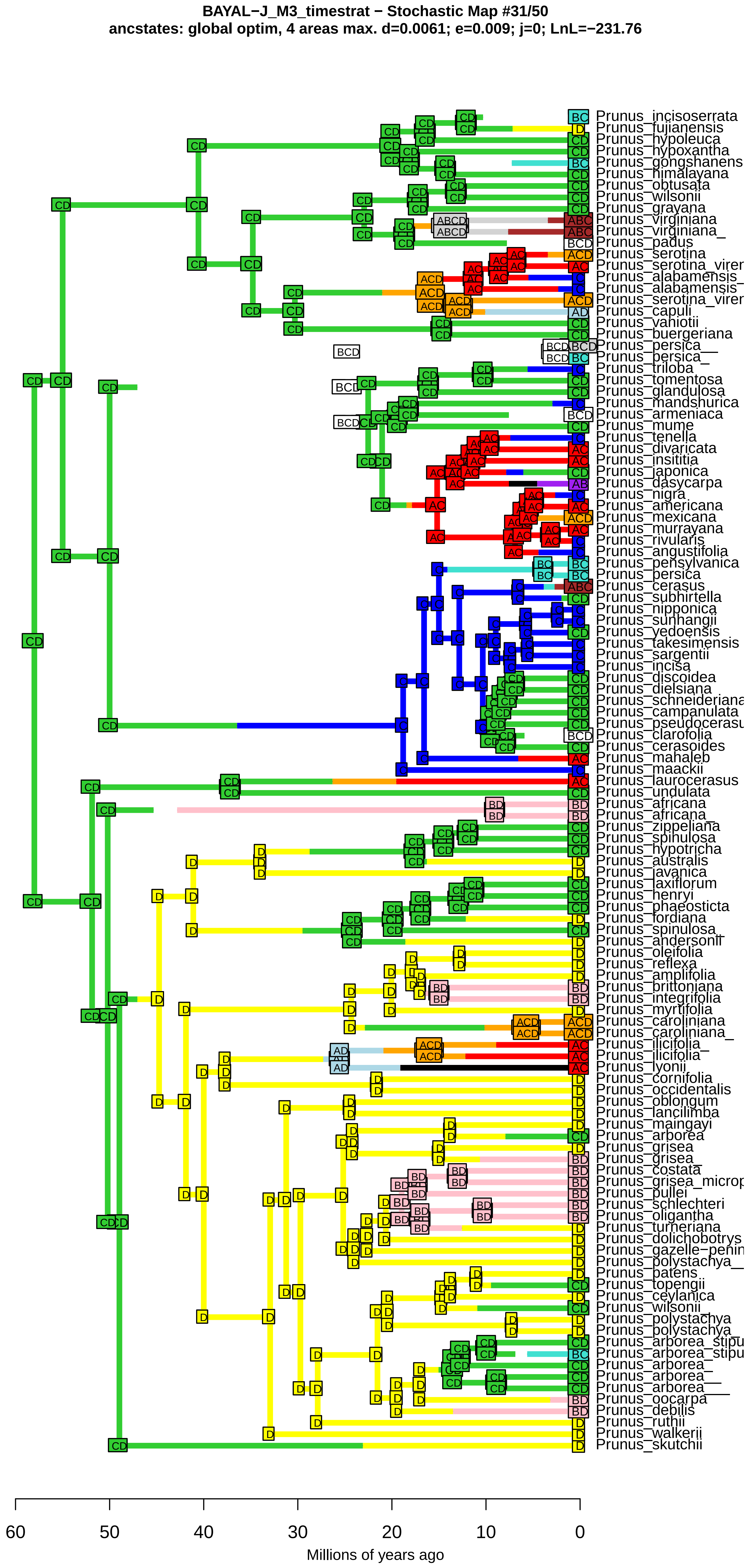
BAYAL−J_M3_timestrat − Stochastic Map #31/50
ancstates: global optim, 4 areas max. d=0.0061; e=0.009; j=0; LnL=−231.76
Prunus_incisoserrata
Prunus_fujianensis
Prunus_hypoleuca
Prunus_hypoxantha
Prunus_gongshanens
Prunus_himalayana
Prunus_obtusata
Prunus_wilsonii
Prunus_grayana
Prunus_virginiana
Prunus_virginiana_
Prunus_padus
Prunus_serotina
Prunus_serotina_virer
Prunus_alabamensis_
Prunus_alabamensis
Prunus_serotina_virer
Prunus_capuli
Prunus_vaniotii
Prunus_buergeriana
Prunus_persica__
Prunus_persica_
Prunus_triloba
Prunus_tomentosa
Prunus_glandulosa
Prunus_mandshurica
Prunus_armeniaca
Prunus_mume
Prunus_tenella
Prunus_divaricata
Prunus_insititia
Prunus_japonica
Prunus_dasycarpa
Prunus_nigra
Prunus_americana
Prunus_mexicana
Prunus_murrayana
Prunus_rivularis
Prunus_angustifolia
Prunus_pensylvanica
Prunus_persica
Prunus_cerasus
Prunus_subhirtella
Prunus_nipponica
Prunus_sunhangii
Prunus_yedoensis
Prunus_takesimensis
Prunus_sargentii
Prunus_incisa
Prunus_discoidea
Prunus_dielsiana
Prunus_schneideriana
Prunus_campanulata
Prunus_pseudocerasu
Prunus_clarofolia
Prunus_cerasoides
Prunus_mahaleb
Prunus_maackii
Prunus_laurocerasus
Prunus_undulata
Prunus_africana
Prunus_africana_
Prunus_zippeliana
Prunus_spinulosa
Prunus_hypotricha
Prunus_australis
Prunus_javanica
Prunus_laxiflorum
Prunus_henryi
Prunus_phaeosticta
Prunus_fordiana
Prunus_spinulosa_
Prunus_andersonii
Prunus_oleifolia
Prunus_reflexa
Prunus_amplifolia
Prunus_brittoniana
Prunus_integrifolia
Prunus_myrtifolia
Prunus_caroliniana
Prunus_caroliniana_
Prunus_ilicifolia_
Prunus_ilicifolia
Prunus_lyonii
Prunus_cornifolia
Prunus_occidentalis
Prunus_oblongum
Prunus_lancilimba
Prunus_maingayi
Prunus_arborea
Prunus_grisea
Prunus_grisea_
Prunus_costata
Prunus_grisea_microp
Prunus_pullei
Prunus_schlechteri
Prunus_oligantha
Prunus_turneriana
Prunus_dolichobotrys
Prunus_gazelle−penin
Prunus_polystachya__
Prunus_patens
Prunus_topengii
Prunus_ceylanica
Prunus_wilsonii_
Prunus_polystachya
Prunus_polystachya_
Prunus_arborea_stipu
Prunus_arborea_stipu
Prunus_arborea_
Prunus_arborea__
Prunus_arborea___
Prunus_oocarpa
Prunus_debilis
Prunus_ruthii
Prunus_walkerii
Prunus_skutchii
60
50
40
30
20
10
0
Millions of years ago

**BAYAL−J_M3_timestrat − Stochastic Map #32/50**
**ancstates: global optim, 4 areas max. d=0.0061; e=0.009; j=0; LnL=−231.76**

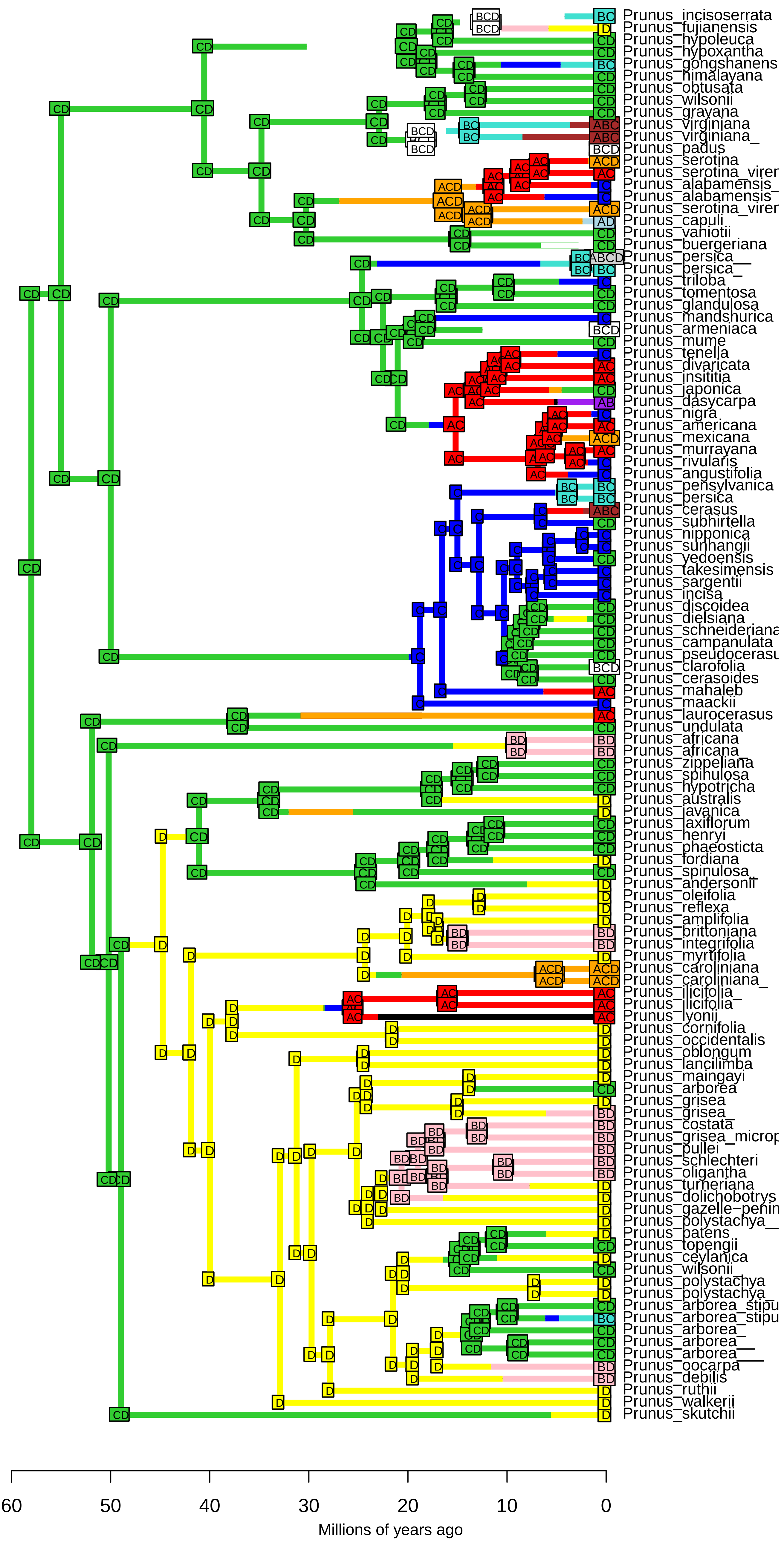

**BAYAL−J_M3_timestrat − Stochastic Map #33/50**
**ancstates: global optim, 4 areas max. d=0.0061; e=0.009; j=0; LnL=−231.76**

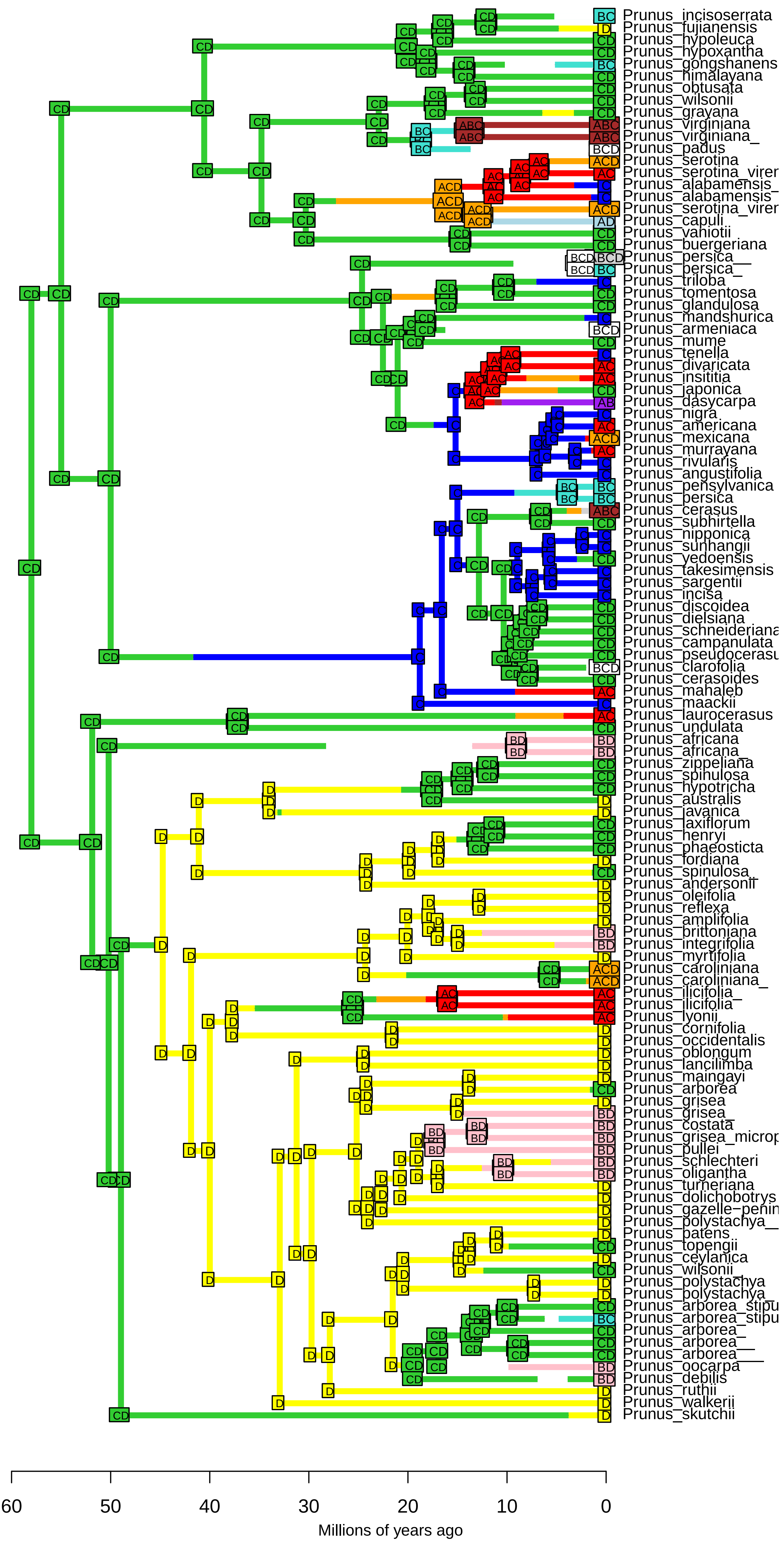

**BAYAL−J_M3_timestrat − Stochastic Map #34/50**
**ancstates: global optim, 4 areas max. d=0.0061; e=0.009; j=0; LnL=−231.76**

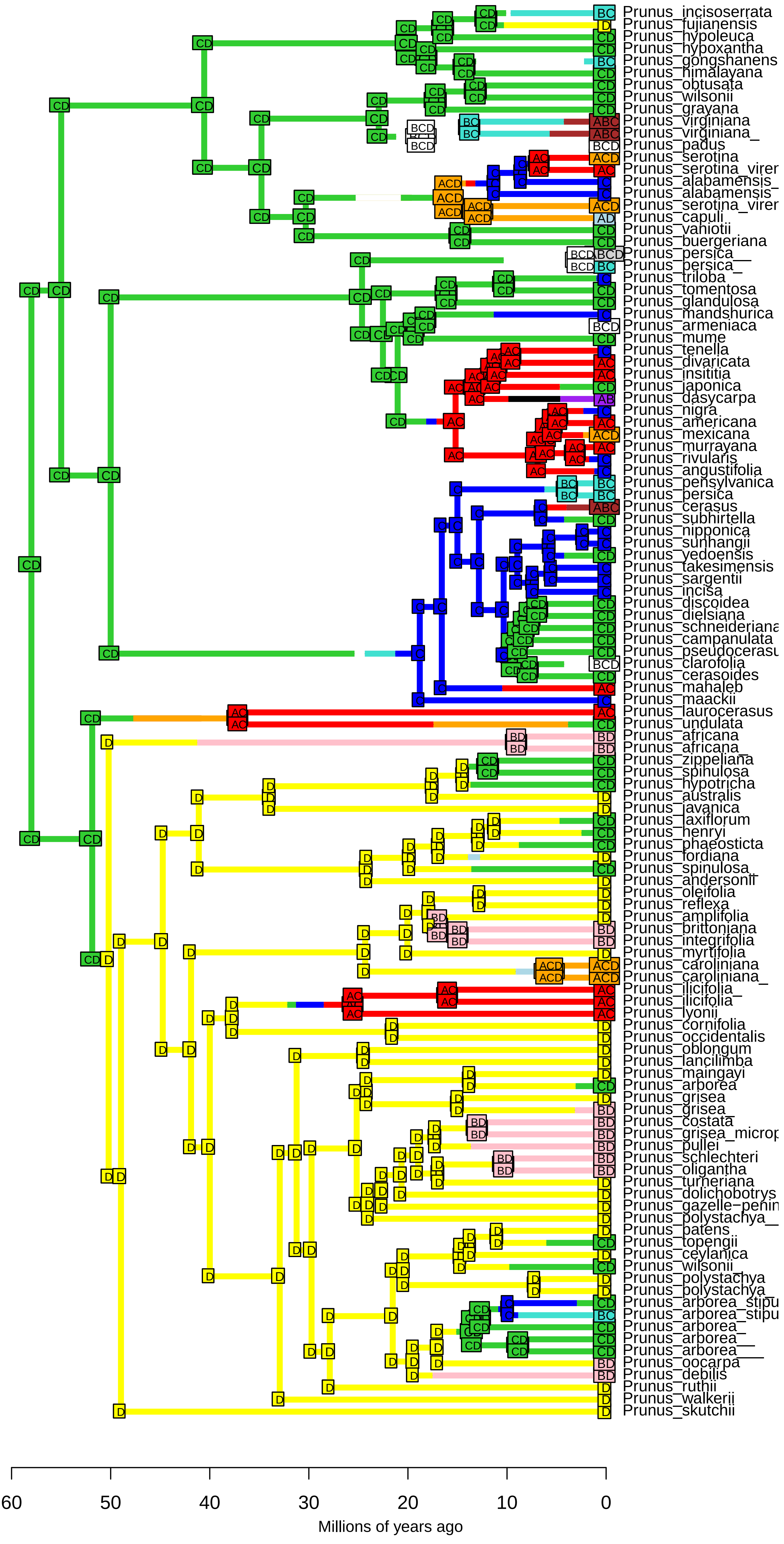

BAYAL−J_M3_timestrat − Stochastic Map #35/50
ancstates: global optim, 4 areas max. d=0.0061; e=0.009; j=0; LnL=−231.76

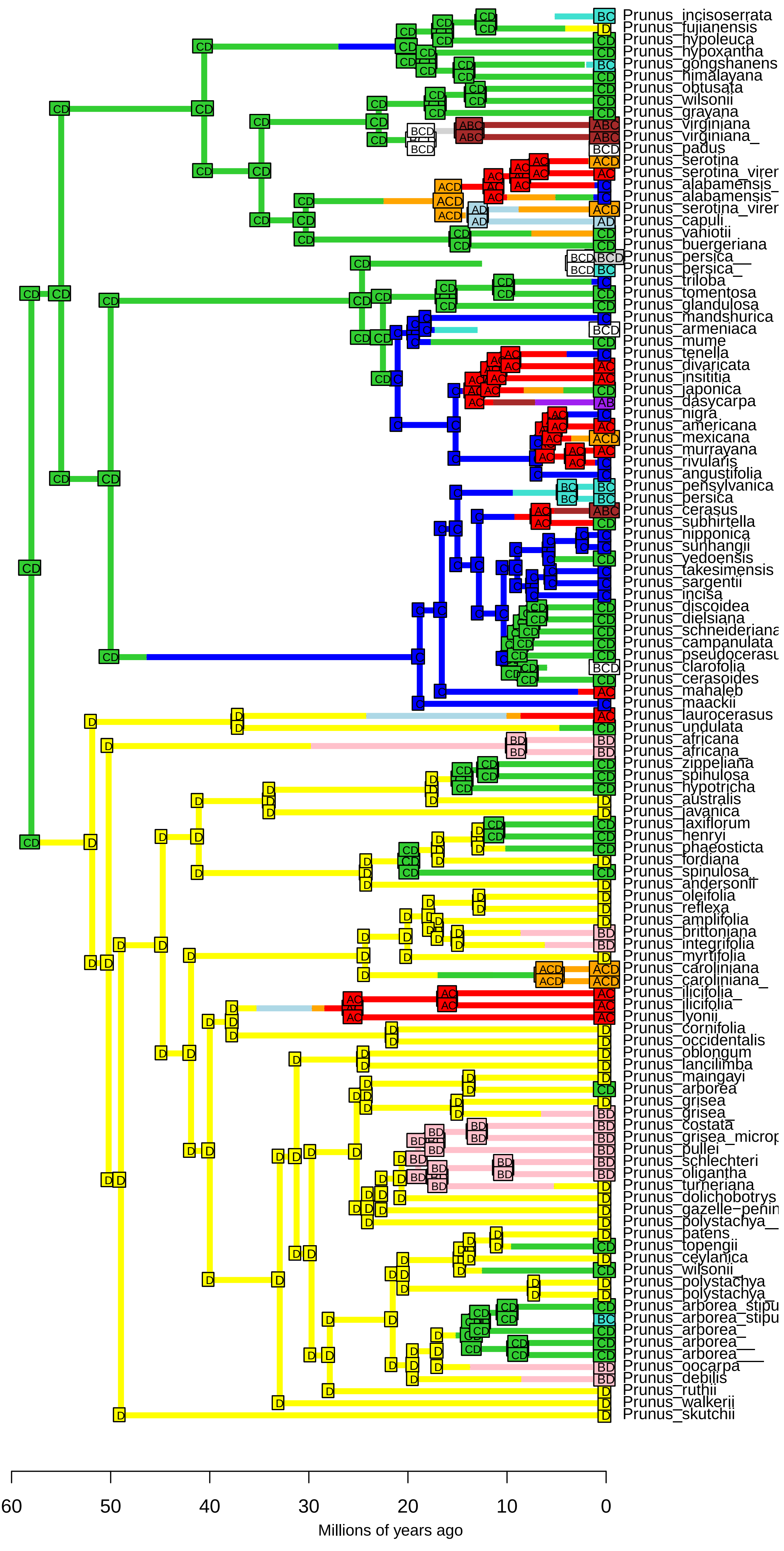

**BAYAL−J_M3_timestrat − Stochastic Map #36/50**
**ancstates: global optim, 4 areas max. d=0.0061; e=0.009; j=0; LnL=−231.76**

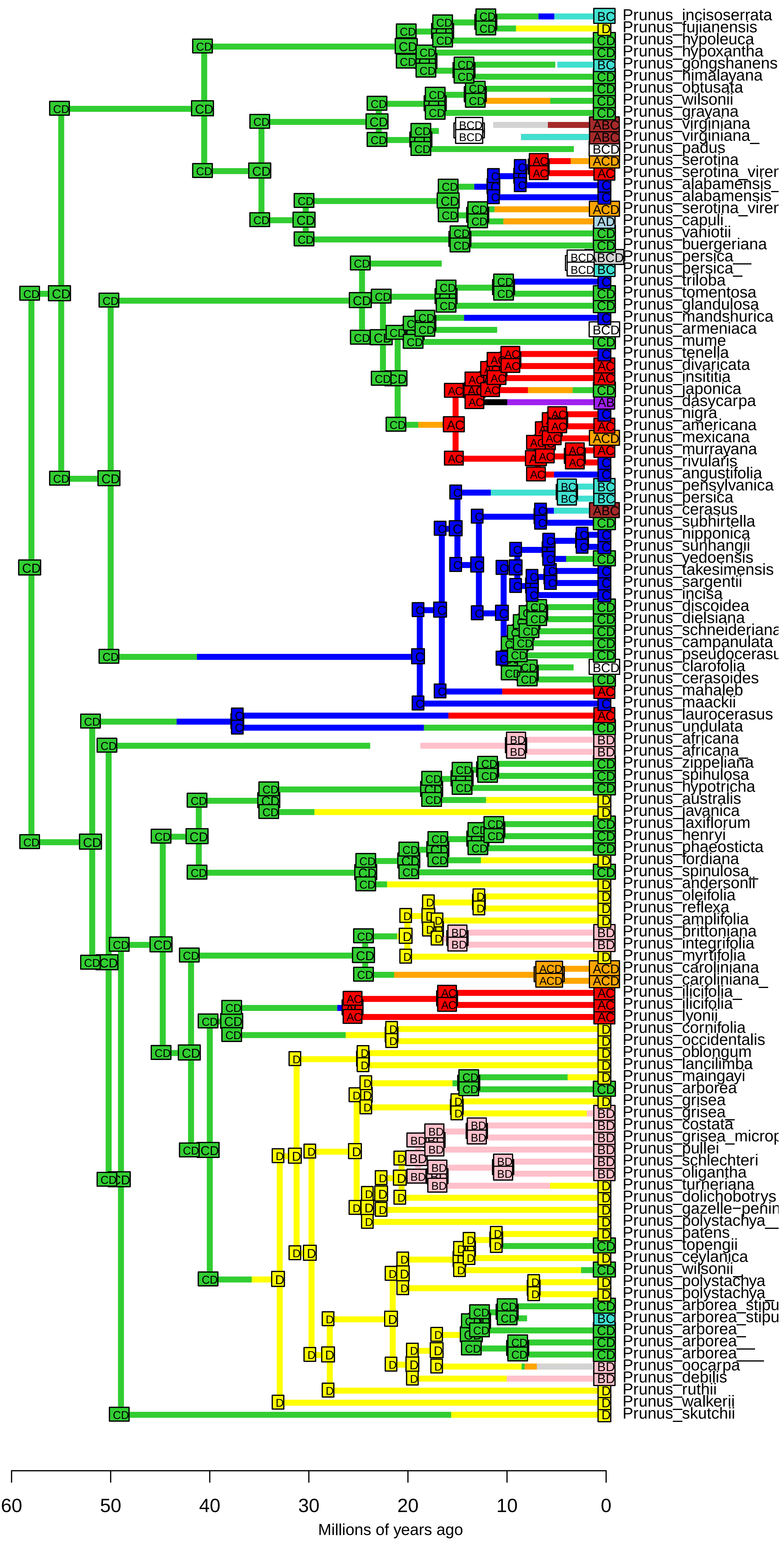

BAYAL−J_M3_timestrat − Stochastic Map #37/50
ancstates: global optim, 4 areas max. d=0.0061; e=0.009; j=0; LnL=−231.76

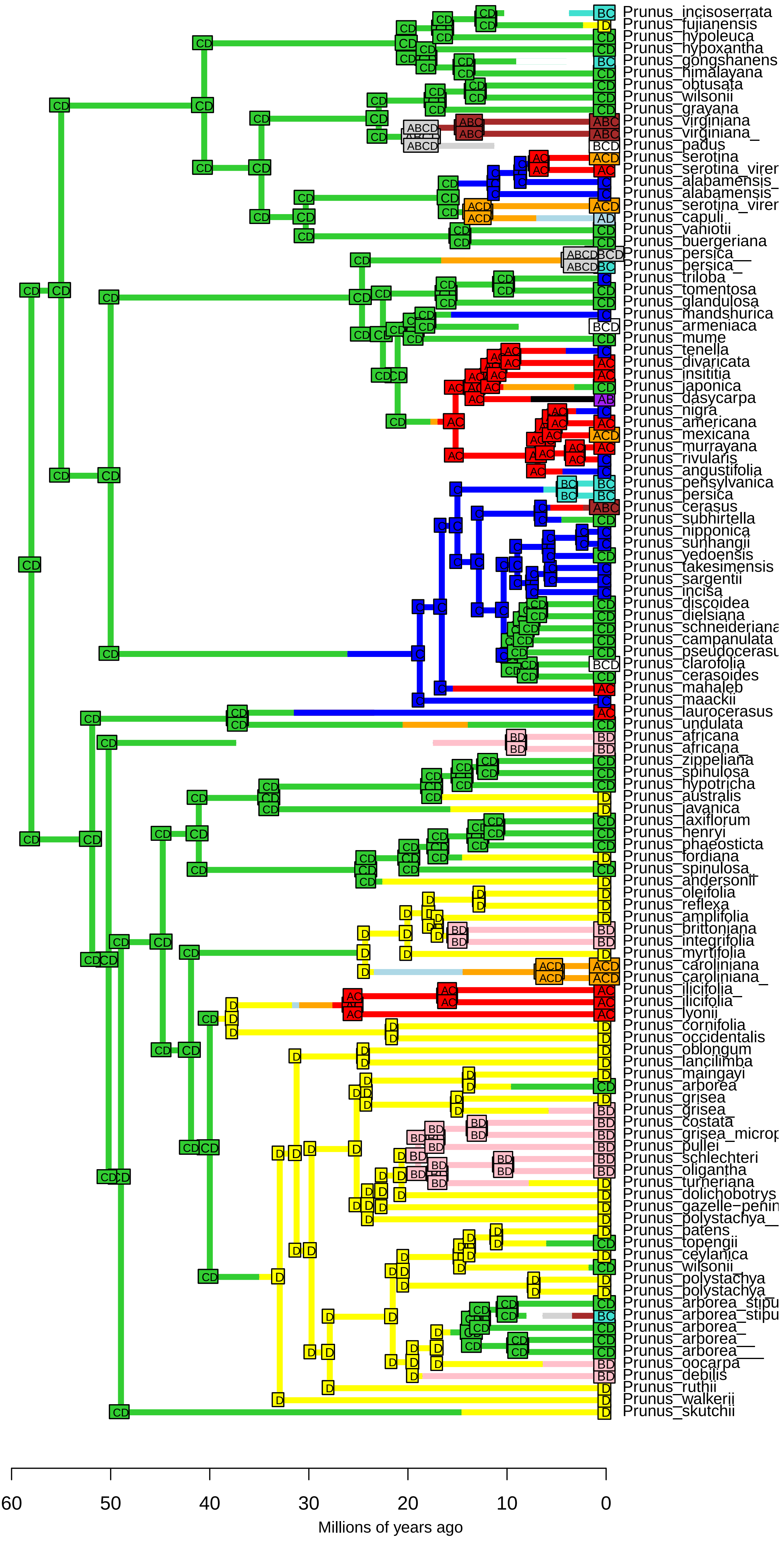

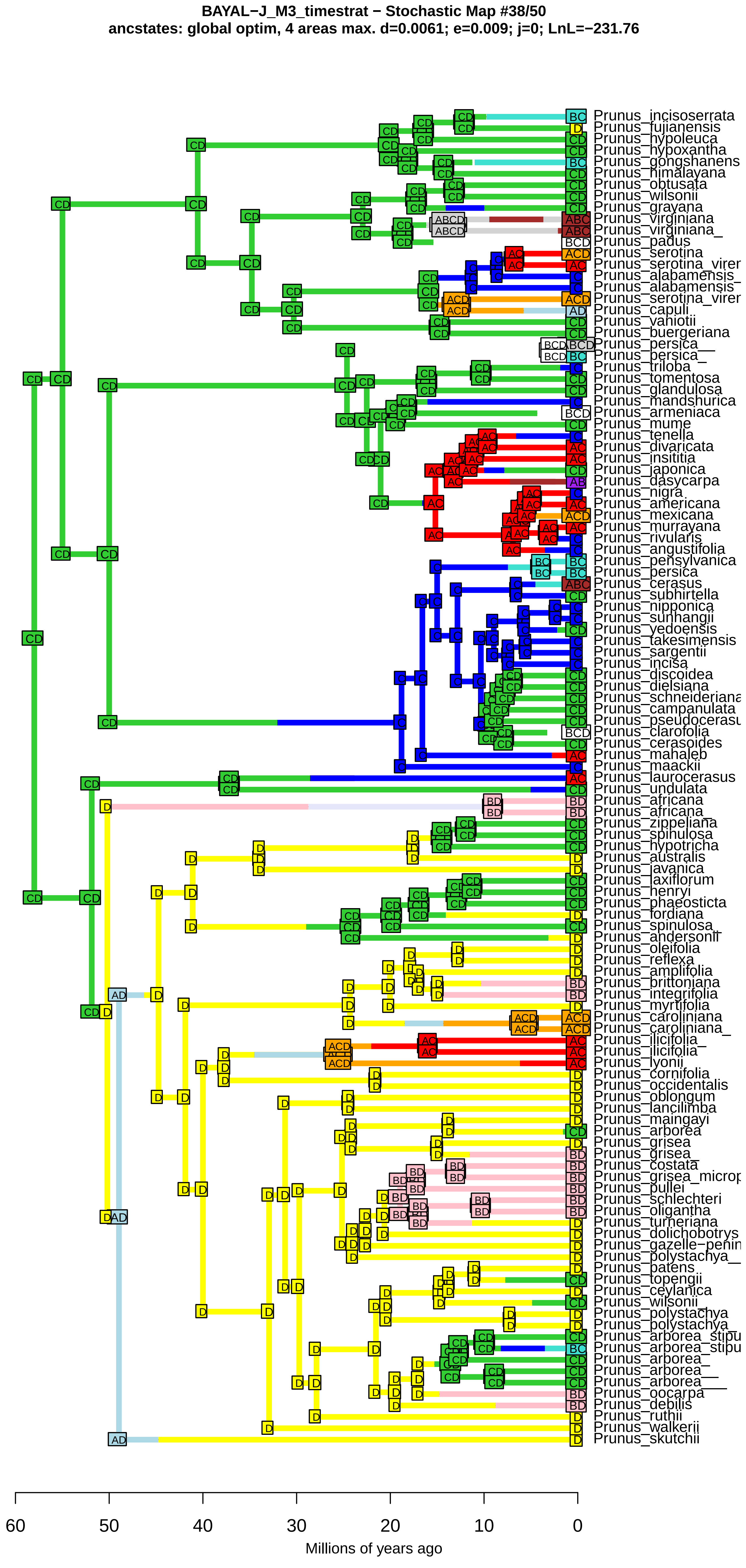
BAYAL−J_M3_timestrat − Stochastic Map #38/50
ancstates: global optim, 4 areas max. d=0.0061; e=0.009; j=0; LnL=−231.76
Prunus_incisoserrata
Prunus_fujianensis
Prunus_hypoleuca
Prunus_hypoxantha
Prunus_gongshanens
Prunus_himalayana
Prunus_obtusata
Prunus_wilsonii
Prunus_grayana
Prunus_virginiana
Prunus_virginiana_
Prunus_padus
Prunus_serotina
Prunus_serotina_virer
Prunus_alabamensis_
Prunus_alabamensis
Prunus_serotina_virer
Prunus_capuli
Prunus_vaniotii
Prunus_buergeriana
Prunus_persica__
Prunus_persica_
Prunus_triloba
Prunus_tomentosa
Prunus_glandulosa
Prunus_mandshurica
Prunus_armeniaca
Prunus_mume
Prunus_tenella
Prunus_divaricata
Prunus_insititia
Prunus_japonica
Prunus_dasycarpa
Prunus_nigra
Prunus_americana
Prunus_mexicana
Prunus_murrayana
Prunus_rivularis
Prunus_angustifolia
Prunus_pensylvanica
Prunus_persica
Prunus_cerasus
Prunus_subhirtella
Prunus_nipponica
Prunus_sunhangii
Prunus_yedoensis
Prunus_takesimensis
Prunus_sargentii
Prunus_incisa
Prunus_discoidea
Prunus_dielsiana
Prunus_schneideriana
Prunus_campanulata
Prunus_pseudocerasu
Prunus_clarofolia
Prunus_cerasoides
Prunus_mahaleb
Prunus_maackii
Prunus_laurocerasus
Prunus_undulata
Prunus_africana
Prunus_africana_
Prunus_zippeliana
Prunus_spinulosa
Prunus_hypotricha
Prunus_australis
Prunus_javanica
Prunus_laxiflorum
Prunus_henryi
Prunus_phaeosticta
Prunus_fordiana
Prunus_spinulosa_
Prunus_andersonii
Prunus_oleifolia
Prunus_reflexa
Prunus_amplifolia
Prunus_brittoniana
Prunus_integrifolia
Prunus_myrtifolia
Prunus_caroliniana
Prunus_caroliniana_
Prunus_ilicifolia_
Prunus_ilicifolia
Prunus_lyonii
Prunus_cornifolia
Prunus_occidentalis
Prunus_oblongum
Prunus_lancilimba
Prunus_maingayi
Prunus_arborea
Prunus_grisea
Prunus_grisea_
Prunus_costata
Prunus_grisea_microp
Prunus_pullei
Prunus_schlechteri
Prunus_oligantha
Prunus_turneriana
Prunus_dolichobotrys
Prunus_gazelle−penin
Prunus_polystachya__
Prunus_patens
Prunus_topengii
Prunus_ceylanica
Prunus_wilsonii_
Prunus_polystachya
Prunus_polystachya_
Prunus_arborea_stipu
Prunus_arborea_stipu
Prunus_arborea_
Prunus_arborea__
Prunus_arborea___
Prunus_oocarpa
Prunus_debilis
Prunus_ruthii
Prunus_walkerii
Prunus_skutchii
60
50
40
30
20
10
0
Millions of years ago

**BAYAL−J_M3_timestrat − Stochastic Map #39/50**
**ancstates: global optim, 4 areas max. d=0.0061; e=0.009; j=0; LnL=−231.76**

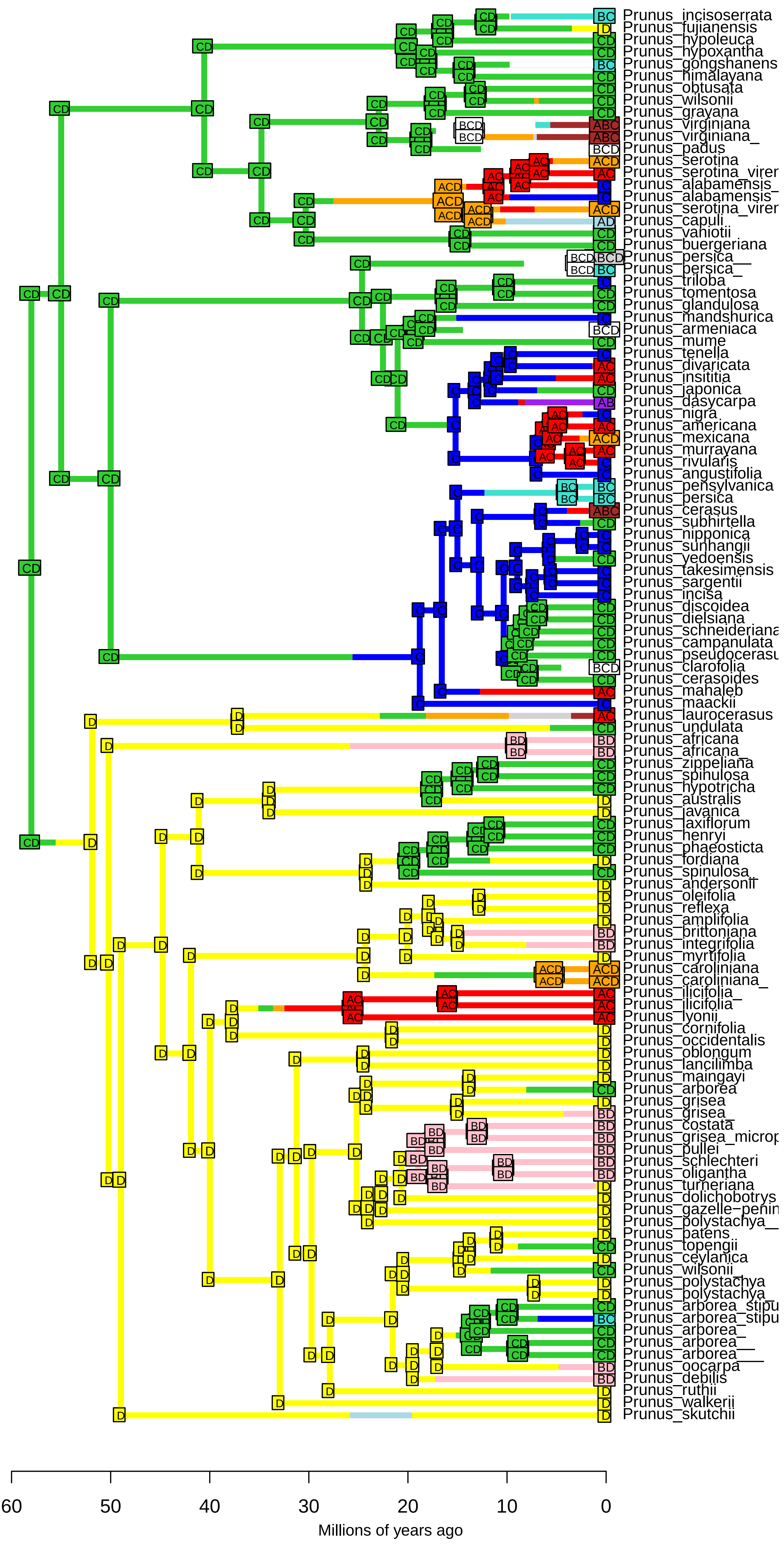

**BAYAL−J_M3_timestrat − Stochastic Map #40/50**
**ancstates: global optim, 4 areas max. d=0.0061; e=0.009; j=0; LnL=−231.76**

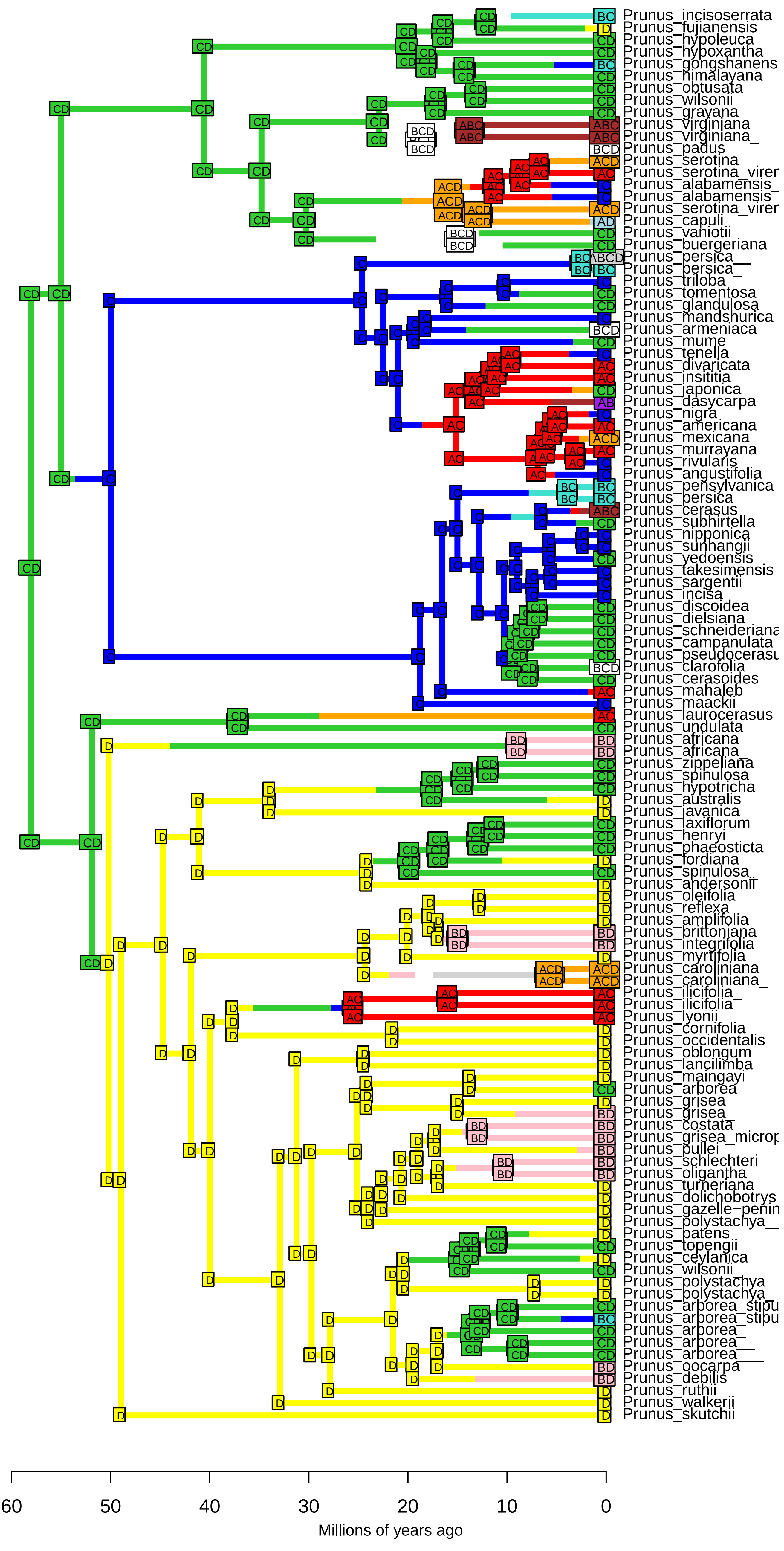

**BAYAL−J_M3_timestrat − Stochastic Map #41/50**
**ancstates: global optim, 4 areas max. d=0.0061; e=0.009; j=0; LnL=−231.76**

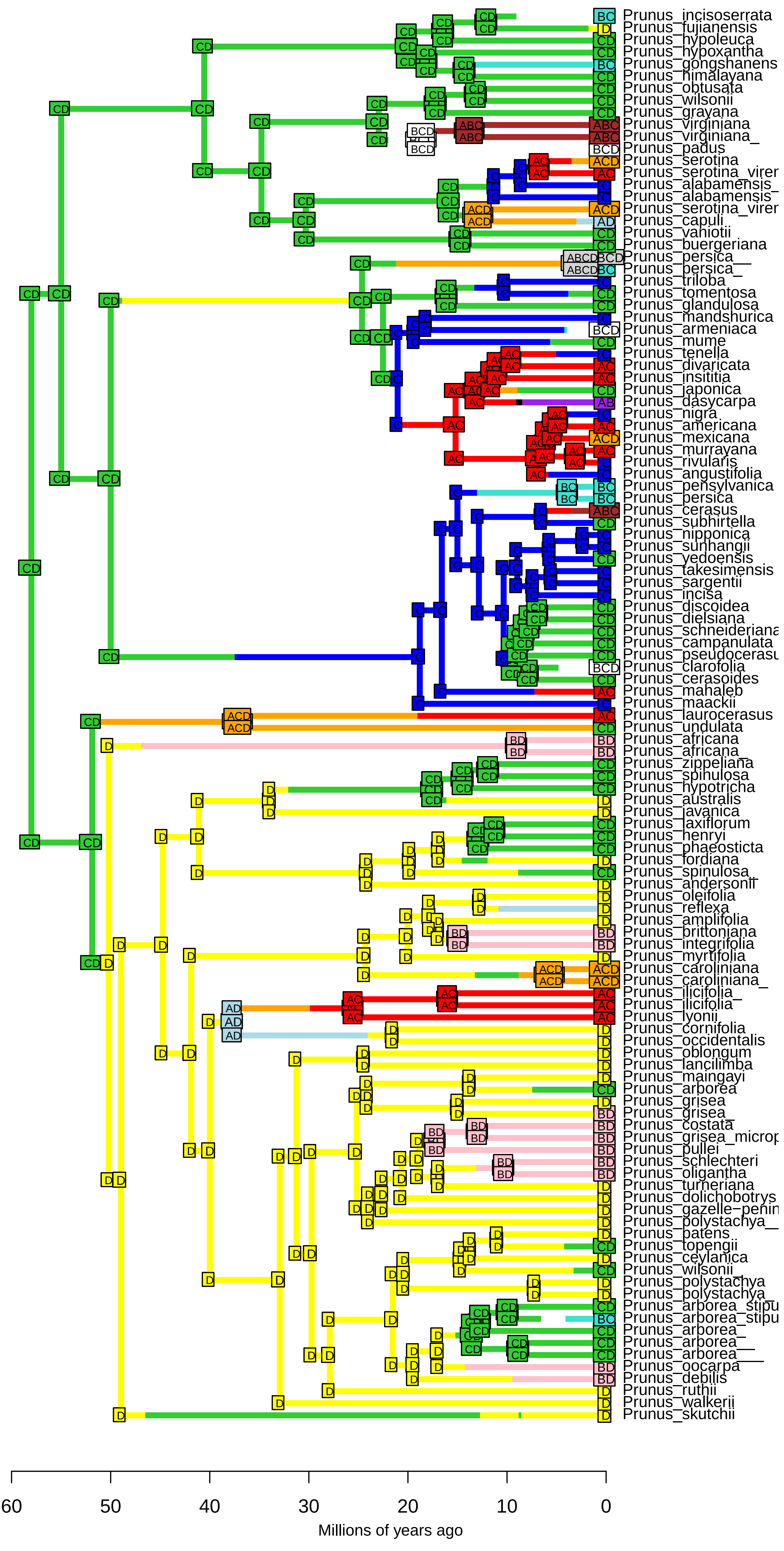

**BAYAL−J_M3_timestrat − Stochastic Map #42/50**
**ancstates: global optim, 4 areas max. d=0.0061; e=0.009; j=0; LnL=−231.76**

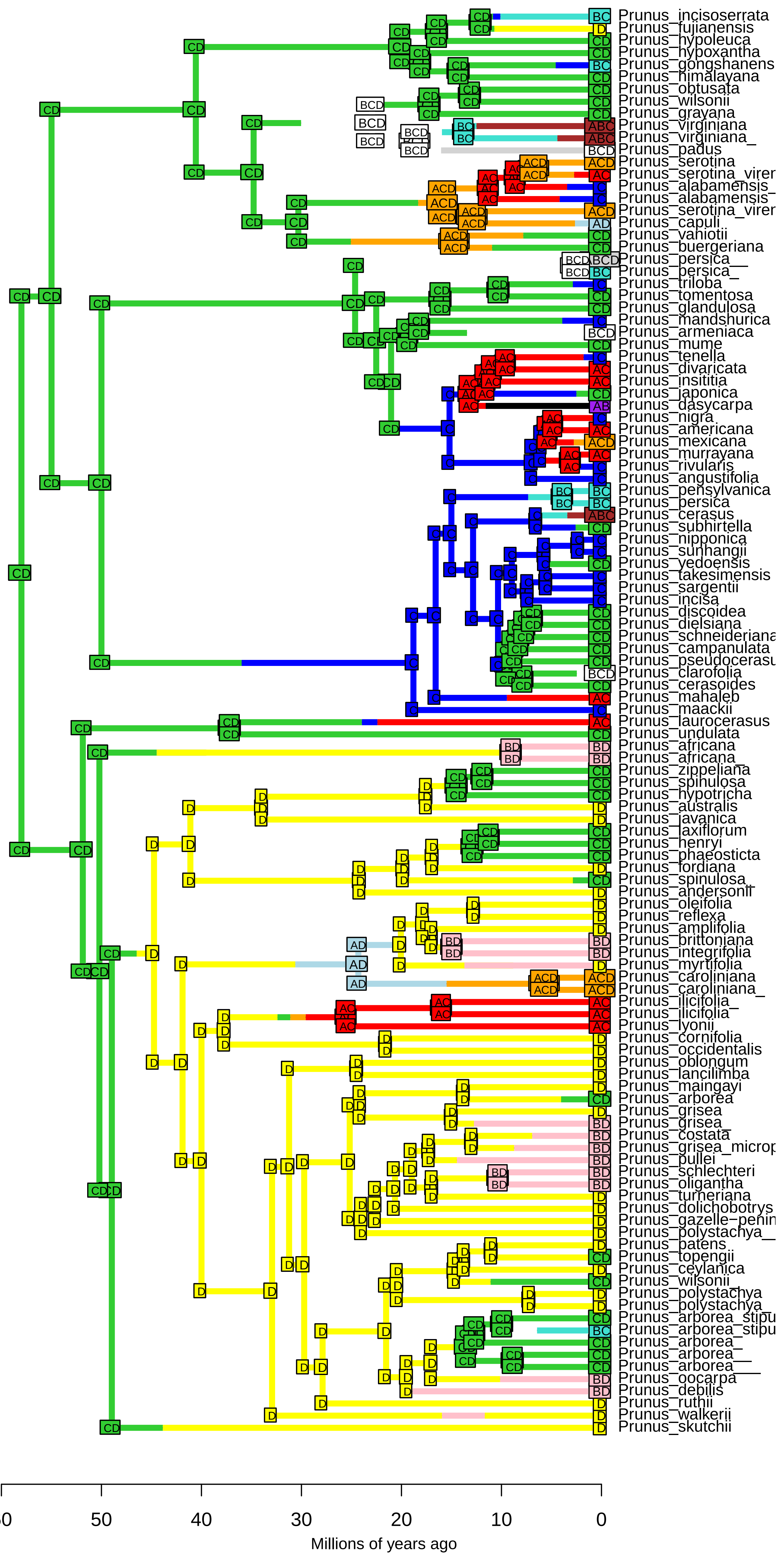

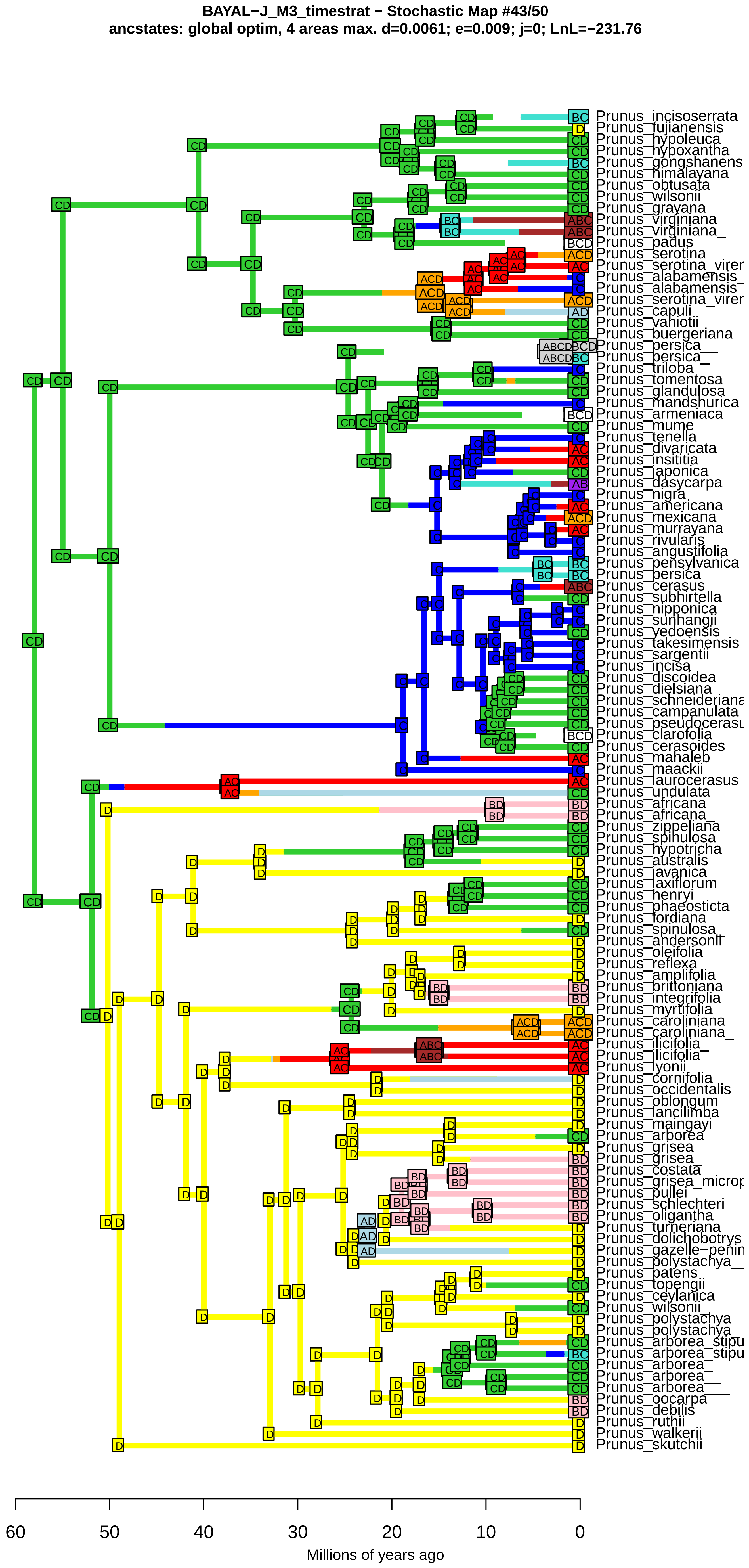
BAYAL−J_M3_timestrat − Stochastic Map #43/50
ancstates: global optim, 4 areas max. d=0.0061; e=0.009; j=0; LnL=−231.76
Prunus_incisoserrata
Prunus_fujianensis
Prunus_hypoleuca
Prunus_hypoxantha
Prunus_gongshanens
Prunus_himalayana
Prunus_obtusata
Prunus_wilsonii
Prunus_grayana
Prunus_virginiana
Prunus_virginiana_
Prunus_padus
Prunus_serotina
Prunus_serotina_virer
Prunus_alabamensis_
Prunus_alabamensis
Prunus_serotina_virer
Prunus_capuli
Prunus_vaniotii
Prunus_buergeriana
Prunus_persica__
Prunus_persica_
Prunus_triloba
Prunus_tomentosa
Prunus_glandulosa
Prunus_mandshurica
Prunus_armeniaca
Prunus_mume
Prunus_tenella
Prunus_divaricata
Prunus_insititia
Prunus_japonica
Prunus_dasycarpa
Prunus_nigra
Prunus_americana
Prunus_mexicana
Prunus_murrayana
Prunus_rivularis
Prunus_angustifolia
Prunus_pensylvanica
Prunus_persica
Prunus_cerasus
Prunus_subhirtella
Prunus_nipponica
Prunus_sunhangii
Prunus_yedoensis
Prunus_takesimensis
Prunus_sargentii
Prunus_incisa
Prunus_discoidea
Prunus_dielsiana
Prunus_schneideriana
Prunus_campanulata
Prunus_pseudocerasu
Prunus_clarofolia
Prunus_cerasoides
Prunus_mahaleb
Prunus_maackii
Prunus_laurocerasus
Prunus_undulata
Prunus_africana
Prunus_africana_
Prunus_zippeliana
Prunus_spinulosa
Prunus_hypotricha
Prunus_australis
Prunus_javanica
Prunus_laxiflorum
Prunus_henryi
Prunus_phaeosticta
Prunus_fordiana
Prunus_spinulosa_
Prunus_andersonii
Prunus_oleifolia
Prunus_reflexa
Prunus_amplifolia
Prunus_brittoniana
Prunus_integrifolia
Prunus_myrtifolia
Prunus_caroliniana
Prunus_caroliniana_
Prunus_ilicifolia_
Prunus_ilicifolia
Prunus_lyonii
Prunus_cornifolia
Prunus_occidentalis
Prunus_oblongum
Prunus_lancilimba
Prunus_maingayi
Prunus_arborea
Prunus_grisea
Prunus_grisea_
Prunus_costata
Prunus_grisea_microp
Prunus_pullei
Prunus_schlechteri
Prunus_oligantha
Prunus_turneriana
Prunus_dolichobotrys
Prunus_gazelle−penin
Prunus_polystachya__
Prunus_patens
Prunus_topengii
Prunus_ceylanica
Prunus_wilsonii_
Prunus_polystachya
Prunus_polystachya_
Prunus_arborea_stipu
Prunus_arborea_stipu
Prunus_arborea_
Prunus_arborea__
Prunus_arborea___
Prunus_oocarpa
Prunus_debilis
Prunus_ruthii
Prunus_walkerii
Prunus_skutchii
60
50
40
30
20
10
0
Millions of years ago

**BAYAL−J_M3_timestrat − Stochastic Map #44/50**
**ancstates: global optim, 4 areas max. d=0.0061; e=0.009; j=0; LnL=−231.76**

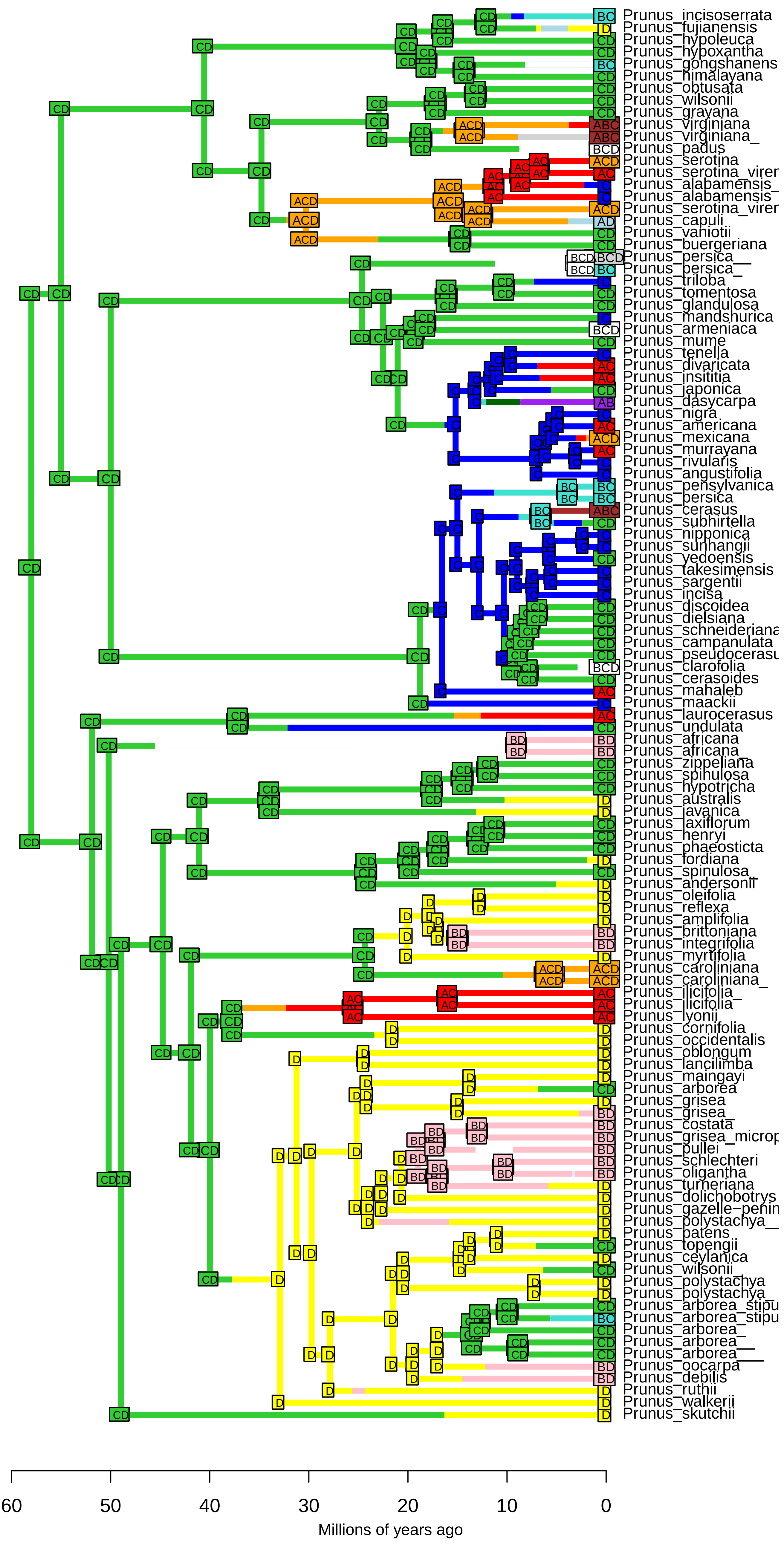

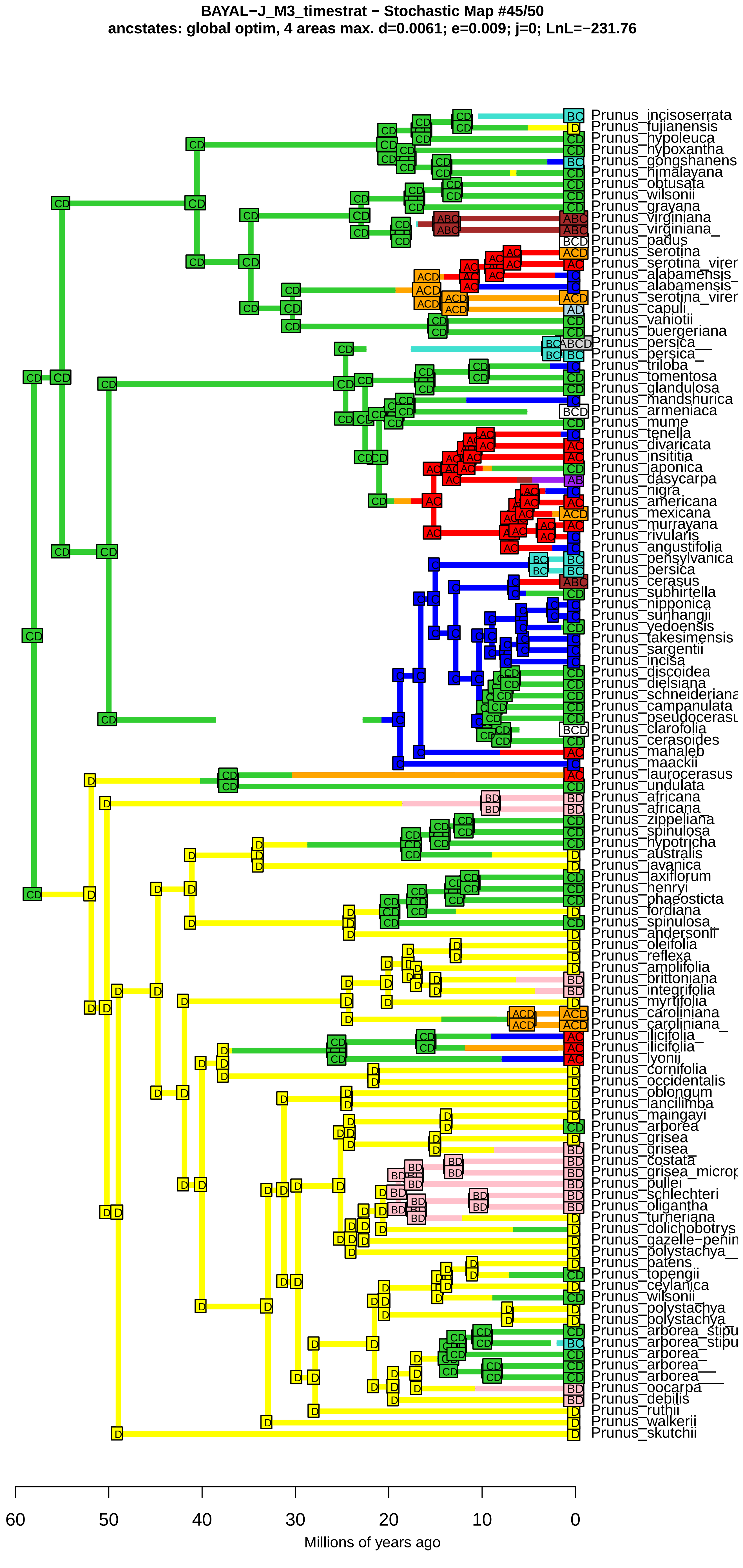
BAYAL−J_M3_timestrat − Stochastic Map #45/50
ancstates: global optim, 4 areas max. d=0.0061; e=0.009; j=0; LnL=−231.76
Prunus_incisoserrata
Prunus_fujianensis
Prunus_hypoleuca
Prunus_hypoxantha
Prunus_gongshanens
Prunus_himalayana
Prunus_obtusata
Prunus_wilsonii
Prunus_grayana
Prunus_virginiana
Prunus_virginiana_
Prunus_padus
Prunus_serotina
Prunus_serotina_virer
Prunus_alabamensis_
Prunus_alabamensis
Prunus_serotina_virer
Prunus_capuli
Prunus_vaniotii
Prunus_buergeriana
Prunus_persica__
Prunus_persica_
Prunus_triloba
Prunus_tomentosa
Prunus_glandulosa
Prunus_mandshurica
Prunus_armeniaca
Prunus_mume
Prunus_tenella
Prunus_divaricata
Prunus_insititia
Prunus_japonica
Prunus_dasycarpa
Prunus_nigra
Prunus_americana
Prunus_mexicana
Prunus_murrayana
Prunus_rivularis
Prunus_angustifolia
Prunus_pensylvanica
Prunus_persica
Prunus_cerasus
Prunus_subhirtella
Prunus_nipponica
Prunus_sunhangii
Prunus_yedoensis
Prunus_takesimensis
Prunus_sargentii
Prunus_incisa
Prunus_discoidea
Prunus_dielsiana
Prunus_schneideriana
Prunus_campanulata
Prunus_pseudocerasu
Prunus_clarofolia
Prunus_cerasoides
Prunus_mahaleb
Prunus_maackii
Prunus_laurocerasus
Prunus_undulata
Prunus_africana
Prunus_africana_
Prunus_zippeliana
Prunus_spinulosa
Prunus_hypotricha
Prunus_australis
Prunus_javanica
Prunus_laxiflorum
Prunus_henryi
Prunus_phaeosticta
Prunus_fordiana
Prunus_spinulosa_
Prunus_andersonii
Prunus_oleifolia
Prunus_reflexa
Prunus_amplifolia
Prunus_brittoniana
Prunus_integrifolia
Prunus_myrtifolia
Prunus_caroliniana
Prunus_caroliniana_
Prunus_ilicifolia_
Prunus_ilicifolia
Prunus_lyonii
Prunus_cornifolia
Prunus_occidentalis
Prunus_oblongum
Prunus_lancilimba
Prunus_maingayi
Prunus_arborea
Prunus_grisea
Prunus_grisea_
Prunus_costata
Prunus_grisea_microp
Prunus_pullei
Prunus_schlechteri
Prunus_oligantha
Prunus_turneriana
Prunus_dolichobotrys
Prunus_gazelle−penin
Prunus_polystachya__
Prunus_patens
Prunus_topengii
Prunus_ceylanica
Prunus_wilsonii_
Prunus_polystachya
Prunus_polystachya_
Prunus_arborea_stipu
Prunus_arborea_stipu
Prunus_arborea_
Prunus_arborea__
Prunus_arborea___
Prunus_oocarpa
Prunus_debilis
Prunus_ruthii
Prunus_walkerii
Prunus_skutchii
60
50
40
30
20
10
0
Millions of years ago

**BAYAL−J_M3_timestrat − Stochastic Map #46/50**
**ancstates: global optim, 4 areas max. d=0.0061; e=0.009; j=0; LnL=−231.76**

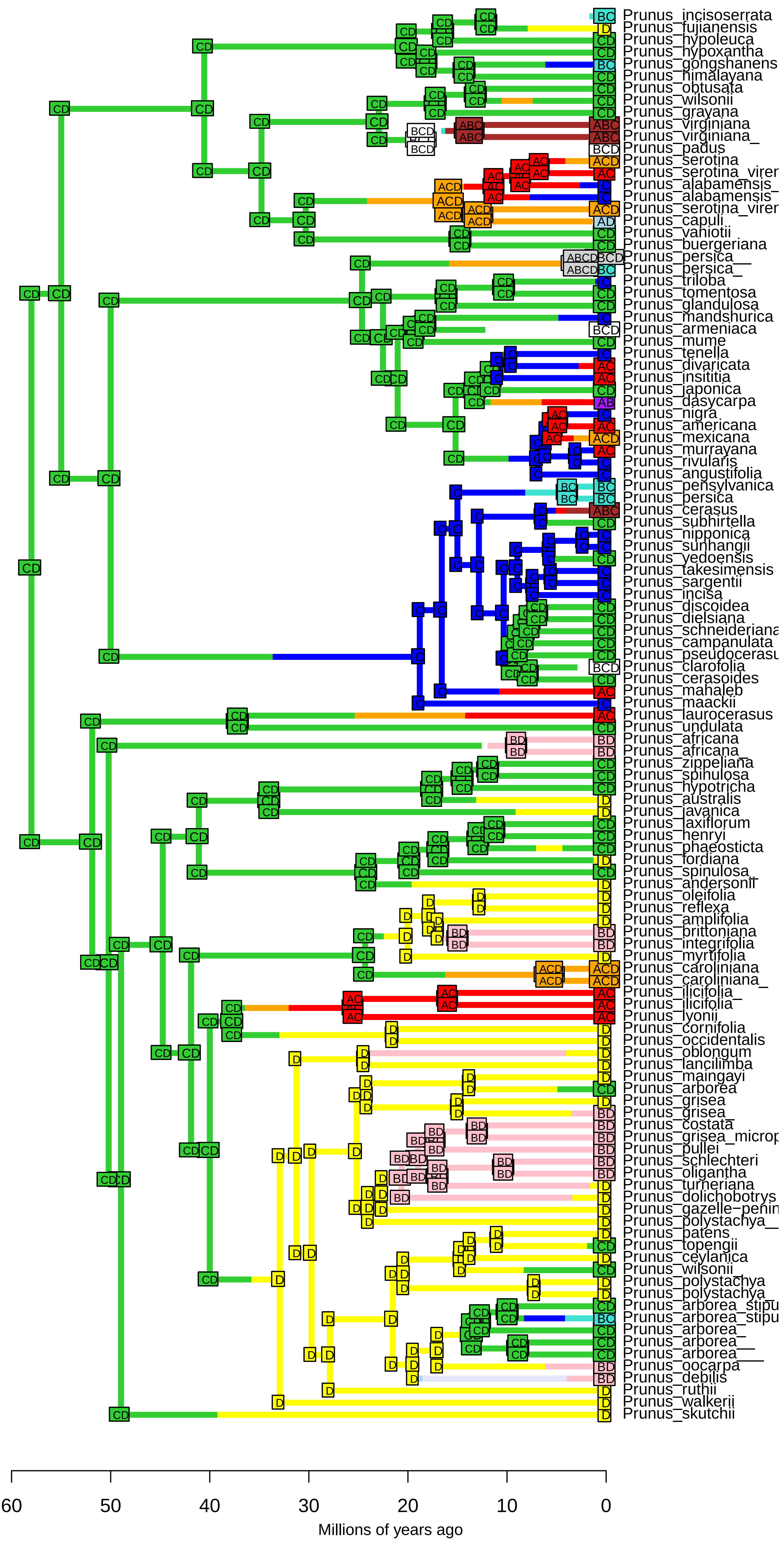

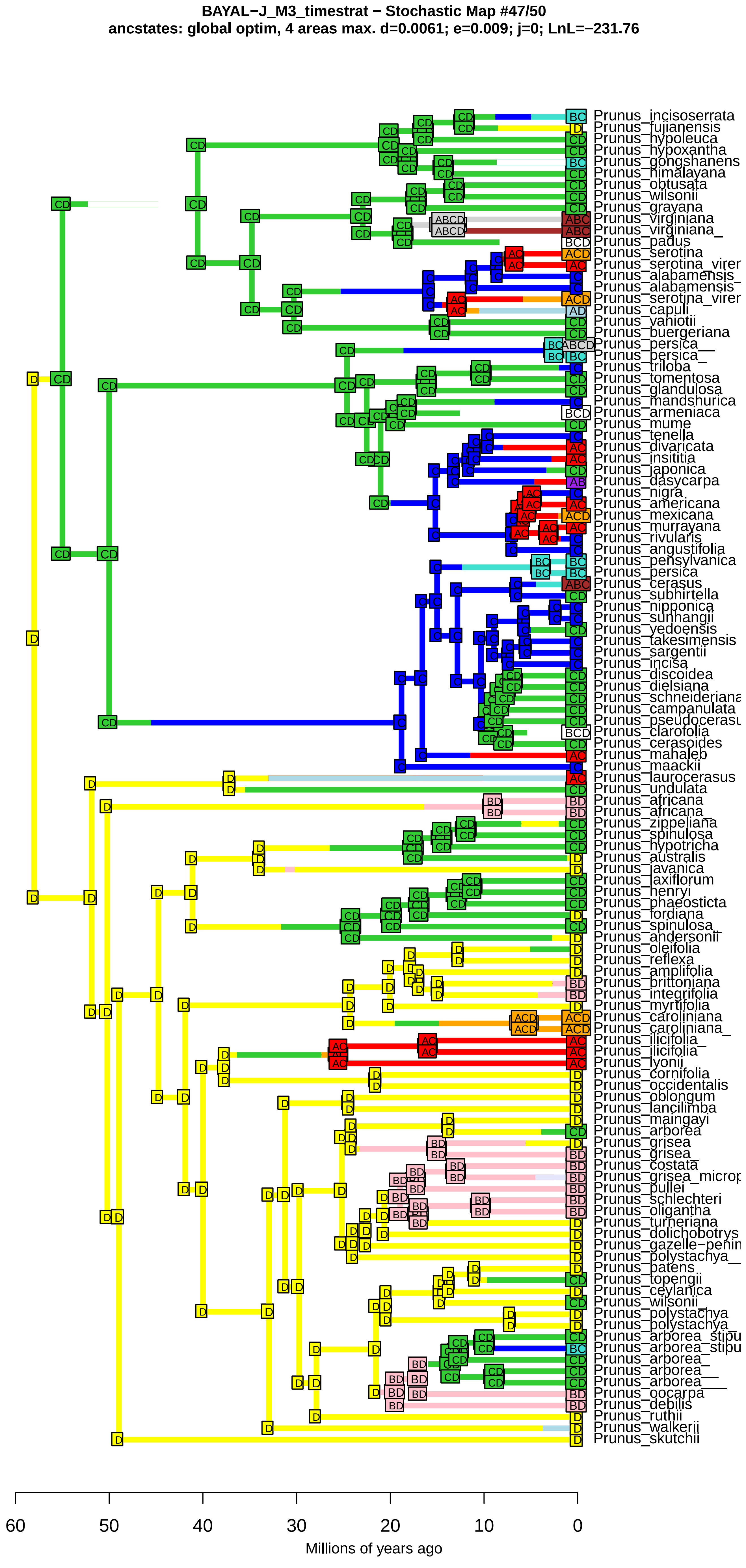
BAYAL−J_M3_timestrat − Stochastic Map #47/50
ancstates: global optim, 4 areas max. d=0.0061; e=0.009; j=0; LnL=−231.76
Prunus_incisoserrata
Prunus_fujianensis
Prunus_hypoleuca
Prunus_hypoxantha
Prunus_gongshanens
Prunus_himalayana
Prunus_obtusata
Prunus_wilsonii
Prunus_grayana
Prunus_virginiana
Prunus_virginiana_
Prunus_padus
Prunus_serotina
Prunus_serotina_virer
Prunus_alabamensis_
Prunus_alabamensis
Prunus_serotina_virer
Prunus_capuli
Prunus_vaniotii
Prunus_buergeriana
Prunus_persica__
Prunus_persica_
Prunus_triloba
Prunus_tomentosa
Prunus_glandulosa
Prunus_mandshurica
Prunus_armeniaca
Prunus_mume
Prunus_tenella
Prunus_divaricata
Prunus_insititia
Prunus_japonica
Prunus_dasycarpa
Prunus_nigra
Prunus_americana
Prunus_mexicana
Prunus_murrayana
Prunus_rivularis
Prunus_angustifolia
Prunus_pensylvanica
Prunus_persica
Prunus_cerasus
Prunus_subhirtella
Prunus_nipponica
Prunus_sunhangii
Prunus_yedoensis
Prunus_takesimensis
Prunus_sargentii
Prunus_incisa
Prunus_discoidea
Prunus_dielsiana
Prunus_schneideriana
Prunus_campanulata
Prunus_pseudocerasu
Prunus_clarofolia
Prunus_cerasoides
Prunus_mahaleb
Prunus_maackii
Prunus_laurocerasus
Prunus_undulata
Prunus_africana
Prunus_africana_
Prunus_zippeliana
Prunus_spinulosa
Prunus_hypotricha
Prunus_australis
Prunus_javanica
Prunus_laxiflorum
Prunus_henryi
Prunus_phaeosticta
Prunus_fordiana
Prunus_spinulosa_
Prunus_andersonii
Prunus_oleifolia
Prunus_reflexa
Prunus_amplifolia
Prunus_brittoniana
Prunus_integrifolia
Prunus_myrtifolia
Prunus_caroliniana
Prunus_caroliniana_
Prunus_ilicifolia_
Prunus_ilicifolia
Prunus_lyonii
Prunus_cornifolia
Prunus_occidentalis
Prunus_oblongum
Prunus_lancilimba
Prunus_maingayi
Prunus_arborea
Prunus_grisea
Prunus_grisea_
Prunus_costata
Prunus_grisea_microp
Prunus_pullei
Prunus_schlechteri
Prunus_oligantha
Prunus_turneriana
Prunus_dolichobotrys
Prunus_gazelle−penin
Prunus_polystachya__
Prunus_patens
Prunus_topengii
Prunus_ceylanica
Prunus_wilsonii_
Prunus_polystachya
Prunus_polystachya_
Prunus_arborea_stipu
Prunus_arborea_stipu
Prunus_arborea_
Prunus_arborea__
Prunus_arborea___
Prunus_oocarpa
Prunus_debilis
Prunus_ruthii
Prunus_walkerii
Prunus_skutchii
60
50
40
30
20
10
0
Millions of years ago

**BAYAL−J_M3_timestrat − Stochastic Map #48/50**
**ancstates: global optim, 4 areas max. d=0.0061; e=0.009; j=0; LnL=−231.76**

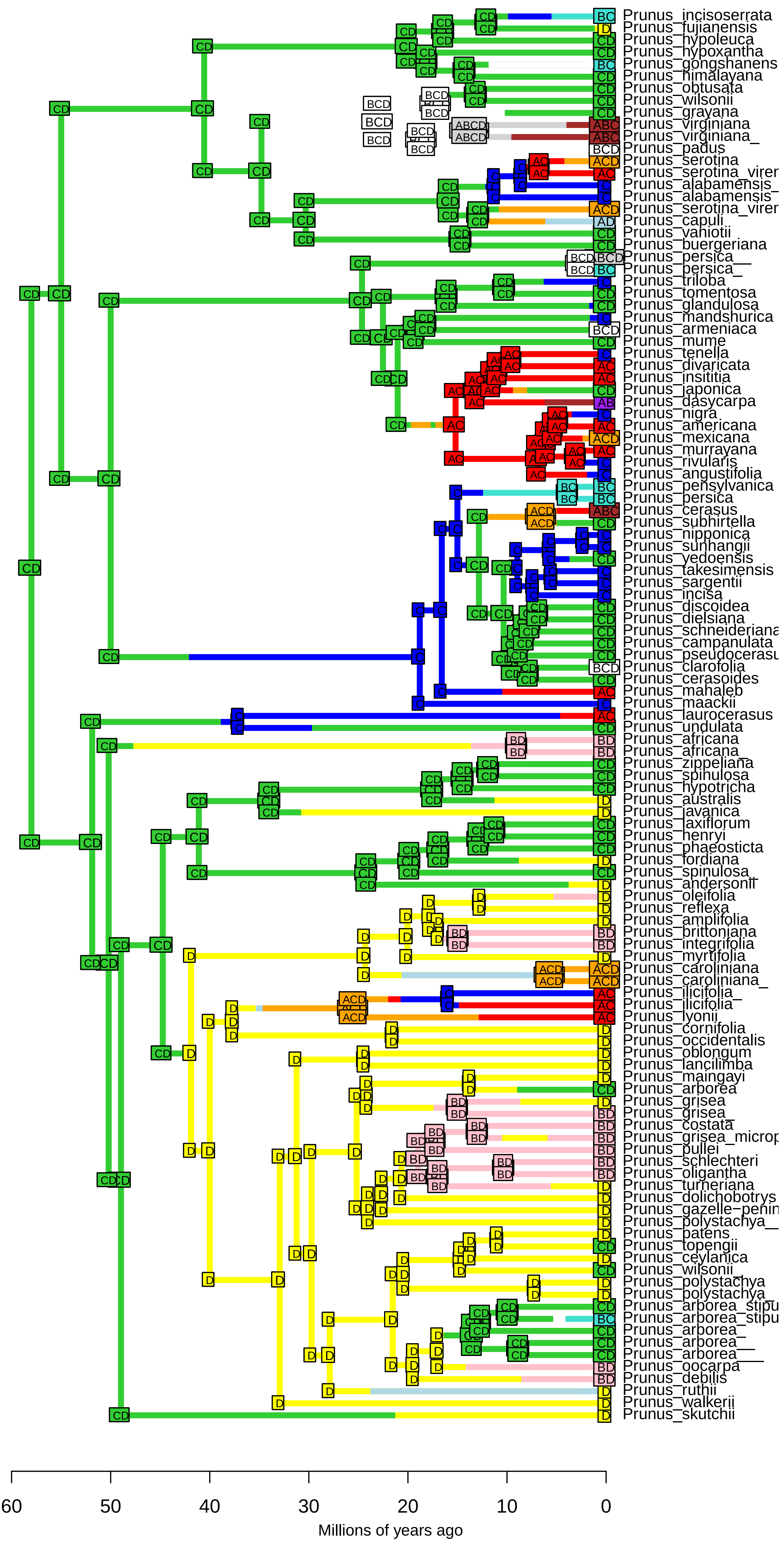

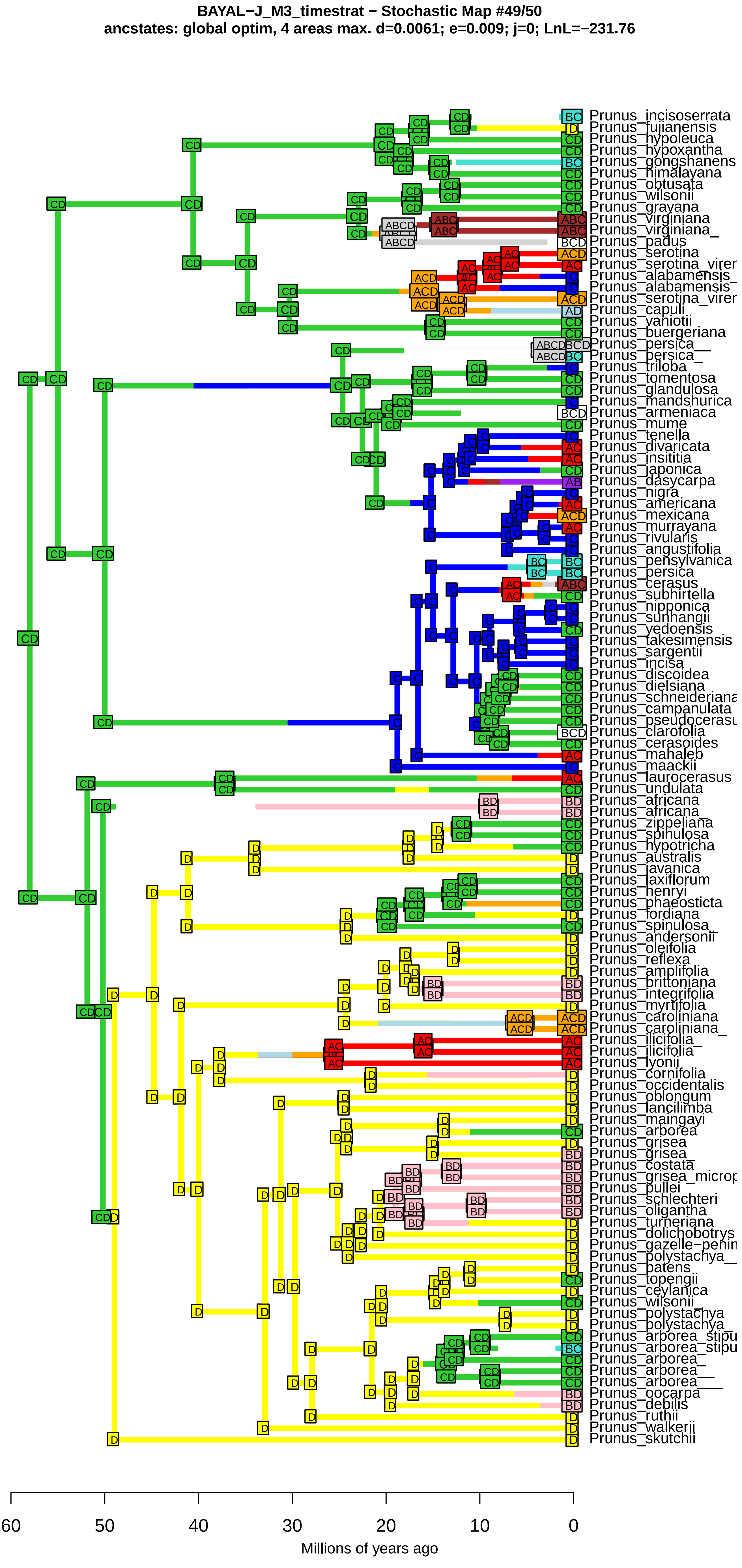
BAYAL−J_M3_timestrat − Stochastic Map #49/50
ancstates: global optim, 4 areas max. d=0.0061; e=0.009; j=0; LnL=−231.76
Prunus_incisoserrata
Prunus_fujianensis
Prunus_hypoleuca
Prunus_hypoxantha
Prunus_gongshanens
Prunus_himalayana
Prunus_obtusata
Prunus_wilsonii
Prunus_grayana
Prunus_virginiana
Prunus_virginiana_
Prunus_padus
Prunus_serotina
Prunus_serotina_virer
Prunus_alabamensis_
Prunus_alabamensis
Prunus_serotina_virer
Prunus_capuli
Prunus_vaniotii
Prunus_buergeriana
Prunus_persica__
Prunus_persica_
Prunus_triloba
Prunus_tomentosa
Prunus_glandulosa
Prunus_mandshurica
Prunus_armeniaca
Prunus_mume
Prunus_tenella
Prunus_divaricata
Prunus_insititia
Prunus_japonica
Prunus_dasycarpa
Prunus_nigra
Prunus_americana
Prunus_mexicana
Prunus_murrayana
Prunus_rivularis
Prunus_angustifolia
Prunus_pensylvanica
Prunus_persica
Prunus_cerasus
Prunus_subhirtella
Prunus_nipponica
Prunus_sunhangii
Prunus_yedoensis
Prunus_takesimensis
Prunus_sargentii
Prunus_incisa
Prunus_discoidea
Prunus_dielsiana
Prunus_schneideriana
Prunus_campanulata
Prunus_pseudocerasu
Prunus_clarofolia
Prunus_cerasoides
Prunus_mahaleb
Prunus_maackii
Prunus_laurocerasus
Prunus_undulata
Prunus_africana
Prunus_africana_
Prunus_zippeliana
Prunus_spinulosa
Prunus_hypotricha
Prunus_australis
Prunus_javanica
Prunus_laxiflorum
Prunus_henryi
Prunus_phaeosticta
Prunus_fordiana
Prunus_spinulosa_
Prunus_andersonii
Prunus_oleifolia
Prunus_reflexa
Prunus_amplifolia
Prunus_brittoniana
Prunus_integrifolia
Prunus_myrtifolia
Prunus_caroliniana
Prunus_caroliniana_
Prunus_ilicifolia_
Prunus_ilicifolia
Prunus_lyonii
Prunus_cornifolia
Prunus_occidentalis
Prunus_oblongum
Prunus_lancilimba
Prunus_maingayi
Prunus_arborea
Prunus_grisea
Prunus_grisea_
Prunus_costata
Prunus_grisea_microp
Prunus_pullei
Prunus_schlechteri
Prunus_oligantha
Prunus_turneriana
Prunus_dolichobotrys
Prunus_gazelle−penin
Prunus_polystachya__
Prunus_patens
Prunus_topengii
Prunus_ceylanica
Prunus_wilsonii_
Prunus_polystachya
Prunus_polystachya_
Prunus_arborea_stipu
Prunus_arborea_stipu
Prunus_arborea_
Prunus_arborea__
Prunus_arborea___
Prunus_oocarpa
Prunus_debilis
Prunus_ruthii
Prunus_walkerii
Prunus_skutchii
60
50
40
30
20
10
0
Millions of years ago

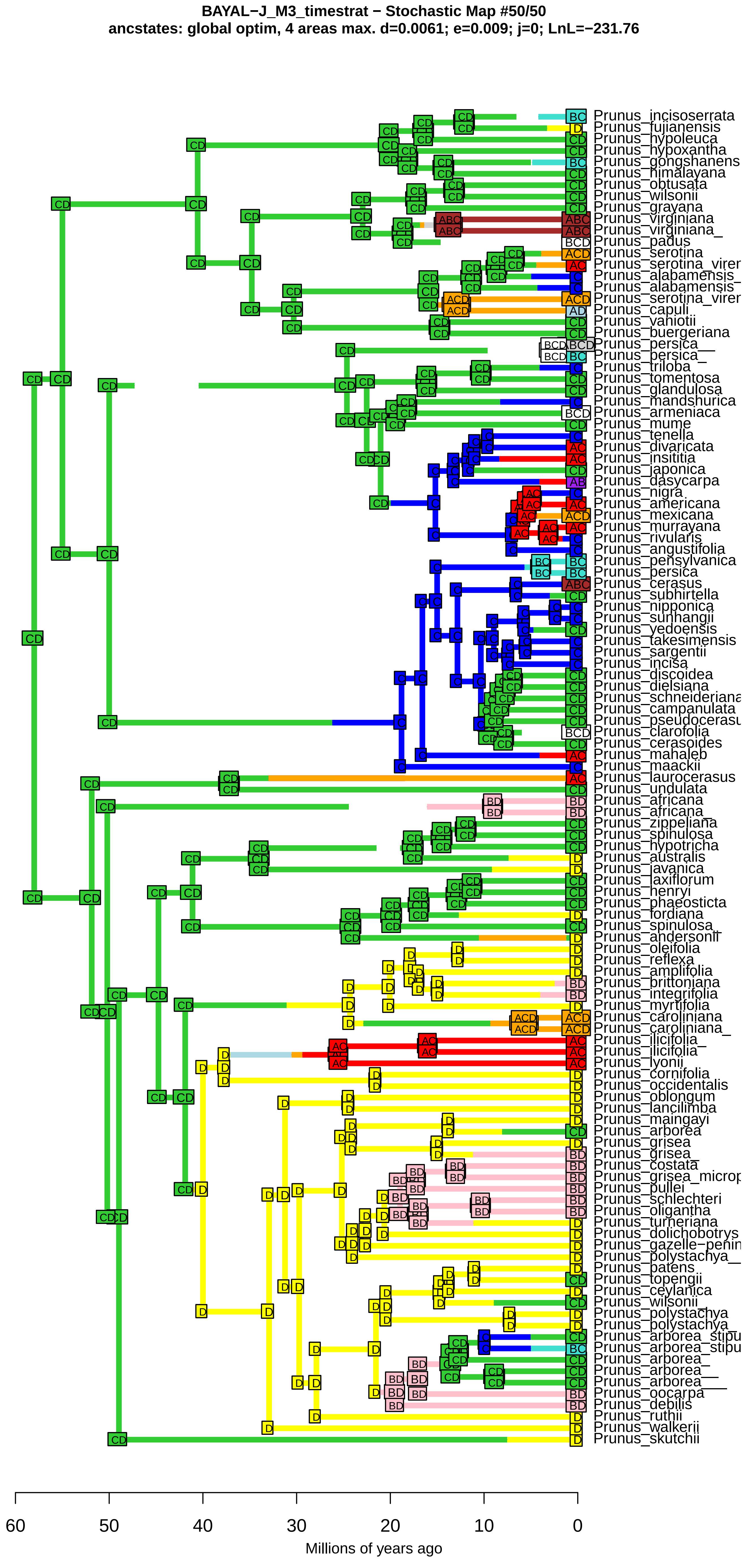
BAYAL−J_M3_timestrat − Stochastic Map #50/50
ancstates: global optim, 4 areas max. d=0.0061; e=0.009; j=0; LnL=−231.76
Prunus_incisoserrata
Prunus_fujianensis
Prunus_hypoleuca
Prunus_hypoxantha
Prunus_gongshanens
Prunus_himalayana
Prunus_obtusata
Prunus_wilsonii
Prunus_grayana
Prunus_virginiana
Prunus_virginiana_
Prunus_padus
Prunus_serotina
Prunus_serotina_virer
Prunus_alabamensis_
Prunus_alabamensis
Prunus_serotina_virer
Prunus_capuli
Prunus_vaniotii
Prunus_buergeriana
Prunus_persica__
Prunus_persica_
Prunus_triloba
Prunus_tomentosa
Prunus_glandulosa
Prunus_mandshurica
Prunus_armeniaca
Prunus_mume
Prunus_tenella
Prunus_divaricata
Prunus_insititia
Prunus_japonica
Prunus_dasycarpa
Prunus_nigra
Prunus_americana
Prunus_mexicana
Prunus_murrayana
Prunus_rivularis
Prunus_angustifolia
Prunus_pensylvanica
Prunus_persica
Prunus_cerasus
Prunus_subhirtella
Prunus_nipponica
Prunus_sunhangii
Prunus_yedoensis
Prunus_takesimensis
Prunus_sargentii
Prunus_incisa
Prunus_discoidea
Prunus_dielsiana
Prunus_schneideriana
Prunus_campanulata
Prunus_pseudocerasu
Prunus_clarofolia
Prunus_cerasoides
Prunus_mahaleb
Prunus_maackii
Prunus_laurocerasus
Prunus_undulata
Prunus_africana
Prunus_africana_
Prunus_zippeliana
Prunus_spinulosa
Prunus_hypotricha
Prunus_australis
Prunus_javanica
Prunus_laxiflorum
Prunus_henryi
Prunus_phaeosticta
Prunus_fordiana
Prunus_spinulosa_
Prunus_andersonii
Prunus_oleifolia
Prunus_reflexa
Prunus_amplifolia
Prunus_brittoniana
Prunus_integrifolia
Prunus_myrtifolia
Prunus_caroliniana
Prunus_caroliniana_
Prunus_ilicifolia_
Prunus_ilicifolia
Prunus_lyonii
Prunus_cornifolia
Prunus_occidentalis
Prunus_oblongum
Prunus_lancilimba
Prunus_maingayi
Prunus_arborea
Prunus_grisea
Prunus_grisea_
Prunus_costata
Prunus_grisea_microp
Prunus_pullei
Prunus_schlechteri
Prunus_oligantha
Prunus_turneriana
Prunus_dolichobotrys
Prunus_gazelle−penin
Prunus_polystachya__
Prunus_patens
Prunus_topengii
Prunus_ceylanica
Prunus_wilsonii_
Prunus_polystachya
Prunus_polystachya_
Prunus_arborea_stipu
Prunus_arborea_stipu
Prunus_arborea_
Prunus_arborea__
Prunus_arborea___
Prunus_oocarpa
Prunus_debilis
Prunus_ruthii
Prunus_walkerii
Prunus_skutchii
60
50
40
30
20
10
0
Millions of years ago
