## Supplemental Table S1 for "A phylogenomic approach, combined with morphological characters gleaned via machine learning, uncovers the hybrid origin and biogeographic diversification of the plum genus"

**Supplemental Table S1.** The accessions used for genomic analysis.

| **Code** | **Accession** | **Genus** | **species** |
| --- | --- | --- | --- |
| A05 | Wen_10703 | *Prunus* | *dolichobotrys* |
| A07 | Wen_11596 | *Pygeum* | *laxiflorum* |
| A09 | Wen_11240 | *Prunus* | *buergeriania* |
| A14 | Wen_11200 | *Prunus* | *campanulata* |
| A15 | Wen_11163 | *Prunus* | *pseudocerasus* |
| A16 | Wen_11471 | *Prunus* | *phaeosticta* |
| A19 | Wen_12139 | *Prunus* | *obtusata* |
| B03 | Wen_12164 | *Prunus* | *undulata* |
| B04 | Wen_12179 | *Prunus* | *grayana* |
| B05 | Wen_12158 | *Prunus* | *spinulosa* |
| B07 | Wen_11759 | *Prunus* | *serotina* |
| B10 | Wen_09061 | *Prunus* | *cerasioides* |
| B17 | Wen_12180 | *Prunus* | *clarofolia* |
| B18 | Wen_12056 | *Prunus* | *schneideriana* |
| B23 | Wen_12071 | *Prunus* | *fujianensis* |
| C14 | Wen_10150 | *Prunus* | *javanica* |
| D01 | Wen_07283 | *Prunus* | *murrayana* |
| D02 | Wen_07298 | *Prunus* | *pensylvanica* |
| D03 | Wen_07307 | *Prunus* | *institia* |
| D04 | Wen_06828 | *Prunus* | *skutchii* |
| D07 | Wen_11112 | *Prunus* | *nigra* |
| D08 | Wen_07275 | *Prunus* | *serotina* |
| D10 | Wen_08509 | *Prunus* | *spinulosa* |
| D11 | Wen_08418 | *Prunus* | *arborea stipulacea* |
| D18 | Nee&Wen_53810 | *Prunus* | *reflexa* |
| D20 | Wen_10469 | *Prunus* | *virginiana* |
| D21 | Wen_11937 | *Prunus* | *takesimensis* |
| D23 | Wen_11936 | *Prunus* | *incisa* |
| E04 | Wen_10630 | *Prunus* | *andersonii* |
| E05 | Wen_08688 | *Prunus* | *capuli* |
| E09 | Wen_08494 | *Prunus* | *dielsiana* |
| E18 | Nee&Wen_53936 | *Prunus* | *brittoniana* |
| E21 | Nee&Wen_53855 | *Prunus* | *oleifolia* |
| F07 | PPM_03489 | *Prunus* | *amplifolia* |
| F08 | Wen_09789 | *Prunus* | *triloba* |
| F13 | Wen_05813 | *Pygeum* | *patens* |
| G05 | Wen_11942 | *Prunus* | *mume* |
| G07 | Wen_10977 | *Prunus* | *fordiana* |
| G13 | Wen_9220 | *Prunus* | *vaniotii* |
| G15 | Wen_11943 | *Physocarpus* | *opulifolius* |
| G18 | Nee&Wen_53925 | *Prunus* | *cerasus* |
| G22 | Wen_6973 | *Prunus* | *cornifolia* |
| H01 | Nee&Wen_53893 | *Prunus* | *integrifolia* |
| H02 | AAH_5444-3*A | *Prunus* | *mandshurica* |
| H05 | Wen_10851 | *Prunus* | *lancilima* |
| H07 | Wen_11932 | *Prunus* | *glandulosa* |
| H09 | Wen_11988 | *Prunus* | *rivularis* |
| H10 | Wen_11836 | *Prunus* | *ruthii* |
| H13 | Wen_10807 | *Prunus* | *ceylanica* |
| H14 | AAH_665-65*A | *Maddenia* | *hypoleuea* |
| H17 | Wen_11892 | *Prunus* | *myrtifolia* |
| H18 | Wen_11946 | *Prunus* | *sargentii* |
| H21 | AAH_237-40*B | *Prunus* | *tenella* |
| H22 | AAH_1372-84*C | *Prunus* | *mahaleb* |
| H24 | Wen_11844 | *Prunus* | *occidentalis* |
| J10 | Wen_12077 | *Prunus* | *hypoxantha* |
| J12 | Wen_11501 | *Prunus* | *japonica* |
| J17 | Wen_10350 | *Prunus* | *divaricata* |
| J20 | Wen_10366 | *Prunus* | *laurocerasus* |
| K01 | Wen_11794 | *Prunus* | *maackii* |
| K10 | Wen_11585 | *Pygeum* | *henryi* |
| K12 | Wen_11284 | *Prunus* | *discoidea* |
| K14 | Wen_11796 | *Prunus* | *yedoensis* |
| K15 | Wen_11795 | *Prunus* | *subhirtella* |
| K16 | Wen_11797 | *Prunus* | *nipponica* |
| K19 | Wen_6226 | *Prunus* | *africana* |
| K22 | Wen_12336 | *Prunus* | *pullei* |
| L01 | Wen_11683 | *Prunus* | *polystachya* |
| L03 | Wen_12400 | *Prunus* | *grisea* |
| L08 | Potter_081112-06 | *Prunus* | *costata* |
| L13 | Wen_11814 | *Prunus* | *arborea* |
| L14 | Wen_11830 | *Prunus* | *maingayi* |
| L15 | Wen_11816 | *Prunus* | *arborea stipulacea* |
| L20 | Wen_11833 | *Prunus* | *patens* |
| L22 | Wen_11708 | *Prunus* | *oocarpa* |
| M02 | Wen_10743 | *Prunus* | *oligantha* |
| M12 | Wen_12318 | *Prunus* | *grisea microphylla* |
| M13 | Wen_10681 | *Prunus* | *grisea* |
| M15 | Wen_12338 | *Prunus* | *schlechteri* |
| M16 | Potter_081120-02 | *Prunus* | *gazelle-peninsulae* |
| N01 | Potter_1005 | *Prunus* | *ilicifolia* |
| N03 | Potter_1014 | *Lyonothamnus* | *floribundus* |
| N04 | Potter_1015 | *Prunus* | *lyonii* |
| PC1 | RGJH_0034 | *Prunus* | *caroliniana* |
| SA60 | RGJH_0044 | *Prunus* | *alabamensis* |
| SV1 | McVaugh_17010 | *Prunus* | *serotina virens* |
| SW16_virginiana | Wen_17122 | *Prunus* | *virginiana* |
| SW17_pensylvanica | Wen_17258 | *Prunus* | *pensylvanica* |
| SW18_pensylvanica | Wen_16988 | *Prunus* | *persica* |
| SW19_mexicana | Wen_17462 | *Prunus* | *mexicana* |
| SW20_angustifolia | Wen_17121 | *Prunus* | *angustifolia* |
| SW21_polystachya | Wen_15049 | *Prunus* | *polystachya* |
| SW22_polystachya | Wen_15018 | *Prunus* | *polystachya* |
| SW24_arborea | Wen_15061 | *Prunus* | *arborea* |
| SW25_arborea | Wen_15017 | *Prunus* | *arborea* |
| SW26_arborea | Wen_15038 | *Prunus* | *arborea* |
| SW27_padus | Wen_17117 | *Prunus* | *padus* |
| SW28_americana | Wen_17127 | *Prunus* | *americana* |
| SW30_africana | Wen_13533 | *Prunus* | *africana* |
| SW31_ilicifolia | Wen_13527 | *Prunus* | *ilicifolia* |
| SW32_caroliniana | Wen_17387 | *Prunus* | *caroliniana* |
| SA124 | RGJH_0100 | *Prunus* | *alabamensis* |
| BBL1 | BOP002749 | *Armeniaca* | *dasycarpa* |
| BBL3 | POC561974 | *Prunus* | *sunhangii* |
| BBL5 | 00589290 | *Laurocerasus* | *australis* |
| BBL6 | 01115996 | *Laurocerasus* | *hypotricha* |
| BBL8 | 01860879 | *Laurocerasus* | *zippeliana* |
| BBL9 | 01682095 | *Pygeum* | *turnerianum* |
| BBL10 | 01892431 | *Padus* | *wilsonii* |
| BBL11 | BOP002795 | *Prunus* | *persica* |
| BBL12 | 01863642 | *Pygeum* | *oblongum* |
| BBL13 | BOP228611 | *Pygeum* | *topengii* |
| BBL14 | 00995621 | *Pygeum* | *wilsonii* |
| BBL15 | 01682097 | *Pygeum* | *walkerii* |
| LZ1 | GLGS30758 | *Prunus* | *gongshanensis* |
| LZ2 | JR304 | *Prunus* | *himalayana* |
| LZ3 | PE01651988 | *Prunus* | *incisoserrata* |
| LZ4 | WX216 | *Prunus* | *tomentosa* |
| NCBI1 | SRR10322054 | *Prunus* | *armeniaca* |
