## Supplemental Table S2 for "A phylogenomic approach, combined with morphological characters gleaned via machine learning, uncovers the hybrid origin and biogeographic diversification of the plum genus"

**Supplemental Table S2.** The NCBI accession numbers for plastomes used to verify the phylogenetic position of the plastomes generated in our study.

| **Species** | **Accession number** |
| --- | --- |
| *Malus domestica* | MK434916.1 |
| *Prunus armeniaca* | NC043901.1 |
| *Prunus avium* | MK622380.1 |
| *Prunus campanulata* | NC044123.1 |
| *Prunus davidiana* | NC039735.1 |
| *Prunus dulcis* | NC034696.1 |
| *Prunus humilis* | NC035880.1 |
| *Prunus mira* | NC040125.1 |
| *Prunus mume* | NC023798.1 |
| *Prunus persica* | NC014697.1 |
| *Prunus pseudocerasus* | NC030599.1 |
| *Prunus salicina* | MH700952.1 |
| *Prunus serotina* | NC036133.1 |
| *Prunus sibirica* | KY101154.1 |
| *Prunus subhirtella* | KP760075.1 |
| *Prunus triloba* | MK790138.1 |
| *Prunus yedoensis* | NC026980.1 |
