## Supplemental Table S3 for "A phylogenomic approach, combined with morphological characters gleaned via machine learning, uncovers the hybrid origin and biogeographic diversification of the plum genus"

**Supplemental Table S3.** Selected key statistics from the HybPiper run for all individuals included in the study.

| **Name** | **Reads Mapped** | **Pct On Target** | **Genes With Seqs** | | **Genes At 25pct** | | **Genes At 50pct** | | **Genes At 75pct** | | **Paralog Warnings Long** | | **Paralog Warnings Depth** | |
| --- | --- | --- | --- | --- | --- | --- | --- | --- | --- | --- | --- | --- | --- | --- |
| Laurocerasus_australis_ BOP048778 | 1418085 | 1.4 | | 600 | | 599 | | 593 | | 585 | | 306 | | 450 |
| Laurocerasus_hypotricha_BOP228534 | 1174708 | 0.8 | | 596 | | 595 | | 592 | | 587 | | 285 | | 426 |
| Laurocerasus_zippeliana_BOP228563 | 941545 | 0.8 | | 601 | | 599 | | 596 | | 592 | | 285 | | 435 |
| Armeniaca_dasycarpa_ BOP002749 | 5084234 | 1 | | 608 | | 608 | | 607 | | 607 | | 52 | | 102 |
| Prunus_persica_ BOP002795 | 4129648 | 0.7 | | 608 | | 609 | | 608 | | 605 | | 42 | | 63 |
| Prunus_gongshanensis_ 308_XZ | 2572283 | 1.4 | | 605 | | 605 | | 600 | | 598 | | 254 | | 459 |
| Prunus_himalayana_ 304_XZ | 1922931 | 1.5 | | 605 | | 604 | | 598 | | 585 | | 77 | | 257 |
| Prunus_incisoserrata_ 901_GS | 1063216 | 0.9 | | 604 | | 603 | | 601 | | 599 | | 314 | | 488 |
| Prunus_tomentosa_216 | 2661467 | 1.3 | | 608 | | 607 | | 606 | | 603 | | 46 | | 63 |
| Prunus_armeniaca_ SRR10322054 | 5344213 | 1.7 | | 579 | | 601 | | 597 | | 577 | | 20 | | 75 |
| Pygeum_turnerianum_ BOP228613 | 1013720 | 1.8 | | 581 | | 577 | | 567 | | 554 | | 189 | | 337 |
| Padus_wilsonii_ BOP228604 | 1072553 | 1.6 | | 607 | | 606 | | 602 | | 599 | | 201 | | 380 |
| Pygeum_oblongum_ BOP228609 | 1025745 | 1.4 | | 598 | | 597 | | 591 | | 580 | | 217 | | 383 |
| Pygeum_topengii_ BOP228611 | 2298490 | 1.3 | | 601 | | 595 | | 590 | | 583 | | 357 | | 460 |
| Pygeum_wilsonii_ BOP228616 | 1441934 | 1.9 | | 598 | | 591 | | 586 | | 575 | | 289 | | 434 |
| Pygeum_walkeri_ BOP228614 | 900709 | 1.4 | | 587 | | 583 | | 571 | | 560 | | 199 | | 345 |
| Prunus_sunhangii_ POC561974 | 1652032 | 1.6 | | 606 | | 606 | | 604 | | 601 | | 48 | | 73 |
| A05_dolichobotrys | 2608722 | 42.9 | | 592 | | 590 | | 585 | | 574 | | 303 | | 427 |
| A07_laxiflorum | 7673465 | 61.2 | | 601 | | 597 | | 592 | | 584 | | 370 | | 469 |
| A09_buergeriania | 1118745 | 17.1 | | 600 | | 598 | | 595 | | 587 | | 266 | | 402 |
| A14_campanulata | 6902872 | 56.9 | | 606 | | 607 | | 606 | | 605 | | 43 | | 86 |
| A15_pseudocerasus | 5511850 | 59.8 | | 608 | | 609 | | 608 | | 608 | | 54 | | 103 |
| A16_phaeosticta | 4995480 | 49 | | 602 | | 595 | | 588 | | 574 | | 240 | | 427 |
| A19_obtusata | 4799782 | 56.4 | | 606 | | 605 | | 602 | | 594 | | 211 | | 406 |
| B03_undulata | 2307484 | 16.2 | | 606 | | 604 | | 602 | | 597 | | 310 | | 451 |
| B04_grayana | 6069413 | 63.5 | | 607 | | 606 | | 605 | | 601 | | 315 | | 498 |
| B05_spinulosa | 3671433 | 38.3 | | 606 | | 606 | | 604 | | 596 | | 293 | | 516 |
| B07_serotina | 9657532 | 63.9 | | 604 | | 602 | | 601 | | 593 | | 396 | | 516 |
| B10_cerasioides | 2025762 | 29.6 | | 607 | | 608 | | 606 | | 600 | | 34 | | 69 |
| B17_clarofolia | 6846026 | 57.7 | | 607 | | 607 | | 607 | | 604 | | 35 | | 99 |
| B18_schneideriana | 656365 | 27.2 | | 605 | | 604 | | 601 | | 596 | | 32 | | 54 |
| B23_fujianensis | 3330676 | 48.5 | | 603 | | 601 | | 596 | | 593 | | 311 | | 472 |
| C14_javanica | 1558214 | 39.6 | | 599 | | 597 | | 587 | | 580 | | 276 | | 416 |
| D01_murrayana | 4353593 | 54.8 | | 608 | | 608 | | 607 | | 604 | | 34 | | 60 |
| D02_pensylvanica | 8482165 | 64.5 | | 606 | | 604 | | 604 | | 600 | | 38 | | 63 |
| D03_institia | 6842396 | 67.7 | | 608 | | 608 | | 607 | | 604 | | 36 | | 78 |
| D04_skutchii | 9529243 | 63.7 | | 603 | | 601 | | 594 | | 585 | | 414 | | 502 |
| D07_nigra | 3351639 | 47.8 | | 606 | | 605 | | 600 | | 586 | | 18 | | 46 |
| D08_serotina | 3988642 | 48.9 | | 603 | | 601 | | 599 | | 591 | | 273 | | 418 |
| D10_spinulosa | 1363580 | 13.9 | | 601 | | 596 | | 588 | | 575 | | 261 | | 401 |
| D11_stipulacea | 4815994 | 52.7 | | 597 | | 595 | | 590 | | 581 | | 334 | | 463 |
| D18_reflexa | 6734141 | 51.2 | | 603 | | 600 | | 597 | | 588 | | 387 | | 477 |
| D20_virginiana | 5286477 | 53.6 | | 607 | | 606 | | 605 | | 602 | | 281 | | 507 |
| D21_takesimensis | 6882800 | 56.1 | | 607 | | 607 | | 604 | | 603 | | 48 | | 78 |
| D23_incisa | 2259795 | 36.6 | | 607 | | 604 | | 598 | | 571 | | 17 | | 43 |
| E04_andersonii | 6649631 | 44.5 | | 599 | | 591 | | 585 | | 559 | | 158 | | 351 |
| E05_capuli | 4600906 | 50.8 | | 602 | | 600 | | 588 | | 564 | | 136 | | 313 |
| E09_dielsiana | 7899040 | 57.4 | | 606 | | 606 | | 598 | | 567 | | 15 | | 48 |
| E18_brittoniana | 1195959 | 17.7 | | 598 | | 597 | | 590 | | 563 | | 165 | | 319 |
| E21_oleifolia | 3793863 | 40.3 | | 603 | | 602 | | 597 | | 588 | | 358 | | 465 |
| F07_amplifolia | 871547 | 18 | | 603 | | 599 | | 592 | | 575 | | 250 | | 398 |
| F08_triloba | 5531316 | 52.9 | | 609 | | 610 | | 608 | | 605 | | 34 | | 138 |
| F13_putenii | 3353165 | 34.1 | | 596 | | 592 | | 585 | | 574 | | 275 | | 419 |
| G05_mume | 4125209 | 50.2 | | 609 | | 609 | | 607 | | 603 | | 42 | | 67 |
| G07_fordiana | 4556468 | 43.7 | | 599 | | 593 | | 588 | | 567 | | 204 | | 390 |
| G13_vaniotii | 5760661 | 55.5 | | 607 | | 604 | | 601 | | 599 | | 401 | | 519 |
| G15_Physocarpus | 587739 | 20.8 | | 506 | | 494 | | 450 | | 377 | | 16 | | 20 |
| G18_cerasus | 3368166 | 28.2 | | 609 | | 609 | | 609 | | 604 | | 65 | | 168 |
| G22_cornifolia | 3228328 | 30.5 | | 600 | | 598 | | 597 | | 589 | | 374 | | 495 |
| H01_integrifolia | 1645963 | 38.1 | | 602 | | 599 | | 595 | | 577 | | 271 | | 414 |
| H02_mandshurica | 620016 | 17.4 | | 603 | | 599 | | 569 | | 501 | | 6 | | 20 |
| H05_lancilima | 825672 | 19.1 | | 593 | | 586 | | 574 | | 535 | | 92 | | 229 |
| H07_glandulosa | 11314888 | 69 | | 604 | | 602 | | 599 | | 597 | | 33 | | 52 |
| H09_rivularis | 3208636 | 30.1 | | 607 | | 604 | | 602 | | 601 | | 38 | | 60 |
| H10_ruthii | 2956050 | 21.3 | | 601 | | 598 | | 592 | | 587 | | 336 | | 455 |
| H13_ceylanica | 597017 | 18.6 | | 584 | | 584 | | 574 | | 553 | | 211 | | 367 |
| H14_hypoleuca | 2380535 | 27.9 | | 605 | | 604 | | 600 | | 593 | | 256 | | 428 |
| H17_myrtifolia | 3255932 | 27.5 | | 604 | | 602 | | 598 | | 591 | | 377 | | 479 |
| H18_sargentii | 4110822 | 47.7 | | 607 | | 605 | | 604 | | 604 | | 38 | | 68 |
| H21_tenella | 2368928 | 28.5 | | 608 | | 607 | | 604 | | 598 | | 34 | | 48 |
| H22_mahaleb | 1095941 | 26.4 | | 606 | | 607 | | 604 | | 599 | | 33 | | 49 |
| H24_occidentalis | 2411325 | 39.6 | | 601 | | 598 | | 592 | | 570 | | 177 | | 391 |
| J10_hypoxantha | 3346929 | 44.6 | | 602 | | 600 | | 596 | | 584 | | 232 | | 394 |
| J12_japonica | 5686107 | 54.7 | | 603 | | 603 | | 602 | | 595 | | 26 | | 45 |
| J17_divaricata | 2369720 | 38 | | 605 | | 605 | | 600 | | 585 | | 14 | | 24 |
| J20_laurocerasus | 5069174 | 49.4 | | 606 | | 607 | | 604 | | 579 | | 89 | | 286 |
| K01_maackii | 8906057 | 68.3 | | 603 | | 604 | | 601 | | 600 | | 41 | | 81 |
| K10_henryi | 6184553 | 54 | | 597 | | 592 | | 583 | | 566 | | 190 | | 380 |
| K12_discoidea | 4929399 | 52.2 | | 608 | | 608 | | 607 | | 605 | | 48 | | 80 |
| K14_yedoensis | 4764016 | 57.3 | | 608 | | 608 | | 607 | | 607 | | 62 | | 120 |
| K15_subhirtella | 7104861 | 64.9 | | 607 | | 608 | | 608 | | 607 | | 36 | | 55 |
| K16_nipponica | 5672093 | 62.4 | | 608 | | 608 | | 604 | | 602 | | 44 | | 62 |
| K19_africana | 5352350 | 59.6 | | 603 | | 604 | | 600 | | 592 | | 333 | | 471 |
| K22_pullei | 4921799 | 44.1 | | 593 | | 589 | | 578 | | 549 | | 163 | | 346 |
| L01_polystachya | 2401957 | 26.2 | | 598 | | 598 | | 586 | | 578 | | 311 | | 425 |
| L03_grisea | 1314243 | 22.6 | | 594 | | 592 | | 583 | | 567 | | 285 | | 407 |
| L08_costata | 5679227 | 62.1 | | 592 | | 591 | | 585 | | 574 | | 335 | | 455 |
| L13_arborea | 4761758 | 46.3 | | 592 | | 589 | | 585 | | 576 | | 293 | | 438 |
| L14_maingayi | 5569102 | 44.7 | | 589 | | 589 | | 583 | | 572 | | 330 | | 450 |
| L15_arborea | 5348687 | 50.7 | | 598 | | 596 | | 590 | | 573 | | 270 | | 433 |
| L20_debilis | 546281 | 20.6 | | 581 | | 580 | | 566 | | 536 | | 146 | | 295 |
| L22_oocarpa | 5661882 | 57.2 | | 597 | | 596 | | 588 | | 580 | | 358 | | 462 |
| M02_oligantha | 6255496 | 51.5 | | 589 | | 590 | | 581 | | 573 | | 367 | | 461 |
| M12_grisea | 1309543 | 16.4 | | 588 | | 587 | | 582 | | 565 | | 288 | | 409 |
| M13_grisea | 1539678 | 17.6 | | 591 | | 587 | | 576 | | 560 | | 241 | | 381 |
| M15_schlechteri | 8100376 | 56.8 | | 593 | | 592 | | 584 | | 572 | | 352 | | 450 |
| M16_gazelle_pen | 1267153 | 33.5 | | 586 | | 583 | | 570 | | 527 | | 147 | | 288 |
| N01_ilicifolia | 4345307 | 53 | | 603 | | 601 | | 598 | | 590 | | 250 | | 453 |
| N03_Lyonothamnus | 1913654 | 36.5 | | 547 | | 534 | | 509 | | 463 | | 236 | | 304 |
| N04_lyonii | 817452 | 20.8 | | 594 | | 593 | | 582 | | 548 | | 158 | | 305 |
| PC1 | 2786081 | 50.9 | | 600 | | 600 | | 596 | | 587 | | 400 | | 472 |
| SA60 | 1307922 | 18.3 | | 596 | | 596 | | 593 | | 584 | | 319 | | 439 |
| SV1 | 8050349 | 71.8 | | 604 | | 604 | | 601 | | 596 | | 405 | | 522 |
| SW16_virginiana | 12775444 | 38.8 | | 607 | | 608 | | 607 | | 606 | | 334 | | 528 |
| SW17_pensylvanica | 13855300 | 47.2 | | 604 | | 603 | | 603 | | 599 | | 41 | | 63 |
| SW18_pensylvanica | 5846194 | 33.8 | | 610 | | 610 | | 610 | | 607 | | 49 | | 62 |
| SW19_mexicana | 10894778 | 42.1 | | 609 | | 609 | | 608 | | 605 | | 37 | | 92 |
| SW20_angustifolia | 8728904 | 44.1 | | 606 | | 606 | | 605 | | 604 | | 43 | | 66 |
| SW21_polystachya | 7285086 | 36.4 | | 598 | | 598 | | 589 | | 582 | | 391 | | 457 |
| SW22_polystachya | 4896998 | 22.2 | | 595 | | 593 | | 588 | | 577 | | 366 | | 457 |
| SW24_arborea | 6920922 | 25 | | 595 | | 595 | | 590 | | 581 | | 392 | | 468 |
| SW25_arborea | 12062465 | 38.7 | | 598 | | 595 | | 588 | | 583 | | 373 | | 475 |
| SW26_arborea | 13163832 | 32.7 | | 595 | | 592 | | 587 | | 581 | | 359 | | 467 |
| SW27_padus | 8059287 | 32.7 | | 607 | | 607 | | 607 | | 607 | | 304 | | 539 |
| SW28_americana | 9873463 | 34.8 | | 608 | | 609 | | 607 | | 604 | | 44 | | 65 |
| SW30_africana | 5017914 | 22.5 | | 603 | | 604 | | 600 | | 589 | | 398 | | 480 |
| SW31_ilicifolia | 7222197 | 31.2 | | 606 | | 603 | | 600 | | 595 | | 408 | | 505 |
| SW32_caroliniana | 9410581 | 29.7 | | 601 | | 598 | | 597 | | 590 | | 416 | | 485 |
| SA124 | 7470133 | 52.3 | | 602 | | 602 | | 600 | | 596 | | 423 | | 525 |
