## Supplemental Table S4 for "A phylogenomic approach, combined with morphological characters gleaned via machine learning, uncovers the hybrid origin and biogeographic diversification of the plum genus"

**Supplemental Table S4.** BioGeoBears AIC and AICc table for the six biogeographic models compared when A) using geography-based biogeographic regions, B) using biome-based biogeographic regions.

**A.**

| **Model** | **LnL** | **Parameters** | **d** | **e** | **j** | **AIC** | **AIC_wt** |
| --- | --- | --- | --- | --- | --- | --- | --- |
| DEC | -189 | 2 | 0.0029 | 0.0017 | 0 | 382 | 2.8e-07 |
| DEC+J | -173.3 | 3 | 0.0016 | 5.7e-10 | 0.015 | 352.6 | 0.67 |
| DIVALIKE | -187.6 | 2 | 0.0032 | 1.0e-12 | 0 | 379.2 | 1.1e-06 |
| DIVALIKE+J | -174 | 3 | 0.0019 | 1.0e-12 | 0.014 | 354.1 | 0.31 |
| BAYAREALIKE | -220.3 | 2 | 0.0033 | 0.016 | 0 | 444.5 | 7.4e-21 |
| BAYAREALIKE+J | -177 | 3 | 0.0012 | 1.0e-07 | 0.021 | 360 | 0.017 |
| **Model** | **LnL** | **Parameters** | **d** | **e** | **j** | **AICc** | **AICc_wt** |
| DEC | -189 | 2 | 0.0029 | 0.0017 | 0 | 382.1 | 3.0e-07 |
| DEC+J | -173.3 | 3 | 0.0016 | 5.7e-10 | 0.015 | 352.8 | 0.67 |
| DIVALIKE | -187.6 | 2 | 0.0032 | 1.0e-12 | 0 | 379.3 | 1.2e-06 |
| DIVALIKE+J | -174 | 3 | 0.0019 | 1.0e-12 | 0.014 | 354.3 | 0.31 |
| BAYAREALIKE | -220.3 | 2 | 0.0033 | 0.016 | 0 | 444.6 | 7.8e-21 |
| BAYAREALIKE+J | -177 | 3 | 0.0012 | 1.0e-07 | 0.021 | 360.2 | 0.017 |

**B.**

| **Model** | **LnL** | **Parameters** | **d** | **e** | **j** | **AIC** | **AIC_wt** |
| --- | --- | --- | --- | --- | --- | --- | --- |
| DEC | -263.2 | 2 | 0.013 | 1.0e-12 | 0 | 530.3 | 1.7e-14 |
| DEC+J | -263.2 | 3 | 0.013 | 1.0e-12 | 1.0e-05 | 532.4 | 6.1e-15 |
| DIVALIKE | -270.8 | 2 | 0.014 | 1.0e-12 | 0 | 545.7 | 7.8e-18 |
| DIVALIKE+J | -270.8 | 3 | 0.014 | 1.0e-12 | 1.0e-05 | 547.7 | 2.9e-18 |
| BAYAREALIKE | -231.8 | 2 | 0.0061 | 0.0089 | 0 | 467.5 | 0.73 |
| BAYAREALIKE+J | -231.8 | 3 | 0.0061 | 0.009 | 1.0e-05 | 469.5 | 0.27 |
| **Model** | **LnL** | **Parameters** | **d** | **e** | **j** | **AICc** | **AICc_wt** |
| DEC | -263.2 | 2 | 0.013 | 1.0e-12 | 0 | 530.5 | 1.7e-14 |
| DEC+J | -263.2 | 3 | 0.013 | 1.0e-12 | 1.0e-05 | 532.6 | 5.9e-15 |
| DIVALIKE | -270.8 | 2 | 0.014 | 1.0e-12 | 0 | 545.8 | 7.9e-18 |
| DIVALIKE+J | -270.8 | 3 | 0.014 | 1.0e-12 | 1.0e-05 | 547.9 | 2.7e-18 |
| BAYAREALIKE | -231.8 | 2 | 0.0061 | 0.0089 | 0 | 467.6 | 0.74 |
| BAYAREALIKE+J | -231.8 | 3 | 0.0061 | 0.009 | 1.0e-05 | 469.7 | 0.26 |
