## Supplemental Table S5 for "A phylogenomic approach, combined with morphological characters gleaned via machine learning, uncovers the hybrid origin and biogeographic diversification of the plum genus"

**Supplemental Table S5.** BioGeoBears likelihood ratio tests to determine if the J parameter results in significantly improved likelihood scores for A) the geography-based biogeographic regions, and B) the biome-based biogeographic regions.

**A.**

| **alt** | **null** | **LnL alt** | **LnL null** | **DFalt** | **DFnull** | **DF** | **D statistic** | **pval** | **test** | **tail** |
| --- | --- | --- | --- | --- | --- | --- | --- | --- | --- | --- |
| DEC+J | DEC | -173.3 | -189 | 3 | 2 | 1 | 31.38 | 2.1e-08 | chi-squared | one-tailed |
| DIVALIKE+J | DIVALIKE | -174 | -187.6 | 3 | 2 | 1 | 27.11 | 1.9e-07 | chi-squared | one-tailed |
| BAYAREALIKE+J | BAYAREALIKE | -177 | -220.3 | 3 | 2 | 1 | 86.54 | 1.4e-20 | chi-squared | one-tailed |

| **alt** | **null** | **AIC1** | **AIC2** | **AICwt1** | **AICwt2** | **AICweight_ratio_model1** | **AICweight_ratio_model2** |
| --- | --- | --- | --- | --- | --- | --- | --- |
| DEC+J | DEC | 352.6 | 382 | 1 | 4.2e-07 | 2393122 | 4.2e-07 |
| DIVALIKE+J | DIVALIKE | 354.1 | 379.2 | 1 | 3.5e-06 | 283796 | 3.5e-06 |
| BAYAREALIKE+J | BAYAREALIKE | 360 | 444.5 | 1 | 4.4e-19 | 2.28e+18 | 4.4e-19 |

**B.**

| **alt** | **null** | **LnL alt** | **LnL null** | **DFalt** | **DFnull** | **DF** | **D statistic** | **pval** | **test** | **tail** |
| --- | --- | --- | --- | --- | --- | --- | --- | --- | --- | --- |
| DEC+J | DEC | -263.2 | -263.2 | 3 | 2 | 1 | -0.0016 | 1 | chi-squared | one-tailed |
| DIVALIKE+J | DIVALIKE | -270.8 | -270.8 | 3 | 2 | 1 | -0.0054 | 1 | chi-squared | one-tailed |
| BAYAREALIKE+J | BAYAREALIKE | -231.8 | -231.8 | 3 | 2 | 1 | -0.005 | 1 | chi-squared | one-tailed |

| **alt** | **null** | **AIC1** | **AIC2** | **AICwt1** | **AICwt2** | **AICweight_ratio_model1** | **AICweight_ratio_model2** |
| --- | --- | --- | --- | --- | --- | --- | --- |
| DEC+J | DEC | 532.4 | 530.3 | 0.27 | 0.73 | 0.37 | 2.72 |
| DIVALIKE+J | DIVALIKE | 547.7 | 545.7 | 0.27 | 0.73 | 0.37 | 2.73 |
| BAYAREALIKE+J | BAYAREALIKE | 469.5 | 467.5 | 0.27 | 0.73 | 0.37 | 2.73 |
