## Supplemental Table S6 for "A phylogenomic approach, combined with morphological characters gleaned via machine learning, uncovers the hybrid origin and biogeographic diversification of the plum genus"

**Supplemental Table S6.** BioGeoBears summary table of the 50 biogeographic stochastic mapping counts using the DEC+J model on the geography-based biogeographic regions (top), and an analogous summary for the biome-based biogeographic regions using the BAYAREALIKE model (bottom).

|  | **Founder** | **a** | **d** | **e** | **Subset** | **Vicariance** |
| --- | --- | --- | --- | --- | --- | --- |
| **Means** | 15.92 | 0 | 19.66 | 0 | 6.02 | 3.48 |
| **Std Devs** | 2.05 | 0 | 1.53 | 0 | 2.44 | 1.61 |
| **Sums** | 796 | 0 | 983 | 0 | 301 | 174 |
|  | **Sympatry** | **All dispersal** | **Ana dispersal** | **All anagenesis** | **All cladogenesis** | **Total events** |
| **Means** | 90.58 | 35.58 | 19.66 | 19.66 | 116 | 135.7 |
| **Std Devs** | 1.91 | 1.51 | 1.53 | 1.53 | 0 | 1.53 |
| **Sums** | 4529 | 1779 | 983 | 983 | 5800 | 6783 |
|  | **Founder** | **a** | **d** | **e** | **Subset** | **Vicariance** |
| **Means** | 0 | 0 | 48.56 | 0 | 0 | 0 |
| **Std Devs** | 0 | 0 | 4.07 | 0 | 0 | 0 |
| **Sums** | 0 | 0 | 2428 | 0 | 0 | 0 |
|  | **Sympatry** | **All dispersal** | **Ana dispersal** | **All anagenesis** | **All cladogenesis** | **Total events** |
| **Means** | 116 | 48.56 | 48.56 | 48.56 | 116 | 164.6 |
| **Std Devs** | 0 | 4.07 | 4.07 | 4.07 | 0 | 4.07 |
| **Sums** | 5800 | 2428 | 2428 | 2428 | 5800 | 8228 |
