## Supplemental Table S7 for "A phylogenomic approach, combined with morphological characters gleaned via machine learning, uncovers the hybrid origin and biogeographic diversification of the plum genus"

**Supplemental Table S7.** BioGeoBears dispersal table showing the average number, and directionality, of all dispersal events for *Prunus* using 50 biogeographic stochastic mapping iterations for the geography-based biogeographic regions (top), and biome-based biogeographic regions (bottom). For each table, the Sink region is denoted by the row names, and the Source region is indicated by the column name.

|  |  | **Source** | | | | | | |
| --- | --- | --- | --- | --- | --- | --- | --- | --- |
|  |  | **N. America** | **Europe** | **W. Asia** | **E. Asia** | **Oceania** | **S. America** | **Africa** |
| **Sink** | **N. America** | 0 | 0.36 | 0.4 | 3.82 | 0.06 | 1.28 | 0.26 |
|  | **Europe** | 0.18 | 0 | 3.08 | 2.76 | 0.06 | 0 | 0.02 |
|  | **W. Asia** | 0.4 | 4.08 | 0 | 3 | 0.12 | 0 | 0.02 |
|  | **E. Asia** | 0.52 | 0.22 | 0.16 | 0 | 2.68 | 0.04 | 0.04 |
|  | **Oceania** | 1 | 0.02 | 0.02 | 5.12 | 0 | 0 | 0 |
|  | **S. America** | 3.96 | 0 | 0 | 0.14 | 1 | 0 | 0.04 |
|  | **Africa** | 0.34 | 0.02 | 0 | 0.28 | 0.08 | 0 | 0 |
|  |  | **Source** | | | |  |  |  |
|  |  | **Dry** | **Cold** | **Temperate** | **Tropical** |  |  |  |
| **Sink** | **Dry** | 0 | 0.95 | 2.94 | 1.55 |  |  |  |
|  | **Cold** | 0.87 | 0 | 1.8 | 1.76 |  |  |  |
|  | **Temperate** | 0.92 | 0.33 | 0 | 2.48 |  |  |  |
|  | **Tropical** | 1.12 | 0.64 | 1.54 | 0 |  |  |  |
